# The tissue-specific autophagic response to nutrient deprivation

**DOI:** 10.1101/2022.11.12.516287

**Authors:** Young Joo Yang, Hilary Grosso Jasutkar, Christopher J. Griffey, Kiryung Kim, Thomas James Melia, Noah Dephoure, Ai Yamamoto

## Abstract

Macroautophagy is a highly adaptable degradative system that is essential for life. Although studies have shown the importance of this pathway across all organ systems, we have little understanding of how discrete tissues might employ autophagy and how this changes during stress. Using an approach to identify quantitatively autophagic cargoes, we sought to identify how cells from the adult liver and brain rely on autophagy under basal conditions and during nutrient deprivation. We find that in addition to the turnover of cell type specific proteins, the different organs relied on autophagy differentially for the turnover of organelles such as mitochondria, peroxisomes and ER. Moreover, in response to nutrient deprivation, although both tissues showed increased cargo capture, cell type- and tissue-specific patterns emerged. Most notably in the brain, we found an increased representation of glial and endothelial cell cargoes, whereas neuronal cargoes were relatively unchanged. In liver, we unexpectedly found a decreased representation of mitochondrial proteins, which represented a shift moving away from the whole mitochondrion turnover to piecemeal. These results indicate how the physiologic context of the different cell types significantly influence autophagy-dependence, and begins to shed insight into how the term ‘autophagy dysfunction’ might be thought of when considering different disease states.

## Introduction

Autophagy is a general term referring to the process of degrading cytoplasmic substrates via the lysosome(Klionsky, 2020). The most conserved and most frequently studied form of autophagy is macroautophagy, which involves the sequestration of cytoplasmic substrates into a transient organelle called the autophagosome. The autophagosome then transports and delivers its cargo into the lysosomal lumen by fusing with the endolysosomal system, permitting cargo-degradation by resident lysosomal hydrolases.

Macroautophagy (herein referred to as autophagy) dysfunction has been implicated in diseases across organ systems, and its outright loss of function is incompatible with life (Mizushima and Levine, 2010). Nonetheless, we understand little about exactly how these different organ systems rely on autophagy and how this reliance might change during stress. In this study, we aimed to understand how cells of an adult organism use autophagy under fully fed and nutrient deprived conditions. Using a quantitative proteomic approach to catalog the cargoes captured by cells of the adult brain and liver, we find that cells of these highly related organs rely on macroautophagy differently, by demonstrating a differential turnover of organelles.

Moreover, in response to nutrient deprivation, we show that the brain indeed mounts and autophagic response, predominantly through the non-neuronal cells, and that in liver, the cells switch from wholesale to piecemeal turnover of mitochondria. In sum, this study illustrates that physiologic context is an important determinant of how cells rely on autophagy, and illustrates the complexity that the seemingly simple concept ‘autophagy dysfunction’ actually implies.

## Results and discussion

Although different approaches have been developed to isolate autophagosomes (Dengjel et al., 2012; Fellinger and Rez, 1990; Furuno et al., 1982; Gao et al., 2010; Le Guerroue et al., 2017; Marzella et al., 1982; Yao et al., 2016)many rely on immunoisolation of LC3 from total membranes. Given that LC3 can be found on different membranes in the cell (Hailey et al., 2010; Heckmann and Green, 2019; Korkhov, 2009; Martinez et al., 2015), immunoisolation from total membranes would not suffice for whole tissue. We therefore returned to an adaptation of a multistep fractionation approach to significantly enrich for autophagic vacuoles (AVs) (Stromhaug et al., 1998) that is combined with an immunoisolation approach to further purify for LC3-positive structures within this fraction (Aber et al., 2022; Jeong et al., 2009) (Figure 1A, S1).

**Figure 1.**
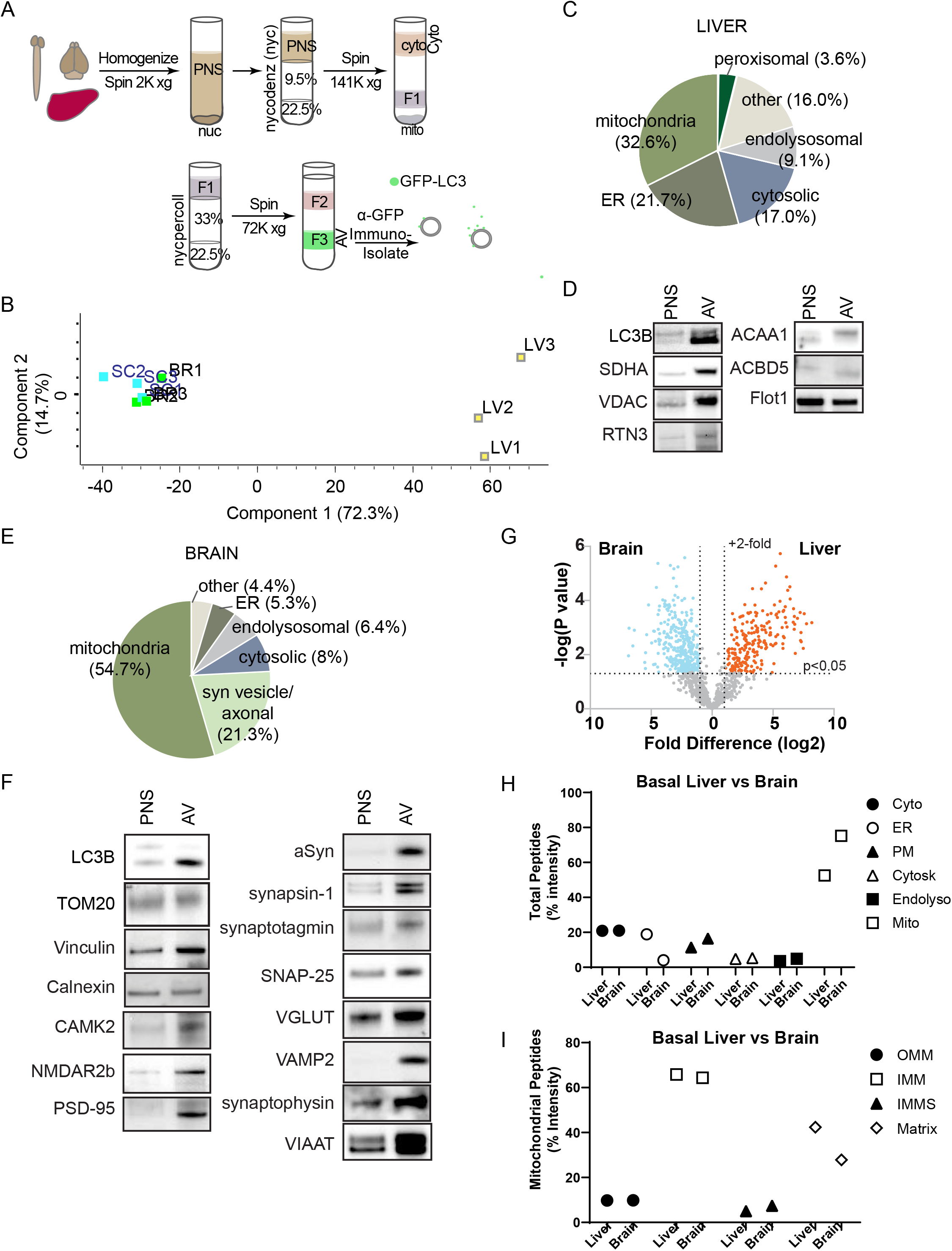
Isolation and characterization of the adult liver and brain AV proteomes. **A**. Schematic of the autophagic vesicle preparation (AV prep). 5 to 6 Spinal cord, brain and liver from 6m/o GFP-LC3 mice are collected, homogenized and fractionated for each AV enrichment, then immunoisolated with anti-GFP magnetic beads. **B**. Principal component analyses (PCA) comparing the AV preps. **C-F**. Pie-chart depiction of subcellular distribution and western blot validation for the presence of a subset of proteins of adult liver (C, D) or brain (E, F) AV proteomes. C, D. Of the 550 proteins identified, 179 were mitochondrial (e.g. SDHA, VDAC), 119 were ER proteins (RTN), 50 were endolysosomal (Flot1) and 20 were peroxisomal proteins (ACAA1, ACBD5). D. The immunoisolated AV fraction is indicated by the presence of LC3B. D. Of the 550 proteins identified, 301 were mitochondrial, followed by 117 synaptic vesicle/axonal proteins. E. Validation of proteomic analyses. In AV, endogenous LC3-II is enriched compared to PNS. TOM20 is a mitochondrial protein, vinculin is cytosolic and calnexin is an ER protein. Both pre (alpha-synuclein, synapsin-1, synaptotagmin, SNAP-25, VGLUT, VAMP2, synaptophysin, VIAAT) and postsynaptic proteins (NMDAR2b, PSD-95) are clearly present in the AV fraction. Representative western blots from n=3 to 4 AV preparations shown. **G**. Volcano plots comparing Liver vs. Brain Proteins greater than 2 fold difference, with a p < 0.05 are considered significant differences and are shown in orange (higher in liver) or blue (higher in brain). Proteins that do not differ (gray) are proteins that are common to the AVs generated across different tissue. **H, I**. A comparison of the two proteomes using peptide intensity and GO analysis-based categorization. H. Quantitative comparison of liver and brain AV proteome based on the intensity of peptides from different intracellular organelles relative to total peptide intensity between liver and brain. In relation to Table S4. Comparison of the cytosolic (Cyto), cytoskeletal (Cytosk), plasmalemmal (PM) and endolysosomal (Endolyso) compartments suggest little difference between the AV proteomes. In contrast, AVs from liver contain proportionately more ER proteins, and AVs from brain contain proportionately more mitochondrial (Mito) proteins. I. Comparison of the outer mitochondrial membrane (OMM), inner mitochondrial membrane (IMM), intermembrane mitochondrial space (IMMS), and matrix proteins. The mitochondrial subcompartment signals were normalized for the total mitochondrial intensity of respective tissue (% Intensity). The proportion of the mitochondrial peptides represented is similar across tissue, suggesting that the increased signal of mitochondrial peptides in C (Brain) is due to an overall greater number of mitochondrial peptides in brain AVs. The classification of peptides in C and D are based on curated lists originally generated from analyses by gene ontology.

For this tissue-based approach, we took advantage of GFP-LC3 mice, which transgenically expresses LC3 broadly, and immunoisolated using anti-GFP magnetic beads (Figure 1A, S1). To quality control our isolation approach, we first analyzed for the presence of both GFP-LC3 and endogenous LC3 across the different fractions from liver with an anti-LC3 antibody (Figure S1A). GFP-LC3 and LC3 enrich in fraction F3, the predicted fraction for autophagic vacuoles (AV)(Stromhaug et al., 1998). Notably, there is also a significant presence in fraction F2, a fraction considered to be enriched in ER microsomes (Strømhaug et al., 1998).

To examine further the difference between fractions F2 and F3, both fractions were individually subject to immunoisolation for GFP-LC3, and probed (Figure S1B). Immunoblotting revealed that immunoisolation enriches for GFP-LC3 as expected, but also enriches for endogenous LC3-II in both fractions. These data suggest that GFP-LC3 and endogenous LC3 co-exist on the same vesicles and do not segregate. Immunoisolation from F3 also leads to the enrichment for autophagy cargo adaptor proteins such as p62, NBR1 and early autophagosome proteins such as Atg16L, WIPI1, Atg9, suggesting that this fraction consists of nascent and mature autophagosomes, although how there is a continued persistence of Atg9 in the presence of an LC3-II enriched fraction is unclear. Atg9 may be inside the autophagosome or could be on the autophagosomal membrane. The lysosomal marker Lamp1 is lost after immunoisolation, indicating that immunoisolation separates the autophagosomes away from lysosomes that are likely persistent in the crude fractions (Strømhaug et al., 1998). In contrast, F2 does not associate with p62 and has a diminished association with WIPI1 and Atg9. Probing for NBR1 and Lamp1 also revealed their absence from F2. These data suggest that a different population of LC3-II positive structures are present in F2, possibly earlier structures, such as the isolation membrane or other structures early in autophagosome biogenesis.

We next confirmed that immunoisolation further enriched for AVs. We performed label-free proteomic analyses and subsequent quantification of fraction F3 (crude) versus an AV fraction after immunoisolation (AV) (Figure S1C,D). Immunoisolated AVs represent a small subset of the crude fraction, reducing the total number of proteins present to about half, ranging from 600 to 1000 proteins. In addition to LAMP1, proteomic analyses indicated that many of the proteins that were no longer present upon immunoisolation were lysosomal proteins such as Npc1, Galns, Gns, Prcp and Pld3 (Table S1, S2 for the complete list). We also performed cryo-EM to determine whether the LC3-positive structures are multilamellar. Crude fraction F3 incubated in the presence and absence of the anti-GFP magnetic beads used for immunoisolation were imaged and assessed for the types of vesicles that were present (Figure S1E, F).

Quantification of all of the vesicles present in the crude fraction revealed that the vast majority of structures in fraction F3 were unilamellar, and did not demonstrate the typical multilamellar structure of AVs. When quantified for structures positive for anti-GFP however, the numbers reversed-the vast majority of multilamellar structures were GFP, and therefore GFP-LC3 positive, whereas only a minority of the unilamellar structures showed positivity. These data further suggest enrichment of AVs by immunoisolation.

Using this approach, we next determined the proteome of AVs isolated from different adult (6 month-old (m/o)) tissues. Three independent AV preparations of liver, brain and spinal cord were isolated and analyzed by TMT Mass Spectrometry (TMT-MS) (Figure S1G). Pearson correlation coefficients indicated a relatively high correlation across sample preparations within the same tissue, especially for brain and spinal cord (Liver, r > 0.75; Brain, r > 0.968; Spinal Cord, r > 0.941) (Figure S1G).

In addition to the strong within tissue correlation, across tissue comparisons indicated that liver AV cargoes differ starkly from those of brain and spinal cord (Lv *vs*. Br, r = 0.49-0.57; Lv *vs*. SC, r = 0.397-0.546), whereas the brain and spinal cord were more similar, showing r-values that were similar to within tissue comparisons (Br *vs*. SC, r = 0.9-0.971). Principal Component Analyses (PCA) echoed this (Figure 1b), indicating that 72.3% of the variance segregated the liver samples, whereas brain and spinal cord samples were virtually indistinguishable by either component. Given the similarity of brain and spinal cord AV proteomes, as well as the limited number of samples generated that hindered validation, we continued our studies only on the liver and the brain.

Although liver specific proteins, such as bile acyl-CoA synthetase (SLC27A5), liver-type aldolase B (ALDOB) and cytochrome P450 enzymes were identified in the AV proteome from liver (Table S3), the majority of proteins identified more generally to ubiquitous organelles such as mitochondria, ER, endolysosomal proteins and peroxisomes (Figure 1C,D). Notably, the ER-phagy adaptor protein RTN3 is present, suggesting that the turnover of ER during basal conditions may be via selective autophagy. The clear presence of adaptor proteins such as p62, OPTN and NBR1 in AV (not shown) indicate that selective autophagy is ongoing during these times. These data indicate that under basal conditions, cells of the liver rely on autophagy to maintain organelles, especially of mitochondria and ER, whose proteins in sum make up greater than half of the proteome. These observations are suggested by early autophagy loss-of-function studies by Komatsu and colleagues, who found that the loss of autophagy in adult hepatocytes leads to hepatomegaly and the intracellular accumulation of membranes and abnormal mitochondria (Komatsu et al., 2005). Given that the inhibition of autophagy in liver can lead to liver cancer (Takamura et al., 2011), these data suggest that basal autophagy in hepatocytes maintain a healthy liver by ensuring that the functional pool of organelles, especially mitochondria and ER, are maintained.

In contrast to the liver AV proteome, in brain samples, there was a clear indication that tissue-specific proteins heavily relied on autophagy, especially synaptic vesicle and axonal proteins (Figure 1E,F). Second only to mitochondrial proteins, both pre- and post-synaptic proteins were identified in the proteome, making up greater than 20% of the total proteome. Given that neurons make up less than half of the cell population of the brain, the preponderance of these proteins in the proteome strongly suggest that neurons heavily rely on autophagy to maintain the synapse (Griffey and Yamamoto, 2022; Grosso Jasutkar and Yamamoto, 2021). Presynaptic proteins predominated, consistent with studies from indicating that autophagosomes can start at the axon tip (Hollenbeck, 1993; Maday and Holzbaur, 2012). Although the presence of axonal autophagosomes have been identified by Novikoff and his early nerve crush studies (Holtzman and Novikoff, 1965), and more recently by Yue (Komatsu et al., 2007; Yue et al., 2002), these data suggest that autophagy plays an important role in axonal maintenance and not only as a protective response in the presence of an acute or prolonged stress.

In contrast to several published reports suggesting a role for autophagy in the degradation of postsynaptic proteins (Nikoletopoulou et al., 2017; Tang et al., 2014; Yan et al., 2018) such as PSD-95 is degraded by autophagy, we did not observe any uniquely postsynaptic proteins in our proteome. One possible candidate is CamKII_α_ (Figure 1F), which plays an active role in dendrites as well as axons. We therefore probed our sample for two postsynaptic proteins, NMDA2R and PSD-95, and readily found their presence (Figure1F). These studies suggest that although there is a very strong prevalence of presynaptic proteins in our proteome, autophagy-mediated degradation of postsynaptic proteins is also occurring, although possibly at a lesser extent. Although there was a strong presence of neuronal proteins in the proteome, we also found indications that glial proteins, especially oligodendrocytes and to a lesser extent, astrocytes, are also degraded in a cell type specific manner by autophagy in the basal brain. In oligodendrocytes, the proteins identified were primarily the myelin sheath proteins, consistent with our study indicating that oligodenodroglial autophagy is important for maintaining the myelin sheath (Aber et al., 2022).

Mitochondrial proteins were by far the most prevalent cargo in the brain AV proteome, making up greater than half. Unlike synaptic vesicle proteins however, the turnover of mitochondria may be a shared feature of cell types found in the brain. Nonetheless, these data strongly suggest that the maintenance of mitochondria represents an important role for autophagy in the adult brain. Although gene knockout studies indicated that the loss of autophagy throughout development leads to the accumulation of abnormal mitochondria in the brain (Liang et al., 2010), our data indicate mitochondrial turnover plays a lasting role well into adulthood. Given how autophagy dysfunction and mitochondrial dysfunction are both implicated in adult onset neurodegenerative disease, a defect in autophagy can diminish mitochondrial turnover, and together with the loss of axonal maintenance, can lead to the degenerative features present across these diseases.

We next generated volcano plots to directly compare the liver AV proteome to the brain AV proteome (Figure 1G). Although there were clear similarities between the two proteomes when looked at individually, the volcano plots revealed that the proteomes were quite distinct. The largest differences between the liver and brain can be reflected in the tissue-specific proteins, such as synaptic vesicle proteins in the brain AV and the P450 enzymes in the liver. Another difference noted in the tissue-specific analyses was how each tissue relied on autophagy differently for the turnover of organelles. To examine this more directly, proteins identified from our proteome were matched to reference organelle groups based on GO cellular component categorization, which is powered by the PANTHER classification system (Mi et al., 2013). Although a valuable resource, the caveat of computational and pattern-matching nature of the PANTHER database is that the predicted subcellular categorizations are not always correct. Therefore, the cellular component reference matching was then manually sorted to correct for such inaccurate categorizations. The peptide intensity of proteins for each curated reference group was then summed and divided by the total intensity detected to generate the % Intensity of intracellular compartments of respective tissue (Figure 1H,Table S3). These values indicate that hepatocytes depend more on autophagy for ER turnover, whereas cells of the brain depend more for mitochondrial turnover.

To gain further insight into why the liver and brain relied on autophagy differently for the turnover of mitochondria, we next used a similar approach to determine how the different regions of the mitochondria might be represented within the AV proteomes. The peptide intensities of outer mitochondrial membrane proteins (OMM), inner mitochondrial membrane proteins (IMM), proteins found in the inter membrane mitochondrial space (IMMS) and mitochondrial matrix proteins were normalized to the total mitochondrial signal of liver or brain (Figure 1I, Table S4).

Overall, the composition of mitochondrial proteins in the AV proteomes were similar across tissue, i.e. the relative proportion of peptides that represent the OMM, IMM, IMMS and matrix were similar. This suggests that the larger portion of mitochondrial proteins in brain AVs compared to liver AVs is not due to a representation of different population of mitochondrial proteins, but rather due to a greater reliance on complete mitochondrial turnover by autophagy in the brain.

Comparative analyses of AV proteomes across different tissues revealed that under basal conditions, liver and brain rely upon autophagy differently to maintain homeostasis. We next determined if we can use the same approach to determine how a physiological stressor might change the AV proteome, and whether those changes occur in a tissue-dependent manner. To do so, 6 m/o mice were subject to a 24hr food restriction with water provided *ad libitum*, after which AVs were collected from liver and brain, then analyzed as described above. Samples from starved and basal conditions per tissue were run via TMT-MS and analyzed (Figure S2, 2, 3, and Table S4 and S5).

In contrast to the difference observed with the comparison in Figure 1, the datasets generated between basal and starved conditions within tissue lead to changes, albeit much smaller ones than between tissue. For example, the comparison between liver and brain, which showed little to no correlation, had coefficients that ranged from (r=0.49-0.57) (Figure S1), whereas between Starve and Basal states within the liver, the correlation coefficients ranged from 0.689 to 0.943 (Figure S2A). Despite the small apparent difference, PCA still found that the 12.5% of the variability across samples was driven by treatment (Figure S2B). These data suggest that in the liver, 24 hours starvation leads to clear and detectable changes in the AV proteome.

Notably, volcano plot analyses comparing the Starved to Basal AV proteomes in liver indicate that although there is increased representation of cargoes upon starvation, most cargoes remain largely unchanged, and markedly, there are also cargoes that significantly decreased (Figure 2A). Initial examination of which proteins decreased in representation during starvation revealed that 15/17 were mitochondrial proteins. Using peptide intensity coupled with a curated GO analysis indicated that although the representation of most organelles and cytosol (including ER) was increased, mitochondria was markedly decreased in response to starvation (Figure 2B). This was unexpected given the well-accepted role of autophagy and the turnover of mitochondria during stress. Further analyses examining which mitochondrial proteins were affected by starvation yielded another unexpected outcome: Rather than a decreased representation of all mitochondrial compartments, starvation led to a reduction specifically of IMM and matrix proteins (Figure 2C). Western blot analyses also confirmed these findings (Figure 2D, E). The fraction of the outer mitochondrial membrane (OMM) protein TOM20 represented in the AVs is similar between basal and starved conditions, whereas, IMM and mitochondrial matrix proteins are clearly decreased upon starvation, supporting the proteomic analyses.

**Figure 2.**
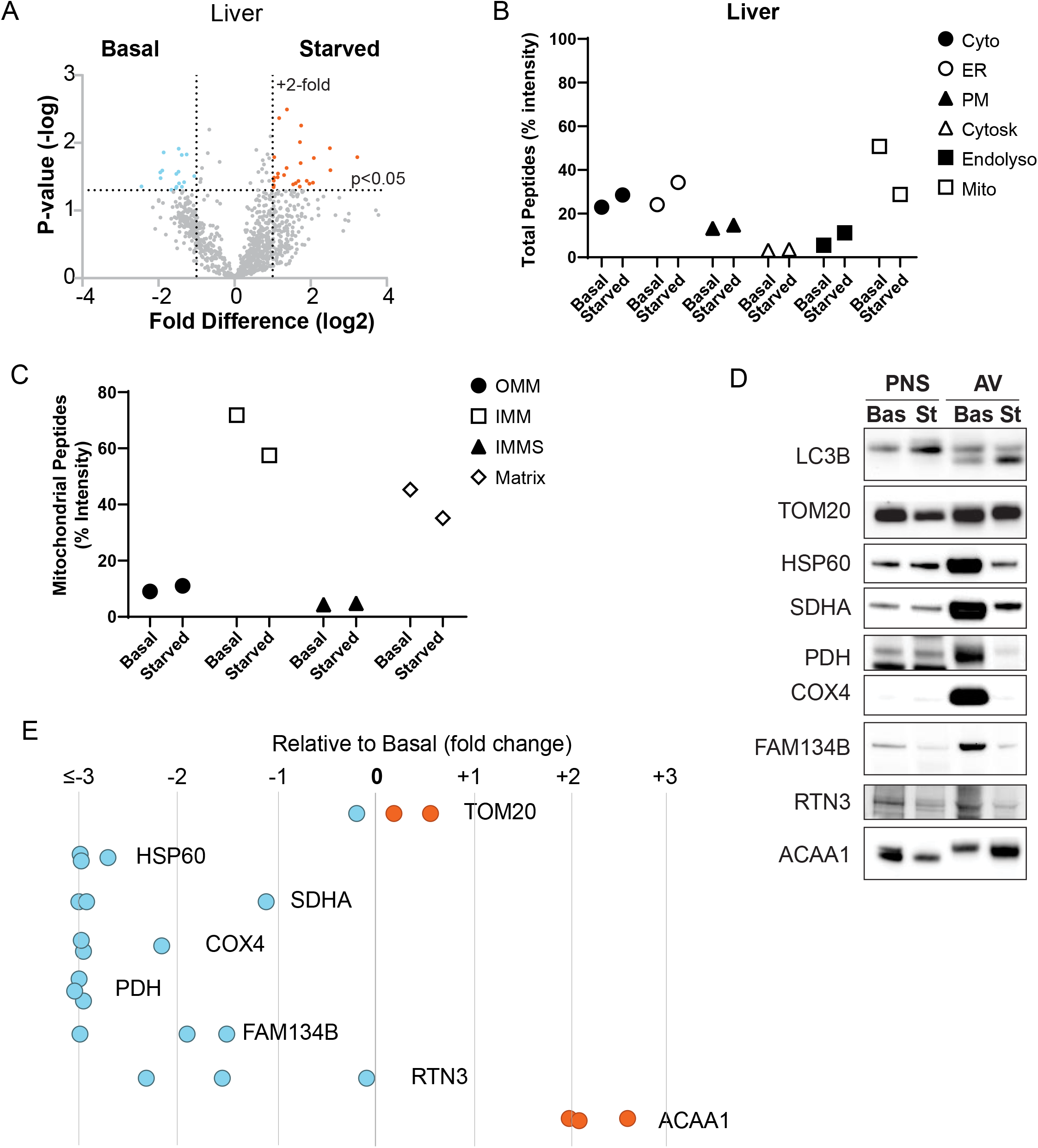
Liver cells move away from the turnover of intact mitochondrion to only the outer membrane in response to nutrient deprivation. **A**. Volcano plot comparing AV proteomes from basal and starved adult liver. Volcano plot of AVs isolated after 24hr nutrient deprivation (Starved) compared to basal conditions. Proteins greater than 2-fold difference, with a p < 0.05 are considered significant differences. Data points increased under starvation are shown in orange whereas decreased are shown in blue. Proteins that do not change are in gray. **B, C**. Decreased representation of IMM and mitochondrial matrix proteins in starved liver AV proteome. B. Quantitative comparison of basal and starved liver AV proteome based on the intensity of peptides as described in Fig. 1, with data in Table S4. Cytosolic (Cyto), endoplasmic reticular (ER), endolysosomal (Endolyso), Plasmalemmal (PM) and cytoskeletal (Cytosk). In contrast, mitochondrial compartments are lost from the proteome of starved AV. Examples of ER proteins include: ACSL1, ACSL5, Tmed10, POR, scarb2, PPIB, PDIA4, CPQ, ALB; Endolysosomal proteins: GRN, ANXA2, ATP6V0D1, CTSB, CTSD, CREG1, LIPA, TTR, GNS, GUSB, Lamp1, Lamp2, PSAP, DPP7, GBA, Man2a1; and Cytosolic proteins: LGMN, SCP2, MYL6, FTL1, ACP5. C. Comparison of the outer mitochondrial membrane (OMM), inner mitochondrial membrane (IMM), intermembrane mitochondrial space (IMMS), and matrix proteins. **D**,**E**. Validation of proteomic data analyses. PNS and AV fractions from basal (Bas) and starved (St) liver. 30 μg of each fraction has been loaded. E. Data shown as fold change relative to Basal level. These values were calculated as the fraction present in AV over total (n=3).The relative abundance to the basal AV fraction is shown in Representative images from the AV preparations are shown. TOM20 is an OMM, HSP60, SDHA, PDH are matrix proteins. COX4 is an IMM protein.

ER proteins represented greater than 20% of the basal liver proteome (Figure 1) and our analyses suggest that these proteins increase in representation upon starvation. ER proteins can be selectively eliminated by autophagy, using adaptor proteins such as FAM134B, RTN3, SEC62 and ATL3 (Grumati et al., 2018). The ER-phagy adaptors FAM134B, RTN3 were not detected in our proteome (Table S2), potentially due to being below detection levels rather than absent. SEC62 and ATL3 were slightly increased. Despite not being found in the proteome, and although ER proteins were increased in general upon starvation, the adaptor proteins FAM134B and RTN3 were among those decreased proteins upon starvation (Figure 2D,E). These data suggest that there is less selective turnover of the ER. Finally, proteins from organelles whose relative turnover increased upon starvation were the peroxisome and the endolysosomal system. Probing for the peroxisomal protein ACAA1 supports these observations (Figure 2D,E). Notable about the endolysosomal proteins is the increase in lysosomal proteins such as LAMP1 and LAMP2. These data suggest that there is an increased number of amphisomes and autolysosomes upon starvation. Taken together, these data indicate that there are distinct changes in the liver AV proteome upon starvation.

### The response of the brain AV proteome to nutrient deprivation

It is well known that in response to acute and prolonged starvation, the brain is uniquely privileged, and its ability to continue to expend energy is maintained at the cost of peripheral organs such as the liver and heart. Nonetheless, whether this privileged status equates with the inability of neural cells to mount an autophagic response is unclear (reviewed in (Griffey and Yamamoto, 2022)).

The Pearson correlation coefficients generated between basal and starved brain AV proteomes indicate a strong correlation between the two proteomes, suggesting little change in response to starvation (basal only: r = 0.943-0.961; starved only: r = 0.936-0.97; basal vs. starved: r = 0.909-0.967) (Figure S2C). Nonetheless, component 1 of PCA indicates that the greatest variability in the data sets is driven by the difference between starved and basal, suggesting that despite the overarching similarity, there are key distinctions in the data sets.

To explore this further, we again directly compared the proteomes. Volcano plot analyses yielded a key observation (Figure 3A). Although there was a small number of neuronal proteins that increased their representation in the starved brain AV proteome (4 out of 60 proteins), a far greater number of proteins uniquely expressed in non-neuronal cells (19 out of 60 proteins) such as Aldh1L1, AldoC, Pfn1,Ckb, Mog and Myh10 were increased. The remaining 37 proteins were proteins common across different cell types.

**Figure 3.**
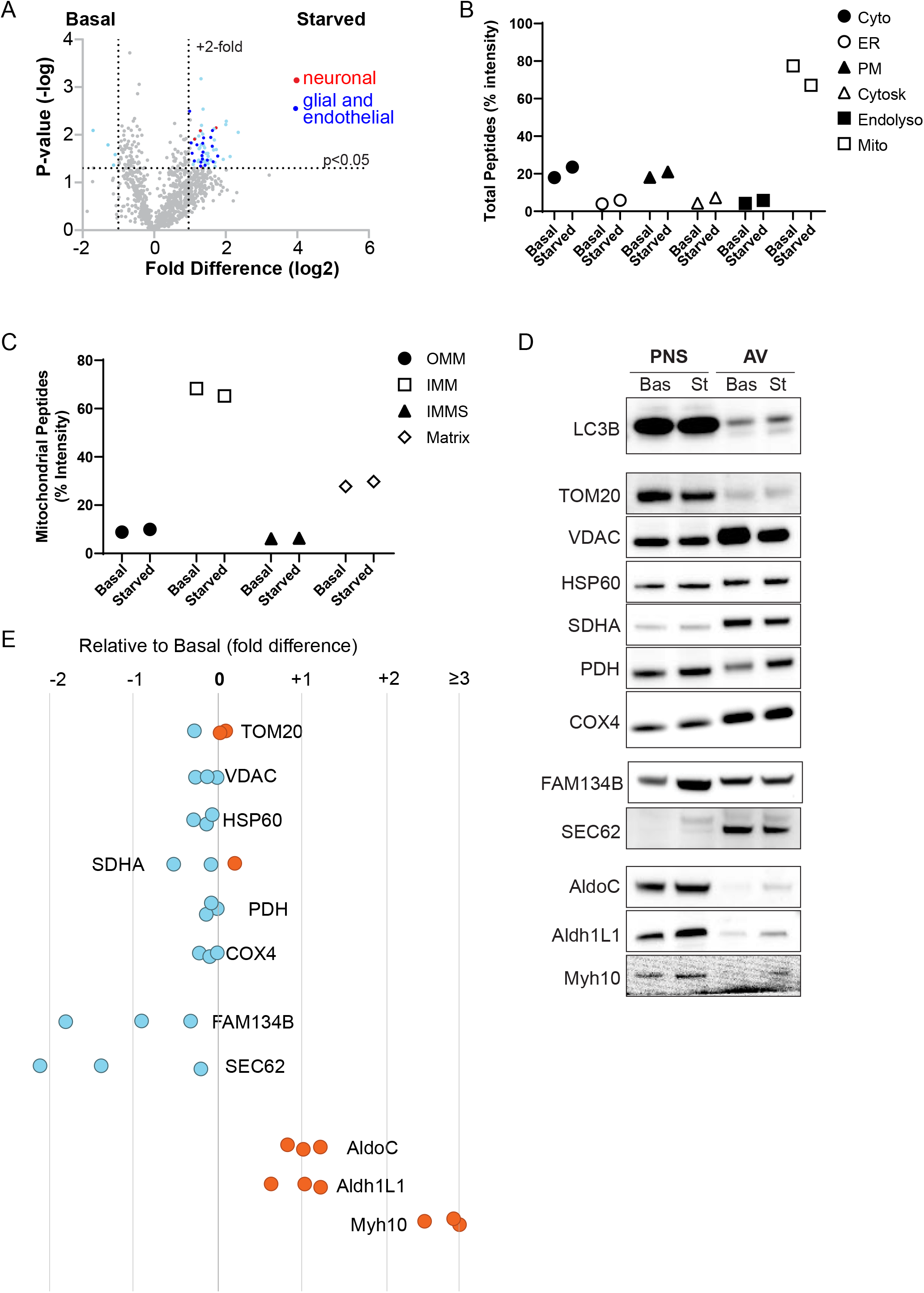
Cell type specific autophagic response to starvation in the brain. **A**. Quantitative comparison of basal and starved brain AV proteome reveals an increased representation of non-neuronal proteins in response to starvation. Comparison of brain AV proteome before vs after 24hr nutrient deprivation. Notably, the majority of cargoes that were increased were not neuronal proteins (indicated in red), but rather proteins exclusive to glial or brain endothelial cells (dark blue). **B**. Comparison of the intensity of peptides from different intracellular organelles relative to total peptide intensity between basal and starved brain (Table S4). The analysis suggests little difference between the conditions. **C**. Comparison of the OMM, IMM, IMMS and mitochondrial matrix between basal and starved brain AV. **D**,**E**. Validation and quantification of brain PNS (30 μg) and AV (30 μg) proteins during basal (Bas) and starved (St) conditions. Mitochondrial markers TOM20, VDAC, HSP60, SDHA, PDH, COX4 represent all regions of the mitochondria. ER phagy adaptor proteins are FAM134B and SEC62. AldoC and Aldh1L1 are astrocytic markers and Myh10 is an endothelial cell marker. D. Data shown as change in level relative to Basal levels in arbitrary units (a.u.). These values were calculated as the fraction present in AV over total (n=3).

Peptide intensity analyses revealed that there was a modestly increased representation of different cellular compartments in the starved brain AV proteome (Figure 3B,C). One exception to this was mitochondria, which showed a decrease, although less so than the change observed in the liver AV proteome (Figure 3B). Examining the representation of the different mitochondrial components, however, showed that unlike the change observed between basal and starved conditions in the liver (Figure 2C), there was little to no change detected in mitochondrial components between basal and starved conditions in the brain (Figure 3C). Immunoblot analyses move in a similar direction with the peptide analyses, showing a modest change in the levels of the different mitochondrial proteins (Figure 3D,E). Notably, the continued presence of both outer and inner components of mitochondria in starved brain AVs implies that whole mitochondrial turnover by autophagy is sustained in brain, unlike in liver, during starvation. Finally, similarly to liver, although the ER proteins modestly increased in peptide intensity upon starvation, probing for the ER-phagy adaptors FAM134B and SEC62 show a decrease, suggesting a potential movement away from selective ER turnover during starvation. Further characterization of other ER proteinsin both brain and liver AV fractions will be necessary to describe fully what is occurring.

A striking observation in Figure 3 is the clear increase of proteins expressed exclusively in non-neuronal cells in response to starvation. Canonical astrocytic proteins AldoC and Aldh1L1 were increased in our statistical analyses, as well as a lesser known brain endothelial cell (EC) marker, Myh10. Immunoblotting analyses indicates that these proteins are upregulated in the starved brain AVs. What’s especially remarkable about this observation is that ECs comprise of only 3-5% of the brain (Daneman et al., 2010; Motoike et al., 2000), and thus to be able to identify this protein by MS, and then validate biochemically suggests that the autophagic response in these cells is quite robust. These data suggest that the protein complexity found in brain-derived AVs increases after starvation, and that this complexity is driven by non-neuronal cells. Moreover, these findings strongly suggest that in response to physiologic starvation, different cell types in the brain mount a cell type specific response to the same physiological stress.

These data may reflect how non-neuronal cells are important for brain metabolism, and how neurons continue to receive nutrients despite starvation in the periphery. The ECs, as part of the neurovascular unit (NVU), line blood vessels and express the transporters responsible for bringing nutrients into the CNS. Astrocytes are also part of the NVU, and they interact with both the ECs and neurons. One possibility is that the autophagic response in these two critical cells types is the means by which the brain remains fed during starvation. Future studies examining the importance of autophagy in these cells during starvation can be achieved by genetic studies that manipulate autophagy in a cell type specific manner. To a larger point, these data also suggest that during disease conditions, autophagy may be affected differently across different neural cell types and can present as differential phenotypes. Future studies that can allow characterization of the AV proteome in a cell type specific manner may help answer these questions.

Although adaptor protein dependent autophagy pathways are increasingly characterized, autophagy is often characterized as a bulk degradation process, especially in response to nutrient deprivation. In light of this, we decided to test whether the proteomes we identified in the AVs were simply a reflection of the expression level of the total tissue proteome. We therefore proteomically characterized the corresponding postnuclear supernatants for the AV preps of both basal and starved liver and brain samples, then compared them to the proteomes of the AVs (Figure S3). Analyses revealed that the greatest correlation was observed between the nature of preparation within tissue (e.g. brain basal AV vs brain starved AV), and there was no correlation between the AV and PNS (e.g. brain basal AV vs brain basal PNS). PCA revealed that 69% of the variability was between the PNS and AV, whereas 20.4% could be accounted for by the differences across tissue. These data indicate that the AV cargo is not a simple reflection of the tissue proteome. Instead these data suggest that AV cargoes are deliberately selected for degradation, and may reflect a greater role for selective autophagy than bulk degradation in both basal and starved conditions.

### Tissue-specific reliance on autophagic degradation of mitochondria in response to starvation: From whole mitochondrial turnover to piecemeal mitochondrial turnover

Overall the data suggests that one of the critical roles for autophagy in both the liver and possibly more importantly in the brain is the turnover of mitochondria. Upon starvation, however, the data suggests that autophagic turnover of mitochondria changes in a tissue-specific manner: In brain there is little change, whereas in liver, autophagic degradation shifts from the wholesale turnover of mitochondria to piecemeal turnover of these organelles. This would indicate that starvation is distinct from other stressors, which often lead to mitochondrial fission and degradation. This re-analyses, however, still raises the intriguing hypothesis that in response to starvation, mitochondrial turnover by autophagy, especially the IMM and matrix proteins, is diminished in liver. To test this hypothesis, we next sought an independent approach by which we could determine if the internal components of the mitochondria were present within the autophagosome. Mitochondrial proteins can be created by the nucleus, but also can be directly created by the mitochondria by transcription and translation of mitochondrial DNA (mtDNA). Thus, we speculated that if mitochondria are present in AVs, we will be able to detect mtDNA by PCR. We performed phenol-chloroform extractions on fraction F3 given that mg quantities of the fraction were necessary to isolate reliably measurable levels of DNA. 100 ng of isolated DNA were then subject to PCR genotyping for four different sequences found on mtDNA: ND2, ND5, ATP6 and COXI (Figure 4).

**Figure 4.**
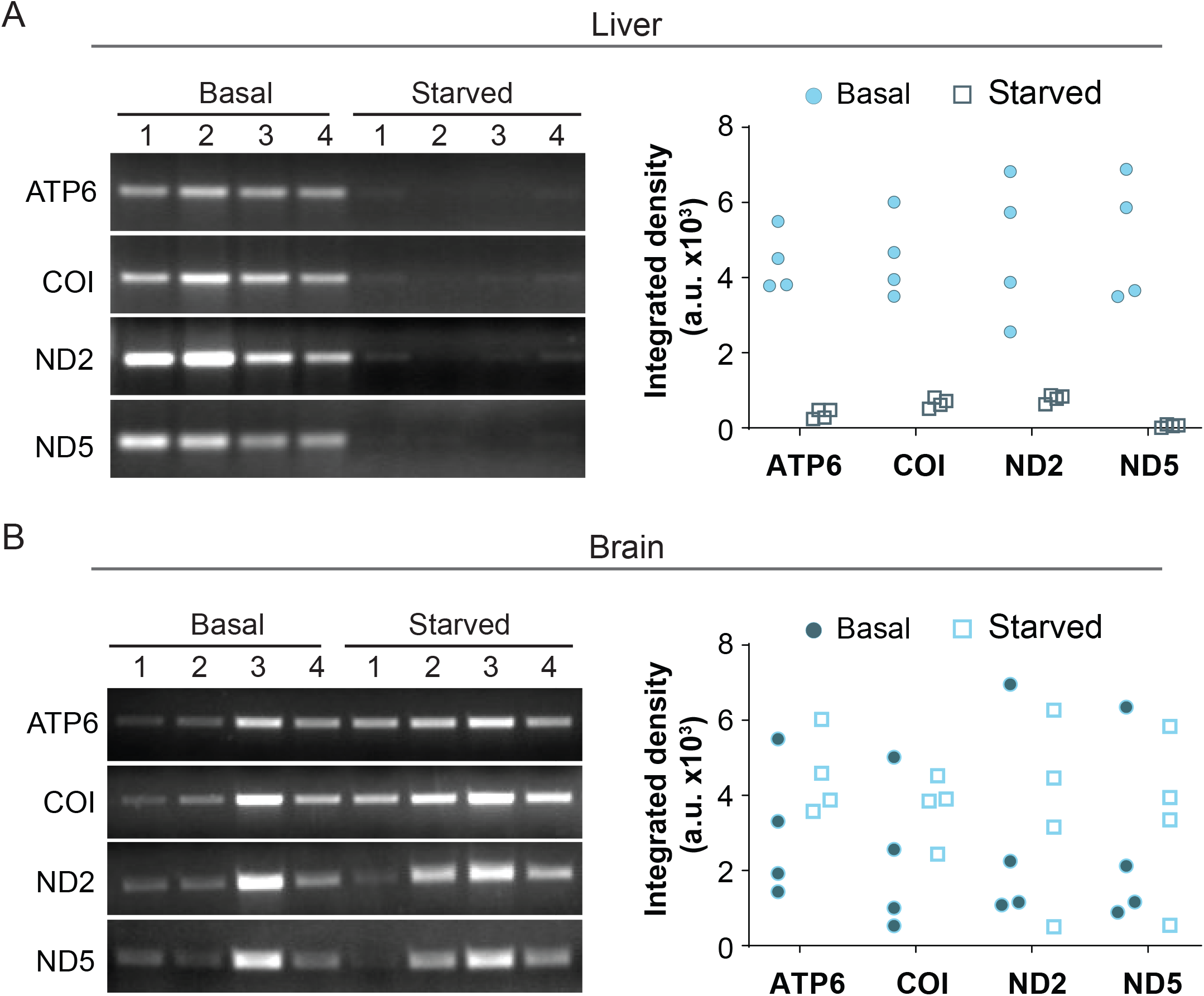
Independent validation of changes in mitochondrial targeting into AVs from starved liver vs. brain. PCR of DNA isolated from AVs indicate the loss of mitochondrial DNA in liver AVs upon starvation. DNA was isolated from the indicated AV fraction (n=4/treatment/tissue) by phenol-chloroform extraction. PCR analyses performed on 100ng of DNA for the presence of 4 mitochondrial genes, ATP6, COX1 (COI), ND2, and ND5. **A**. Results from Liver AVs. Two-sample t-test reveals a significantly decreased presence of mtDNA products in starved liver AVs (ATP6: p < 0.001, COI: p < 0.001, ND2: p = 0.006, ND5: p = 0.001). **B**. Results from Brain AVs. Two sample t-test reveals no significant difference between basal and starved conditions. (ATP6: p = 0.2151, COI: p = 0.2504, ND2: p = 0.704, ND5: p = 0.6551).

Under basal conditions, the presence of all four DNA sequences could be detected in AV fractions from both liver and brain (Figure 4). Although starvation did not significantly affect the presence of mtDNA in brain AVs, there was a significant reduction in their presence in the starved liver AVs. The loss of mtDNA supports the hypothesis that upon starvation, autophagy in hepatocytes transition from turning over intact mitochondria to either not turning over mitochondria at all or only partially in a piecemeal fashion. To distinguish between the two scenarios it would be necessary to perform observational studies in tissue or cells. First, studies suggest that upon starvation, mitochondria tubularize rather than fragment (Hailey et al., 2010; Rambold et al., 2011). We would confirm this in the liver, to determine if this happens in adult hepatocytes after 24hrs starvation. Second, using approaches that tag OMM proteins versus IMM proteins, we could determine if upon starvation the OMM proteins continue to enter into an acidified compartment whereas IMMs do not.

In contrast to the liver AVs, the sustained presence of mtDNA in brain AV suggest that there is little change in mitochondria turnover upon physiologic starvation. Whether this is a shared phenomenon across the different neural cell types or the neuronal response is dominating the proteome is unclear. Using approaches that allow us to determine the autophagosome proteome in a cell type specific manner would be particularly valuable to address this point.

What made the PCR experiments possible was that there were detectable levels of DNA present in the AV fraction, and although there was a loss of mtDNA in the starved liver AV fractions, we still were able to isolate comparable levels of DNA. One question this observation raises is what are the other sources of DNA in the autophagosomes? Several studies suggest that autophagy plays an important role in the DNA damage response (DDR) across phyla (Vessoni et al., 2013). The relationship between autophagy and DDR is complex, but reports suggest that damaged extranuclear DNA is found in autophagosomes (Bonne et al., 1999; Eriksson et al., 2003; Park et al., 2009). In addition to DNA, however, it is likely there is RNA present in the autophagosomes as well. We have not pursued this direction currently, but treatment with DNAse rather than RNAse followed by sequencing might provide interesting insights, especially in the starvation response. Given the prevalence of ribosomal proteins in all of the proteomes, as well as the potential for stress body turnover by autophagy (Monahan et al., 2016; Silva et al., 2019), this might also be important for neurodegenerative diseases such as ALS.

The role of autophagy in health and disease has been a question since the discovery of the pathway in the 1960’s. By optimizing the isolation of AVs and using a quantitative proteomic approach, we have found that tissues can rely on autophagy differently, gaining insight into why different organs might rely upon autophagy to maintain their health. For example, we find that the adult brain uses autophagy to turnover mitochondria and to maintain axonal and synaptic vesicle homeostasis, and thus it becomes less surprising that autophagy is implicated across very different adult onset neurodegenerative diseases. The introduction of starvation as a stressor also revealed not only how tissues might rely upon autophagy differently, but how discrete cell types within a given tissue might use autophagy differently in response to a given stress. This has strong implications therapeutically, given that simple upregulation or inhibition of autophagy might have highly unpredictable consequences. Taken together, by using this unbiased approach we have gained a better appreciation of the complex relationship this pathway has in a physiological context, and opened up new avenues of research to refine the often used statement, ‘autophagy is an essential pathway that plays an important role in health and disease.’

## Author contributions

YJY and HGJ led work on the liver and brain, respectively. YJY, HGJ, CJG, KK, AG, TJM, ND, AY generated the data; YJY, HGJ, ND, AY analyzed the data; YJY, HGJ, AY wrote the paper)

