## Supplemental Methods for "The tissue-specific autophagic response to nutrient deprivation"

### Materials and Methods

**Animals**

All experiments were reviewed and approved by the Columbia University Medical Center’s

Institutional Animal Care and Use Committee (IACUC). Mice were bred and housed in facilities

at the William Black Medical Research Building. Same-sex animals of mixed genotypes are

housed four to five per cage in a humidity- and temperature-controlled room, and mice are

provided food and water ad libitum. Mice are maintained on a 12 h light/dark cycle (lights on at

7:00 A.M.) *GFP-LC3B (GFPLC3+/+)* (Mizushima, 2009; Mizushima et al., 2004) was obtained from Dr. Noboru Mizushima (Tokyo Medical and Dental University, Tokyo, Japan).

**Genotyping**

*DNA isolation*

21-day-old mice were ear punched for identification and genotyping. The ear punches were incubated at 55°C overnight in 0.5 mg/ml proteinase K made into 500 μl lysis buffer (50 mM Tris-HCl pH 8.0, 100 mM EDTA pH 8.0, 100 mM NaCl, 1% SDS). DNA was extracted by adding 350ul isopropyl alcohol (Fisher Scientific) to the digested sample, which was then centrifuged at 14,000 rpm for 30 min. The supernatant was carefully removed, the pellet was washed with 800ul 80% ethanol and spun at 14,000 rpm for 10 min. The supernatant was removed and the DNA pellet was air-dried for 5 min at room temperature. Samples were resuspended in 50ul of 10mM Tris-HCL pH 8.0.

#### PCR

Each PCR reaction is 20 μl, consisting of 1 μl genomic DNA, 10 μM primer, 8 μl 5 PRIME HotMasterMix (5 PRIME GmbH), and water up to volume. The HotMasterMix contains Taq DNA polymerase, MgCl2, and dNTPs. Primers and PCR conditions is described below:

| Gene | Primer target | Primer sequence (5'-3') | Reagent | V per reaction (uL) | Temp | Time (m:ss) | # of cycle |
| --- | --- | --- | --- | --- | --- | --- | --- |
| GFP-LC3B | 1 | ATA ACT TGC TGG CCT TTC CAC T | primer 1 | 1 | 94 | 4:00 | 1 |
|  | 2 | CGG GCC ATT TAC CGT AAG TTA T | primer 2 | 1 | 94 | 0:30 | 30 |
|  | 3 | GCA GCT CAT TGC TGT TCC TCA A | primer 3 | 1 | 60 | 0:30 | 30 |
|  |  |  | mastermix | 8 | 72 | 1:00 | 30 |
|  |  |  | ddH2O | 21 | 72 | 1:00 | 1 |
|  |  |  | gDNA | 1 |  |  |  |

### Autophagic vesicle isolation (AV prep)

Protocol based on Strømhaug et al., 1998. For starvation, the animals were deprived of food 24 hours prior to experiment but were provided with water *ad libitum.* Each AV prep is a pool of 4-6 animals. Liver, brain and spinal cord of adult mice between the ages of 5.5 months – 6.5 months were collected and weighed. The tissues were homogenized in 1x SHB (0.25M sucrose, 10mM HEPES pH7.3, 1mM EDTA) with 1x protease inhibitor cocktail (PI) then spun at 4C, 2000 rpm, 10 minutes on Eppendorf 5810 centrifuge for nuclear pellet removal. The post nuclear homogenate (PNS) was first subjected to rate-zonal centrifugation on a discontinuous Nycodenz gradient (bottom - 22.5% Nycodenz+1x PI (Nyc) / 9.5% Nyc / PNS -top) at 144430xg for 1h. The layer at the 22.5% Nyc -9.5% Nyc interface was collected and subjected to rate-zonal centrifugation on a discontinuous Nyc/Percoll gradient (bottom – 22.5% Nyc / 33% Percoll / collected fraction – top) at 77670xg for 30 minutes. The two layers were collected separately. The collected lower fraction was measured and x0.7 Volume of 60% iodixanol (Optiprep) was added. The lower fraction + 60% iodixanol mixture was subjected to a discontinuous gradient (bottom – sample + 60% iodixanol / 30% iodixanol / 1x SHB) and centrifuged at 74870 xg for 30 minutes. This final step removed residual Percoll from previous gradients. The final band collected from this gradient was used as the crude AV fraction.

**Immuno-isolation**

The fractions from the AV prep were quantified via Bradford assay. 250ug of AV fraction was brought up to a total volume of 50uL using 1x SHB (0.25M sucrose, 10mM HEPES pH7.3, 1mM EDTA) + 1x Protease inhibitor cocktail (PI) then incubated for 2 hours at 4°C with 50uL of uMACS beads (Miltenyi). The magnetic columns were attached to MultiStand and prewashed with 200uL 1xSHB+1xPI. The sample + bead mixture was run through the magnetic column twice with gravity flow. The columns were then rinsed by gravity flow with 500uL 1x SHB + PI. The columns were released from the MultiStand and eluted into 1.5mL Microcentrifuge tubes with 50uL 1x SHB + 1xPI. A 200uL pipette was used to retrieve all residual eluate from column, which also consisted of beads.

**Mass spectrometry**

Total protein was harvested from AV crude or immunoisolates by standard trichloroacetic acid (TCA) method followed by acetone precipitation. Airdried protein pellets were resuspended then reduced then alkylated, then diluted to a final concentration of 2M urea prior to digestion with Endoproteinase Lys C (LysC) for 6 hours – over night at 37°C. Samples were further diluted to a final concentration of 1M urea and were digested with Trypsin at 37°C overnight, shaking. Upon desalting, digests were TMT labeled (10 channels). Peptides were analyzed using Thermo Orbitrap Fusion Tribrid mass spectrometer.

Proteomic profiling was done using LC-MS/MS with TMT 10. Proteomic data was analyzed using Perseus (version 1.6.2.3). Only proteins with 3 or more unique matching peptides were taken for further analysis. The raw intensity of each protein was corrected by total intensity prior to comparison and then log2 transformed to achieve normal distribution.

**Proteomic analysis**

Mass spectra were processed, filtered and quantified using SEQUEST-based software pipeline. (Huttlin et al., 2010). A filter of at least 2 or more peptides for each protein identification was set before cell component analysis. Further data analysis (including PCA and creation of volcano plots) were done using Perseus (MaxQuant, version 1.6.2.3). The raw intensity of each protein was corrected by total intensity prior to comparison and then log2 transformed to achieve normal distribution for subsequent statistical analysis.

**Mitochondrial DNA extraction and PCR**

MitochondrialDNA (mtDNA) was extracted as follows: Total DNA was isolated via phenol/chloroform extraction. The retrieved aqueous phase was treated with RNase (final concentration 100ug/ml), incubated at 37°C for 45 minutes. The isolation was followed with Ethanol precipitation, incubated in -80c for 1h, then spun down. DNA quality and quantity was measured by Nanodrop. mtDNA detection by PCR: DNA input of 100ng per PCR reaction. PCR was 18 cycles, annealing temp 60C, primer concentration 0.4um each. Primers used are as below:

ATP6: forward - TTGCCCACTTCCTTCCACAA reverse - AGGAGGGTGAATACGTAGGCT

COI: forward - TACTATTCGGAGCCTGAGCG reverse - GATTTCCGGCTAGAGGTGGG

ND2: forward – ATAGGGGCATGAGGAGGACT reverse - TGGAAGGCCTCCTAGGGATAG

ND5: forward - CCCAGCTACTACCATCATTCAAGT reverse – GATGGTTTGGGAGATTGGTTGATGT.

The four genes are encoded on the mitochondrial genome.

**Antibodies**

The following antibodies were used for immunoblotting: rabbit anti LC3B (1:1000, Abcam ab48394), rabbit anti ACAA1 (1:1000, Thermo PA5-29956), rabbit anti ACBD5 (1:1000, Thermo, PA5-88011), rabbit anti COX IV (1:1000, CST #4850), rabbit anti FAM134B (1:500, Abcam ab15755), rabbit anti Hsp60 (1:1000, CST #12165), rabbit anti Myh10 (1:1000, Abcam ab684), guinea pig anti p62 (1:1000, Progen GP62-C), rabbit anti p62 (1:1000 Abcam ab109012), rabbit anti Pyruvate dehydrogenase (1:1000, CST #3205), rabbit anti RTN3 (1:1000, Thermo PA5-79941), rabbit anti SDHA (1:1000, CST #11998), rabbit anti SEC62 (1:1000, Bethyl Laboratories A303-981A), mouse anti TOM20 (1:1000, BD Sciences 612278), rabbit anti VDAC (1:1000, CST #4661), rabbit anti vinculin (1:1000, Invitrogen 700062), rabbit anti NBR1 (1:1000, CST #9891), rabbit anti-LC3A (1:1000, Abcam ab52768), rabbit anti-GABARAP (1:1000, Abcam, ab13733), rabbit anti-GABARAPL1 (1:500, Abcam, ab86497), rabbit anti-GABARAPL2 (1:1000, Abcam, ab126607). Synaptic proteins: rabbit anti Synapsin-1 (1:2000, Stressgen VAP-SC060), rabbit anti CAMK2 (1:1000, CST #3362), rabbit anti SNAP25 (1:1000, CST #5308), mouse anti Synaptophysin (1:2000, Chemicon #5258), mouse anti Synaptotagmin (1:1000, Chemicon MAB 5200), rabbit anti VAMP2 (1:1000, Chemicon ab5826), rabbit anti VIAAT (1:1000, Invitrogen PA5-27569), rabbit anti VGLUT1 (1:1000, Synaptic systems #135-302), rat anti NCAM (1:2000, Millipore MAB5272), rabbit anti NMDAR2b (1:1000, Sigma M-265), mouse anti PSD-95 (1:1000, Chemicon MAB1596)
