## Supplemental Figures and Legends for "The tissue-specific autophagic response to nutrient deprivation"

Yang and Grosso Jasutkar et al Suppl. Fig. 1

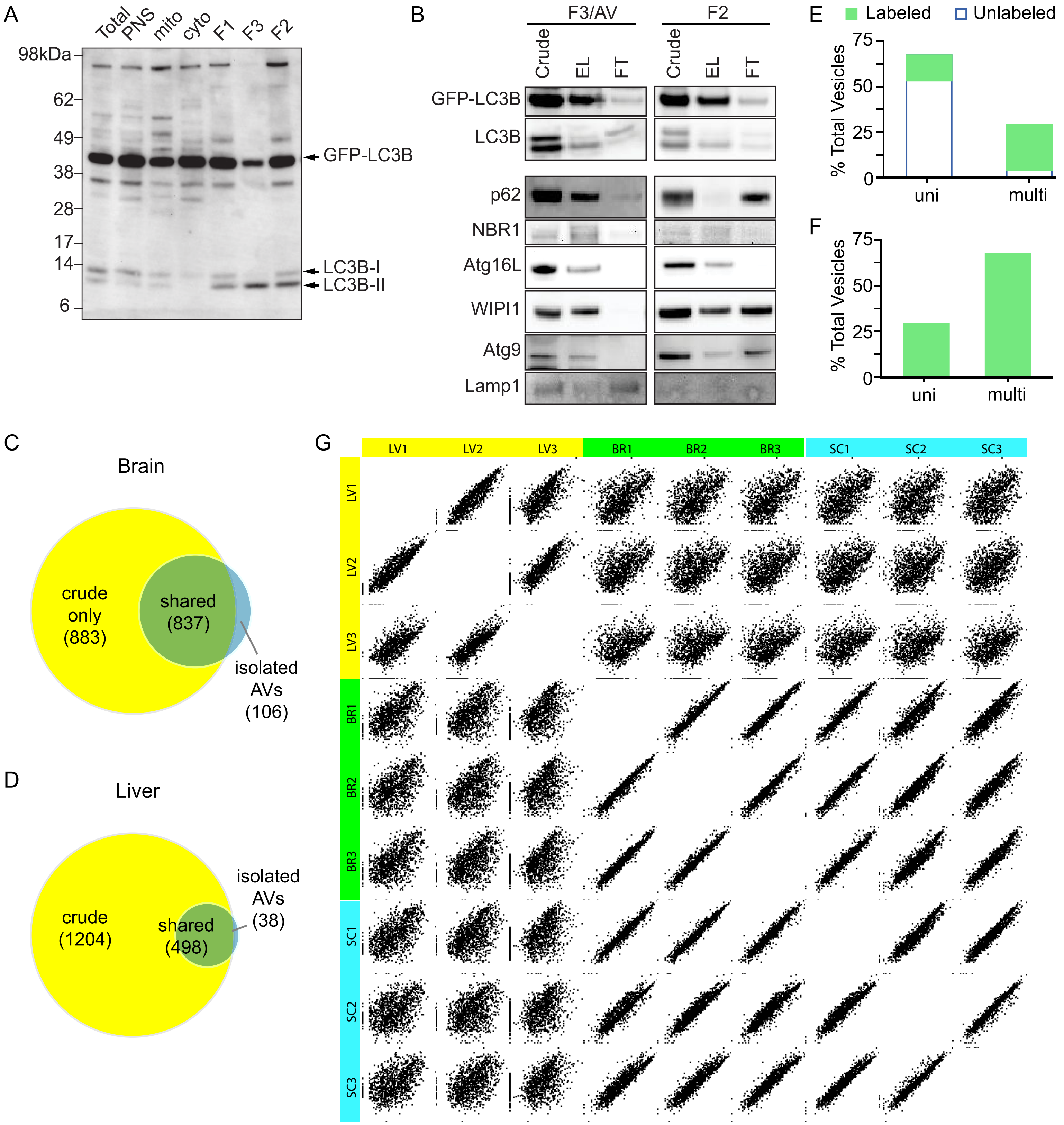

Yang and Grosso Jasutkar et al Figure S2

LIVER

A

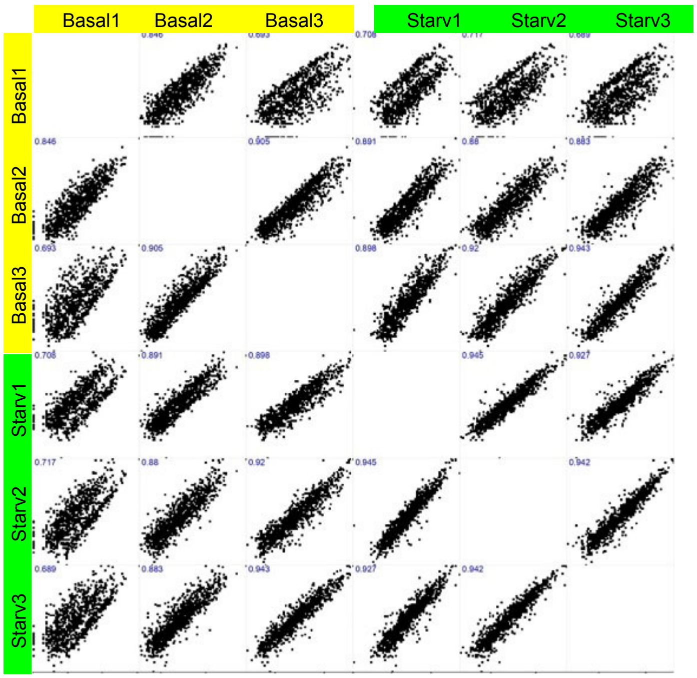


B

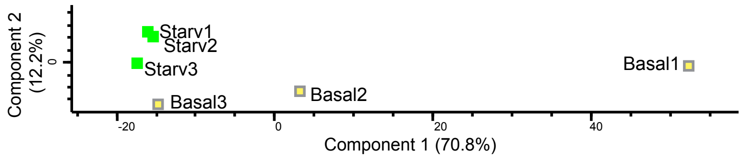


BRAIN

C

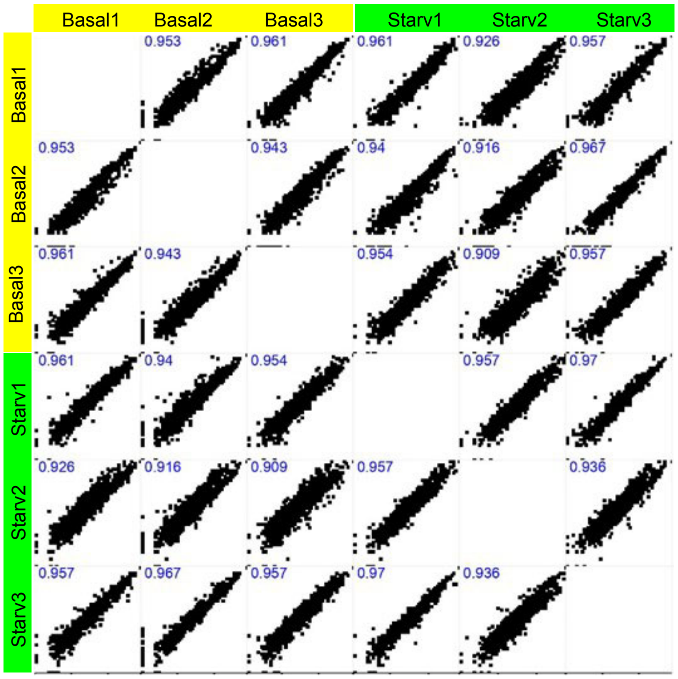


D

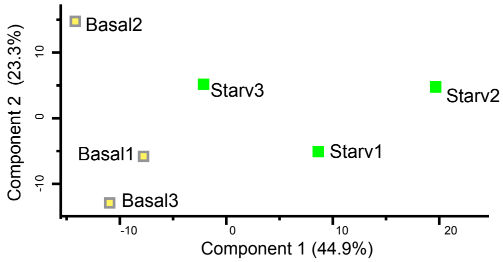

Yang and Grosso Jasutkar et al Suppl. Fig. 3

A

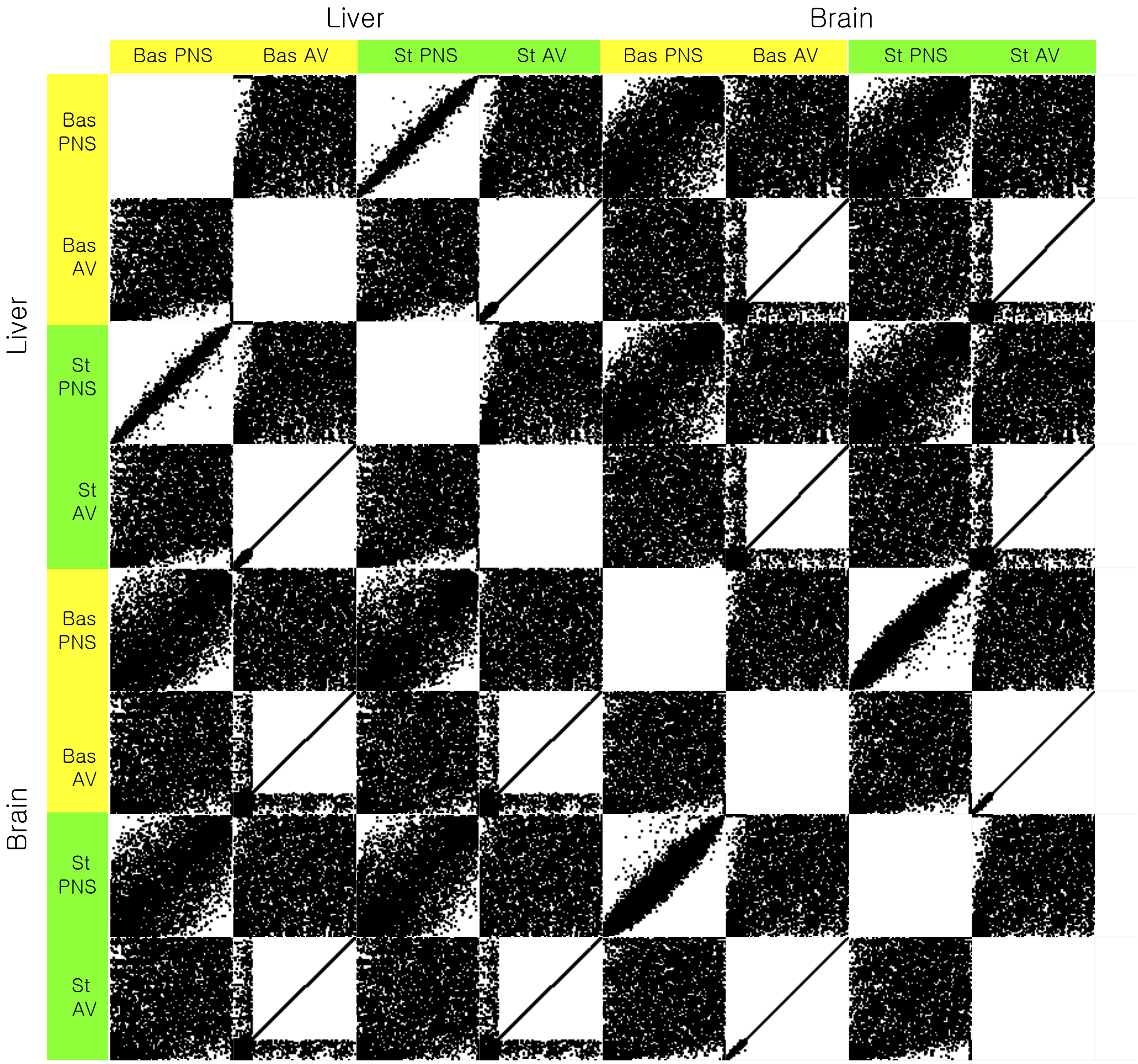


B

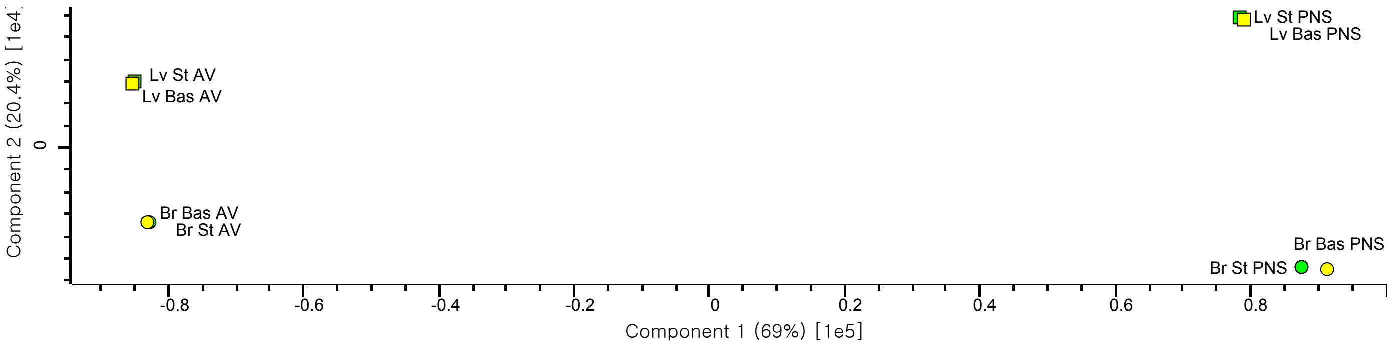

**Supplemental Figure 1. Initial characterization of immunoisolated AVs from brain and liver of GFP-LC3 mice.**

**A.** Endogenous and GFP-LC3B levels in AV prep fractions. Cytosolic fraction, devoid of membranous structures only contains cytosolic form LC3B-I. F3/AV fraction presents an enrichment of LC3B-II, the autophagosomal membrane bound form.

**B.** Western blot analysis of immunoisolation of fractions F3 and F2. Immunoisolation of F3 selectively enriches LC3B-II positive structures. F3 is positive for autophagy cargo adaptor proteins such as p62 and NBR1, and early autophagosome proteins such as Atg16L, WIPI1, and Atg9. Lysosomal marker Lamp1 is lost after immunoisolation. Immunoisolation of F2 finds both GFP-LC3B and endogenous LC3B-II, indicative of the presence of autophagic membranes, but are not associated with p62. Mildly positive for Atg16L, WIPI1 and Atg9.

**C, D.** Venn diagram summarizing the proteomic contents identified from the crude vs. immunoisolated AV preps after TMT-MS from 3 identical preps in brain and liver.

**E, F.** Quantification of unilamellar versus multilammelar vesicles E. relative to the total number of vesicles E. labeled (green) vs. unlabeled (white) or F. labeled alone.

**G.** Summary of MS LC/LC data generated for AVs isolated from wildtype liver (LV), brain (BR) and spinal cord (SC). a. Pearson correlations. Sample LV3 and to a lesser extent LV2 were quite variable and are currently being re-processed (see Figure III.). The correlation within brain AV preps and within SC AV preps were quite strong. In addition, the correlation between brain and spinal cord was equally strong indicating a high similarity in cargo between these two tissues.

Supplemental Figure 2. Initial analyses of the basla and starved immunoisolated AV proteomes for liver and brain.

Scatterplots of proteomic data (A, C) and PCA (B, D) of liver (A, B) and brain (C, D) samples. Pearson correlation coefficients are embedded into the scatterplots.

Supplemental Figure 3. The AV and PNS proteomes are distinct under basal and starved conditions for liver and brain.

A. Scatterplot and B. PCA comparing AV and PNS proteomes of basal and starved, liver and brain samples. Comparisons reveal the greatest similarity between type of preparation (AV vs PNS), then the tissue (liver vs brain).
