## Supplemental Tables for "The tissue-specific autophagic response to nutrient deprivation"

**Table S1: Crude (Pre-immunoisolation) proteome of Liver, Brain and Spinal Cord AVs**

Only proteins that were detected with more than 3 unique peptides are listed.

The numerical values are peptide intensity normalized to total intensity of each sample.

| uniprot | gene symbol | Intensity of Lv1 | Intensity of Lv2 | Intensity of Lv3 | Intensity of Br1 | Intensity of Br2 | Intensity of Br3 | Intensity of SC1 | Intensity of SC2 | Intensity of SC3 |
|---|---|---|---|---|---|---|---|---|---|---|
| Q99MN9 | *Pccb* | 74.7 | 115.7 | 113.3 | 3340 | 3304.1 | 3111.6 | 1863.8 | 1231.2 | 1062.2 |
| P61027 | *Rab10* | 610.5 | 547.8 | 463 | 4401.3 | 4166 | 4115.9 | 3235.9 | 1815 | 2032 |
| Q9CR57 | *Rpl14* | 132.6 | 376.1 | 311.9 | 9020.7 | 8316.2 | 8394.6 | 7117.3 | 3125.5 | 3914.1 |
| Q6PIC6 | *Atp1a3* | 896.5 | 584.5 | 550.8 | 12179.9 | 12606.7 | 11376 | 9534.8 | 5464.6 | 6222.6 |
| P05063 | *Aldoc* | 889.6 | 674.4 | 409.1 | 5785.6 | 5863.1 | 5416.8 | 5644.2 | 3023.2 | 3415.6 |
| Q99L04 | *Dhrs1* | 20.1 | 20.2 | 15.2 | 177.3 | 200 | 187.2 | 140.3 | 545.7 | 331.7 |
| P55264 | *Adk* | 1142 | 979.5 | 621.6 | 5204 | 5250.3 | 4940.6 | 5257.2 | 4629.8 | 3665.2 |
| P80318 | *Cct3* | 31.9 | 45 | 33.6 | 780.3 | 708.3 | 815.7 | 633.3 | 374.9 | 385.3 |
| Q9WUM3 | *Coro1b* | 12.3 | 24.6 | 29.8 | 1298.1 | 1383.1 | 1507.2 | 1286.3 | 3096.8 | 2834.8 |
| Q99PU5 | *Acsbg1* | 885.8 | 745.9 | 792 | 4153.2 | 4690.7 | 4392.4 | 2806.6 | 2496.7 | 1991.8 |
| Q8C1B7 | ***42989*** | 3694.6 | 1987.1 | 1133.6 | 23102.8 | 21392 | 22636.9 | 28378.7 | 12166.8 | 14880.6 |
| Q61207 | *Psap* | 61.1 | 59.2 | 42.9 | 764 | 887.9 | 783.9 | 1629.7 | 2920.6 | 2442.1 |
| P62962 | *Pfn1* | 153.1 | 127.3 | 97.7 | 510.7 | 526.4 | 480.2 | 348.8 | 266.9 | 241.2 |
| Q9DCG6 | *Pbld1* | 10264.2 | 9792 | 8947.2 | 2099.2 | 2482.4 | 1927.7 | 2452.6 | 1273.7 | 1818.5 |
| Q9JIL4 | *Pdzk1* | 84.4 | 65.5 | 43.5 | 414.1 | 442.5 | 387.6 | 384.3 | 231.5 | 256.7 |
| Q91V14 | *Slc12a5* | 24.8 | 34.9 | 20.5 | 303.9 | 283 | 336.6 | 191.5 | 182 | 138.6 |
| P31650 | *Slc6a11* | 2160.3 | 2156.8 | 1709.1 | 17663.1 | 18349 | 15452.4 | 14875.4 | 10329.8 | 9527.5 |
| UPSP:K1CN_HUMAN | | 185.3 | 130.8 | 108.7 | 2040.3 | 2370.8 | 2003.4 | 3703.5 | 5665.6 | 5008.3 |
| P29391 | *Ftl1* | 610.8 | 316.2 | 224.8 | 2534.3 | 2442.7 | 2387.1 | 2367.7 | 1390.2 | 1473.7 |
| Q8BMF4 | *Dlat* | 1634 | 1275.1 | 947.3 | 6171.7 | 7011 | 6846.4 | 3811.5 | 3335.5 | 2898.8 |
| P23492 | *Pnp* | 130.8 | 336.4 | 531.8 | 13157.2 | 16117 | 14252.5 | 9424.4 | 10875.5 | 7295.7 |
| P35486 | *Pdha1* | 23912 | 27431.6 | 28819.1 | 2943.4 | 1503.1 | 1557.2 | 2478.3 | 1391.6 | 1797.7 |
| P51881 | *Slc25a5* | 629.6 | 584.8 | 445.4 | 4697.9 | 5654.9 | 5056.2 | 4301.8 | 3779.4 | 3720.7 |
| Q6PHZ2 | *Camk2d* | 35.2 | 68.9 | 79.5 | 2093.1 | 1711 | 1819.7 | 1426.1 | 885.5 | 736.6 |
| Q91WN4 | *Kmo* | 6.9 | 23 | 16.9 | 260.5 | 266.7 | 221 | 338.1 | 718.2 | 589.2 |
| Q91YW3 | *Dnajc3* | 89.3 | 212.8 | 243.4 | 3341.8 | 4162.2 | 3714.8 | 4739.5 | 9121.4 | 8163.3 |
| Q61282 | *Acan* | 7.1 | 22.1 | 19.7 | 345.7 | 433.3 | 401.1 | 206.1 | 111.7 | 125.3 |
| P47911 | *Rpl6* | 201.3 | 113.4 | 63 | 900.2 | 1034.4 | 1038.1 | 887 | 597.4 | 574 |
| Q9Z1P6 | *Ndufa7* | 988.5 | 823.8 | 579.7 | 4644.8 | 5619.1 | 5397.4 | 5021.6 | 4247.2 | 5073.6 |
| Q03265 | *Atp5a1* | 355.3 | 200.4 | 117.9 | 1855.3 | 1703.4 | 1569.2 | 2398.3 | 1250.3 | 1397.2 |
| P80316 | *Cct5* | 19.6 | 22.5 | 36.1 | 334.9 | 313.2 | 267.3 | 410.8 | 684 | 595 |
| Q9ESP1 | *Sdf2l1* | 274.5 | 143.8 | 117.8 | 956.2 | 1093.3 | 1047.8 | 807.6 | 643.9 | 609.5 |
| Q64176 | *Ces1e* | 530.2 | 565.1 | 634.8 | 3283.5 | 3103 | 2676.8 | 4051.1 | 2127.8 | 2264.9 |
| Q6PIE5 | *Atp1a2* | 1316.7 | 916.4 | 546 | 6640.5 | 7587.7 | 8150.7 | 6439.5 | 4021 | 4897.7 |
| Q3UIU2 | *Ndufb6* | 53.5 | 87 | 69.7 | 209.1 | 204.7 | 198.3 | 251.9 | 505.7 | 427.3 |
| Q9D0M5 | *Dynll2* | 354 | 395.9 | 456.8 | 3.4 | 9.9 | 10.5 | 8.4 | 3.5 | 5 |
| P17879 | *Hspa1b* | 41 | 33.8 | 27.5 | 314.5 | 394.9 | 381 | 597.9 | 686.9 | 1143 |
| O88696 | *Clpp* | 0 | 11.7 | 20.1 | 146.4 | 182.5 | 170.6 | 265.7 | 406.4 | 394 |
| Q9Z2Q6 | ***42983*** | 3471.9 | 2633.6 | 1898.1 | 13100.4 | 15490.3 | 13345.1 | 11966.1 | 8682.2 | 8281.8 |
| P35564 | *Canx* | 149.3 | 75 | 62.4 | 1028.6 | 821.2 | 935.7 | 1052.9 | 934.5 | 916.9 |
| O08749 | *Dld* | 837.8 | 433.1 | 299 | 2940 | 3100.8 | 2749.8 | 2830 | 1554.7 | 1709.9 |

| | | | | | | | | | | |
|---|---|---|---|---|---|---|---|---|---|---|
| P16675 | ***Ctsa*** | 3338.4 | 1988 | 1346.4 | 10215.4 | 10461.8 | 9670.8 | 9223.5 | 4901.9 | 5710.4 |
| Q8C5H8 | ***Nadk2*** | 651.6 | 470.3 | 445.2 | 3238.2 | 2744.6 | 2665.3 | 2697.1 | 1543.6 | 1693.2 |
| Q9WVA4 | ***Tagln2*** | 140.7 | 174 | 140.6 | 8119.1 | 9002.4 | 10629.8 | 8699.1 | 5404.3 | 6019.4 |
| O08692 | ***Ngp*** | 20.2 | 58.5 | 47.2 | 882.5 | 1052.3 | 818 | 1101.3 | 1996.3 | 1693.8 |
| Q64459 | ***Cyp3a11*** | 1368 | 1683.3 | 1786.9 | 49.9 | 80 | 36 | 36.7 | 18.1 | 51.7 |
| Q8BH95 | ***Echs1*** | 2200.3 | 1899.2 | 2248.2 | 775.5 | 778.6 | 707 | 1314.8 | 720.7 | 985.7 |
| Q8R0Y6 | ***Aldh1l1*** | 4.5 | 64.5 | 127.7 | 747.4 | 871 | 719.1 | 809.5 | 725.9 | 663.1 |
| Q9WU79 | ***Prodh*** | 1768.9 | 938.3 | 722.8 | 5389.7 | 5954.1 | 6100.2 | 5153.6 | 3412.8 | 4071.7 |
| Q61838 | ***Pzp*** | 1284.3 | 778.9 | 455.8 | 4270.1 | 3856.2 | 4092.4 | 4355.9 | 2503.3 | 2682.2 |
| P32261 | ***Serpinc1*** | 597.2 | 360.1 | 209.4 | 1731.6 | 1740.6 | 1761.2 | 2212.5 | 1230.7 | 1117.7 |
| Q8VCC2 | ***Ces1*** | 62.9 | 97.6 | 107.5 | 1377.2 | 1506.9 | 1146.5 | 1468.2 | 1294.7 | 1022.6 |
| P26041 | ***Msn*** | 46.5 | 60.9 | 45.7 | 197.8 | 232 | 192.8 | 166 | 113.8 | 135.8 |
| Q9CR84 | ***Atp5g1*** | 18.6 | 46.1 | 43.9 | 148 | 160.2 | 166.9 | 178.8 | 164.7 | 179.1 |
| P63328 | ***Ppp3ca*** | 897.5 | 775.3 | 724.1 | 7827.8 | 10064.4 | 8306.1 | 6451.3 | 5770.8 | 4963.3 |
| P47738 | ***Aldh2*** | 1904.9 | 1041.5 | 675.8 | 8002.9 | 9965.8 | 9353.5 | 9861.6 | 6802.4 | 7433.4 |
| P21279 | ***Gnaq*** | 356.1 | 330.7 | 377.5 | 924.3 | 856.7 | 780.3 | 829.9 | 472.7 | 629.7 |
| Q8JZZ0 | ***Ugt3a2*** | 24.5 | 68.5 | 69.6 | 495.6 | 597.2 | 470.9 | 539.9 | 2676.6 | 1279 |
| P13595 | ***Ncam1*** | 45.4 | 256.2 | 234 | 1993.5 | 1663.5 | 2159.4 | 2134.8 | 4611.2 | 3795.5 |
| P50518 | ***Atp6v1e1*** | 757.6 | 375.7 | 327 | 2205.3 | 2587.1 | 2462 | 1869.3 | 1289 | 1393.2 |
| Q9D0K2 | ***Oxct1*** | 17.6 | 56.9 | 58.9 | 791.6 | 816 | 615.4 | 712.6 | 2161.6 | 1080.2 |
| Q9QXX4 | ***Slc25a13*** | 143.3 | 85.4 | 77.2 | 1986.2 | 2509.7 | 2702.7 | 1540.4 | 1209.3 | 1381.2 |
| P61028 | ***Rab8b*** | 3719 | 4416.5 | 4445.4 | 1207.3 | 486.7 | 695.1 | 933.8 | 647.1 | 692.7 |
| P29699 | ***Ahsg*** | 455.8 | 448.5 | 455.8 | 188.5 | 87.1 | 111.1 | 136.3 | 118.9 | 105.5 |
| P63030 | ***Mpc1*** | 3382.5 | 2153 | 1523.4 | 10669.3 | 11098.1 | 9302.3 | 10541.3 | 5767.4 | 5994.3 |
| Q61885 | ***Mog*** | 43.5 | 72.5 | 96 | 271.1 | 258.1 | 237.7 | 277.2 | 456.4 | 428.7 |
| P20357 | ***Map2*** | 711.7 | 676.8 | 574.4 | 2891.4 | 2431.6 | 2297.6 | 2515 | 1308.3 | 1512.6 |
| P48962 | ***Slc25a4*** | 1041.4 | 740.2 | 648.8 | 4688.8 | 6212.9 | 5503.5 | 4096.4 | 2888 | 3564.4 |
| Q91XD4 | ***Ftcd*** | 5.6 | 37.6 | 35.7 | 868.8 | 1058.8 | 766.9 | 939 | 2872.8 | 2101.2 |
| Q60936 | ***Coq8a*** | 126.2 | 734.5 | 1744.1 | 10306.6 | 12907.7 | 10127 | 16097.6 | 55725.4 | 31093 |
| Q91VM9 | ***Ppa2*** | 1208.1 | 782.9 | 449.8 | 3053.3 | 3384.6 | 3266.3 | 3356.7 | 2552 | 2068.1 |
| P61255 | ***Rpl26*** | 14.3 | 29.2 | 26.5 | 184 | 234.5 | 246.2 | 415.6 | 1212.1 | 944.2 |
| O35465 | ***Fkbp8*** | 1752.5 | 1196.9 | 1184.3 | 9111.4 | 7625.1 | 6995.5 | 12015.2 | 5096.3 | 5788.5 |
| Q60931 | ***Vdac3*** | 2701.6 | 1702.1 | 1355 | 8204.2 | 10324.4 | 9872.8 | 7378.9 | 5151.2 | 6316.1 |
| Q9CPV4 | ***Glod4*** | 435.4 | 342 | 190.6 | 1283.1 | 1520 | 1290 | 1201.8 | 945.6 | 872.5 |
| O35604 | ***Npc1*** | 617.8 | 275.5 | 238.3 | 1637.2 | 1828.6 | 1657.2 | 1648.7 | 1112.5 | 1178.8 |
| P18760 | ***Cfl1*** | 23.5 | 44 | 35.7 | 271.9 | 350.3 | 270.7 | 295.1 | 393.7 | 359.8 |
| P17426 | ***Ap2a1*** | 9946.6 | 11934.4 | 12160.3 | 3645.8 | 1755.3 | 2610.1 | 2481.7 | 2754.8 | 2610.9 |
| O89053 | ***Coro1a*** | 109.9 | 127.2 | 113.7 | 4489.2 | 6133.6 | 6257.2 | 2187.8 | 1557.8 | 1667.6 |
| P62075 | ***Timm13*** | 94.3 | 68.9 | 47.3 | 714.8 | 537.6 | 714.7 | 527.1 | 341.1 | 444.9 |
| Q9WV98 | ***Timm9*** | 67.7 | 60.7 | 66.9 | 1125.1 | 1590.7 | 1445.8 | 1667 | 2701.3 | 3195.9 |
| Q8VC12 | ***Uroc1*** | 8.7 | 9.6 | 11.5 | 141.6 | 167.8 | 199.4 | 400.8 | 564.7 | 689.5 |
| Q8VCU1 | ***Ces3b*** | 0 | 85.9 | 97.8 | 389.6 | 460.7 | 407.5 | 744.1 | 1393.1 | 1175.5 |
| Q9WUB3 | ***Pygm*** | 246.9 | 147.4 | 109.7 | 567 | 602.6 | 559.9 | 611.8 | 354.5 | 448.1 |
| Q8K0E8 | ***Fgb*** | 294.5 | 166.6 | 140 | 670.9 | 746.5 | 763.3 | 582.4 | 530.4 | 489.8 |
| P56480 | ***Atp5b*** | 929.2 | 471.8 | 333.8 | 2700.8 | 2495.2 | 2349.7 | 2610 | 1442.1 | 1521.2 |
| Q9D051 | ***Pdhb*** | 19.9 | 23.3 | 21.8 | 276.1 | 347.1 | 249.8 | 534 | 965.4 | 766.5 |
| P30275 | ***Ckmt1*** | 965.1 | 419.6 | 419.1 | 2756.4 | 3429.3 | 3338.1 | 2644.4 | 2159.9 | 2525 |

| | | | | | | | | | | |
|---|---|---|---|---|---|---|---|---|---|---|
| Q99K67 | ***Aass*** | 1535 | 2139.6 | 2146.6 | 74.8 | 88.9 | 40.7 | 52.5 | 27 | 44.4 |
| P24549 | ***Aldh1a1*** | 2675.5 | 1460.4 | 925.1 | 7836.7 | 9053 | 9887.6 | 7423 | 5179.6 | 6315.9 |
| P26645 | ***Marcks*** | 442 | 432 | 201 | 1231.4 | 1474.3 | 1300.7 | 1059.5 | 869 | 680.9 |
| Q78JT3 | ***Haao*** | 677.7 | 650.1 | 593.4 | 209.2 | 96.9 | 241.5 | 176.5 | 121.9 | 194.4 |
| Q9D0S9 | ***Hint2*** | 1745.1 | 1052.1 | 726.6 | 5011.5 | 6332.1 | 6196 | 4835.8 | 3657 | 4306.2 |
| Q91YT0 | ***Ndufv1*** | 2391 | 1623.3 | 863.1 | 5928.2 | 5739.9 | 5549.6 | 6397.4 | 4071.2 | 4228.3 |
| Q9R0K7 | ***Atp2b2*** | 32.9 | 31.7 | 55.1 | 310.8 | 425.5 | 339.9 | 390.8 | 1676.3 | 955 |
| P68510 | ***Ywhah*** | 242.3 | 524.7 | 688 | 2811.8 | 3636.9 | 3748.3 | 4966.8 | 7958.9 | 7835.4 |
| Q9JII6 | ***Akr1a1*** | 2205.2 | 1806.3 | 1521.6 | 37.8 | 77.3 | 41.7 | 53.5 | 24.1 | 54.4 |
| Q505D7 | ***Opa3*** | 3716.5 | 2168.9 | 1679.8 | 22442.6 | 17694.8 | 16851.6 | 22663.1 | 9105.2 | 9767.3 |
| P17563 | ***Selenbp1*** | 38.8 | 39.1 | 52.9 | 114.7 | 125.3 | 142.9 | 278.5 | 343.9 | 369 |
| Q8R1I1 | ***Uqcr10*** | 1850 | 1462.5 | 2018.7 | 311.6 | 241.2 | 360.7 | 337.6 | 250.4 | 453.9 |
| Q9ERD7 | ***Tubb3*** | 894.6 | 531.5 | 292.3 | 2479.8 | 3123 | 2986.4 | 2577.4 | 1800.3 | 2090.4 |
| P61620 | ***Sec61a1*** | 1037.6 | 523.8 | 378.3 | 3538.1 | 3352.1 | 2798.4 | 2946.4 | 1885.5 | 1649.3 |
| D3YZV8 | ***Ccdc8*** | 64.9 | 307.8 | 199.5 | 789.5 | 869.9 | 828.2 | 875.1 | 3440.6 | 2134.3 |
| Q07417 | ***Acads*** | 447.8 | 190.2 | 119.5 | 1336.1 | 1143 | 1218.1 | 1303.4 | 772.2 | 717.1 |
| P10649 | ***Gstm1*** | 911.9 | 568.5 | 416.8 | 2073 | 2419.6 | 2098.4 | 2172 | 1337.4 | 1508.2 |
| Q91XE8 | ***Tmem205*** | 83.6 | 115.7 | 68.4 | 3112.4 | 2372.5 | 3563.5 | 2511 | 1441.4 | 1092.1 |
| P63011 | ***Rab3a*** | 1110.4 | 504.1 | 268.2 | 3451.2 | 3928.3 | 3119.7 | 4037.1 | 2477.6 | 2368.4 |
| Q80X85 | ***Mrps7*** | 10531.3 | 5590.7 | 3653.9 | 23330.2 | 23499.9 | 24400 | 21814.9 | 11768.6 | 12110.1 |
| Q3TWL2 | ***Tmem55b*** | 757.2 | 913.4 | 1138 | 26.5 | 25.8 | 14.4 | 19.3 | 8.6 | 10.1 |
| P13707 | ***Gpd1*** | 352.8 | 168.8 | 106 | 1182.4 | 971.6 | 978.4 | 1198.7 | 611.8 | 491.7 |
| O70435 | ***Psma3*** | 230.1 | 155.4 | 112.1 | 696.7 | 550.6 | 686.5 | 664.5 | 323 | 308.4 |
| Q71RI9 | ***Kyat3*** | 4429.3 | 2802.3 | 1891.7 | 9039.2 | 10108.6 | 9830 | 8769 | 5693.1 | 6131.7 |
| Q9ERE7 | ***Mesdc2*** | 27.9 | 30.6 | 39.4 | 301.6 | 441.9 | 357.5 | 321.4 | 831.8 | 659.4 |
| Q8JZS9 | ***Mrpl48*** | 0 | 6.8 | 10.6 | 80.6 | 99.3 | 118.9 | 275.4 | 255 | 534.4 |
| Q922Q1 | ***42796*** | 38.2 | 47.6 | 48.2 | 273.6 | 379.2 | 288.9 | 352.6 | 642.3 | 571.2 |
| Q9JIK9 | ***Mrps34*** | 6.7 | 6.4 | 4.1 | 136.9 | 150.5 | 200.5 | 532.9 | 1021.7 | 932.2 |
| Q3TYD4 | ***Arsg*** | 19.5 | 55.2 | 64.9 | 2377.3 | 2557.3 | 1684.3 | 825.5 | 2823.3 | 1131.2 |
| P46638 | ***Rab11b*** | 420.2 | 267.2 | 108.6 | 1228.3 | 1463.9 | 1619.2 | 1079.6 | 978.6 | 1173.4 |
| O08601 | ***Mttp*** | 215.7 | 177.8 | 123 | 2065.3 | 3103.7 | 2859.6 | 3114.7 | 7680.1 | 5740.4 |
| P30115 | ***Gsta3*** | 1011.3 | 1538.4 | 1236.4 | 35 | 63 | 52.5 | 36.7 | 93.6 | 80.9 |
| Q60605 | ***Myl6*** | 787.2 | 614.7 | 383.6 | 2417.7 | 2943.8 | 2222.9 | 2514.3 | 1221.5 | 1577.9 |
| Q811I0 | ***Atpaf1*** | 0 | 11 | 17.9 | 263.4 | 401.8 | 376.1 | 397.7 | 909 | 1008.8 |
| Q8BMF3 | ***Me3*** | 11.2 | 38.3 | 65.4 | 323.2 | 407.9 | 290.2 | 284.3 | 1107.9 | 560.4 |
| Q04519 | ***Smpd1*** | 2651.3 | 1825.3 | 1079.1 | 5370.2 | 5906.4 | 5547.3 | 6928.9 | 3811.1 | 4269.8 |
| Q61335 | ***Bcap31*** | 56.7 | 64.1 | 55.1 | 544.6 | 683.8 | 822.5 | 1958.6 | 1531.1 | 3522 |
| O70251 | ***Eef1b*** | 317.6 | 235.5 | 129.9 | 713.9 | 842.1 | 720.4 | 708.6 | 609.2 | 573.2 |
| Q9CPW2 | ***Fdx2*** | 619.7 | 345.3 | 231.2 | 1601.8 | 2128.6 | 1878.6 | 1553.2 | 1239.9 | 1312.9 |
| Q91XE0 | ***Glyat*** | 2348.4 | 3368.2 | 3583.8 | 188.2 | 182.9 | 153.5 | 187.2 | 130.4 | 188.8 |
| P24472 | ***Gsta4*** | 1039.6 | 1202.7 | 1569.2 | 66.8 | 94.5 | 35.9 | 50.2 | 72 | 82.1 |
| P08032 | ***Spta1*** | 1033.9 | 775.7 | 398.7 | 4447.2 | 6478.5 | 5379.1 | 2391.4 | 2319.4 | 2065.2 |
| Q91VN4 | ***Chchd6*** | 271.6 | 126.9 | 143.2 | 990.1 | 832.4 | 1167.8 | 1027.8 | 580.9 | 778.3 |
| P23953 | ***Ces1c*** | 11 | 25.7 | 29.6 | 1220.8 | 1406.2 | 889.6 | 1493 | 3842.9 | 2287.3 |
| Q01339 | ***Apoh*** | 374.2 | 499.1 | 408.6 | 146.8 | 124.1 | 145.2 | 127.1 | 231.6 | 231.6 |
| Q8BLK3 | ***Lsamp*** | 5241.9 | 7462.3 | 7984 | 581.9 | 405.1 | 506.5 | 969 | 838.9 | 1099.3 |

| | | | | | | | | | | |
|---|---|---|---|---|---|---|---|---|---|---|
| Q6GSS7 | ***Hist2h2aa1; Hist2h2aa2*** | 462.9 | 206.3 | 134.8 | 1095.1 | 1325.9 | 1149.9 | 859.9 | 672.6 | 742.6 |
| Q9CPQ8 | ***Atp5l*** | 6.8 | 28.9 | 26.6 | 1084.8 | 1370.7 | 875.2 | 952.4 | 2083 | 1515.3 |
| Q91Z53 | ***Grhpr*** | 2158.9 | 1215.7 | 961.8 | 7808.2 | 6131.7 | 8538.5 | 6928.4 | 3277.8 | 4244.9 |
| Q6R0H7 | ***Gnas*** | 367.6 | 43.6 | 25.6 | 1016.2 | 1078.8 | 1241.4 | 748.8 | 319.3 | 401.4 |
| Q8CGK3 | ***Lonp1*** | 653.6 | 801.4 | 769.2 | 293.6 | 84.3 | 127.4 | 230.2 | 131.2 | 140.5 |
| Q9CQN1 | ***Trap1*** | 1596.9 | 1044 | 530.2 | 3881.3 | 3459.8 | 3392.9 | 3150.1 | 2361.3 | 2135.5 |
| O09114 | ***Ptgds*** | 865.5 | 1332.7 | 979 | 25.4 | 31.2 | 29.6 | 28.6 | 27.9 | 38 |
| P03899 | ***Mtnd3*** | 400.7 | 380.5 | 321.2 | 100.3 | 65.5 | 158.1 | 91.7 | 106.2 | 108.5 |
| Q8K2B3 | ***Sdha*** | 2729 | 1677.5 | 1307 | 5934.7 | 7855.3 | 7047.7 | 6773.8 | 4955.2 | 6003.4 |
| P05201 | ***Got1*** | 209.5 | 153.2 | 165.6 | 49.9 | 35.9 | 55.9 | 27.9 | 31.5 | 36.9 |
| P34928 | ***Apoc1*** | 7638.2 | 10640.8 | 7120 | 494 | 495.2 | 875.9 | 595.9 | 520.1 | 1230.1 |
| Q32Q92 | ***Acot6*** | 8.4 | 39.5 | 54 | 2042.5 | 2841.4 | 1818.4 | 2088.7 | 5622.9 | 3838.9 |
| UPSP:K1CJ_HUMAN | | 86.3 | 432.1 | 417.4 | 1300.5 | 1667.2 | 1754.6 | 2347.9 | 5121.4 | 4298.2 |
| P48774 | ***Gstm5*** | 618.7 | 360.8 | 290.4 | 2418.6 | 3600 | 2836.9 | 1888 | 1997.8 | 1904.8 |
| P23780 | ***Glb1*** | 1896.6 | 2104 | 1496.4 | 560.6 | 634.9 | 525.5 | 573 | 728.7 | 631.5 |
| UPSP:K2C1_HUMAN | | 335.8 | 316 | 168.9 | 905.7 | 1245.8 | 1236.6 | 1225.9 | 942.7 | 972.9 |
| Q9CR68 | ***Uqcrfs1*** | 25.1 | 72.3 | 77.4 | 1660.7 | 2708.2 | 2112.9 | 2530.1 | 5604.4 | 4534.8 |
| P63158 | ***Hmgb1*** | 227.8 | 249.8 | 289.2 | 445.2 | 546.3 | 488.3 | 388.1 | 1122 | 890.9 |
| Q9D1G1 | ***Rab1b*** | 3478.9 | 1764.6 | 845.4 | 7586.5 | 9165.5 | 8021.1 | 8246.4 | 5735.7 | 5986.6 |
| P15636 | | 9.9 | 48.5 | 62.2 | 281 | 333.7 | 225.8 | 253.3 | 568.6 | 397.7 |
| P62717 | ***Rpl18a*** | 12.1 | 15.4 | 13.1 | 490.6 | 428.6 | 297 | 382.1 | 644.8 | 450.7 |
| P12265 | ***Gusb*** | 802 | 471 | 286.6 | 2585.4 | 3795.6 | 3046.1 | 3301.1 | 2035 | 2239.3 |
| P01867 | ***Igh-3*** | 32.6 | 69.5 | 110.1 | 507.9 | 505.6 | 349.7 | 398.6 | 871 | 664.6 |
| Q9QYR9 | ***Acot2*** | 3015.5 | 1804.1 | 1270.8 | 5755.1 | 7236.6 | 6812.5 | 6087.5 | 4259.7 | 3413.3 |
| Q2TPA8 | ***Hsdl2*** | 186 | 91.8 | 86.9 | 564.5 | 821.7 | 645.1 | 618.6 | 406.4 | 419 |
| P47740 | ***Aldh3a2*** | 83.7 | 457.9 | 756.8 | 1765.3 | 2092.3 | 2215.6 | 1987.5 | 5483.3 | 4490.3 |
| P63017 | ***Hspa8*** | 5.1 | 8.9 | 9 | 277.6 | 450.5 | 328.3 | 351.5 | 774.8 | 678.6 |
| P62204 | ***Calm1; Calm2; Calm3*** | 500.8 | 461.7 | 598.4 | 170.1 | 130.4 | 236.2 | 126.3 | 114 | 162.6 |
| P16332 | ***Mut*** | 3708.1 | 2254.3 | 1137.9 | 7474.8 | 8983.9 | 7998.8 | 7305.7 | 4830.7 | 4571.9 |
| P07356 | ***Anxa2*** | 556.7 | 918.2 | 688.5 | 11.7 | 33.5 | 11 | 9.5 | 5.2 | 15 |
| P51660 | ***Hsd17b4*** | 131 | 196.9 | 237.6 | 1837.2 | 2960.4 | 2828 | 2380.7 | 5025.4 | 5822.3 |
| Q99JB2 | ***Stoml2*** | 0 | 10.7 | 5.6 | 145.3 | 246.5 | 206.3 | 262.5 | 788.1 | 455.8 |
| Q3TCN2 | ***Plbd2*** | 534.9 | 746.3 | 538.9 | 116.7 | 104.5 | 173.9 | 147.7 | 196.3 | 222 |
| P51174 | ***Acadl*** | 1414.4 | 1909.3 | 1171.4 | 94.9 | 112.4 | 73.5 | 83.2 | 105.9 | 117.4 |
| Q8BK72 | ***Mrps27*** | 519.7 | 877.4 | 786.3 | 34.2 | 42.4 | 29.4 | 31 | 18.5 | 29 |
| Q80YN3 | ***Bcas1*** | 116.8 | 106.2 | 70.5 | 502.6 | 758 | 550.1 | 676.2 | 621.4 | 521.3 |
| Q8K0Z7 | ***Taco1*** | 20.5 | 34 | 55.7 | 1014.4 | 1142.2 | 665.3 | 959.6 | 3114.5 | 1699.8 |
| P21278 | ***Gna11*** | 1829.6 | 1945 | 2931.2 | 23.1 | 48.7 | 6.9 | 22.5 | 3.3 | 26.3 |
| Q99L47 | ***St13*** | 992 | 1337.7 | 1123.2 | 458.6 | 143.6 | 278.7 | 292.1 | 251.3 | 255 |
| Q8R086 | ***Suox*** | 156.6 | 88.8 | 53 | 640.2 | 521.4 | 426.6 | 538.5 | 299.9 | 275.6 |
| Q8BWT1 | ***Acaa2*** | 4749.3 | 7273.9 | 7654.9 | 1131.8 | 576 | 803.8 | 864.7 | 689.3 | 811 |
| Q60930 | ***Vdac2*** | 2990.7 | 1712.7 | 914.6 | 8061.6 | 12264.9 | 11461 | 9067.6 | 7344 | 8298.4 |
| Q9DC70 | ***Ndufs7*** | 559.6 | 426.7 | 245.3 | 1100.8 | 1325.6 | 1060.3 | 985 | 746.7 | 698.9 |
| Q91V41 | ***Rab14*** | 1550.2 | 911.6 | 539.5 | 3315.6 | 2977.6 | 2791.2 | 3093.4 | 1814.1 | 1816.8 |
| Q921I1 | ***Tf*** | 392.7 | 391.9 | 322.9 | 1065.5 | 1621.9 | 1405.1 | 955.1 | 1125.3 | 1013.6 |

| | | | | | | | | | | |
|---|---|---|---|---|---|---|---|---|---|---|
| Q9WUR2 | ***Eci2*** | 189.3 | 235.4 | 303.9 | 2406.2 | 4042.2 | 3945.8 | 6076.9 | 30498.2 | 12887.6 |
| P13020 | ***Gsn*** | 99 | 74.5 | 41.8 | 225.7 | 205.1 | 177.8 | 155.4 | 101.5 | 79.1 |
| Q8VCR2 | ***Hsd17b13*** | 231.9 | 125.4 | 100.3 | 603.6 | 467 | 475 | 594.6 | 286.5 | 352.1 |
| Q8BKZ9 | ***Pdhx*** | 1883.6 | 1074.9 | 650.9 | 4930.2 | 7569.1 | 6399.8 | 8720.5 | 6969.3 | 7990.9 |
| G5E829 | ***Atp2b1*** | 2058.4 | 766.3 | 603.2 | 3960.9 | 4875.7 | 4238.3 | 3559 | 2306.8 | 2439.3 |
| Q91W43 | ***Gldc*** | 530.7 | 683.9 | 615.5 | 288 | 101 | 142.2 | 199.7 | 153.6 | 136.5 |
| P21107 | ***Tpm3*** | 8022.1 | 4128.1 | 2887.7 | 13941.1 | 15211.4 | 14512.4 | 14784.8 | 7581.8 | 7343.7 |
| P26040 | ***Ezr*** | 736.1 | 534.1 | 576.1 | 214.5 | 166.8 | 258.9 | 184.3 | 157 | 211 |
| Q9QYA2 | ***Tomm40*** | 746.9 | 595.9 | 983.3 | 135.7 | 58.2 | 55.9 | 150.4 | 69.6 | 80.7 |
| P19536 | ***Cox5b*** | 3011.9 | 1215.6 | 909.2 | 5518.4 | 5871.2 | 5622.9 | 5524.8 | 3635.5 | 3977 |
| Q9CXN7 | ***Pbld2*** | 8650.6 | 12087.3 | 11168.6 | 4323.8 | 1488.9 | 2010.7 | 2932.7 | 1830.7 | 1694.3 |
| Q9Z0X1 | ***Aifm1*** | 2508.8 | 2489.5 | 3989.4 | 29.2 | 34.3 | 41.8 | 52 | 23.7 | 49.4 |
| Q9Z1S5 | ***42981*** | 231.2 | 157.2 | 97.8 | 948.5 | 828 | 609.7 | 889.5 | 480.6 | 497.4 |
| P84091 | ***Ap2m1*** | 63.8 | 35.3 | 15.1 | 164.3 | 127.9 | 137.7 | 220.1 | 126.5 | 144.3 |
| P58281 | ***Opa1*** | 2475.5 | 1800.7 | 2336.2 | 952 | 439 | 734.9 | 868.6 | 477.3 | 664.3 |
| P46460 | ***Nsf*** | 1303.8 | 814.7 | 739.5 | 12602 | 8279.9 | 8113.1 | 10833.7 | 4218.4 | 5125.9 |
| Q922Q8 | ***Lrrc59*** | 643.6 | 1169.4 | 927.6 | 21.4 | 25.8 | 11.3 | 6 | 4 | 8.2 |
| Q99K51 | ***Pls3*** | 37 | 110.2 | 133 | 331.8 | 482.2 | 388.6 | 511 | 2491.7 | 1260.9 |
| Q8R1S0 | ***Coq6*** | 760.7 | 1279.9 | 1271.5 | 93.1 | 124 | 86.3 | 102.9 | 325.4 | 176.7 |
| P80314 | ***Cct2*** | 888.8 | 1125.1 | 1026.1 | 541.6 | 265.9 | 409.8 | 426.2 | 353.6 | 348 |
| Q64105 | ***Spr*** | 625.2 | 1114.6 | 839.2 | 51.2 | 43.1 | 40.5 | 143.5 | 63 | 64.9 |
| O08709 | ***Prdx6*** | 883.9 | 1645.5 | 1267.7 | 7.5 | 18.8 | 9.2 | 15.9 | 4 | 14.4 |
| P41216 | ***Acsl1*** | 4729.3 | 6759.2 | 3789.4 | 80.2 | 137 | 139.4 | 169.1 | 228.9 | 269.3 |
| UPSP:K1CX_HUMAN | | 2357.8 | 974.8 | 675.8 | 4834.1 | 5309.7 | 4192.5 | 4274.5 | 3436.1 | 2307.6 |
| Q9DD20 | ***Mettl7b*** | 1773.2 | 2658.6 | 3291.2 | 109.3 | 90.9 | 81 | 72.9 | 63.5 | 94 |
| Q3UJU9 | ***Rmdn3*** | 1965.7 | 592.8 | 577 | 3827.9 | 3633.6 | 3558.3 | 3113 | 2194.6 | 2325.4 |
| Q7TSJ2 | ***Map6*** | 305.4 | 201.6 | 157.9 | 2039.7 | 3424.5 | 3622 | 1771 | 1954.6 | 2566.6 |
| P68254 | ***Ywhaq*** | 165.3 | 162.6 | 140.7 | 4124.2 | 7335.5 | 5026.2 | 4651.6 | 6421.1 | 5063.2 |
| Q08331 | ***Calb2*** | 24.9 | 20.2 | 48.4 | 262.7 | 453.4 | 325.4 | 240.6 | 238.9 | 220.1 |
| P68372 | ***Tubb4b*** | 20.1 | 21.4 | 22.3 | 612.8 | 1153.1 | 1041.8 | 1575.4 | 6019.6 | 2730.5 |
| Q9CQA3 | ***Sdhb*** | 1281.2 | 902.3 | 981.9 | 389.3 | 233.4 | 410.3 | 302.2 | 254.4 | 308.8 |
| Q8VEB4 | ***Pla2g15*** | 28.5 | 37.4 | 70.7 | 404.8 | 732.4 | 590.7 | 967.8 | 2175.6 | 1832.7 |
| P70404 | ***Idh3g*** | 35.5 | 146.7 | 158 | 375.5 | 318.6 | 348.2 | 445.6 | 744.6 | 636.1 |
| Q80ZS3 | ***Mrps26*** | 0 | 43.1 | 34.8 | 808.8 | 1389.3 | 885.1 | 1027.6 | 3146.4 | 2138.2 |
| Q91WG0 | ***Ces2c*** | 553.8 | 374.4 | 417.2 | 107.9 | 86.2 | 170.9 | 55.1 | 88.8 | 126.3 |
| Q9QZW0 | ***Atp11c*** | 275.9 | 213.4 | 149.5 | 9.1 | 11.8 | 20.5 | 17.3 | 10.3 | 44.8 |
| Q9JKD3 | ***Scamp5*** | 3.1 | 1.8 | 10 | 1000.9 | 1582.9 | 897.3 | 834.9 | 1145.2 | 702 |
| Q9D273 | ***Mmab*** | 2946.1 | 4812.8 | 2965.8 | 296.4 | 217.5 | 187.4 | 216.2 | 125 | 135.5 |
| P42125 | ***Eci1*** | 438.9 | 666.3 | 705.9 | 184.4 | 85.2 | 117.4 | 112.8 | 100.8 | 123.1 |
| P08113 | ***Hsp90b1*** | 1808.7 | 1095.9 | 656.3 | 3689.1 | 5318.5 | 4111.1 | 3387.7 | 2682.8 | 2386.4 |
| Q3UUI3 | ***Them4*** | 28.9 | 88.8 | 87.9 | 388.4 | 690.1 | 599.4 | 508.6 | 1315.5 | 1233.7 |
| Q60932 | ***Vdac1*** | 231.7 | 1355.7 | 1801.7 | 4286.3 | 6660 | 5720.8 | 5674.6 | 14687.9 | 11371.7 |
| P51150 | ***Rab7a*** | 25.1 | 14.3 | 19.9 | 85 | 128.8 | 149.2 | 182.6 | 558.5 | 589.9 |
| P53026 | ***Rpl10a*** | 321.5 | 227 | 258.9 | 133.9 | 90.5 | 104.6 | 138.4 | 66.4 | 97 |
| Q8VEM8 | ***Slc25a3*** | 208.5 | 168.5 | 140.1 | 303.7 | 382.3 | 408.5 | 318.9 | 219.3 | 242.1 |
| Q7TNG8 | ***Ldhd*** | 1511 | 2824.9 | 2948.6 | 58.3 | 57.1 | 38.6 | 45.4 | 43.7 | 56.3 |

| | | | | | | | | | | |
|---|---|---|---|---|---|---|---|---|---|---|
| Q9D855 | ***Uqcrb*** | 569.4 | 1054.8 | 1093.3 | 37.3 | 45.8 | 28.3 | 28.1 | 5.2 | 30.2 |
| Q63880 | ***Ces3a*** | 10.7 | 101.4 | 209.3 | 490.2 | 787.6 | 769.1 | 658 | 1671.6 | 2075.9 |
| P24270 | ***Cat*** | 827.3 | 913.9 | 1455 | 57.2 | 66.8 | 47.1 | 42.8 | 39.6 | 53.5 |
| Q5FW60 | ***Mup20*** | 4050.6 | 5622.8 | 7746.1 | 322.1 | 273.8 | 340.8 | 272.9 | 166.2 | 265.1 |
| Q9CPQ1 | ***Cox6c*** | 776.4 | 1450.1 | 1137.6 | 146.4 | 92.9 | 122.9 | 134.1 | 70.6 | 83.6 |
| P50429 | ***Arsb*** | 307.2 | 375.7 | 571.3 | 16.3 | 16.4 | 9 | 11.7 | 12.3 | 10.1 |
| P97427 | ***Crmp1*** | 877.3 | 1438.5 | 1766.6 | 46.6 | 37.3 | 31.6 | 35 | 12.9 | 27.6 |
| P06330 | | 24.2 | 81.1 | 126.5 | 245.1 | 338.8 | 270 | 316.7 | 475.9 | 358.5 |
| Q8C196 | ***Cps1*** | 7483.4 | 11152.1 | 13305.2 | 2368.3 | 1260.6 | 1977.3 | 1556.8 | 1536.8 | 1757.2 |
| P47802 | ***Mtx1*** | 336.2 | 637.4 | 649 | 23.6 | 17.4 | 34.2 | 23.9 | 18.1 | 29.3 |
| P01027 | ***C3*** | 3012.6 | 2094.4 | 1431.9 | 7026.7 | 8414.6 | 11300.5 | 6558 | 5406.3 | 5554.2 |
| Q64516 | ***Gk*** | 743.4 | 459.3 | 266.1 | 2680.9 | 1784.4 | 1842.3 | 2157.4 | 1172.6 | 1025 |
| Q9DBM2 | ***Ehhadh*** | 81.2 | 226.8 | 264.1 | 636.1 | 477.6 | 543.4 | 669.4 | 1381.2 | 993.4 |
| P07901 | ***Hsp90aa1*** | 96.3 | 404.8 | 756.2 | 1448.8 | 1899.3 | 1485.6 | 2042.5 | 4002 | 2867.5 |
| Q99JI6 | ***Rap1b*** | 0 | 3.5 | 0 | 47.4 | 97.9 | 82.9 | 149.1 | 305.3 | 244.6 |
| P12970 | ***Rpl7a*** | 1389.6 | 2274.9 | 2852.3 | 56.4 | 55 | 47.4 | 78.8 | 50 | 82 |
| Q8BH04 | ***Pck2*** | 4.4 | 2.3 | 4.8 | 29 | 44.4 | 25.3 | 54.1 | 239.9 | 116 |
| Q9D710 | ***Tmx2*** | 627.5 | 265.5 | 180.6 | 1005.7 | 1105.3 | 1032.9 | 979.8 | 649.3 | 662.9 |
| Q99LC5 | ***Etfa*** | 3772.1 | 2117.1 | 1082.1 | 6120.8 | 6075.3 | 6349.8 | 9295 | 4085.3 | 4601.6 |
| P26043 | ***Rdx*** | 378.7 | 195.4 | 113 | 1703.2 | 1102.4 | 1083.6 | 1304.2 | 659.6 | 647.5 |
| P60879 | ***Snap25*** | 3172.8 | 6565.8 | 6093.8 | 64.8 | 114.2 | 81.9 | 95.9 | 103.8 | 138.1 |
| Q7TMM9 | ***Tubb2a*** | 523.8 | 331.9 | 175.8 | 2130.9 | 1495.5 | 1316 | 1727.5 | 950.7 | 723.9 |
| Q9Z1G4 | ***Atp6v0a1*** | 290.2 | 197.9 | 138.5 | 514.6 | 802.4 | 745 | 507.9 | 390.2 | 525.6 |
| O08553 | ***Dpysl2*** | 338.1 | 175.5 | 107.6 | 771.2 | 641.5 | 556.6 | 529.1 | 279.2 | 337.8 |
| Q8CAK1 | ***Iba57*** | 120.8 | 431.7 | 1096.9 | 2163.4 | 2924.2 | 2152.2 | 3475.6 | 10906.4 | 6329.7 |
| P54071 | ***Idh2*** | 598.4 | 1205.2 | 832.2 | 24.4 | 24.3 | 26.5 | 28.3 | 30.6 | 33.1 |
| Q61233 | ***Lcp1*** | 1258.1 | 2275.8 | 2678.3 | 48.8 | 42.7 | 47.4 | 61.4 | 34.2 | 45.1 |
| Q9CZR8 | ***Tsfm*** | 13.6 | 40.2 | 24.8 | 769 | 1497.9 | 943 | 1341.2 | 4764.3 | 3028.8 |
| Q925I1 | ***Atad3*** | 183.1 | 139.4 | 146.3 | 1772.2 | 3323.5 | 2098.4 | 2535.1 | 2390.9 | 2808.2 |
| Q8BFP9 | ***Pdk1*** | 528 | 297.5 | 211.4 | 1473.8 | 2486.6 | 1677 | 1275 | 1160.7 | 1056.5 |
| Q922B1 | ***Macrod1*** | 9374 | 6029.5 | 12798.2 | 177.1 | 310.6 | 126.6 | 159.5 | 371.2 | 368.1 |
| Q91WC3 | ***Acsl6*** | 5789.7 | 6270 | 10786 | 172.5 | 115.9 | 114.2 | 223.7 | 100.3 | 207.1 |
| Q60872 | ***Eif1a*** | 57.1 | 40 | 54.2 | 121.2 | 168.6 | 202.7 | 212.3 | 346.3 | 240 |
| P11438 | ***Lamp1*** | 26.1 | 16.9 | 23.9 | 421.8 | 263.4 | 225.6 | 584.5 | 1080 | 690.1 |
| Q9QYR6 | ***Map1a*** | 162.5 | 253.8 | 242.9 | 87.1 | 84.2 | 83.6 | 96.8 | 57.1 | 66.1 |
| P63044 | ***Vamp2*** | 3845.5 | 1545.3 | 967.2 | 8639.6 | 6399 | 6887.9 | 7845.6 | 3989.1 | 4181.4 |
| Q9ET22 | ***Dpp7*** | 486.2 | 925.8 | 1015.4 | 47.5 | 37.6 | 56.8 | 69.3 | 33.6 | 52.5 |
| Q8BHN3 | ***Ganab*** | 1397.8 | 520.7 | 295.5 | 2725.9 | 2152.8 | 2626.9 | 2065.8 | 1114.4 | 1173 |
| Q922U2 | ***Krt5*** | 721.8 | 1454 | 1556.9 | 24.8 | 36.9 | 13.4 | 19.1 | 12.8 | 29.9 |
| P10107 | ***Anxa1*** | 91 | 233.9 | 300.4 | 512.8 | 518.2 | 649.1 | 508.9 | 705.8 | 617 |
| P61922 | ***Abat*** | 182.3 | 115.8 | 84.7 | 301.6 | 252 | 272.4 | 270.2 | 135.9 | 162.6 |
| P24369 | ***Ppib*** | 128.7 | 504.7 | 919.7 | 1792.1 | 2451.3 | 1736.4 | 2087 | 3638.3 | 2812.4 |
| Q91YI0 | ***Asl*** | 1874.5 | 3072.3 | 3598.7 | 513.7 | 546.8 | 513 | 451.7 | 428.1 | 456.1 |
| O08997 | ***Atox1*** | 569.1 | 331.9 | 272.6 | 875.4 | 863.2 | 763.5 | 890.7 | 494.5 | 464.3 |
| P40936 | ***Inmt*** | 74.3 | 250.6 | 389.6 | 626.4 | 813.6 | 711.6 | 722.8 | 1428.2 | 1032.6 |
| P16858 | ***Gapdh*** | 1186.1 | 1001.3 | 864.5 | 550.4 | 305.5 | 542.5 | 305.4 | 271 | 364.2 |
| Q9D6R2 | ***Idh3a*** | 99.1 | 67.1 | 68.7 | 224.8 | 305.7 | 400.4 | 680.5 | 1866.4 | 1570.4 |

| | | | | | | | | | | |
|---|---|---|---|---|---|---|---|---|---|---|
| P11588 | ***Mup1*** | 2430.2 | 4578 | 4529.1 | 606.6 | 756.2 | 656 | 538.4 | 415.4 | 528.6 |
| O08917 | ***Flot1*** | 397.1 | 865 | 831.8 | 26.1 | 20.1 | 28.4 | 27.6 | 23.8 | 26.2 |
| Q9CPZ8 | ***Cmc1*** | 672.6 | 515 | 631.1 | 391.4 | 379.5 | 407.5 | 415 | 330.5 | 376.3 |
| Q9D8T7 | ***Slirp*** | 1017.8 | 433.1 | 385.5 | 2384.6 | 4220 | 2958.7 | 2414.6 | 1974.1 | 2179.6 |
| Q9WTP7 | ***Ak3*** | 4.7 | 20.6 | 12.6 | 242.8 | 519.2 | 349.1 | 329.4 | 438.6 | 564 |
| Q3UMR5 | ***Mcu*** | 67.6 | 53.8 | 36.7 | 108 | 88.3 | 107.4 | 128.6 | 86.1 | 89.6 |
| Q9Z2Z6 | ***Slc25a20*** | 191.2 | 80.3 | 101.8 | 375.7 | 271.4 | 320.9 | 216 | 99.6 | 313.7 |
| P08905 | ***Lyz2*** | 255.4 | 118.2 | 79 | 1111.7 | 706.2 | 651.2 | 556.4 | 297.8 | 275.3 |
| Q99JR1 | ***Sfxn1*** | 1554.6 | 2827.3 | 1550.7 | 76.9 | 163.9 | 151.4 | 164 | 238.6 | 250.7 |
| P45591 | ***Cfl2*** | 459.2 | 273.1 | 238.5 | 935.6 | 705.9 | 689.6 | 1127.1 | 532.4 | 617 |
| Q9DD18 | ***Dtd1*** | 220.1 | 168.5 | 136.7 | 315.5 | 440.3 | 347.4 | 271.1 | 261.8 | 243.3 |
| Q6P3A8 | ***Bckdhb*** | 186.4 | 738 | 1151.6 | 2803.6 | 1921.1 | 2240 | 2685.6 | 3999.4 | 4060.4 |
| Q8BFR5 | ***Tufm*** | 9.2 | 42.5 | 64.6 | 1036.9 | 1678.3 | 793.9 | 725.9 | 2651.6 | 1335.2 |
| P28474 | ***Adh5*** | 4258.9 | 1928.3 | 1252.6 | 7981.8 | 7392.8 | 5992.1 | 7554 | 4273.3 | 3790.6 |
| P70441 | ***Slc9a3r1*** | 795.4 | 1719.3 | 1782.9 | 52.5 | 79.4 | 80.7 | 20.7 | 11.5 | 128.3 |
| P17717 | ***Ugt2b17*** | 418.9 | 206.3 | 224.9 | 600 | 886 | 730.4 | 822.6 | 2028 | 1600.3 |
| P47934 | ***Crat*** | 912.7 | 1875.5 | 1241.2 | 121.3 | 162 | 144 | 128.4 | 163.8 | 159.1 |
| Q99KI0 | ***Aco2*** | 1810.3 | 2028.4 | 3389 | 427.6 | 264.4 | 220.4 | 228.3 | 200.7 | 152.2 |
| Q9DBG3 | ***Ap2b1*** | 88.9 | 61.1 | 36.2 | 133.1 | 156.2 | 124.1 | 311.2 | 170.1 | 200.1 |
| Q9DB73 | ***Cyb5r1*** | 511.5 | 388.4 | 308.9 | 134.4 | 95.6 | 178.6 | 109.2 | 109 | 142.7 |
| P04919 | ***Slc4a1*** | 185.7 | 1225.4 | 1195 | 2719.4 | 4947.9 | 4531.4 | 5806.5 | 13393.8 | 11647 |
| Q8CAQ8 | ***Immt*** | 108.9 | 399.7 | 473.9 | 756.2 | 1001.5 | 1077.8 | 716.4 | 2138 | 2177.5 |
| P54116 | ***Stom*** | 519.5 | 947.8 | 1227.6 | 49 | 44.2 | 28.9 | 32 | 37.2 | 42.3 |
| P08003 | ***Pdia4*** | 27.3 | 110.4 | 95.9 | 1259.3 | 2387 | 3035.9 | 2199.8 | 5779.9 | 5400.9 |
| P62814 | ***Atp6v1b2*** | 249.3 | 136.9 | 126.4 | 321.6 | 414.9 | 366.8 | 347.9 | 244.7 | 263 |
| P19246 | ***Nefh*** | 25.1 | 44.7 | 42.3 | 121.1 | 240.2 | 234.5 | 216.8 | 606 | 622.3 |
| Q8BJ64 | ***Chdh*** | 3295 | 1670.1 | 932.8 | 5660.4 | 4937.2 | 4640.6 | 5109 | 3376.3 | 3007.5 |
| P28843 | ***Dpp4*** | 1662 | 1175.2 | 890.6 | 322.4 | 330.2 | 319.3 | 428.1 | 213.3 | 366 |
| Q9CRB8 | ***Mtfp1*** | 1211.2 | 3024.4 | 2650.2 | 21.9 | 61.8 | 50.6 | 58 | 49.2 | 61.8 |
| P00329 | ***Adh1*** | 6545.6 | 5023.4 | 3651.4 | 1396.5 | 2045.2 | 1297.4 | 2149.4 | 1423.4 | 1798.6 |
| P61804 | ***Dad1*** | 2436.4 | 1155.4 | 525.6 | 4977.1 | 3460.1 | 4253.6 | 4728 | 2440.7 | 2473.6 |
| P43274 | ***Hist1h1e*** | 592.1 | 637.3 | 1181 | 39.1 | 40.7 | 54.4 | 63.8 | 127.6 | 88.9 |
| O88441 | ***Mtx2*** | 716.3 | 1162.9 | 1451 | 295.8 | 167.4 | 262.8 | 194.1 | 151.8 | 197.6 |
| O88587 | ***Comt*** | 288.9 | 345.5 | 204.2 | 136.9 | 60.4 | 74.9 | 102.4 | 86.7 | 87.5 |
| Q9D0G0 | ***Mrps30*** | 607.1 | 1131.3 | 842.7 | 331.3 | 73.7 | 146.4 | 130.6 | 26.3 | 74.4 |
| Q8BMS4 | ***Coq3*** | 648.1 | 403.1 | 243.2 | 1903.7 | 1289.1 | 1165.4 | 1832.9 | 863.2 | 872.4 |
| Q99KB8 | ***Hagh*** | 809.2 | 443.3 | 350.8 | 1124.6 | 1197.9 | 1032.3 | 1799.2 | 823.9 | 1131.1 |
| Q64374 | ***Rgn*** | 8.5 | 40.4 | 54.2 | 108.6 | 206.6 | 208.6 | 197.3 | 517.8 | 490.8 |
| Q9DCN2 | ***Cyb5r3*** | 1392.1 | 690.1 | 643.2 | 1919.9 | 2886.8 | 3132.5 | 1861.8 | 2246.8 | 2568.1 |
| Q8CHT0 | ***Aldh4a1*** | 179.5 | 1372.5 | 2412.6 | 3928.5 | 6967.2 | 6148.8 | 5216.3 | 14822.1 | 11908.9 |
| Q91V61 | ***Sfxn3*** | 66.2 | 59.8 | 183.7 | 1117 | 2244 | 2820.9 | 2260.3 | 4657.9 | 5392.5 |
| O35143 | ***Atpif1*** | 987 | 1011.7 | 456.5 | 1731 | 2229.5 | 1591.8 | 2952.5 | 2908 | 2323.1 |
| P62874 | ***Gnb1*** | 7599.2 | 5902 | 3041.7 | 295.5 | 465.1 | 262.3 | 426.7 | 300.9 | 410.9 |
| Q80TB8 | ***Vat1l*** | 91.3 | 67.9 | 36.4 | 158.9 | 273.4 | 196.4 | 115.7 | 172.5 | 150.3 |
| Q9JHI5 | ***Ivd*** | 236.4 | 850.6 | 1152.1 | 1696.5 | 2167 | 2495.4 | 2212.1 | 4604.9 | 4723.3 |
| P56379 | ***Mp68*** | 2015 | 1159.4 | 869.6 | 42.5 | 21 | 10.6 | 23.9 | 7.7 | 20.2 |
| O88935 | ***Syn1*** | 360.4 | 573.7 | 878.4 | 18.2 | 25.5 | 41.7 | 27.7 | 44 | 44.5 |

| | | | | | | | | | | |
|---|---|---|---|---|---|---|---|---|---|---|
| P10854 | ***Hist1h2bm*** | 495.1 | 241.8 | 177.7 | 775.1 | 1464.9 | 1394.3 | 803.7 | 1129.1 | 1252.3 |
| P17742 | ***Ppia*** | 136.7 | 590.8 | 1083.1 | 1630.3 | 2027 | 1613 | 2381.3 | 4434.4 | 3210.9 |
| P16546 | ***Sptan1*** | 552.8 | 822.8 | 1233.6 | 114.7 | 117.6 | 146.9 | 104.1 | 160.3 | 150.3 |
| P12023 | ***App*** | 136.4 | 81 | 54.3 | 587.1 | 343.6 | 326.3 | 493.9 | 181 | 249 |
| Q8BWQ1 | ***Ugt2a3*** | 119 | 300.7 | 334.9 | 1516 | 3321.4 | 2008.2 | 3957.5 | 7553.8 | 7897.3 |
| Q9CQV8 | ***Ywhab*** | 4216.2 | 8729.2 | 4642.4 | 493.4 | 613.8 | 334.6 | 449.6 | 319.7 | 334 |
| P00920 | ***Ca2*** | 10.3 | 39.2 | 37.9 | 148.1 | 188 | 311.4 | 184.9 | 484.6 | 591.9 |
| Q3TC72 | ***Fahd2*** | 1632.8 | 1019.5 | 936.5 | 376.5 | 410.8 | 337.6 | 348 | 171 | 188.9 |
| Q61102 | ***Abcb7*** | 2856.4 | 1723.9 | 1155 | 71.9 | 81.9 | 71.1 | 76.4 | 18 | 71.4 |
| P49935 | ***Ctsh*** | 423.7 | 233.5 | 251.2 | 88.7 | 85.2 | 58.8 | 117.2 | 69.1 | 81.6 |
| O35405 | ***Pld3*** | 14.7 | 21.5 | 68.3 | 231.4 | 571.9 | 397.2 | 372.3 | 1076 | 825.7 |
| Q9CZS1 | ***Aldh1b1*** | 2297.3 | 6369.8 | 5732.1 | 213.1 | 175.8 | 185.1 | 321.4 | 184.2 | 251.2 |
| P28663 | ***Napb*** | 70.8 | 176.7 | 203.2 | 2543.3 | 6238.2 | 3564.9 | 4690.6 | 18529 | 11302.3 |
| Q8BIJ6 | ***Iars2*** | 4560.5 | 2727.5 | 2078.9 | 393.2 | 519.9 | 437.2 | 702.9 | 444.5 | 732.8 |
| Q7TQD2 | ***Tppp*** | 1646.5 | 2420.1 | 4104.6 | 100 | 184.9 | 81.5 | 86.3 | 52.3 | 89.7 |
| Q9CRB6 | ***Tppp3*** | 542.7 | 308.7 | 202.1 | 1196.1 | 2384.4 | 1473.8 | 1162 | 1253.1 | 1135.8 |
| Q9CRB9 | ***Chchd3*** | 9622.6 | 15089 | 20039.7 | 5535 | 2325.8 | 2917.3 | 3368.3 | 2687.8 | 2988.1 |
| Q9JKC6 | ***Cend1*** | 16.1 | 96 | 126.7 | 675.1 | 1456.1 | 765.6 | 1074.4 | 6136.7 | 2292.8 |
| P14206 | ***Rpsa*** | 2325.7 | 1478.3 | 871.8 | 52.1 | 56.2 | 66.4 | 72.9 | 37.3 | 43.7 |
| Q9WV54 | ***Asah1*** | 2339.8 | 1481.8 | 877 | 69.8 | 39.1 | 58.4 | 54.8 | 49.3 | 66.9 |
| P48320 | ***Gad2*** | 184.3 | 255.4 | 263.8 | 362.4 | 412.3 | 320.1 | 548.3 | 282.8 | 411.3 |
| P99024 | ***Tubb5*** | 78.8 | 291.2 | 739.9 | 1057.5 | 1084.9 | 1040.3 | 1211.4 | 2026.8 | 1720.4 |
| Q6IRU5 | ***Cltb*** | 49.6 | 172.6 | 319.7 | 477.5 | 944.1 | 808.6 | 1020.7 | 3669.8 | 2185.3 |
| Q3UV17 | ***Krt76*** | 720.2 | 694.5 | 344.8 | 203.4 | 80.1 | 143.4 | 124.5 | 93.2 | 118.4 |
| P40142 | ***Tkt*** | 48.1 | 75.7 | 114.3 | 230.5 | 143 | 205.6 | 312.7 | 416.2 | 404.8 |
| Q9CWZ7 | ***Napg*** | 227.7 | 368.3 | 277.3 | 151.2 | 144 | 144.1 | 142.3 | 196.9 | 204.5 |
| P32020 | ***Scp2*** | 23.5 | 50.6 | 60.3 | 158.8 | 395 | 319.4 | 505.1 | 2980.8 | 1535.6 |
| Q9QXD6 | ***Fbp1*** | 1097.1 | 1855.8 | 2339.7 | 594.4 | 349.4 | 493.9 | 369 | 365.5 | 422.2 |
| P01899 | ***H2-D1*** | 177.6 | 259.8 | 362.6 | 89.8 | 70.2 | 77.9 | 88.1 | 119.1 | 101.5 |
| Q9JLJ2 | ***Aldh9a1*** | 7335.2 | 3881.5 | 3012.8 | 207.4 | 171.5 | 188 | 262.5 | 105 | 202.7 |
| P47857 | ***Pfkm*** | 738.8 | 484.1 | 429.2 | 211.2 | 133 | 256.1 | 181.5 | 176.8 | 191.9 |
| P43276 | ***Hist1h1b*** | 565.2 | 1349.4 | 1636 | 76.9 | 76.3 | 99.2 | 73.3 | 83.6 | 125.5 |
| Q9CZP5 | ***Bcs1l*** | 35.9 | 63 | 128.1 | 254 | 587.1 | 413.9 | 406.7 | 1029.9 | 904.1 |
| O55143 | ***Atp2a2*** | 1399.9 | 4309.5 | 3563.8 | 135.2 | 100.5 | 86 | 116.4 | 23.2 | 110.8 |
| P08226 | ***Apoe*** | 0 | 46.9 | 67.2 | 108.7 | 228.6 | 213.1 | 169.9 | 486.4 | 453.4 |
| P56135 | ***Atp5j2*** | 288 | 894.2 | 801.7 | 20.5 | 20.1 | 15.8 | 14.1 | 9.6 | 19.3 |
| P47963 | ***Rpl13*** | 2679.4 | 1730.5 | 1327.2 | 5136.2 | 3823.2 | 3433.2 | 6390.3 | 2550.9 | 3023 |
| P61161 | ***Actr2*** | 1004.9 | 818.8 | 436.9 | 150.7 | 235.4 | 146.1 | 238.2 | 345.2 | 334.7 |
| Q8VCT4 | ***Ces1d*** | 263.7 | 649.5 | 685.3 | 38.5 | 37.9 | 122.1 | 49.1 | 25.4 | 80.5 |
| P06837 | ***Gap43*** | 1587.9 | 1124.5 | 507 | 3994.1 | 9346.5 | 5991.7 | 3794.7 | 6939.7 | 5909.5 |
| O35658 | ***C1qbp*** | 757.9 | 708.8 | 454.8 | 229.3 | 381.9 | 241.3 | 318 | 273.2 | 259.5 |
| P08752 | ***Gnai2*** | 1292.4 | 3859.9 | 2499.8 | 51.4 | 74.5 | 56.5 | 85.7 | 27.4 | 53 |
| O35887 | ***Calu*** | 259.9 | 149.9 | 85.8 | 354 | 331.3 | 327.7 | 453.4 | 235.4 | 261.7 |
| Q9DCS9 | ***Ndufb10*** | 162.5 | 99.7 | 196.3 | 63.3 | 39.5 | 62 | 43.4 | 61.9 | 59.4 |
| P59017 | ***Bcl2l13*** | 55.8 | 71.6 | 66.2 | 678.2 | 1466.9 | 2072 | 1451.4 | 3809 | 4260.2 |
| P14148 | ***Rpl7*** | 965 | 1202.1 | 535.5 | 338.4 | 170.7 | 197.7 | 217.4 | 173.5 | 196.1 |
| Q8BUV3 | ***Gphn*** | 917.4 | 654.6 | 422.8 | 141.8 | 249.6 | 153.1 | 192.1 | 227.4 | 234.1 |

| | | | | | | | | | | |
|---|---|---|---|---|---|---|---|---|---|---|
| Q63886 | ***Ugt1a1*** | 346.4 | 218.6 | 140.9 | 572.9 | 1059.1 | 659.7 | 544.5 | 458.6 | 473.6 |
| O08599 | ***Stxbp1*** | 133.5 | 628.4 | 777.1 | 1562.2 | 2976.4 | 1751.3 | 1325.9 | 5349.9 | 2453.7 |
| Q91VT4 | ***Cbr4*** | 161.4 | 69.7 | 58.9 | 205.5 | 415.6 | 364.1 | 371.6 | 330.3 | 361.1 |
| Q9R0P5 | ***Dstn*** | 893.7 | 2510.9 | 2505.8 | 294.4 | 119.1 | 220.3 | 101.5 | 43 | 82.6 |
| UPSP:K2CE_HUMAN | | 716.3 | 762.1 | 394 | 302 | 134.1 | 210.9 | 162.6 | 100.6 | 127.4 |
| P60766 | ***Cdc42*** | 162.8 | 470 | 293.3 | 29.2 | 13.4 | 21.6 | 12.5 | 37.2 | 19.1 |
| Q9DBT9 | ***Dmgdh*** | 609.6 | 1700.1 | 1349 | 123.2 | 107.7 | 276.8 | 126.8 | 60.9 | 208.5 |
| Q64332 | ***Syn2*** | 603.5 | 367.4 | 173.1 | 877.8 | 826.6 | 721.6 | 976.8 | 569.5 | 572 |
| Q8QZT1 | ***Acat1*** | 2788.7 | 1914.6 | 906.1 | 134.7 | 145.2 | 91.5 | 137.7 | 65.1 | 96 |
| Q9D0M3 | ***Cyc1*** | 13691.5 | 8061.4 | 4442.8 | 166.4 | 201.6 | 193.5 | 126.1 | 171.5 | 317.1 |
| P28661 | ***42982*** | 33142.2 | 19572.4 | 12058.4 | 1732.3 | 2420.7 | 2052.2 | 2918.2 | 1413 | 2449.7 |
| B5X0G2 | ***Mup17*** | 1574.4 | 1206.9 | 767.8 | 400.1 | 508.4 | 408.1 | 610.3 | 415.5 | 560.7 |
| P17932 | ***Rpl32-ps*** | 3098.5 | 1491.4 | 997.3 | 3893.2 | 8302.6 | 7259.3 | 4741.4 | 6343.9 | 6628.1 |
| Q8BGD8 | ***Coa6*** | 127.6 | 87.7 | 51.5 | 173.2 | 162.7 | 148.4 | 176.1 | 148.1 | 127.1 |
| P26039 | ***Tln1*** | 9748.5 | 4620 | 2698.5 | 25631 | 15403.2 | 14686.3 | 22998.6 | 11102.9 | 9614.9 |
| Q8BGA8 | ***Acsm5*** | 3699.3 | 2431.4 | 1361.9 | 404.6 | 402.3 | 335.1 | 518.9 | 285.1 | 364.4 |
| Q91ZA3 | ***Pcca*** | 893.7 | 3193 | 2745.3 | 86.7 | 92.9 | 63.9 | 174 | 177.7 | 181.7 |
| Q9CZL5 | ***Pcbd2*** | 10972 | 5708 | 3813.9 | 176.6 | 204.5 | 132.8 | 196.5 | 89.1 | 206.2 |
| Q6Q477 | ***Atp2b4*** | 417.8 | 546.9 | 849.9 | 126.2 | 207.3 | 245.1 | 187.7 | 332.4 | 289.3 |
| P56394 | ***Cox17*** | 14.3 | 76.8 | 165.4 | 209.1 | 415.4 | 316.8 | 403.4 | 2474.6 | 1085 |
| Q99LP6 | ***Grpel1*** | 1348.8 | 5027.6 | 4256.6 | 110.1 | 104.2 | 50.6 | 73 | 13.7 | 106.7 |
| P34927 | ***Asgr1*** | 1065.1 | 544.7 | 394.2 | 42.1 | 38.6 | 43.4 | 50.9 | 20.9 | 37.1 |
| P12960 | ***Cntn1*** | 566.6 | 2085.9 | 1825.2 | 62.3 | 48.5 | 49.7 | 93 | 81.6 | 91.3 |
| Q8K215 | ***Lyrm4*** | 1223.4 | 1603.2 | 2398 | 730.6 | 617.9 | 703.1 | 1138.4 | 1273.9 | 1363 |
| Q9DB77 | ***Uqcrc2*** | 139.7 | 543 | 814.2 | 1417.2 | 1449.8 | 947.7 | 1560.6 | 3186.4 | 1952.9 |
| Q4LDG0 | ***Slc27a5*** | 224.8 | 814.7 | 667.1 | 31.8 | 32.2 | 27.6 | 66.8 | 54.5 | 53.1 |
| P97823 | ***Lypla1*** | 588.3 | 732.5 | 559.4 | 244.8 | 464.6 | 401.3 | 402.9 | 320.6 | 475 |
| P20060 | ***Hexb*** | 2231.2 | 1107.1 | 784.5 | 66 | 48.9 | 33 | 62.6 | 23.9 | 61.7 |
| Q8VEH3 | ***Arl8a*** | 178.2 | 240.6 | 221 | 162.5 | 47.4 | 62 | 106.2 | 49.6 | 86.4 |
| Q6PB66 | ***Lrpprc*** | 20371.2 | 10769.9 | 6370.6 | 190.4 | 192 | 111.5 | 147.2 | 46.4 | 235.9 |
| Q99LB7 | ***Sardh*** | 7703.7 | 4481.9 | 3338.2 | 8728 | 10055.2 | 8977.2 | 8786.8 | 5167.7 | 6085.7 |
| Q9DBL1 | ***Acadsb*** | 272.2 | 677.4 | 885.4 | 54.5 | 50.2 | 111.8 | 64.7 | 35.4 | 87.2 |
| Q9CQ91 | ***Ndufa3*** | 517.3 | 326.3 | 173 | 27 | 57.9 | 43.1 | 53.5 | 58.8 | 80 |
| Q06890 | ***Clu*** | 172.1 | 455.5 | 597.4 | 42.9 | 31.7 | 45.7 | 56.3 | 39.4 | 49.1 |
| P61164 | ***Actr1a*** | 1810.5 | 1600.7 | 993.6 | 578 | 851.3 | 742.9 | 826.7 | 688.8 | 874.7 |
| P11983 | ***Tcp1*** | 473 | 326.6 | 141.9 | 548 | 866.6 | 1064.7 | 2066.2 | 2066.5 | 3721.4 |
| Q61644 | ***Pacsin1*** | 99.3 | 504.4 | 630.1 | 1084.5 | 1367.2 | 781.3 | 866.4 | 2777.3 | 1335.9 |
| P70245 | ***Ebp*** | 230.7 | 972.2 | 976.5 | 9.1 | 21.4 | 26.1 | 63.9 | 47 | 69.3 |
| O08807 | ***Prdx4*** | 3315 | 2152.3 | 1458.7 | 673.7 | 911.8 | 680.4 | 1188.8 | 699.8 | 978 |
| Q8BP92 | ***Rcn2*** | 1521.1 | 5341.6 | 5667.6 | 613.7 | 106.3 | 411.7 | 374.6 | 90.3 | 470 |
| Q9R013 | ***Ctsf*** | 4319.1 | 3592.2 | 1667.4 | 827.2 | 1088.5 | 937 | 1307.3 | 865.7 | 994.5 |
| P54227 | ***Stmn1*** | 748.3 | 1186.6 | 958.7 | 702.5 | 401 | 477 | 549.4 | 750.2 | 636 |
| Q8BFR4 | ***Gns*** | 2073.8 | 8183.6 | 6919.7 | 746.2 | 211 | 473 | 393.2 | 114.2 | 347.2 |
| Q9JLZ3 | ***Auh*** | 437.8 | 1104.7 | 1640.5 | 83.4 | 78 | 92.8 | 124.1 | 133.1 | 191.5 |
| Q921G7 | ***Etfdh*** | 8428.8 | 4723.3 | 3592.4 | 1551.4 | 1699.7 | 1277.1 | 2346.9 | 1137.6 | 1637.4 |
| P10639 | ***Txn*** | 690.3 | 561.7 | 570.7 | 479.2 | 515.2 | 453.6 | 691.7 | 399.1 | 499.6 |

| | | | | | | | | | | |
|---|---|---|---|---|---|---|---|---|---|---|
| Q9DCW4 | ***Etfb*** | 1657 | 775 | 433.4 | 1733.5 | 2258.1 | 2199.6 | 1495.3 | 1509.6 | 1315 |
| Q9DCM2 | ***Gstk1*** | 267.1 | 1060.4 | 1205.2 | 43.6 | 32.6 | 38.5 | 33.6 | 31.7 | 37.4 |
| Q9R0H0 | ***Acox1*** | 26062.4 | 12898.2 | 10377.5 | 2770.7 | 3459.1 | 2781 | 4560.3 | 2228.8 | 3474.3 |
| Q8K4F5 | ***Abhd11*** | 674.6 | 1896.8 | 1560.5 | 559.1 | 202.3 | 207.5 | 330.1 | 89.1 | 180.4 |
| Q99JY0 | ***Hadhb*** | 1800.6 | 648.2 | 556.2 | 5532 | 3514.8 | 2316.4 | 4282.3 | 1785.3 | 1754.2 |
| Q571F8 | ***Gls2*** | 1684.1 | 815.1 | 582.3 | 122.4 | 119 | 95.3 | 146.9 | 59.6 | 83.7 |
| Q9JMA7 | ***Cyp3a41a; Cyp3a41b*** | 863.6 | 3785.7 | 2816 | 225.6 | 110.5 | 133.9 | 121.7 | 41.2 | 132.4 |
| Q9DB05 | ***Napa*** | 6850.4 | 3946.5 | 2190 | 579.4 | 814.6 | 704.1 | 797 | 549.4 | 780.1 |
| Q62442 | ***Vamp1*** | 3945.6 | 2716.4 | 2033 | 4954.7 | 4741.4 | 4014.2 | 3738.2 | 2366.7 | 2346.5 |
| Q8CC88 | ***Vwa8*** | 3005.5 | 1720.9 | 1177 | 556.3 | 517.4 | 512.9 | 457.2 | 188.5 | 232.9 |
| P50544 | ***Acadvl*** | 318.3 | 356.8 | 224.2 | 146.8 | 223.7 | 165.6 | 254.5 | 203.3 | 234.9 |
| Q9CQ75 | ***Ndufa2*** | 464.6 | 1655.4 | 1645.4 | 250.1 | 109.1 | 265.2 | 169.6 | 63.6 | 217.7 |
| Q01853 | ***Vcp*** | 1753.9 | 1117.9 | 626.4 | 263.7 | 400.7 | 255.3 | 406.7 | 352.8 | 366.3 |
| P00186 | ***Cyp1a2*** | 426.3 | 955.1 | 336 | 46.2 | 111.7 | 45.7 | 88.8 | 42.5 | 44.8 |
| P15626 | ***Gstm2*** | 4155.9 | 3703.8 | 1551.3 | 600.9 | 1342.7 | 1001.7 | 579 | 943.6 | 948.2 |
| Q8BW75 | ***Maob*** | 294.3 | 906 | 784.6 | 246.3 | 110.6 | 142.5 | 154.4 | 45.8 | 101.9 |
| O88986 | ***Gcat*** | 1070.1 | 987.1 | 818.1 | 775.2 | 786 | 720.7 | 644.4 | 584.3 | 525.7 |
| Q62261 | ***Sptbn1*** | 312.9 | 996.6 | 790.1 | 221.5 | 133.6 | 178.2 | 157.4 | 125.8 | 248.4 |
| P97807 | ***Fh*** | 5129.5 | 23429.9 | 27194 | 1246.8 | 1179.2 | 1145.8 | 2420 | 2959.4 | 2830.1 |
| Q9CW42 | ***42795*** | 0 | 4.9 | 3.1 | 61.4 | 202.3 | 81.3 | 124.4 | 549.5 | 349.8 |
| Q9CZU6 | ***Cs*** | 698.9 | 3318.3 | 3711.5 | 151.7 | 96.4 | 247.5 | 145.7 | 27.5 | 227.9 |
| Q9D2G2 | ***Dlst*** | 482.5 | 1320.7 | 2231.3 | 51.3 | 84.6 | 40.8 | 90.8 | 70.8 | 127.3 |
| Q8VDQ8 | ***Sirt2*** | 979.8 | 497.5 | 279.2 | 44.5 | 73.8 | 62.7 | 67.8 | 70.1 | 91.3 |
| Q8VDT9 | ***Mrpl50*** | 1048.6 | 483.1 | 297.9 | 43.1 | 41.1 | 31.4 | 51.7 | 26.8 | 58.1 |
| Q64521 | ***Gpd2*** | 3337.4 | 1683 | 1105.6 | 323.1 | 411.6 | 319 | 331.3 | 278.3 | 349.5 |
| O88844 | ***Idh1*** | 498.5 | 2629.2 | 2402.9 | 203.4 | 91.3 | 122.9 | 103.1 | 82.1 | 134.8 |
| Q60648 | ***Gm2a*** | 286 | 156.1 | 107.8 | 48.2 | 50.6 | 50.1 | 43.3 | 43.2 | 54.3 |
| P05202 | ***Got2*** | 755.8 | 3572.2 | 3740.3 | 310 | 157.6 | 309.7 | 225 | 85.9 | 346.7 |
| P20108 | ***Prdx3*** | 3603.9 | 3147.2 | 1560.9 | 963.5 | 1577.3 | 875.5 | 1132 | 1260.8 | 970.5 |
| P16460 | ***Ass1*** | 128.8 | 372.6 | 603.5 | 34.4 | 24.6 | 26.1 | 38.7 | 40.1 | 30.7 |
| Q8JZQ2 | ***Afg3l2*** | 2410.9 | 1185.2 | 547.1 | 34.4 | 39.1 | 16.8 | 32.4 | 4.4 | 40.2 |
| P35700 | ***Prdx1*** | 53.3 | 299.3 | 229.7 | 690.4 | 566.7 | 1457.9 | 2714.5 | 4206.1 | 9609.9 |
| P25688 | ***Uox*** | 16 | 127.9 | 390.3 | 352.6 | 673.1 | 572.6 | 502.9 | 1245.9 | 1295.5 |
| Q07797 | ***Lgals3bp*** | 267.1 | 161.9 | 107.9 | 316.5 | 300.6 | 272.5 | 241.2 | 141 | 146.9 |
| O09159 | ***Man2b1*** | 484.4 | 1613 | 575.4 | 21.5 | 7.8 | 17.5 | 55.8 | 19.1 | 21.1 |
| P61264 | ***Stx1b*** | 1278.3 | 1156.1 | 1644.7 | 1016.7 | 1051 | 898.9 | 1454.7 | 799.6 | 960.3 |
| Q3URE1 | ***Acsf3*** | 222.7 | 812.4 | 817.5 | 97.1 | 91.9 | 221.1 | 118.3 | 39.7 | 140.2 |
| Q3U5Q7 | ***Cmpk2*** | 2740.3 | 1055.1 | 758.8 | 20.5 | 35.8 | 100 | 43.7 | 16.9 | 355.4 |
| P52825 | ***Cpt2*** | 315.6 | 1824.5 | 2188.7 | 144 | 81 | 78.8 | 158.4 | 78.1 | 92.6 |
| O08705 | ***Slc10a1*** | 29560.6 | 12337.7 | 7472.2 | 712.6 | 943.7 | 787.9 | 1351.7 | 739.2 | 1273.3 |
| Q99M71 | ***Epdr1*** | 450.2 | 179.8 | 213.4 | 116.3 | 49.8 | 66.7 | 80.3 | 53.2 | 68.4 |
| E9PV24 | ***Fga*** | 327.5 | 1564.6 | 1876.6 | 220.5 | 75.3 | 162.3 | 108.2 | 20.8 | 124 |
| Q9JHU4 | ***Dync1h1*** | 957.8 | 601 | 301.8 | 182.5 | 192.4 | 166.1 | 245.4 | 118.6 | 160.4 |
| P62918 | ***Rpl8*** | 591.5 | 313.6 | 219.1 | 799.5 | 611.8 | 603.3 | 706.9 | 305.4 | 380.1 |
| Q64475 | ***Hist1h2bb*** | 99.2 | 449.9 | 234.3 | 27.8 | 28.2 | 22.4 | 37.3 | 32.9 | 31.2 |
| Q9JKR6 | ***Hyou1*** | 6896.1 | 4065.1 | 2296.4 | 1142.7 | 1575.4 | 1321.7 | 2029.7 | 1191.7 | 1869.6 |

| | | | | | | | | | | |
|---|---|---|---|---|---|---|---|---|---|---|
| Q9CQI6 | ***Cotl1*** | 6048.5 | 2813 | 1352.3 | 225.6 | 266.6 | 223 | 253 | 131.4 | 204.7 |
| P01901 | ***H2-K1*** | 30.4 | 149.2 | 233.8 | 235.2 | 293.6 | 330.7 | 393.8 | 615 | 599.8 |
| P38647 | ***Hspa9*** | 548 | 2414.3 | 1560.6 | 417.4 | 114.4 | 288.9 | 315.2 | 155.5 | 362.8 |
| Q9QXF8 | ***Gnmt*** | 1543.3 | 15452.5 | 17635.1 | 135.5 | 207.4 | 148.8 | 211.3 | 246.5 | 261.3 |
| Q80XL6 | ***Acad11*** | 9922.5 | 5820.7 | 3938 | 2736.4 | 2702.8 | 2265.7 | 4029.6 | 1830.5 | 2567.9 |
| UPSP:K22E_HUMAN | | 10.3 | 19.5 | 14.2 | 595.2 | 1763.3 | 459.5 | 347.1 | 1800.3 | 496.2 |
| P17182 | ***Eno1*** | 9286.4 | 5614.4 | 3140.2 | 1703.1 | 2260.8 | 2060.3 | 3134.8 | 2391.3 | 2664.2 |
| P35293 | ***Rab18*** | 574.9 | 925.3 | 776.9 | 1180.4 | 1004.8 | 929.7 | 782.8 | 1394.5 | 1028.7 |
| P20029 | ***Hspa5*** | 121.7 | 433.7 | 418.7 | 2190.2 | 498.6 | 1691.1 | 589.9 | 135.5 | 581 |
| Q61176 | ***Arg1*** | 314.7 | 932.7 | 994.1 | 217.1 | 204.7 | 350.1 | 222.8 | 217.2 | 274.6 |
| Q9DCX2 | ***Atp5h*** | 1327.1 | 940.6 | 555.2 | 420.8 | 477.5 | 452.8 | 451.6 | 258.9 | 357 |
| P53657 | ***Pklr*** | 7225.1 | 5560.4 | 2706.9 | 1702.6 | 2637.1 | 2330.7 | 2283.2 | 1889 | 2004.5 |
| P52760 | ***Hrsp12*** | 7070.6 | 5655.8 | 2954.3 | 2029.8 | 3077.7 | 2423.8 | 4211.6 | 3278.7 | 3350.5 |
| P26443 | ***Glud1*** | 419.7 | 1494.1 | 1822.6 | 404.1 | 175.7 | 357.4 | 208.4 | 88.2 | 289.9 |
| Q9JLT4 | ***Txnrd2*** | 160.5 | 864.8 | 596 | 141.9 | 58.1 | 73.9 | 122.1 | 26.9 | 104.7 |
| Q9DBG1 | ***Cyp27a1*** | 139.2 | 623.1 | 687.2 | 111.1 | 94.8 | 134.4 | 107.2 | 73 | 88.2 |
| Q99LX0 | ***Park7*** | 406.8 | 156.5 | 84.2 | 418 | 420.5 | 432.1 | 248.4 | 134.3 | 153.1 |
| O35490 | ***Bhmt*** | 52.3 | 409.9 | 174.9 | 7620.1 | 910.9 | 4457.6 | 3589.9 | 608.5 | 3660.6 |
| Q9CXJ4 | ***Abcb8*** | 1133.9 | 650 | 428.3 | 1034.2 | 1582.8 | 1249.4 | 927.9 | 1277.6 | 1133 |
| Q8BLE7 | ***Slc17a6*** | 33.6 | 44.1 | 106.8 | 107.6 | 481.6 | 712.4 | 567.6 | 321.7 | 412.7 |
| Q00623 | ***Apoa1*** | 2192 | 1135.5 | 515.5 | 246.5 | 263.4 | 258.3 | 212.9 | 196.9 | 177 |
| Q9JHS4 | ***Clpx*** | 86825.5 | 37585.5 | 13310.6 | 723 | 678 | 482.9 | 617.2 | 230.5 | 858.3 |
| P68373 | ***Tuba1c*** | 341.9 | 6799.6 | 8296 | 23.2 | 60.9 | 60.3 | 71 | 35.2 | 86.3 |
| Q8VCW8 | ***Acsf2*** | 4592.6 | 2173.8 | 1460.6 | 3905 | 5543.7 | 5702.5 | 4293.3 | 3260.9 | 3759.4 |
| Q3UNZ8 | | 8318.3 | 4327.7 | 2553.1 | 1201.7 | 1855.2 | 1431.1 | 3532.2 | 2122.4 | 2431.3 |
| Q8BK48 | ***Ces2e*** | 411.2 | 240.1 | 151 | 478 | 421.5 | 399.9 | 321.7 | 207 | 236.2 |
| Q04447 | ***Ckb*** | 807.9 | 3856 | 2880.3 | 1031.6 | 203.7 | 550.5 | 633 | 172.9 | 731.9 |
| Q3UEG6 | ***Agxt2*** | 13192.1 | 8052.1 | 4999.4 | 3700.6 | 4399.6 | 3265.3 | 5406.8 | 2389.7 | 3493 |
| Q7TMR0 | ***Prcp*** | 99.8 | 343.4 | 554.3 | 626.1 | 737.4 | 521.1 | 463.6 | 1480.5 | 722.6 |
| Q9QXV0 | ***Pcsk1n*** | 4393.2 | 2826.2 | 1518.1 | 1134.1 | 1413.6 | 1109 | 2389.7 | 1339.2 | 1964.1 |
| Q7TQI3 | ***Otub1*** | 120.6 | 284.6 | 219.3 | 338.7 | 673.5 | 344 | 289.9 | 1339.3 | 581.7 |
| P05064 | ***Aldoa*** | 2410.8 | 8165 | 10682.6 | 1579.7 | 1923.9 | 2774.1 | 1677.6 | 3340.3 | 3174.7 |
| P24456 | ***Cyp2d10*** | 294.4 | 146.4 | 99.1 | 339.8 | 303.7 | 270.2 | 408.5 | 172 | 223.4 |
| O08539 | ***Bin1*** | 7342.9 | 3216.2 | 2482.1 | 1129.6 | 1526.4 | 1279.2 | 1662.9 | 1076.4 | 1534.1 |
| Q8QZR3 | ***Ces2a*** | 82.8 | 353.5 | 367.2 | 90 | 30.4 | 110.3 | 52 | 16.2 | 83.9 |
| Q8K0D5 | ***Gfm1*** | 160.3 | 410.7 | 523.9 | 201.8 | 80.7 | 137.6 | 185.4 | 82.5 | 308.9 |
| Q61425 | ***Hadh*** | 87.7 | 380.7 | 907.7 | 809 | 1030 | 1018.9 | 1170.7 | 2093 | 2242 |
| P50516 | ***Atp6v1a*** | 544.5 | 407.5 | 243.6 | 143 | 249.9 | 247.8 | 267.9 | 233.1 | 339.9 |
| Q6IRU2 | ***Tpm4*** | 14811 | 8665.4 | 5883.9 | 4332.4 | 5134.3 | 4336.7 | 6082 | 3117.3 | 4700 |
| P52196 | ***Tst*** | 685.4 | 2193.6 | 2694.5 | 1002 | 188.1 | 593.6 | 403.9 | 103.8 | 517.8 |
| P97742 | ***Cpt1a*** | 8496.4 | 5594.5 | 3514.3 | 2619.9 | 3490.2 | 2972.9 | 5219 | 3876.3 | 4245.8 |
| P50285 | ***Fmo1*** | 314 | 940.3 | 1612.9 | 208.4 | 130.2 | 328.3 | 115.1 | 49.3 | 342.5 |
| P62242 | ***Rps8*** | 1530.8 | 958.7 | 411.3 | 1468.4 | 1917.2 | 1545.8 | 1337 | 833.5 | 712.5 |
| UPSP:K2C5_HUMAN | | 1444.5 | 5306.9 | 4360.8 | 2007.6 | 670.2 | 1374.3 | 894.2 | 506.9 | 1652.2 |
| P11499 | ***Hsp90ab1*** | 198.1 | 945.7 | 2743.6 | 1958.7 | 4051.6 | 3498.4 | 3068.9 | 7839.3 | 5788.8 |
| P18242 | ***Ctsd*** | 1767.2 | 756.3 | 550.7 | 302 | 351.8 | 264.5 | 276.8 | 164.6 | 263 |

| | | | | | | | | | | |
|---|---|---|---|---|---|---|---|---|---|---|
| O88741 | ***Gdap1*** | 480.2 | 923.6 | 2163.2 | 233.1 | 248.1 | 206.2 | 420 | 388.1 | 456.7 |
| P19157 | ***Gstp1*** | 526.5 | 233.4 | 141.9 | 479.7 | 575.1 | 528.2 | 728.6 | 461.4 | 516.6 |
| Q9CQ54 | ***Ndufc2*** | 219.8 | 929.2 | 2132.8 | 35 | 47.4 | 15.2 | 26.5 | 10.2 | 29.4 |
| P11714 | ***Cyp2d9*** | 306.2 | 1159 | 1084.9 | 318.3 | 372.5 | 310 | 441.2 | 757.5 | 521.7 |
| P68134 | ***Acta1*** | 26.5 | 101.6 | 105.4 | 626.4 | 2819.7 | 784 | 1557.2 | 8632.7 | 4191.3 |
| P46660 | ***Ina*** | 125.5 | 100 | 264.9 | 207.2 | 1257.1 | 2070.4 | 919.2 | 755.7 | 1075 |
| P38060 | ***Hmgcl*** | 402 | 277.1 | 211.2 | 507 | 411.5 | 363.8 | 397.3 | 256.4 | 263.8 |
| Q99PT1 | ***Arhgdia*** | 925.8 | 3758.9 | 4503 | 1480.1 | 341.9 | 1007.3 | 654.8 | 145 | 821.8 |
| Q5RKZ7 | ***Mocs1*** | 111.9 | 72.9 | 183.3 | 134 | 1061.9 | 1718.4 | 597.4 | 600.8 | 696.1 |
| Q9JJI8 | ***Rpl38*** | 283.1 | 1252.6 | 284.9 | 13.6 | 17.8 | 6.7 | 10.8 | 2.6 | 13.9 |
| P62880 | ***Gnb2*** | 773.5 | 5438.1 | 11655.7 | 145.2 | 264.7 | 65.3 | 93.9 | 44.5 | 85.8 |
| P99028 | ***Uqcrh*** | 541.4 | 219.2 | 120.2 | 335.1 | 992.3 | 857 | 347 | 798.7 | 638.1 |
| Q8CI94 | ***Pygb*** | 344.7 | 224.9 | 105.5 | 328.1 | 391 | 347.5 | 369.4 | 233.4 | 247.7 |
| P11798 | ***Camk2a*** | 423.7 | 1538.1 | 2058.3 | 1937.8 | 2247.2 | 2864.7 | 2806.6 | 5790.9 | 5569.7 |
| O55022 | ***Pgrmc1*** | 283.1 | 803 | 93.4 | 6.4 | 8.6 | 5.6 | 4.9 | 3.5 | 0 |
| P15508 | ***Sptb*** | 356.1 | 791 | 965.9 | 344.8 | 298.5 | 453.7 | 451.5 | 515.7 | 501.2 |
| Q9CQ62 | ***Decr1*** | 939.7 | 477.9 | 209 | 132 | 189.5 | 144.9 | 207.8 | 179.5 | 174.8 |
| Q9DCT2 | ***Ndufs3*** | 44.8 | 151.8 | 494.4 | 288.2 | 736.7 | 852.2 | 397.5 | 1317.5 | 1618 |
| P11589 | ***Mup2*** | 869 | 232.5 | 168.2 | 538.8 | 1192.4 | 1600.7 | 710.5 | 759.6 | 1006.3 |
| Q9WUM5 | ***Suclg1*** | 1028.6 | 6250.5 | 3480 | 1419 | 252.9 | 753.2 | 554.5 | 95.3 | 532.7 |
| P14211 | ***Calr*** | 53.6 | 120.1 | 243.9 | 215.8 | 259.7 | 248.7 | 208.9 | 481.2 | 375.6 |
| Q9JM76 | ***Arpc3*** | 24.6 | 107.6 | 266.1 | 2.6 | 10 | 6.6 | 5 | 4.7 | 5.1 |
| P17710 | ***Hk1*** | 5250.5 | 3581.7 | 2616.1 | 2125.7 | 2666.5 | 2463.1 | 1868 | 1603.3 | 1547.3 |
| Q9WV96 | ***Timm10b*** | 540.7 | 485.6 | 274.9 | 342.9 | 238.4 | 257.1 | 234.8 | 318.7 | 216.4 |
| P12710 | ***Fabp1*** | 213.3 | 1381.9 | 3284.6 | 33.6 | 68.5 | 31.6 | 30.3 | 2.1 | 30.7 |
| O55125 | ***Nipsnap1*** | 215.9 | 2302.2 | 4634.9 | 111.1 | 175.7 | 133.4 | 104.6 | 68.1 | 100.5 |
| P09671 | ***Sod2*** | 80.6 | 95 | 209.1 | 118.7 | 929.2 | 1597.5 | 469.7 | 568.3 | 801.5 |
| P58252 | ***Eef2*** | 664.3 | 378.3 | 236.2 | 596.7 | 669 | 689.9 | 446.3 | 377.9 | 349.3 |
| Q68FD5 | ***Cltc*** | 800 | 3083.6 | 8735.7 | 71.8 | 153.8 | 26.4 | 27.5 | 18.8 | 52.1 |
| P17047 | ***Lamp2*** | 187.8 | 604.4 | 1435.2 | 1233 | 1533.1 | 1438.4 | 1435.5 | 2565 | 2168.9 |
| P03995 | ***Gfap*** | 107.4 | 588.5 | 1234.3 | 33.2 | 83.1 | 109.7 | 68.8 | 125.3 | 176.4 |
| O35114 | ***Scarb2*** | 93.9 | 458.4 | 311.8 | 151.2 | 65.1 | 80.4 | 86.3 | 35.7 | 75.7 |
| Q91X83 | ***Mat1a*** | 193.7 | 216.7 | 87 | 257.9 | 308.7 | 196.1 | 289.7 | 249.2 | 135.7 |
| P17183 | ***Eno2*** | 3322.3 | 18948.4 | 4619.4 | 193.9 | 421.5 | 565.7 | 590 | 668.2 | 897.2 |
| P53810 | ***Pitpna*** | 293.4 | 700.6 | 1441.9 | 179.3 | 262.7 | 267.4 | 271 | 232 | 294.3 |
| P34914 | ***Ephx2*** | 128.6 | 101.4 | 151.3 | 158.5 | 174.1 | 138.9 | 121.7 | 99.2 | 94.5 |
| P19783 | ***Cox4i1*** | 561.2 | 1191.3 | 2778.8 | 2726.4 | 2550.5 | 2631.2 | 3162.2 | 4996.9 | 3637.6 |
| Q99KE1 | ***Me2*** | 123.7 | 741.8 | 1990.7 | 18.1 | 31 | 6.4 | 16.1 | 5.2 | 11.1 |
| Q9DCS3 | ***Mecr*** | 127.6 | 621.8 | 591.1 | 199.5 | 91.4 | 212.3 | 110.1 | 76.9 | 184 |
| Q9JM62 | ***Reep6*** | 201.3 | 1453 | 3841.7 | 33 | 36.6 | 35.3 | 56 | 30.7 | 44.1 |
| Q9D7A8 | ***Armc1*** | 415.5 | 1302.3 | 2150.9 | 617.4 | 245.9 | 420.6 | 253.1 | 268.3 | 461 |
| P17427 | ***Ap2a2*** | 1394.7 | 3377.5 | 7818.5 | 6438.3 | 7616 | 8608.6 | 8366.9 | 14875.2 | 14517.7 |
| Q9QWR8 | ***Naga*** | 297.9 | 54 | 58.5 | 222.7 | 368.2 | 837.4 | 238.8 | 168.6 | 345 |
| Q8VDD5 | ***Myh9*** | 327.1 | 179 | 359 | 253.8 | 2072.3 | 4169.9 | 1089.4 | 1203.6 | 1567.7 |
| P67984 | ***Rpl22*** | 1167.4 | 3379.5 | 5531.9 | 1532.3 | 957.2 | 1283 | 1493.1 | 1508.2 | 1600 |
| Q9D819 | ***Ppa1*** | 345.9 | 2697.9 | 7544.3 | 23.6 | 31.5 | 41 | 65.8 | 51.8 | 74.9 |
| Q61035 | ***Hars*** | 250.5 | 678.1 | 336.7 | 176.2 | 244.1 | 196.2 | 254.9 | 205.7 | 197.4 |

| | | | | | | | | | | |
|---|---|---|---|---|---|---|---|---|---|---|
| Q9Z2I9 | ***Sucla2*** | 92.7 | 499.4 | 1380 | 32.8 | 43 | 33.9 | 36.7 | 23 | 21.6 |
| Q9DCM0 | ***Ethe1*** | 89.4 | 374.8 | 1140.4 | 42.4 | 17.9 | 16.8 | 26.6 | 18 | 25 |
| Q9JK42 | ***Pdk2*** | 87.2 | 551 | 1345.7 | 63.8 | 65.3 | 78.4 | 79.2 | 108.2 | 100.8 |
| Q9D379 | ***Ephx1*** | 185.5 | 790.3 | 198.2 | 82 | 71.8 | 55.5 | 89.6 | 84.8 | 75.3 |
| Q9D3D9 | ***Atp5d*** | 835 | 3392.7 | 10820.2 | 206.7 | 266.5 | 191.5 | 196.9 | 136.7 | 218.2 |
| Q62420 | ***Sh3gl2*** | 108.8 | 153.7 | 383 | 368.6 | 346.8 | 339.5 | 432.6 | 409.6 | 465.1 |
| Q8CIM7 | ***Cyp2d26*** | 86.4 | 377.5 | 578.9 | 113.5 | 127.6 | 118.5 | 90.4 | 79.3 | 85.8 |
| P68369 | ***Tuba1a*** | 816.5 | 2998.4 | 7041.4 | 663.2 | 704.1 | 803.9 | 978 | 1395.5 | 1491.4 |
| P39053 | ***Dnm1*** | 145.3 | 406.7 | 374.8 | 2108.2 | 295.5 | 1049.4 | 855.6 | 140.6 | 759.2 |
| P58044 | ***Idi1*** | 503.3 | 1180.5 | 1419.6 | 2234.7 | 990 | 2242.6 | 1221.2 | 421.9 | 1894.1 |
| P14152 | ***Mdh1*** | 528.6 | 2309.7 | 1807.1 | 1067.5 | 195.7 | 616.6 | 480.9 | 99.7 | 468 |
| P35505 | ***Fah*** | 185.4 | 1302.8 | 3947.1 | 50.3 | 61.6 | 88.8 | 72.9 | 103.1 | 121.8 |
| P63101 | ***Ywhaz*** | 350.2 | 2775.2 | 6674.3 | 226.5 | 407.2 | 514.3 | 334.6 | 637.5 | 681.8 |
| P08553 | ***Nefm*** | 113.1 | 498.1 | 640.6 | 105.1 | 191.5 | 205.5 | 263.4 | 508.2 | 591.4 |
| P41105 | ***Rpl28*** | 147.1 | 827.5 | 2481.4 | 50.1 | 73.1 | 99.7 | 40.5 | 12.3 | 146.1 |
| Q9CQ69 | ***Uqcrq*** | 342.9 | 1368.8 | 4633.9 | 82.6 | 68.7 | 162.9 | 141.7 | 103.5 | 226 |
| P62259 | ***Ywhae*** | 237.7 | 1286.9 | 2944.2 | 2191.2 | 2962.3 | 3243.3 | 2934.5 | 5525.2 | 5314.3 |
| Q7TQF7 | ***Amph*** | 1405 | 759.7 | 518.9 | 1410.4 | 1321.9 | 1202.7 | 2514.7 | 1059.7 | 1427.2 |
| Q99KR7 | ***Ppif*** | 627.3 | 3116.9 | 11305.9 | 102.6 | 197.3 | 139.5 | 136.8 | 152.7 | 228.9 |
| Q9DCJ5 | ***Ndufa8*** | 16706.8 | 9016.2 | 4427.8 | 4090.1 | 5071.8 | 4724.8 | 4849 | 2906.2 | 3850.2 |
| Q6ZQI3 | ***Mlec*** | 96.6 | 614.9 | 1346.3 | 171.1 | 120.8 | 156.9 | 180.1 | 304.9 | 239.5 |
| P27773 | ***Pdia3*** | 1816.2 | 1164 | 599.7 | 1478.5 | 1957.8 | 1818.1 | 1332 | 1020 | 1164.1 |
| Q9WVJ3 | ***Cpq*** | 162.9 | 1312.1 | 4703.7 | 54.5 | 100.8 | 8.2 | 91 | 30.3 | 32.9 |
| Q9D1Q6 | ***Erp44*** | 89.2 | 415.5 | 713.9 | 526.6 | 722.9 | 886.1 | 749.4 | 1760 | 1579 |
| Q8R3V5 | ***Sh3glb2*** | 117.4 | 538.1 | 798.8 | 261.5 | 160.4 | 156.8 | 316.1 | 412.3 | 430.8 |
| O35129 | ***Phb2*** | 23.7 | 78.4 | 144.4 | 105.5 | 147.8 | 160.5 | 162 | 279.5 | 293.7 |
| Q8BP40 | ***Acp6*** | 1536.8 | 6628.4 | 6028.3 | 3706.6 | 650.1 | 1910.7 | 1659.7 | 316.2 | 1679.6 |
| P28798 | ***Grn*** | 104.5 | 402.4 | 1049.8 | 109 | 125.5 | 118.3 | 175.4 | 248.7 | 254.7 |
| Q8BH59 | ***Slc25a12*** | 87.6 | 678.6 | 2302.9 | 49.4 | 50.9 | 117.4 | 82 | 122 | 174.9 |
| Q99M87 | ***Dnaja3*** | 33.5 | 145.8 | 221.5 | 61.1 | 42.2 | 61.2 | 64.4 | 50 | 98.2 |
| P08551 | ***Nefl*** | 215.8 | 694.1 | 1903.8 | 345.3 | 120.6 | 217.7 | 194.5 | 43.1 | 91 |
| Q8VCR7 | ***Abhd14b*** | 372.6 | 190.6 | 165.5 | 155.3 | 145.3 | 154.7 | 214.1 | 101.9 | 162.1 |
| P40630 | ***Tfam*** | 4.9 | 157.7 | 128 | 140.6 | 205 | 156.4 | 156.9 | 405.3 | 209 |
| Q9CQQ7 | ***Atp5f1*** | 114 | 626 | 2145.4 | 66.7 | 107.5 | 162.4 | 124.5 | 271.9 | 339.3 |
| Q9CPU0 | ***Glo1*** | 58.9 | 252.5 | 295.6 | 76 | 75.4 | 140.3 | 117.6 | 156.6 | 201.8 |
| Q5FW57 | ***Gm4952*** | 156.8 | 908.6 | 3111.8 | 132.8 | 216.9 | 146.6 | 186.3 | 290.7 | 293.1 |
| P15864 | ***Hist1h1c*** | 166.7 | 173.8 | 154.4 | 144.2 | 240.6 | 255.4 | 389.3 | 773 | 594.4 |
| P08228 | ***Sod1*** | 228.3 | 506.8 | 1232.4 | 237.4 | 274.6 | 232.4 | 303.5 | 190 | 290.9 |
| O55100 | ***Syngr1*** | 178.4 | 1509.5 | 4098.4 | 301.6 | 347.4 | 454.2 | 309.1 | 398.6 | 570.3 |
| Q571E4 | ***Galns*** | 4780.2 | 2547.4 | 1487.8 | 3211.5 | 5153 | 5218 | 3506.8 | 3188.9 | 3857.2 |
| Q8BVI4 | ***Qdpr*** | 375.7 | 2092.8 | 5264.1 | 3476.4 | 5132.2 | 5411.3 | 4666.1 | 8430.1 | 8223 |
| Q920A5 | ***Scpep1*** | 224.2 | 946.5 | 2842.8 | 232 | 273.1 | 350 | 405.9 | 917.8 | 901.6 |
| Q99P72 | ***Rtn4*** | 119.4 | 385 | 556.2 | 257.2 | 114.9 | 150.2 | 153.1 | 135.5 | 105.1 |
| B2RSH2 | ***Gnai1*** | 173.5 | 437.9 | 963.6 | 209.6 | 209.9 | 226.1 | 260.6 | 286.5 | 327 |
| Q91XV3 | ***Basp1*** | 70.2 | 522.4 | 1682.6 | 111.7 | 128.6 | 127.2 | 207.3 | 277.4 | 270.7 |
| Q9DCP2 | ***Slc38a3*** | 130.2 | 312.2 | 218.1 | 244.2 | 396.9 | 293.8 | 237.3 | 535.4 | 364.1 |
| P16406 | ***Enpep*** | 237.7 | 616.8 | 1116.6 | 323.7 | 321 | 329.6 | 337.9 | 506.9 | 470.7 |

| | | | | | | | | | | |
|---|---|---|---|---|---|---|---|---|---|---|
| Q91WD5 | ***Ndufs2*** | 560.7 | 1128 | 1249 | 964.6 | 314.1 | 541.9 | 463 | 225 | 538.6 |
| P07759 | ***Serpina3k*** | 32.4 | 73.8 | 79 | 454.6 | 63.3 | 135.7 | 510.2 | 201.1 | 549.3 |
| Q8JZR0 | ***Acsl5*** | 394.1 | 2598.7 | 9882.3 | 401.7 | 784 | 615.5 | 528.4 | 1872 | 1350 |
| Q9DBG6 | ***Rpn2*** | 4455 | 2530.6 | 2116.1 | 2044.4 | 2253.2 | 2020.8 | 2245 | 1178.9 | 1524.2 |
| Q69ZS6 | ***Sv2c*** | 116.9 | 326.9 | 635.2 | 183.8 | 159.3 | 156.9 | 170.8 | 298.1 | 239.2 |
| P62821 | ***Rab1A*** | 385.9 | 2400.4 | 6879.6 | 427.5 | 945.9 | 954.7 | 676.7 | 1638.7 | 1681.2 |
| P02088 | ***Hbb-b1*** | 13 | 112.8 | 215.2 | 40.1 | 34.8 | 44.6 | 34.4 | 81.6 | 70.2 |
| Q3UHB1 | ***Nt5dc3*** | 580.6 | 312 | 171.1 | 600.4 | 483.5 | 460.9 | 517.5 | 278 | 273 |
| Q78IK4 | ***Apool*** | 81.1 | 344.1 | 891.7 | 155.4 | 170.4 | 94.4 | 149.8 | 254.3 | 162.7 |
| Q05920 | ***Pc*** | 4265 | 1726.1 | 899.3 | 934.3 | 950 | 1207.4 | 465.2 | 275.4 | 428.5 |
| E9PUL5 | ***Prrt2*** | 21.1 | 135.5 | 625.6 | 19.3 | 40.3 | 37.1 | 19.1 | 18.5 | 26.4 |
| Q921H8 | ***Acaa1a*** | 21.1 | 245.5 | 698.4 | 56.9 | 76.7 | 99.8 | 79.3 | 165.3 | 143 |
| Q9Z1J3 | ***Nfs1*** | 75.5 | 167.6 | 319.7 | 238.4 | 318.9 | 279.9 | 273 | 339 | 414.8 |
| P14094 | ***Atp1b1*** | 65.4 | 100.6 | 83.8 | 180 | 52.2 | 174.6 | 67.2 | 25.1 | 108.5 |
| P97872 | ***Fmo5*** | 156 | 551.9 | 545.4 | 345.7 | 92.7 | 273.5 | 166.9 | 34.6 | 183.9 |
| P61982 | ***Ywhag*** | 5181.8 | 3355 | 1777.9 | 1797.6 | 2708 | 2193.5 | 3031.1 | 2366.5 | 2641.3 |
| Q9DBJ1 | ***Pgam1*** | 773.6 | 1902 | 2023.1 | 3209.1 | 1347.9 | 2525.8 | 2933.5 | 4022.8 | 3311.8 |
| P68368 | ***Tuba4a*** | 556 | 3079.8 | 4829 | 2396.2 | 477.3 | 792.1 | 995.3 | 168.3 | 370.7 |
| Q8C650 | ***42988*** | 11383.4 | 6044.9 | 3914.9 | 9662.1 | 10614.5 | 8883.6 | 9659.1 | 5554.9 | 5724.8 |
| D3Z7P3 | ***Gls*** | 70.1 | 331 | 743.6 | 264.8 | 75.1 | 103.9 | 120.5 | 46.5 | 63.5 |
| P52480 | ***Pkm*** | 283.2 | 1298 | 3570.3 | 2113.8 | 3221.4 | 3400.3 | 2932.8 | 5440.9 | 5862 |
| O70362 | ***Gpld1*** | 1356.5 | 1085.3 | 622.2 | 754.4 | 818.3 | 760.5 | 861.6 | 499 | 583 |
| P70349 | ***Hint1*** | 76.1 | 260.2 | 915.4 | 93.9 | 157.7 | 142 | 117.8 | 211.9 | 255.9 |
| P14131 | ***Rps16*** | 214 | 942.1 | 956.9 | 357.5 | 464.5 | 460.8 | 385.7 | 927.9 | 755.6 |
| O35643 | ***Ap1b1*** | 69.6 | 249.8 | 561.3 | 96.1 | 138.1 | 163.3 | 76.4 | 151.7 | 125.1 |
| P60710 | ***Actb*** | 321.1 | 320.4 | 365.4 | 255 | 498.4 | 569.6 | 423.9 | 1338.2 | 1078.9 |
| P62274 | ***Rps29*** | 41.2 | 121.7 | 185.2 | 99.2 | 31.2 | 66.8 | 55.5 | 78.8 | 68.4 |
| O55126 | ***Gbas*** | 1208.5 | 668.2 | 379.2 | 1025.4 | 1108.3 | 928.4 | 1610.1 | 809 | 1036.6 |
| P11610 | ***Cd1d2*** | 8 | 47.2 | 181.6 | 70.9 | 193.9 | 182.6 | 152.4 | 344.3 | 553.9 |
| Q99PL5 | ***Rrbp1*** | 1098.2 | 7615.4 | 6152.5 | 4776.4 | 1030.9 | 1877.3 | 2828 | 378.9 | 843.6 |
| Q60759 | ***Gcdh*** | 30.1 | 369.7 | 280.5 | 294.3 | 401.7 | 319.6 | 340.9 | 846 | 641.9 |
| P22315 | ***Fech*** | 239.6 | 847.9 | 609.1 | 541.4 | 104.5 | 386.2 | 201.4 | 47 | 267.4 |
| Q9CQW2 | ***Arl8b*** | 146.7 | 620.6 | 334.2 | 334.6 | 71.6 | 227.6 | 129.2 | 35.1 | 163.6 |
| P50136 | ***Bckdha*** | 53.7 | 172.7 | 261.4 | 190.8 | 246 | 232 | 177.8 | 417.2 | 338.8 |
| Q9CR62 | ***Slc25a11*** | 273.8 | 173.9 | 110.5 | 194.3 | 268.2 | 247 | 139.8 | 107.8 | 132.2 |
| P62908 | ***Rps3*** | 141.9 | 863.6 | 2199.1 | 1223.9 | 1867.6 | 1969.5 | 2036.5 | 3753.8 | 3531.5 |
| Q9Z2I8 | ***Suclg2*** | 168.1 | 851.3 | 1239.9 | 714.1 | 1515.8 | 1147.8 | 1117.9 | 3044 | 2507.5 |
| P57780 | ***Actn4*** | 2320.3 | 2536.6 | 1218.8 | 1266.5 | 1853.7 | 1684.9 | 1388 | 1480.4 | 1500.3 |
| P42932 | ***Cct8*** | 1650.6 | 948.7 | 469.7 | 805.4 | 622.3 | 660.4 | 925.5 | 466.9 | 576.6 |
| Q9D1H6 | ***Ndufaf4*** | 840.8 | 2060.9 | 3190.9 | 1983.8 | 544.3 | 1343.7 | 725.1 | 210 | 1080.3 |
| P45952 | ***Acadm*** | 2970.2 | 1772.5 | 1214.2 | 2697.1 | 2467.9 | 2274.6 | 2129.6 | 1446.6 | 1255.5 |
| P63040 | ***Cplx1*** | 8404.1 | 4118.1 | 2752.3 | 3739.6 | 3474.5 | 3347.7 | 4418.1 | 2418.8 | 3310.8 |
| P62267 | ***Rps23*** | 2610.6 | 1213.7 | 615.3 | 991.7 | 983.8 | 831.1 | 1011.4 | 420.6 | 568.8 |
| Q8BMS1 | ***Hadha*** | 2546.9 | 1269.5 | 725.3 | 1221.9 | 1007 | 793.5 | 941.3 | 451.3 | 485.7 |
| P46656 | ***Fdx1*** | 110.6 | 634.4 | 2281.9 | 388 | 372.7 | 476.1 | 556.3 | 770.9 | 986.3 |
| P09411 | ***Pgk1*** | 3311.1 | 8887.2 | 7647.2 | 23249.3 | 2350.8 | 11357.4 | 9141.3 | 1340.6 | 6592.6 |
| Q9WVA2 | ***Timm8a1*** | 579.2 | 1632.6 | 710.4 | 771.2 | 364.2 | 832.5 | 1561.7 | 349.2 | 1580.4 |

| | | | | | | | | | | |
|---|---|---|---|---|---|---|---|---|---|---|
| P29758 | ***Oat*** | 63.7 | 384.9 | 321 | 301.6 | 418.3 | 319.2 | 333.8 | 1052.6 | 651 |
| Q61699 | ***Hsph1*** | 302.5 | 86.9 | 1012 | 364.9 | 554.6 | 1946.8 | 523.7 | 1010.6 | 2990.7 |
| P84099 | ***Rpl19*** | 276.9 | 329.6 | 353.1 | 399.8 | 151.4 | 207.3 | 264.1 | 158.3 | 181.2 |
| P35441 | ***Thbs1*** | 669.3 | 285.5 | 192.2 | 612.9 | 465.7 | 463.1 | 616.1 | 322.3 | 327.3 |
| Q99J99 | ***Mpst*** | 384 | 798.4 | 609.2 | 845.1 | 593.3 | 706.9 | 506.9 | 706.4 | 680.6 |
| UPSP:K1CI_HUMAN | | 149.2 | 328.6 | 904.8 | 419.7 | 806.7 | 815.9 | 767.5 | 2103.7 | 1999.2 |
| P62270 | ***Rps18*** | 2466 | 1705.3 | 987.9 | 1343.6 | 1458.6 | 1298.6 | 2122.1 | 1200.1 | 1634.3 |
| Q9Z2I0 | ***Letm1*** | 2296.3 | 1484.1 | 866.8 | 1123.1 | 1301.5 | 1194.4 | 1914.9 | 1125.4 | 1646 |
| P51863 | ***Atp6v0d1*** | 1291.1 | 817.1 | 406.5 | 497.2 | 673.1 | 698.8 | 697.4 | 493.2 | 635 |
| P47962 | ***Rpl5*** | 26.7 | 39.8 | 32.1 | 52 | 24.1 | 44.5 | 762 | 1762.7 | 3762.2 |
| Q9CPY7 | ***Lap3*** | 669.1 | 387.4 | 212.2 | 292.5 | 368.5 | 281.1 | 281.5 | 233.3 | 227.3 |
| Q9CQJ8 | ***Ndufb9*** | 36 | 243.4 | 609.3 | 424.9 | 471.7 | 397.1 | 539 | 1139.7 | 870.9 |
| P50171 | ***Hsd17b8*** | 1591.5 | 872.7 | 461.9 | 524.2 | 883.5 | 711.8 | 696 | 1023.1 | 904.7 |
| Q8BH24 | ***Tm9sf4*** | 359.4 | 2972 | 2852.9 | 2490.5 | 525.2 | 745.1 | 761.5 | 219.5 | 364.6 |
| Q9R0P3 | ***Esd*** | 198.6 | 990.9 | 2030.9 | 442.9 | 721.6 | 811.6 | 537.8 | 1129.8 | 1097.3 |
| P47915 | ***Rpl29*** | 336.6 | 1074.9 | 1652.4 | 1047.1 | 266.4 | 737.9 | 280.6 | 95.9 | 627.4 |
| P11352 | ***Gpx1*** | 358.4 | 1230 | 1326.7 | 785 | 782.7 | 640.7 | 967.7 | 1938.9 | 1525 |
| Q64133 | ***Maoa*** | 396.8 | 241.6 | 140.7 | 392.1 | 294.2 | 278.1 | 367.4 | 147.3 | 165.2 |
| P14873 | ***Map1b*** | 142.5 | 623.4 | 1430 | 292.3 | 472.7 | 572.3 | 500.2 | 618.3 | 617.4 |
| P48758 | ***Cbr1*** | 283.3 | 206.4 | 109.6 | 134.3 | 199.7 | 146.2 | 115.1 | 106.5 | 90.3 |
| Q9JJJ3 | ***Aqp9*** | 625.3 | 3060.8 | 2927.5 | 5544.8 | 946.7 | 3493.5 | 1965.7 | 347.4 | 1712.9 |
| P15105 | ***Glul*** | 826.7 | 2854.5 | 2612.4 | 3795.6 | 1261 | 3412.3 | 1837.6 | 714.4 | 3398.6 |
| P51910 | ***Apod*** | 130.2 | 473 | 126.5 | 179.5 | 94.5 | 200.6 | 316.4 | 74.4 | 263.2 |
| O55131 | ***42985*** | 25022.9 | 12822.3 | 8306.6 | 20642.7 | 19141 | 17149 | 18960.2 | 10863.7 | 11524.9 |
| Q9D924 | ***Isca1*** | 7016.8 | 4385.7 | 3138 | 3905.2 | 4132.1 | 4096.2 | 5294.3 | 2413.1 | 3956.1 |
| O70318 | ***Epb41l2*** | 747.6 | 521.6 | 298.4 | 565.4 | 375.1 | 325.8 | 512.8 | 586.6 | 498 |
| O35488 | ***Slc27a2*** | 343.9 | 1427.4 | 921.3 | 1586.3 | 461.8 | 1632 | 817.7 | 312.2 | 1245.5 |
| Q8BGT5 | ***Gpt2*** | 213.9 | 1081.7 | 3831.7 | 676.8 | 962.7 | 1285.1 | 1353.9 | 2092.4 | 2372.3 |
| P28652 | ***Camk2b*** | 94.1 | 620.3 | 515.5 | 455.8 | 618.3 | 485.2 | 579.5 | 1348.1 | 1013.6 |
| Q91VD9 | ***Ndufs1*** | 58.6 | 322.7 | 1171.2 | 195.6 | 360 | 362.7 | 299.2 | 944.2 | 684.3 |
| Q9WTP6 | ***Ak2*** | 51.6 | 166 | 318.1 | 190.4 | 245.3 | 245.5 | 257.3 | 409.7 | 389.3 |
| P09103 | ***P4hb*** | 1344.6 | 890.5 | 484.5 | 668.3 | 836.8 | 762.8 | 823.2 | 539.4 | 709 |
| Q91YQ5 | ***Rpn1*** | 65.8 | 249 | 506.1 | 149.3 | 188.4 | 253.4 | 351.2 | 684.6 | 729.3 |
| P61458 | ***Pcbd1*** | 7918.4 | 5072.8 | 3012.7 | 6070.7 | 2502.1 | 4404 | 5245 | 3860.8 | 3252.6 |
| Q8VCH0 | ***Acaa1b*** | 615.4 | 304.6 | 141.4 | 556.8 | 336 | 430.3 | 350.6 | 210.6 | 167.8 |
| Q8R164 | ***Bphl*** | 152.5 | 86.5 | 57.3 | 109.6 | 114.9 | 119.6 | 105.8 | 65.9 | 70 |
| P60761 | ***Nrgn*** | 6355 | 5001 | 3058.5 | 3578.4 | 5027.6 | 4044.5 | 4835.4 | 2825.6 | 3908.1 |
| P14824 | ***Anxa6*** | 499.1 | 276.6 | 180.3 | 181.8 | 295.9 | 307.2 | 157.6 | 162.4 | 171.6 |
| Q3ULD5 | ***Mccc2*** | 3225.6 | 1592.1 | 921.7 | 2325.4 | 2220.3 | 2330 | 2501.8 | 1002.3 | 1470.4 |
| P40124 | ***Cap1*** | 22.1 | 96.7 | 113.6 | 106.6 | 87.5 | 85.6 | 97.2 | 302.2 | 160.2 |
| Q80TJ1 | ***Cadps*** | 369.9 | 626.7 | 1219.3 | 595 | 628.6 | 580.9 | 852.5 | 1086.5 | 1111.7 |
| Q91W90 | ***Txndc5*** | 344.2 | 1326.7 | 2452.1 | 1453.2 | 1903.4 | 1760.7 | 1548 | 3119.8 | 2266.9 |
| P50247 | ***Ahcy*** | 419.9 | 1826 | 1398 | 1355.7 | 371.3 | 1109.5 | 610 | 189.7 | 849.2 |
| P18872 | ***Gnao1*** | 588.2 | 355.3 | 274.2 | 358 | 500.6 | 523.4 | 351.5 | 303.8 | 300.9 |
| O08677 | ***Kng1*** | 81.4 | 113.3 | 602.2 | 46.5 | 53.2 | 387.3 | 70.9 | 64.1 | 576 |
| Q9D5T0 | ***Atad1*** | 1444.2 | 654.3 | 287.3 | 353.2 | 671.8 | 805.7 | 293.1 | 629.8 | 771.9 |

| | | | | | | | | | | |
|---|---|---|---|---|---|---|---|---|---|---|
| P63001 | ***Rac1*** | 8.8 | 99.4 | 135.1 | 77.1 | 102.8 | 121.4 | 70.1 | 166.9 | 99.4 |
| Q8CG76 | ***Akr7a2*** | 534.3 | 3360.8 | 1985.6 | 2751.3 | 565 | 1030.8 | 1233.7 | 252.1 | 586.7 |
| Q8JZN5 | ***Acad9*** | 1693.6 | 841.1 | 434.1 | 1375.5 | 1101.6 | 1004.1 | 1886.2 | 901.1 | 1080 |
| Q91Y97 | ***Aldob*** | 285.8 | 2262.3 | 4361.8 | 1660.8 | 2020.4 | 1665.7 | 1198.1 | 4441.5 | 2129.8 |
| P07724 | ***Alb*** | 943.5 | 522.2 | 362.3 | 458.2 | 621.5 | 523.8 | 405.8 | 325.9 | 380.1 |
| P04939 | ***Mup3*** | 579 | 585.5 | 2634.1 | 894.2 | 935.7 | 3248.1 | 1159.7 | 1811 | 4558.1 |
| Q922D8 | ***Mthfd1*** | 1951.9 | 7302.7 | 13674.7 | 7761.8 | 9762 | 9641.5 | 9014 | 18046.5 | 12602.2 |
| P54869 | ***Hmgcs2*** | 1175.7 | 398.3 | 301.7 | 443.1 | 986.6 | 833.9 | 505.8 | 792.5 | 796.3 |
| P56213 | ***Gfer*** | 12657.1 | 5621.8 | 2585.3 | 11248.1 | 6562.5 | 7023.8 | 9142.4 | 4166.4 | 4173.8 |
| Q9DC69 | ***Ndufa9*** | 456.8 | 406.8 | 181.6 | 363 | 401.1 | 382.3 | 402.7 | 279.3 | 307.2 |
| Q8QZS1 | ***Hibch*** | 993.9 | 470.3 | 326.4 | 512.8 | 585.9 | 453.4 | 460.2 | 296.4 | 358.1 |
| Q9CZW5 | ***Tomm70*** | 3501.7 | 18273.7 | 18200.2 | 21846.7 | 4204.7 | 5458.4 | 5880.2 | 1258.5 | 2169.9 |
| P84086 | ***Cplx2*** | 402 | 2392.6 | 5958.4 | 1421.9 | 2761.2 | 2771.1 | 2571.9 | 6617.2 | 5611.5 |
| P31786 | ***Dbi*** | 312.9 | 300.9 | 418.5 | 186.9 | 379.5 | 387.1 | 440.3 | 1079.7 | 884.3 |
| Q8R191 | ***Syngr3*** | 362.3 | 1360.5 | 1368.2 | 1471 | 245.4 | 878.2 | 868.1 | 169.6 | 943.6 |
| Q9DAS9 | ***Gng12*** | 173.3 | 73.4 | 75.7 | 102.7 | 128.8 | 121.8 | 164.1 | 304.2 | 291.7 |
| P63038 | ***Hspd1*** | 5275.1 | 2912.5 | 1757.3 | 3365.5 | 3214.6 | 2388.2 | 4382.9 | 2111.9 | 2364.5 |
| P62897 | ***Cycs*** | 4898.3 | 1825.7 | 1455.8 | 2447.6 | 2984 | 3732.5 | 1600.7 | 1208.3 | 1523.9 |
| Q8BWF0 | ***Aldh5a1*** | 888.8 | 2588.1 | 3034.4 | 3189.9 | 530.6 | 1931.5 | 1305.1 | 261.3 | 1725.2 |
| P19096 | ***Fasn*** | 2552.2 | 257.6 | 9738.2 | 238.9 | 376.3 | 8613.8 | 401.8 | 244 | 12998.7 |
| P16330 | ***Cnp*** | 1716.5 | 6273 | 5852.5 | 12909.4 | 1708.4 | 2200.8 | 2077 | 567.9 | 892.3 |
| Q99MR8 | ***Mccc1*** | 189.8 | 611.4 | 673.1 | 768.9 | 101.3 | 419.7 | 286.6 | 47.3 | 320.3 |
| Q64442 | ***Sord*** | 407.2 | 591.1 | 1409.3 | 357.1 | 803.1 | 991.4 | 740.6 | 1371 | 1648.3 |
| P37040 | ***Por*** | 487 | 405.3 | 285.9 | 305.9 | 442.2 | 380.2 | 341.5 | 252.8 | 357.2 |
| Q9CYW4 | ***Hdhd3*** | 89 | 228.8 | 262.4 | 217.6 | 191.9 | 208.5 | 255.3 | 342.2 | 289.7 |
| Q9D7B6 | ***Acad8*** | 5183.4 | 2617.5 | 1720.9 | 3350.4 | 3574.4 | 3309.5 | 3267 | 1790 | 2279.6 |
| Q9EQ20 | ***Aldh6a1*** | 113.9 | 415.7 | 1017.5 | 317.2 | 392 | 666.5 | 384.9 | 845.9 | 952.3 |
| P08249 | ***Mdh2*** | 1701.6 | 887 | 560.7 | 1267.9 | 1080.5 | 996.3 | 1224 | 518.6 | 614.8 |
| O09111 | ***Ndufb11*** | 452 | 311 | 157.9 | 297.8 | 341.6 | 327.9 | 284.4 | 262.5 | 386.2 |
| P21460 | ***Cst3*** | 530.1 | 993.1 | 1069.2 | 747.9 | 998.4 | 747.6 | 797.8 | 1838.7 | 1264.4 |
| Q80XN0 | ***Bdh1*** | 50.6 | 139.6 | 309.2 | 168.9 | 187.4 | 183 | 178.2 | 246.6 | 255.8 |
| P62806 | ***Hist1h4a; Hist1h4b; Hist1h4c; Hist1h4d; Hist1h4f; Hist1h4h; Hist1h4i; Hist1h4j; Hist1h4k; Hist1h4m; Hist2h4a; Hist4h4*** | 100.1 | 232.9 | 224.4 | 351.7 | 78 | 174.2 | 114.4 | 101.8 | 134.8 |
| Q9D6F9 | ***Tubb4a*** | 92 | 378.8 | 634.6 | 417.6 | 287.3 | 485.7 | 315.3 | 659.3 | 809.2 |
| O54734 | ***Ddost*** | 159.6 | 197.5 | 448.4 | 63.9 | 64.1 | 781.3 | 130.4 | 124.9 | 1234.6 |
| P43006 | ***Slc1a2*** | 23.7 | 46.1 | 102.1 | 60.4 | 65.9 | 54 | 90.9 | 51.6 | 68.5 |
| P60867 | ***Rps20*** | 72 | 453.1 | 524.2 | 252.3 | 403.8 | 341.2 | 322 | 816.1 | 532.5 |
| Q8R4N0 | ***Clybl*** | 115.2 | 502.5 | 294.6 | 476 | 161.2 | 321.1 | 249.2 | 509.6 | 337.8 |
| Q9EP69 | ***Sacm1l*** | 77.5 | 303.4 | 625.2 | 263.3 | 291.4 | 402 | 300.4 | 647.2 | 567.6 |
| P29416 | ***Hexa*** | 1354.6 | 749.2 | 658.2 | 850.7 | 1104.8 | 871.1 | 925.8 | 690.4 | 932.6 |
| Q9R1T4 | ***42984*** | 15343.2 | 7959 | 6353.2 | 7446.8 | 11231.6 | 10155.7 | 17305.1 | 11246.7 | 16831.3 |

| | | | | | | | | | | |
|---|---|---|---|---|---|---|---|---|---|---|
| Q80SW1 | ***Ahcyl1*** | 2247.2 | 1184.9 | 725.1 | 1447.3 | 1478.7 | 1334.9 | 2114.1 | 1099.6 | 1402 |
| Q02053 | ***Uba1*** | 139.1 | 706 | 1847.9 | 568.6 | 922.7 | 1083.4 | 1194.5 | 2112.9 | 2089.3 |
| Q62048 | ***Pea15*** | 152.2 | 100.1 | 70.2 | 115 | 116.2 | 86.4 | 145.4 | 68.5 | 85.7 |
| Q9R1Q8 | ***Tagln3*** | 572.4 | 2147.2 | 5768.6 | 1361.5 | 3366.2 | 3501.3 | 1338.8 | 4310.6 | 4032.7 |
| P67778 | ***Phb*** | 3515.5 | 428.2 | 11038.8 | 278.6 | 711.2 | 14860.8 | 557.2 | 419.2 | 23098.1 |
| Q9CWF2 | ***Tubb2b*** | 1509.8 | 995.6 | 755.8 | 1405.5 | 610.4 | 1295.1 | 1365.4 | 746.6 | 761.3 |
| P50428 | ***Arsa*** | 2633 | 7719.2 | 8603 | 11013.9 | 1931.9 | 5588.8 | 4558.5 | 878.7 | 4376.4 |
| Q8VDN2 | ***Atp1a1*** | 13151.7 | 6740.2 | 4407.3 | 9295.3 | 7867.6 | 7067.4 | 8160.4 | 4298.9 | 4420.9 |

**Table S2: Immunoisolated proteome of Liver, Brain and Spinal cord Avs**

Only proteins that were detected with more than 3 unique peptides are listed.

The numerical values are peptide intensity normalized to total intensity of each sample.

| uniprot | gene symbol | Intensity of Lv1 | Intensity of Lv2 | Intensity of Lv3 | Intensity of Br1 | Intensity of Br2 | Intensity of Br3 | Intensity of SC1 | Intensity of SC2 | Intensity of SC3 |
|---|---|---|---|---|---|---|---|---|---|---|
| Q8C163 | ***Exog*** | 36.7 | 28 | 36 | 735.4 | 739.2 | 711.4 | 656 | 419.3 | 371.7 |
| Q3V3R1 | ***Mthfd1l*** | 23.3 | 30.9 | 26.5 | 248.6 | 238.6 | 236.9 | 245.6 | 125 | 134.3 |
| Q60597 | ***Ogdh*** | 2200.8 | 2177.6 | 1996.6 | 9662.5 | 10561.8 | 10099.2 | 12402.3 | 7168 | 8611.1 |
| P35486 | ***Pdha1*** | 1062.2 | 887.6 | 1000.9 | 6466.5 | 7083 | 7093.8 | 7048.3 | 4176.7 | 4478.9 |
| P99029 | ***Prdx5*** | 890.7 | 876.7 | 732.5 | 6854.3 | 7515.1 | 7622.8 | 6050.1 | 5892.5 | 4171.5 |
| O55126 | ***Gbas*** | 262 | 250.1 | 307.6 | 2730.8 | 3064 | 2974.4 | 2200.1 | 1408.6 | 1446.3 |
| Q9CQR4 | ***Acot13*** | 453.8 | 325.5 | 239.3 | 3519.6 | 3179.5 | 3180.6 | 2497.8 | 1923.5 | 1576.7 |
| Q9D924 | ***Isca1*** | 19 | 11.1 | 11.4 | 210.1 | 194 | 184 | 177.7 | 106 | 109.3 |
| P17710 | ***Hk1*** | 204.5 | 455.2 | 860 | 12950.4 | 11309.6 | 11735.2 | 11833.7 | 8462.5 | 8994.5 |
| Q8BKZ9 | ***Pdhx*** | 715.1 | 531.8 | 228.7 | 5580.3 | 6168.1 | 5426.1 | 5143.7 | 3198.6 | 3151.8 |
| Q9WTT4 | ***Atp6v1g2*** | 93.9 | 54.8 | 51.6 | 364.8 | 364.1 | 387.6 | 360.8 | 883.7 | 607 |
| Q62277 | ***Syp*** | 20.9 | 70 | 100.4 | 991.9 | 1156.5 | 1045.6 | 2084.5 | 2605.6 | 3397.8 |
| Q8BWF0 | ***Aldh5a1*** | 875.6 | 735.7 | 472.2 | 4929.3 | 4678.7 | 4309.1 | 5289.7 | 4331.5 | 3525.9 |
| Q9R0P9 | ***Uchl1*** | 8.4 | 19.3 | 0 | 166.5 | 180.6 | 193.2 | 321.1 | 761.6 | 671.7 |
| P84086 | ***Cplx2*** | 186.2 | 75.8 | 131.2 | 992.7 | 986.9 | 1110.4 | 943.3 | 1128.9 | 1118.7 |
| P47911 | ***Rpl6*** | 1886.9 | 2184.6 | 2043.5 | 447.6 | 266.1 | 366.1 | 378.7 | 208.6 | 288.7 |
| O35658 | ***C1qbp*** | 167.7 | 126.6 | 87.6 | 550.4 | 581.4 | 535.5 | 502.3 | 422.5 | 283 |
| Q9CQ92 | ***Fis1*** | 224.8 | 157.4 | 113.8 | 761.6 | 738.4 | 700.3 | 584 | 667.7 | 790.1 |
| Q64521 | ***Gpd2*** | 1832.5 | 1447 | 905.8 | 7073.4 | 7616.1 | 6716.5 | 8555.5 | 6441.1 | 7645.4 |
| Q9WUR9 | ***Ak4*** | 118 | 70.2 | 22.3 | 685.4 | 660.2 | 763.5 | 394 | 284.2 | 291.6 |
| O35143 | ***Atpif1*** | 110.5 | 131 | 177.3 | 3453 | 3506.2 | 2853 | 3407.1 | 2448.3 | 1629.2 |
| Q6PIE5 | ***Atp1a2*** | 197.8 | 274.5 | 446.5 | 3086 | 3629.4 | 3737 | 9462.1 | 29452.9 | 15744.2 |
| Q9D051 | ***Pdhb*** | 1041.5 | 853.9 | 446.2 | 7210.5 | 7559.1 | 6199.8 | 6875.5 | 4487.5 | 3713.3 |
| Q9WV96 | ***Timm10b*** | 63.2 | 59.2 | 77.9 | 211.6 | 234.1 | 203.6 | 138.9 | 125.7 | 103.7 |
| Q9ESW4 | ***Agk*** | 174.7 | 239.4 | 145.9 | 565.6 | 579.4 | 565.8 | 538.5 | 376.4 | 418.4 |
| O35435 | ***Dhodh*** | 71.9 | 65.2 | 75.8 | 221.5 | 217.4 | 189.5 | 185.2 | 157.7 | 123.2 |
| Q9R0X4 | ***Acot9*** | 64.2 | 97.8 | 114.1 | 474.9 | 575.7 | 508.9 | 480.8 | 329.5 | 316.3 |
| P28740 | ***Kif2a*** | 25.9 | 21.4 | 30.7 | 83.3 | 99.9 | 91.3 | 113.7 | 116.8 | 149.4 |
| Q99LC3 | ***Ndufa10*** | 2280.6 | 1240 | 777.2 | 7228.7 | 8011 | 7167.1 | 7738.7 | 4546.3 | 5836.7 |
| Q91VS7 | ***Mgst1*** | 2200.6 | 2880.2 | 2295.2 | 53.4 | 47 | 52 | 51.3 | 63.6 | 72 |
| Q9D6R2 | ***Idh3a*** | 810.7 | 598.3 | 1175.1 | 8266.9 | 8948.6 | 10730.3 | 7387.6 | 5550.2 | 5113.3 |
| Q9CQX8 | ***Mrps36*** | 304.3 | 258.3 | 85 | 1963.3 | 2355.7 | 1850.4 | 2353.2 | 1682.5 | 1459.3 |
| Q9DCZ4 | ***Apoo*** | 410.7 | 293 | 154.9 | 1975.9 | 1816.7 | 1573.8 | 1826 | 1050.8 | 1022.4 |
| Q9Z2I9 | ***Sucla2*** | 2752.8 | 1773.7 | 1557.2 | 8541.7 | 10584 | 9596.3 | 9724.9 | 6317.4 | 6756.6 |
| Q91V14 | ***Slc12a5*** | 19.7 | 26.8 | 16 | 259.7 | 347.2 | 292.2 | 349.8 | 630.3 | 529.4 |
| Q04447 | ***Ckb*** | 56.2 | 131.6 | 77.2 | 558.5 | 680.3 | 712 | 739.3 | 2706.5 | 1981.9 |
| Q9D3D9 | ***Atp5d*** | 360.2 | 196 | 196.9 | 935.6 | 1046.7 | 924.3 | 981.2 | 685 | 740.8 |
| O08749 | ***Dld*** | 2660.4 | 1778.1 | 1620.2 | 7852 | 9602.7 | 9541.8 | 7503.5 | 5480.3 | 5595.1 |
| D3Z7P3 | ***Gls*** | 74.4 | 149.7 | 301.9 | 3509.5 | 3882.3 | 4736.4 | 1780.5 | 971.4 | 1044.3 |
| P11798 | ***Camk2a*** | 29.9 | 59.2 | 40.6 | 389.5 | 460.9 | 347.6 | 197 | 257.1 | 290.1 |
| P30275 | ***Ckmt1*** | 84.5 | 234.2 | 469.3 | 9264.6 | 7144.3 | 9919 | 8016.9 | 6071.9 | 6602.2 |
| Q3TC72 | ***Fahd2*** | 550.8 | 561.8 | 692.7 | 1670.4 | 2022.4 | 2055.9 | 1823.9 | 1347.9 | 1289.1 |

| | | | | | | | | | | |
|---|---|---|---|---|---|---|---|---|---|---|
| Q8BMF4 | ***Dlat*** | 694.4 | 739.5 | 498.5 | 6095.8 | 5699 | 4511 | 6077.9 | 3894.7 | 3376.1 |
| Q91WD5 | ***Ndufs2*** | 715.1 | 629.8 | 608.1 | 3357.5 | 3409.1 | 2636.2 | 3390.9 | 2096.2 | 2059.2 |
| Q5IRJ6 | ***Slc30a9*** | 81.6 | 108.6 | 106.9 | 459.1 | 467.1 | 363.9 | 507.8 | 385.1 | 334.4 |
| P46096 | ***Syt1*** | 56.3 | 94.6 | 184.4 | 1024.4 | 1337.4 | 1046.6 | 2102 | 2441.9 | 4497.3 |
| P63044 | ***Vamp2*** | 22.3 | 63.3 | 137.9 | 558.8 | 535.9 | 453.2 | 291.3 | 963.1 | 557 |
| Q6PIC6 | ***Atp1a3*** | 337.3 | 1179.9 | 2101.1 | 9916 | 9479.4 | 7758 | 19949.1 | 41604.1 | 33503.9 |
| Q3UUI3 | ***Them4*** | 146.7 | 65.5 | 146.6 | 585.8 | 715 | 768.1 | 538.7 | 512.9 | 396.7 |
| Q9D2G2 | ***Dlst*** | 1727.7 | 1248.3 | 830.8 | 5433.1 | 5952.7 | 4727.1 | 5807.5 | 3916.3 | 3549.9 |
| Q7TQD2 | ***Tppp*** | 21.4 | 39.6 | 43.8 | 1011.7 | 1460.5 | 1271.5 | 900.8 | 2298.7 | 1882.6 |
| P47708 | ***Rph3a*** | 10.8 | 7.3 | 11.3 | 102.2 | 144.9 | 129.7 | 315.4 | 522.7 | 704.5 |
| P61922 | ***Abat*** | 839.3 | 733 | 477.4 | 2567.2 | 2636.9 | 3220.9 | 4329.9 | 2772.1 | 3837.7 |
| Q91WC3 | ***Acsl6*** | 49.4 | 43.5 | 55.9 | 354.1 | 486.9 | 389.2 | 767.6 | 482.6 | 754.1 |
| Q9R0K7 | ***Atp2b2*** | 14.4 | 12.9 | 39.8 | 216.2 | 287.4 | 298.3 | 484.2 | 775.5 | 1098.1 |
| P50516 | ***Atp6v1a*** | 422.7 | 923.1 | 518.7 | 2149.1 | 2041.7 | 2309.6 | 2207.7 | 3788.1 | 4106 |
| Q7TSJ2 | ***Map6*** | 22.7 | 38.3 | 26.3 | 366.8 | 373.9 | 268.6 | 467.2 | 708.7 | 666.5 |
| O08601 | ***Mttp*** | 871.8 | 1061.1 | 1240.7 | 94.6 | 107.9 | 113.1 | 110.2 | 70.9 | 91.5 |
| Q9CZW5 | ***Tomm70*** | 2458.8 | 1844.6 | 1627.3 | 6042.7 | 6264.5 | 5107 | 4392.6 | 3238.4 | 2861.4 |
| Q9CZR8 | ***Tsfm*** | 109.5 | 111.4 | 44.7 | 374.7 | 467 | 482.1 | 372.8 | 262.2 | 274.5 |
| P33267 | ***Cyp2f2*** | 1431.4 | 1760.6 | 1205.9 | 60.1 | 78.4 | 71.8 | 46 | 35.5 | 69.5 |
| P23242 | ***Gja1*** | 15.6 | 24.5 | 52.6 | 181.7 | 173.1 | 221.6 | 211 | 299.3 | 410.9 |
| Q9WV98 | ***Timm9*** | 401.5 | 144.7 | 128.2 | 1248.8 | 1117.7 | 1043.9 | 705 | 738 | 520.2 |
| P54869 | ***Hmgcs2*** | 13757.4 | 9899.3 | 9903.6 | 287 | 307.4 | 297.1 | 259.8 | 125.1 | 322.3 |
| Q99LX0 | ***Park7*** | 166.1 | 215.7 | 204.7 | 556.7 | 619.2 | 726.4 | 548.7 | 768.3 | 641.5 |
| P14231 | ***Atp1b2*** | 22.8 | 50.5 | 51.7 | 293.4 | 260 | 360.7 | 1256.9 | 1407.9 | 2160.3 |
| O09111 | ***Ndufb11*** | 724.6 | 466.8 | 383.6 | 1662.6 | 1813.5 | 1507.3 | 1817.2 | 1120.3 | 1025.5 |
| Q91VN4 | ***Chchd6*** | 196.9 | 202.1 | 328.1 | 2968.5 | 3463.1 | 2356 | 3047.6 | 1256.1 | 1405.7 |
| Q9QXV0 | ***Pcsk1n*** | 32.1 | 52.7 | 56.9 | 245.4 | 226.2 | 307.1 | 375.7 | 1451.4 | 596.1 |
| P62897 | ***Cycs*** | 3443.8 | 3043.6 | 1199.5 | 12077.4 | 10020.5 | 9955.1 | 8141.7 | 9296.8 | 5757 |
| Q9CZU6 | ***Cs*** | 1049.8 | 1078.3 | 939.2 | 7693.1 | 5751.4 | 8208.4 | 5673.8 | 5893.5 | 5386.2 |
| Q64332 | ***Syn2*** | 23.3 | 35.4 | 47.3 | 544.9 | 721 | 815.9 | 736.9 | 1287 | 1548.5 |
| P54227 | ***Stmn1*** | 5.7 | 22 | 0 | 217.3 | 308 | 226.8 | 250.7 | 638.6 | 406 |
| P01831 | ***Thy1*** | 47.1 | 97.8 | 159.7 | 604.9 | 600.4 | 466.7 | 999.2 | 1276.6 | 1546 |
| Q9DCS9 | ***Ndufb10*** | 1165.8 | 648.2 | 406.1 | 2877.8 | 3262.7 | 2735 | 3160.3 | 1814.7 | 1912.7 |
| Q91VD9 | ***Ndufs1*** | 4639.2 | 3135.9 | 1356.3 | 12234.8 | 13585.3 | 11277.6 | 13691.5 | 8390.3 | 8526.8 |
| Q91V61 | ***Sfxn3*** | 35 | 68.6 | 232.1 | 3140.4 | 2623.3 | 2076.6 | 1706.3 | 997.4 | 896.6 |
| P99028 | ***Uqcrh*** | 588.8 | 530 | 296.1 | 2832.9 | 2240 | 2102.9 | 2775.4 | 1811.5 | 1608.6 |
| P56480 | ***Atp5b*** | 5130.5 | 4053.7 | 1293 | 12293.8 | 13170.8 | 13261.8 | 12225.7 | 9888.7 | 8628 |
| Q91XR9 | ***Gpx4*** | 92.9 | 92 | 132 | 264.6 | 316.9 | 262.8 | 341.7 | 200.7 | 283.4 |
| Q03265 | ***Atp5a1*** | 9042.4 | 7436.7 | 3014.3 | 25272.1 | 26255.1 | 21681.4 | 26106.6 | 17412.6 | 16334 |
| Q9JLZ3 | ***Auh*** | 574.3 | 542.7 | 375.8 | 4799.5 | 4402.8 | 3254.7 | 4695.2 | 3942.3 | 2980.5 |
| P47934 | ***Crat*** | 62.8 | 72.6 | 0 | 362.8 | 399.1 | 290.8 | 433.1 | 257.9 | 299.7 |
| P56593 | ***Cyp2a12*** | 921.2 | 1358.2 | 972.7 | 13.4 | 14.9 | 19.2 | 10.9 | 18.4 | 26 |
| Q01768 | ***Nme2*** | 125.5 | 193.9 | 178.9 | 422.9 | 469.3 | 541.5 | 396 | 1004.5 | 863.8 |
| P47963 | ***Rpl13*** | 766.9 | 993.8 | 717.2 | 194.5 | 142.4 | 135.2 | 153.3 | 99.9 | 130.7 |
| P97872 | ***Fmo5*** | 1403.7 | 2231.1 | 1912.8 | 19 | 29.6 | 18 | 26.1 | 15.6 | 36.7 |
| Q8BMF3 | ***Me3*** | 46.5 | 64.7 | 109.8 | 781.1 | 759.4 | 1083.9 | 451.3 | 334.1 | 338.4 |
| P70404 | ***Idh3g*** | 140.3 | 99.5 | 172 | 1442 | 1701.7 | 2183.5 | 1372.1 | 812.8 | 1097.3 |

| | | | | | | | | | | |
|---|---|---|---|---|---|---|---|---|---|---|
| P97807 | ***Fh*** | 2948 | 1699.7 | 994.6 | 7328 | 7528.2 | 6261.2 | 6945.4 | 5975.9 | 4153.6 |
| Q99LY9 | ***Ndufs5*** | 552.1 | 143.1 | 273.2 | 2316 | 2406.3 | 1724.5 | 2418.2 | 1538.7 | 1401.2 |
| Q8BIJ6 | ***Iars2*** | 444.8 | 296.4 | 565 | 1002.4 | 1035.7 | 1006.5 | 1135.1 | 592.1 | 875.4 |
| P46460 | ***Nsf*** | 100 | 246.9 | 656 | 1776.6 | 1664.1 | 2057 | 2436.1 | 2803.6 | 4568.1 |
| Q8BP92 | ***Rcn2*** | 76.5 | 54.2 | 59.4 | 324.3 | 235.8 | 334.4 | 346.2 | 744.9 | 524.1 |
| Q60930 | ***Vdac2*** | 1135.6 | 1141.8 | 673.2 | 5197.7 | 6597 | 4599.4 | 7441.9 | 3162.1 | 4999.2 |
| Q791T5 | ***Mtch1*** | 110.4 | 131.8 | 113 | 308.1 | 308 | 242.4 | 331.7 | 174.4 | 250.3 |
| P41216 | ***Acsl1*** | 4128.8 | 6407.1 | 4891.4 | 354.9 | 334 | 194.2 | 335.3 | 170.2 | 192.2 |
| P28663 | ***Napb*** | 11.6 | 0 | 32.7 | 175.7 | 121.5 | 169 | 134.9 | 166.3 | 231 |
| Q922H2 | ***Pdk3*** | 16.3 | 22.2 | 101.2 | 499.2 | 621.1 | 418.3 | 466.2 | 295.2 | 355.6 |
| P20108 | ***Prdx3*** | 819.9 | 666 | 497.8 | 1767.4 | 1747.9 | 2202.9 | 1956.7 | 1650.3 | 1790.5 |
| Q9CPP6 | ***Ndufa5*** | 1114 | 514.4 | 379.5 | 2464.3 | 2849.2 | 2375.3 | 1801.1 | 1825.8 | 1548.7 |
| Q9QZD8 | ***Slc25a10*** | 490.6 | 348.6 | 396.8 | 131.7 | 111.5 | 103.5 | 170.8 | 118 | 111 |
| P58281 | ***Opa1*** | 1755.9 | 1074.7 | 659.6 | 4840.1 | 5800.5 | 4271.4 | 4667.9 | 2424.1 | 2851 |
| P43024 | ***Cox6a1*** | 198 | 88.3 | 120.3 | 663.6 | 759.8 | 528.9 | 652.6 | 462.8 | 431.5 |
| Q64516 | ***Gk*** | 132.9 | 156.8 | 138.9 | 1330.8 | 1612.7 | 1022.4 | 2033.7 | 1126.8 | 1383.4 |
| Q9D6J6 | ***Ndufv2*** | 1081.4 | 568.6 | 563.6 | 2812 | 2990.5 | 2248.5 | 3059.3 | 1925.4 | 1853.4 |
| Q9CQV8 | ***Ywhab*** | 92.8 | 252.3 | 233.5 | 668.3 | 917.5 | 853.3 | 822.9 | 1857.3 | 1819.7 |
| Q9JKC6 | ***Cend1*** | 6.4 | 20.8 | 216.1 | 4122.2 | 4024.4 | 2555.3 | 4482.9 | 3641.4 | 2701.7 |
| P62743 | ***Ap2s1*** | 29.7 | 59.3 | 7.9 | 149.2 | 150.2 | 130 | 119.7 | 323.9 | 235.2 |
| Q9CRB6 | ***Tppp3*** | 44 | 89 | 71.5 | 332.6 | 414.2 | 505.2 | 701.8 | 2492.2 | 1625.4 |
| Q60932 | ***Vdac1*** | 2013.3 | 2280.8 | 1467 | 15684.4 | 19023.1 | 12072.7 | 15025 | 9949 | 9627 |
| O08599 | ***Stxbp1*** | 82.9 | 150.1 | 300.2 | 3044.7 | 3912.2 | 2425 | 3717.3 | 8888.7 | 6636.9 |
| Q8K0T0 | ***Rtn1*** | 25.2 | 110.8 | 55.1 | 306.3 | 287.3 | 240.3 | 158.7 | 334 | 310 |
| Q9DCT2 | ***Ndufs3*** | 945.5 | 494.6 | 269.9 | 3708.1 | 3853.4 | 2669.6 | 3856.6 | 2373.9 | 2261.6 |
| P56391 | ***Cox6b1*** | 2197.2 | 1215.1 | 1059 | 4744.8 | 4903.6 | 3983.2 | 5059.6 | 3481 | 2982.5 |
| Q5M8N4 | ***Sdr39u1*** | 85.7 | 69.1 | 144.2 | 568.8 | 669.8 | 444.5 | 595.4 | 487.8 | 449.3 |
| P50518 | ***Atp6v1e1*** | 95.2 | 397.6 | 278.8 | 1225.3 | 940.8 | 1033.2 | 995.2 | 2588.6 | 1761.8 |
| Q8BVE3 | ***Atp6v1h*** | 20.5 | 127.4 | 54.1 | 309.8 | 267.3 | 306.8 | 290.1 | 610.2 | 588 |
| P60879 | ***Snap25*** | 18.3 | 51 | 86.8 | 676 | 733.1 | 455 | 690.8 | 2004.3 | 1065 |
| P61264 | ***Stx1b*** | 57.2 | 153.2 | 245 | 2433.5 | 3132.8 | 1911 | 3431.1 | 8749 | 5315.6 |
| Q9D6F9 | ***Tubb4a*** | 24.2 | 37 | 8.1 | 300.5 | 498.7 | 397 | 500.6 | 919.4 | 1304.1 |
| Q91YT0 | ***Ndufv1*** | 1568.6 | 1059.2 | 535.1 | 4579.5 | 4227.3 | 3377.2 | 4980.7 | 2735.8 | 3107.2 |
| Q9D1H6 | ***Ndufaf4*** | 80 | 68.7 | 105.8 | 207.8 | 203.1 | 167.5 | 182.9 | 153.2 | 140.2 |
| Q8K3J1 | ***Ndufs8*** | 1203.4 | 693 | 369.2 | 2433.5 | 2828.9 | 2397.5 | 2639.6 | 1636.1 | 1667.6 |
| Q9Z1S5 | ***42981*** | 9.7 | 30.1 | 52.4 | 546.1 | 620.2 | 371 | 461.2 | 1048.8 | 616.8 |
| P62073 | ***Timm10*** | 457.1 | 237.3 | 72.9 | 1373.1 | 1104.8 | 1084.6 | 830.9 | 731.6 | 527.6 |
| O08553 | ***Dpysl2*** | 227.5 | 418.2 | 509.7 | 2867.4 | 4693.6 | 3877.5 | 4232.7 | 11423.2 | 8600.6 |
| P62761 | ***Vsnl1*** | 32.1 | 44.4 | 39.7 | 344.3 | 565.7 | 440.6 | 563.4 | 2158.4 | 1180.9 |
| Q9CZ13 | ***Uqcrc1*** | 3804.5 | 2831 | 1273.1 | 7352.5 | 7367.7 | 8119.3 | 7254.4 | 5246.6 | 4776 |
| Q8BK30 | ***Ndufv3*** | 422.3 | 241.9 | 180.1 | 1056 | 1098.6 | 825.4 | 1062.5 | 695.7 | 546.3 |
| Q91VM9 | ***Ppa2*** | 430.1 | 262.6 | 295.9 | 715.6 | 745.4 | 873.6 | 826.1 | 704.1 | 672.2 |
| P00920 | ***Ca2*** | 45.9 | 24.8 | 36.9 | 179.6 | 286.2 | 249.9 | 277 | 855.8 | 661.9 |
| P61982 | ***Ywhag*** | 20.4 | 81.4 | 127.3 | 280.4 | 342 | 363.4 | 529.1 | 1174.2 | 1062.9 |
| Q9CQ75 | ***Ndufa2*** | 758.3 | 339 | 155.2 | 1902 | 2131.8 | 1609.5 | 1503.4 | 1137.1 | 1074.6 |
| P50285 | ***Fmo1*** | 664.5 | 1159.4 | 1056.6 | 28.2 | 17.9 | 15.3 | 20.8 | 16.1 | 20.7 |
| Q62452 | ***Ugt1a9*** | 3051.3 | 4621.8 | 2857 | 33.6 | 58.8 | 49.8 | 56.5 | 23.1 | 44.8 |

| | | | | | | | | | | |
|---|---|---|---|---|---|---|---|---|---|---|
| P56393 | ***Cox7b*** | 234.7 | 104.3 | 72.7 | 463.8 | 465.1 | 428.8 | 568.9 | 334.1 | 222.6 |
| P12960 | ***Cntn1*** | 24.8 | 15.2 | 53.4 | 254.9 | 274.1 | 172.6 | 323.5 | 539.1 | 370.5 |
| Q9JIS5 | ***Sv2a*** | 14.9 | 14.5 | 37.1 | 177.1 | 255.8 | 299.6 | 582.2 | 478.6 | 1161.3 |
| Q61102 | ***Abcb7*** | 153.3 | 121.3 | 82.4 | 239.8 | 243.4 | 246.8 | 225.8 | 193.5 | 188.8 |
| Q9QYR6 | ***Map1a*** | 30.6 | 58.2 | 0 | 208.4 | 279.9 | 188.4 | 369.8 | 489.6 | 571.7 |
| Q925N0 | ***Sfxn5*** | 216.1 | 176.2 | 186.2 | 2433.6 | 2635.7 | 1547.7 | 2966.5 | 2232 | 2385.2 |
| O88741 | ***Gdap1*** | 88.9 | 301.1 | 232.7 | 1663.7 | 1395.5 | 1036 | 1533.9 | 1281.7 | 1077 |
| Q9D6J5 | ***Ndufb8*** | 401.8 | 224 | 296.7 | 1297.6 | 1171.1 | 878.2 | 1292.2 | 843.7 | 804.5 |
| Q9Z1G4 | ***Atp6v0a1*** | 73.9 | 390.8 | 450.3 | 1036.3 | 1016.4 | 981.8 | 1530.1 | 2496.3 | 2397.3 |
| O88712 | ***Ctbp1*** | 22.3 | 25.1 | 0 | 74 | 109.3 | 99.2 | 97.7 | 253.3 | 197.8 |
| P52503 | ***Ndufs6*** | 1316 | 657.8 | 359.2 | 2700 | 2905.7 | 2405.9 | 2915.6 | 2015.1 | 1825.4 |
| Q99KI0 | ***Aco2*** | 3294.3 | 2173.7 | 2226.2 | 17245.4 | 18320.8 | 26894.7 | 14054.8 | 13570.5 | 13254 |
| Q8R127 | ***Sccpdh*** | 15.7 | 32.2 | 11.2 | 330.8 | 324.3 | 195.4 | 411.4 | 440.9 | 441.9 |
| Q3UMR5 | ***Mcu*** | 145.4 | 178.8 | 200.1 | 1333 | 1284.5 | 816.9 | 945.4 | 486.3 | 528.8 |
| Q9ERD7 | ***Tubb3*** | 18 | 49.2 | 36.8 | 239.7 | 409.9 | 334.6 | 450 | 925.3 | 924.2 |
| P48962 | ***Slc25a4*** | 145.8 | 221.6 | 426.7 | 4926.4 | 3153.5 | 3164.2 | 5047.6 | 3059.8 | 3948.1 |
| O88935 | ***Syn1*** | 533.2 | 588.2 | 404.6 | 2311.2 | 3755.2 | 3636.4 | 4128.4 | 9425.3 | 8305 |
| P56382 | ***Atp5e*** | 720.8 | 375.5 | 22 | 1740 | 1492.8 | 1605.8 | 1131 | 1351.1 | 917.9 |
| Q8BH59 | ***Slc25a12*** | 1263.1 | 1422.7 | 2443.2 | 9740.2 | 7420.4 | 6369.4 | 7425 | 4309.4 | 4644 |
| Q8BWM0 | ***Ptges2*** | 76.6 | 53.3 | 88.7 | 157.4 | 168.5 | 136.3 | 136.8 | 96.2 | 122.9 |
| O55131 | ***42985*** | 44.9 | 200.9 | 224.3 | 623.5 | 771.3 | 550.8 | 620.2 | 1360.5 | 922.4 |
| Q60931 | ***Vdac3*** | 1047.6 | 903.8 | 658.5 | 4046.4 | 4176.6 | 2700.2 | 6510.8 | 2758.1 | 3805.5 |
| Q9CQJ8 | ***Ndufb9*** | 618.7 | 362.5 | 276 | 1459.2 | 1438.5 | 1084.7 | 1438.3 | 926.2 | 877.2 |
| P31650 | ***Slc6a11*** | 18.9 | 33.3 | 34.8 | 422.8 | 690.1 | 770.6 | 1459.5 | 2957.9 | 2776.3 |
| P63011 | ***Rab3a*** | 87.1 | 162 | 451.2 | 1030.6 | 1221.5 | 905.2 | 1459.7 | 2863.9 | 3131.4 |
| P26645 | ***Marcks*** | 20.3 | 81.4 | 26 | 166.4 | 218.4 | 239.3 | 121.6 | 331.8 | 203.7 |
| Q9CWZ7 | ***Napg*** | 140.4 | 581.8 | 519.4 | 1456.2 | 1266.3 | 1712.4 | 1123.6 | 1773.7 | 2174.3 |
| P62814 | ***Atp6v1b2*** | 146.7 | 806.9 | 498.7 | 1676.3 | 1510.7 | 1828.7 | 1695.9 | 3353.4 | 3367.4 |
| O88441 | ***Mtx2*** | 126.6 | 84.9 | 0 | 320.6 | 387.5 | 290.6 | 313 | 155.3 | 181.5 |
| Q9D6M3 | ***Slc25a22*** | 1454.5 | 877.4 | 804.8 | 4672.5 | 3640.9 | 3131 | 2370.4 | 1199.2 | 1564.5 |
| Q8BWT1 | ***Acaa2*** | 21916.7 | 12838.5 | 15002 | 1108.8 | 1734 | 1329.4 | 1673.8 | 1077.8 | 1747 |
| Q7TNG8 | ***Ldhd*** | 1166.4 | 651.3 | 883.9 | 76 | 82.1 | 70.9 | 87.6 | 80.5 | 72.5 |
| Q03517 | ***Scg2*** | 31.2 | 34.1 | 82.8 | 211.7 | 170.2 | 253.9 | 283.9 | 1286.1 | 488.8 |
| Q99PJ0 | ***Ntm*** | 8.6 | 37.1 | 12.9 | 349.6 | 451.9 | 246.9 | 260.6 | 788.2 | 445.2 |
| O08539 | ***Bin1*** | 9.9 | 28.1 | 42.1 | 248.2 | 454.2 | 373.7 | 452.7 | 992.6 | 892 |
| P05201 | ***Got1*** | 12.8 | 75.3 | 14.9 | 162.1 | 213.1 | 244 | 146.3 | 573.9 | 560.4 |
| Q9Z1J3 | ***Nfs1*** | 179.5 | 138.8 | 92.7 | 292.9 | 356.2 | 294.4 | 347.5 | 199.1 | 235.6 |
| Q62442 | ***Vamp1*** | 14.5 | 22.7 | 40.3 | 117.6 | 123.5 | 179.2 | 255.2 | 686.8 | 568.8 |
| Q8BFR5 | ***Tufm*** | 5118.6 | 2830.4 | 6834.4 | 10998.6 | 13389 | 12122 | 11157.2 | 7069.7 | 7932.5 |
| Q9CZ42 | ***Naxd*** | 27.5 | 28.1 | 16.5 | 101.2 | 125.2 | 168.8 | 94.4 | 87.9 | 97.5 |
| P13595 | ***Ncam1*** | 40.5 | 110.6 | 206.3 | 660.2 | 862.6 | 535.7 | 770.3 | 1996.5 | 1415.2 |
| Q9DC70 | ***Ndufs7*** | 1021.6 | 630.3 | 259.4 | 2407.2 | 2306 | 1801.1 | 2629.9 | 1469.2 | 1732.7 |
| P39053 | ***Dnm1*** | 97.9 | 193.7 | 356.8 | 1604.9 | 2900.5 | 2211.3 | 2284.9 | 4967.9 | 5542 |
| Q9R0H0 | ***Acox1*** | 651.5 | 888 | 500.9 | 73.5 | 90.6 | 84.2 | 87.7 | 96.9 | 105.8 |
| Q8JZQ2 | ***Afg3l2*** | 2097.5 | 1328.1 | 1105 | 3009.9 | 3613.4 | 3589 | 3253.6 | 1735.8 | 2330.2 |
| O08585 | ***Clta*** | 108.7 | 340.4 | 174.8 | 622.2 | 607.9 | 780 | 1427.4 | 2096.5 | 2362.5 |
| Q9CQ54 | ***Ndufc2*** | 1114.1 | 738 | 364.8 | 3190.4 | 2864.7 | 2131.6 | 2007.1 | 1913.1 | 1363.7 |

| | | | | | | | | | | |
|---|---|---|---|---|---|---|---|---|---|---|
| P05063 | ***Aldoc*** | 54.9 | 91.6 | 135 | 391.4 | 399.3 | 595.4 | 511.6 | 2234.9 | 1338.5 |
| Q9CQZ5 | ***Ndufa6*** | 1186.7 | 517.6 | 419.8 | 4386.3 | 4631.4 | 2798.4 | 3733 | 3131.3 | 2365.6 |
| Q8R404 | ***Mic13*** | 207.3 | 98.1 | 29.8 | 452.7 | 653.3 | 485.1 | 412.7 | 383.3 | 318.9 |
| Q9CQZ6 | ***Ndufb3*** | 645.3 | 251.8 | 463 | 1437.6 | 1559.4 | 1106.6 | 1503.8 | 921.4 | 1004.3 |
| P97427 | ***Crmp1*** | 51.6 | 110.2 | 300.3 | 808.4 | 1306.6 | 938.2 | 1100.5 | 2835.9 | 1919.2 |
| Q99P72 | ***Rtn4*** | 23.4 | 45.1 | 0 | 139.3 | 129.4 | 93.7 | 120.4 | 182.7 | 198.2 |
| O35633 | ***Slc32a1*** | 18.7 | 29.9 | 19.2 | 131.7 | 227.4 | 240.9 | 347.1 | 872 | 930.5 |
| Q9CQC7 | ***Ndufb4*** | 596.2 | 366.8 | 181.2 | 1934.4 | 1741.4 | 1218.7 | 1820.4 | 1305.9 | 1027.8 |
| Q7TMF3 | ***Ndufa12*** | 856.3 | 521.4 | 419.5 | 2320.1 | 2363.3 | 1564 | 2257.8 | 1728.9 | 1301.8 |
| Q9Z1G3 | ***Atp6v1c1*** | 22.9 | 98.6 | 26.1 | 296.9 | 239.1 | 195.9 | 216.8 | 569.6 | 455.5 |
| Q9Z2Q6 | ***42983*** | 37.9 | 55.3 | 53.3 | 905.1 | 1349.6 | 723.7 | 794.7 | 1690.1 | 1205.5 |
| P14869 | ***Rplp0*** | 689.5 | 834.5 | 969.7 | 449.6 | 280.2 | 297.9 | 418.2 | 327.6 | 362.7 |
| Q9CR21 | ***Ndufab1*** | 1178.5 | 468.5 | 1814 | 3343.4 | 3326.7 | 2949.2 | 2126.6 | 2477.7 | 1419.1 |
| Q9R1Q8 | ***Tagln3*** | 4.4 | 10.6 | 48.6 | 129.5 | 203.8 | 230.8 | 178.3 | 734.5 | 429.9 |
| Q9CR68 | ***Uqcrfs1*** | 3031.4 | 1410.3 | 609.8 | 5822.3 | 8447.2 | 6593.1 | 6843.8 | 4803.8 | 5231.2 |
| Q9DB20 | ***Atp5o*** | 3296.9 | 1709.7 | 618.2 | 7898.7 | 7943.9 | 5737 | 6755.2 | 5905.6 | 4275.1 |
| P16125 | ***Ldhb*** | 25.9 | 58.5 | 42.4 | 501.6 | 768.6 | 997.3 | 572.8 | 2074.3 | 2067.5 |
| Q9CQH3 | ***Ndufb5*** | 1617.2 | 963 | 280 | 3804.3 | 3944.9 | 2829.9 | 3386.2 | 2350.1 | 2030.9 |
| P08553 | ***Nefm*** | 35.1 | 54.7 | 49.3 | 241.1 | 457.1 | 397.8 | 1068.2 | 702.1 | 1474.7 |
| P12787 | ***Cox5a*** | 1574.2 | 393.9 | 174.9 | 3080.2 | 3867.6 | 2909.5 | 2862.1 | 2461.6 | 2116.4 |
| Q9CXZ1 | ***Ndufs4*** | 1937.5 | 823.6 | 270.5 | 4258.6 | 4756.5 | 3362.7 | 4683 | 3078.4 | 2693.4 |
| Q9DCJ5 | ***Ndufa8*** | 1771.6 | 711.7 | 334.7 | 3108.9 | 3359 | 2913.5 | 3263.4 | 1927.7 | 1924.3 |
| Q8C1B7 | ***42989*** | 30.5 | 75.7 | 237.3 | 528.9 | 712.1 | 479 | 573.9 | 1127.6 | 907.6 |
| Q9CQ91 | ***Ndufa3*** | 413.9 | 149.8 | 57.3 | 1458.4 | 1363.3 | 865.3 | 768.6 | 1107.9 | 716 |
| P00186 | ***Cyp1a2*** | 414.7 | 734.9 | 790.6 | 85.7 | 82.8 | 86.1 | 69.3 | 52.7 | 63.6 |
| Q9DCB8 | ***Isca2*** | 50.6 | 31.6 | 0 | 105.9 | 111.7 | 147.5 | 105.1 | 78.8 | 85.5 |
| P63101 | ***Ywhaz*** | 98.5 | 270.7 | 279.5 | 483 | 640.8 | 602 | 354 | 1035 | 783.6 |
| Q61578 | ***Fdxr*** | 93.2 | 49.3 | 22.3 | 155.8 | 160.4 | 146.7 | 147 | 131.5 | 125.2 |
| Q8BGH2 | ***Samm50*** | 233.5 | 219.6 | 70.3 | 638.3 | 767.3 | 484.9 | 720.1 | 308.5 | 524.1 |
| Q9CR61 | ***Ndufb7*** | 1016 | 518.2 | 295.6 | 1543.2 | 1930 | 1749.1 | 1457.3 | 1131.8 | 1081 |
| P52480 | ***Pkm*** | 179.6 | 594.1 | 669.4 | 1445.4 | 1819 | 2310.5 | 1670.9 | 4703.4 | 4208.1 |
| Q62420 | ***Sh3gl2*** | 77.9 | 130.9 | 251.5 | 765.3 | 1461.4 | 1334.5 | 994.4 | 2745 | 2517.1 |
| Q9WV55 | ***Vapa*** | 2129.8 | 1708 | 1438.8 | 910.1 | 634 | 581.4 | 601.1 | 630.5 | 735 |
| Q9DCS3 | ***Mecr*** | 325.9 | 131.5 | 137.6 | 484.7 | 504.6 | 577 | 473 | 377.9 | 414.1 |
| Q7TQF7 | ***Amph*** | 28.2 | 42.1 | 89.3 | 236.4 | 433.4 | 323.4 | 371.3 | 817 | 715.2 |
| P53395 | ***Dbt*** | 4565.8 | 3751.4 | 2959.4 | 1340.8 | 1804.8 | 1142.8 | 1521.4 | 1078.6 | 1201.5 |
| P0DN34 | ***Ndufb1*** | 567.5 | 93.3 | 100.5 | 1029.5 | 1107.5 | 916.3 | 419.7 | 763.9 | 474.4 |
| Q61598 | ***Gdi2*** | 20.2 | 107.3 | 34.7 | 171.6 | 224.5 | 246 | 179.9 | 531.9 | 351.3 |
| P01027 | ***C3*** | 208.2 | 402.7 | 237.3 | 9.6 | 6.1 | 7.7 | 18.7 | 9.9 | 24.2 |
| Q9D2R6 | ***Coa3*** | 156.7 | 62.9 | 60.5 | 318.1 | 382 | 250.8 | 228.4 | 212.9 | 163.3 |
| Q9Z1P6 | ***Ndufa7*** | 2863.7 | 2086.7 | 891.6 | 7878.7 | 8089.5 | 5088.5 | 7423.6 | 5036 | 3916.5 |
| Q921I1 | ***Tf*** | 6127.6 | 6420 | 3088.7 | 514.9 | 259.4 | 435.4 | 461.5 | 722.1 | 862.2 |
| Q9ERS2 | ***Ndufa13*** | 1087.3 | 653.7 | 435.6 | 3137.3 | 2934.6 | 1895.9 | 2834.3 | 2009 | 1549.2 |
| Q63886 | ***Ugt1a1*** | 1062.9 | 2435.4 | 1899.4 | 1.5 | 8.7 | 11.3 | 3.7 | 12.5 | 20.3 |
| Q9DBG1 | ***Cyp27a1*** | 3595.6 | 2429.9 | 1644.6 | 45.7 | 47.5 | 24.8 | 70.5 | 21.8 | 69.1 |
| Q9CQQ7 | ***Atp5f1*** | 5311.9 | 4546.1 | 1606.9 | 11033.2 | 10199.2 | 8421.8 | 9842.8 | 7631.1 | 6244.3 |
| Q3UIU2 | ***Ndufb6*** | 100.2 | 81.8 | 36.2 | 317.7 | 344.6 | 199 | 341.8 | 174.1 | 163.5 |

| | | | | | | | | | | |
|---|---|---|---|---|---|---|---|---|---|---|
| P05202 | ***Got2*** | 3773.8 | 1965.5 | 1500.2 | 11294.4 | 6675.1 | 10984.7 | 5963.9 | 7325.1 | 5526.2 |
| Q9DB77 | ***Uqcrc2*** | 3106.9 | 2085.2 | 1147.4 | 9037.4 | 8932.7 | 5453.9 | 9045.3 | 6315.6 | 5222.6 |
| Q9D855 | ***Uqcrb*** | 3609.6 | 1479.7 | 753.1 | 5953.5 | 6579.2 | 5382 | 5723.1 | 4894.9 | 3778.4 |
| P51881 | ***Slc25a5*** | 4429.2 | 3500.6 | 4649.3 | 7384.9 | 6227.7 | 6069.2 | 8186.6 | 5240 | 8865.8 |
| O88587 | ***Comt*** | 1215.7 | 1904.5 | 931.4 | 92.6 | 100 | 111.8 | 96.9 | 116.5 | 131.4 |
| Q922B1 | ***Macrod1*** | 130.4 | 76.1 | 70.7 | 193.2 | 222.9 | 282.9 | 210.3 | 123.7 | 139.9 |
| Q91VR2 | ***Atp5c1*** | 3976.6 | 2584.9 | 1234.9 | 8636.6 | 8006.7 | 5911 | 9349.1 | 5025.3 | 5208.2 |
| Q61335 | ***Bcap31*** | 797.9 | 869.1 | 463.3 | 213.6 | 136.8 | 126.1 | 169.6 | 261.3 | 198.2 |
| Q9CYH2 | ***Fam213a*** | 194.7 | 238.3 | 60 | 437.6 | 518.6 | 387.4 | 432.6 | 538.1 | 430.8 |
| P16330 | ***Cnp*** | 296.4 | 456.5 | 488.7 | 2481.9 | 3807 | 1990.9 | 5161.2 | 8479.5 | 8845.2 |
| Q9DC69 | ***Ndufa9*** | 1294.7 | 915.2 | 558.3 | 5333.3 | 4413.9 | 2816 | 4521.6 | 2954.1 | 2654.1 |
| P47754 | ***Capza2*** | 14.5 | 56.7 | 55.1 | 105.4 | 151.7 | 118.5 | 115.9 | 226.7 | 164.8 |
| Q9DCX2 | ***Atp5h*** | 5757.9 | 2876.7 | 1643.7 | 9106.7 | 9397.7 | 8103.9 | 8520.7 | 7287.5 | 5209.2 |
| P47857 | ***Pfkm*** | 7.4 | 41.7 | 49.4 | 150.7 | 294.4 | 304.9 | 318.9 | 648.4 | 853.3 |
| P46660 | ***Ina*** | 54 | 102 | 73.1 | 244.7 | 482.7 | 411 | 660.7 | 487.2 | 911.4 |
| Q59J78 | ***Ndufaf2*** | 706.9 | 386.8 | 235.6 | 1036.6 | 1294.4 | 1049.2 | 1523.1 | 1154.4 | 989.5 |
| Q9D0M3 | ***Cyc1*** | 3167.5 | 2304.3 | 942.2 | 6857.2 | 6187.5 | 4735.3 | 6410.1 | 4508.2 | 4032.6 |
| Q05421 | ***Cyp2e1*** | 2643.3 | 6119.4 | 3978.9 | 10.2 | 31.2 | 31.4 | 39.4 | 16.5 | 41.5 |
| P47740 | ***Aldh3a2*** | 1087.8 | 1542.6 | 866.1 | 360.1 | 272.7 | 353.1 | 348.4 | 558.3 | 464.2 |
| P17751 | ***Tpi1*** | 66.8 | 286.7 | 235.4 | 440.3 | 547 | 585.3 | 584 | 1299.8 | 1165.8 |
| P08113 | ***Hsp90b1*** | 9877.3 | 12337.4 | 6066.8 | 2275.5 | 1032.8 | 1902 | 1678.9 | 1377 | 1671.9 |
| P06837 | ***Gap43*** | 2.4 | 32.7 | 96.2 | 877.8 | 1209.1 | 521.9 | 634 | 2128.3 | 902.5 |
| P19783 | ***Cox4i1*** | 1396.8 | 839.2 | 367.5 | 3136.5 | 3409.5 | 2125.8 | 3587.5 | 2429.7 | 2151.9 |
| Q9CR62 | ***Slc25a11*** | 402.9 | 338.2 | 281.8 | 2009.4 | 1526.9 | 1022.7 | 1693.9 | 942.7 | 1210.9 |
| Q9CWX2 | ***Ndufaf1*** | 253.8 | 173.7 | 100.6 | 374.5 | 407.2 | 335.3 | 345.2 | 265.2 | 248.9 |
| Q8JZR0 | ***Acsl5*** | 1022.5 | 1955.2 | 2539.2 | 52.8 | 100.4 | 82.3 | 65.8 | 74 | 74.9 |
| P62880 | ***Gnb2*** | 52.3 | 224.5 | 306.5 | 676.6 | 964.4 | 580.6 | 852.8 | 1721.7 | 1276.3 |
| Q9CXT8 | ***Pmpcb*** | 429.2 | 244.6 | 37.7 | 741.9 | 756.3 | 634.2 | 719.5 | 451.7 | 450.3 |
| Q62425 | ***Ndufa4*** | 2798.9 | 1491 | 484.1 | 6727.9 | 6713.6 | 4148.6 | 5361.2 | 5139.5 | 3393.5 |
| Q80TJ1 | ***Cadps*** | 19.8 | 26.7 | 0 | 63.2 | 110 | 134.1 | 132.2 | 286.5 | 439.9 |
| Q9DB15 | ***Mrpl12*** | 944.8 | 560.8 | 399.5 | 1404.1 | 1589.8 | 1202.6 | 990.3 | 891.4 | 532.5 |
| P41105 | ***Rpl28*** | 465.3 | 569.1 | 925.2 | 135 | 87 | 107.2 | 106 | 70 | 109.1 |
| P62918 | ***Rpl8*** | 608.4 | 1383.1 | 1076.9 | 196.2 | 118 | 119.2 | 135.6 | 106 | 136.7 |
| P05064 | ***Aldoa*** | 143.3 | 1144.5 | 590.3 | 1678.4 | 2758.3 | 2631.8 | 1646 | 6198.9 | 3985.1 |
| P07759 | ***Serpina3k*** | 395.8 | 645.9 | 281.8 | 22 | 19.4 | 37.1 | 55.6 | 50.1 | 119.7 |
| Q64133 | ***Maoa*** | 654.5 | 884.3 | 838.8 | 2337.6 | 2101.2 | 1373.9 | 3595.2 | 3021.7 | 2955.8 |
| Q9CPQ1 | ***Cox6c*** | 2496.9 | 904.2 | 584.2 | 4717.7 | 5101.7 | 3322 | 2747.1 | 3166.6 | 2466 |
| Q9WV92 | ***Epb41l3*** | 29.8 | 73.1 | 18.2 | 110.9 | 149.1 | 106.4 | 151.6 | 353.5 | 229.3 |
| Q00897 | ***Serpina1d*** | 785.3 | 1274.4 | 525.7 | 31 | 21.9 | 32.7 | 50 | 57 | 118.2 |
| Q9QXE0 | ***Hacl1*** | 336.8 | 665.4 | 898.7 | 11.6 | 15.9 | 18.8 | 16.8 | 15.5 | 31.7 |
| Q9CRB8 | ***Mtfp1*** | 50 | 42.1 | 33 | 274.9 | 211.6 | 126.4 | 108.3 | 137.7 | 107.9 |
| P97742 | ***Cpt1a*** | 2500.4 | 2090 | 1063.2 | 296.1 | 319.7 | 239.2 | 571.5 | 386.7 | 527.5 |
| P48771 | ***Cox7a2*** | 631 | 251.1 | 108.6 | 1696.9 | 2119.9 | 1055.8 | 935.8 | 1227.7 | 854.4 |
| Q8VEM8 | ***Slc25a3*** | 2323.1 | 2368.8 | 1105.3 | 5976.9 | 4715.2 | 3701.1 | 5592.5 | 3427.2 | 3907.5 |
| Q921H8 | ***Acaa1a*** | 1143.9 | 2928.7 | 2496.1 | 224.4 | 194.1 | 229.5 | 223.2 | 216.4 | 290.7 |
| P18872 | ***Gnao1*** | 34.3 | 89.9 | 111.9 | 1252.2 | 1041 | 474.6 | 944.1 | 2161.3 | 1263.4 |
| P08551 | ***Nefl*** | 142 | 121.9 | 338 | 472.5 | 891 | 815.3 | 2401.1 | 1264.3 | 3098.5 |

| | | | | | | | | | | |
|---|---|---|---|---|---|---|---|---|---|---|
| Q64458 | ***Cyp2c29*** | 876.9 | 2035.3 | 2335.6 | 112.2 | 114.8 | 216.8 | 61.4 | 86.7 | 79.8 |
| Q9CPQ8 | ***Atp5l*** | 1137.6 | 414.8 | 324.1 | 3902.2 | 3494.3 | 1830.4 | 1526.2 | 3086.8 | 1558.3 |
| Q9QYA2 | ***Tomm40*** | 264.7 | 261.7 | 72.7 | 630.5 | 775.6 | 440 | 621.9 | 312.2 | 405 |
| Q9CQ69 | ***Uqcrq*** | 1665.2 | 817.2 | 471.9 | 3102 | 2939.9 | 2058.1 | 2055.2 | 1855.8 | 1584.4 |
| P21460 | ***Cst3*** | 18.4 | 28.2 | 26.5 | 126.2 | 75.7 | 172.8 | 62.9 | 103 | 105.2 |
| P58252 | ***Eef2*** | 147.6 | 191.4 | 281.5 | 52.3 | 72.6 | 69.4 | 60.8 | 77.6 | 90.3 |
| Q9Z2I0 | ***Letm1*** | 2945.6 | 1731.8 | 1022.1 | 3986.8 | 4415.7 | 3684 | 3774.2 | 2911.1 | 2757.2 |
| Q8C196 | ***Cps1*** | 92879.7 | 49220.2 | 37398.7 | 706.5 | 683.3 | 577.9 | 683.4 | 327.4 | 1026.6 |
| O88451 | ***Rdh7*** | 1302.7 | 3640 | 2333.6 | 46.2 | 61.4 | 65.5 | 74.2 | 47.9 | 86.3 |
| Q3UEG6 | ***Agxt2*** | 770.6 | 629.2 | 264 | 28.4 | 23.5 | 27.8 | 28.3 | 29.6 | 42.5 |
| Q9D0K2 | ***Oxct1*** | 176.7 | 338.2 | 959.6 | 3269.5 | 3200.8 | 6428.5 | 1777.6 | 1913.7 | 2036.9 |
| Q60759 | ***Gcdh*** | 2814.5 | 1628.9 | 1300.2 | 280.9 | 342.2 | 303.6 | 349.6 | 225.1 | 270.7 |
| P19536 | ***Cox5b*** | 2945 | 1726.2 | 955.1 | 4255.3 | 4600.6 | 3576.6 | 4830.8 | 2961.6 | 2663.6 |
| P62204 | ***Calm1; Calm2; Calm3*** | 162.9 | 651.2 | 524.1 | 911 | 1357.1 | 1104.6 | 619.6 | 2423.5 | 1453.5 |
| P47915 | ***Rpl29*** | 1425.7 | 1358.8 | 2806.9 | 285.1 | 177 | 207.6 | 191.3 | 114.4 | 185.9 |
| Q9CXW2 | ***Mrps22*** | 134.9 | 109.7 | 90.1 | 238.4 | 263 | 165.7 | 224.1 | 145.6 | 119.4 |
| Q91XV3 | ***Basp1*** | 10.5 | 141.1 | 267.4 | 3322.4 | 5672.1 | 2164.4 | 2286.2 | 5969 | 2545.2 |
| Q8VCT4 | ***Ces1d*** | 7410.1 | 10329 | 3483.4 | 216.5 | 268.1 | 275.2 | 259.7 | 263.5 | 383.4 |
| Q8CAQ8 | ***Immt*** | 8942.1 | 5555.1 | 3284.6 | 13645.3 | 18618.2 | 11990.9 | 17054.7 | 7646.9 | 9536.7 |
| Q06185 | ***Atp5i*** | 1448.3 | 383.8 | 359.5 | 1799.4 | 3210.3 | 2948.7 | 1307.9 | 1988.3 | 1672.6 |
| Q9CY27 | ***Tecr*** | 809.3 | 952 | 1820.8 | 139.2 | 98.8 | 110.7 | 106 | 158.9 | 104.5 |
| P24369 | ***Ppib*** | 3130.5 | 3657.4 | 1642.3 | 1004.3 | 493.6 | 551.9 | 751.6 | 980.5 | 749 |
| P08249 | ***Mdh2*** | 2950.3 | 1526.9 | 1610.1 | 13805.3 | 10262.5 | 23862 | 8498.5 | 10966.6 | 9300.7 |
| Q99PG0 | ***Aadac*** | 1207.2 | 2554.4 | 1098.6 | 37.4 | 18.9 | 25.6 | 25.7 | 25.6 | 39 |
| Q61548 | ***Snap91*** | 211.3 | 194.5 | 255.6 | 310.3 | 471 | 490.9 | 449.8 | 1060.6 | 1084.5 |
| P51660 | ***Hsd17b4*** | 585 | 858 | 402.2 | 135.2 | 156.2 | 199.2 | 203.8 | 310.8 | 364.7 |
| Q8VDQ8 | ***Sirt2*** | 47.9 | 82.2 | 8.9 | 151.8 | 237.6 | 135.2 | 255.7 | 609.9 | 429.1 |
| Q922Q8 | ***Lrrc59*** | 1589.3 | 1515.6 | 671.8 | 403.9 | 107.4 | 161.7 | 187.7 | 121.8 | 133.1 |
| Q8R3V5 | ***Sh3glb2*** | 8 | 29.7 | 73 | 97.2 | 163.1 | 192 | 226.4 | 524.5 | 472.5 |
| Q8BMS1 | ***Hadha*** | 12072.1 | 8151.8 | 6548.2 | 3026 | 3947.8 | 3080.7 | 5543.2 | 3987.7 | 5080.3 |
| Q99N96 | ***Mrpl1*** | 68.8 | 60.6 | 37.4 | 107.4 | 123.3 | 83.7 | 95 | 66.9 | 56.1 |
| P57746 | ***Atp6v1d*** | 9.1 | 45 | 20.9 | 176.9 | 108.1 | 84.5 | 163 | 244.8 | 225.6 |
| P11352 | ***Gpx1*** | 1197.2 | 883.3 | 484.1 | 132.9 | 154.6 | 217 | 149 | 128.1 | 166.5 |
| Q05920 | ***Pc*** | 12546.2 | 6515.1 | 11428.7 | 3439.2 | 4471.3 | 4013.7 | 4720.2 | 2409.3 | 3575.8 |
| Q8CI94 | ***Pygb*** | 51.8 | 163.8 | 110.1 | 194.2 | 243.4 | 257.4 | 334.3 | 796.7 | 764.8 |
| Q9DCY0 | ***Keg1*** | 1120 | 597.3 | 447.9 | 56.2 | 50 | 61.6 | 36.7 | 41.7 | 68.8 |
| Q8VCU1 | ***Ces3b*** | 1324.5 | 1065.7 | 406.4 | 46 | 39.7 | 38.4 | 49.3 | 38.7 | 31.1 |
| O55022 | ***Pgrmc1*** | 2881.9 | 7185.8 | 3860.8 | 532.9 | 359.8 | 320.6 | 302.1 | 368.2 | 253.5 |
| P99024 | ***Tubb5*** | 79.8 | 166.3 | 596.3 | 732.7 | 936.1 | 822.7 | 970.5 | 1526.5 | 1883.3 |
| Q8CHT0 | ***Aldh4a1*** | 6284.8 | 5263.3 | 2178.2 | 624.4 | 683.8 | 628.5 | 960.9 | 630.8 | 712.3 |
| Q9QXX4 | ***Slc25a13*** | 9129.8 | 7935.7 | 2735.9 | 489.5 | 444.9 | 301.1 | 586.4 | 300.8 | 431.7 |
| Q9WU79 | ***Prodh*** | 1838.7 | 1154.5 | 893.7 | 381.6 | 482.9 | 326 | 582.6 | 311.9 | 472 |
| P40630 | ***Tfam*** | 1019.6 | 727.9 | 302.8 | 2018.9 | 1844.9 | 1212.1 | 2024.2 | 951.4 | 858.2 |
| Q8K411 | ***Pitrm1*** | 163.6 | 139.2 | 54.7 | 216.8 | 217.2 | 256.6 | 251.3 | 194.9 | 204.1 |
| Q5U458 | ***Dnajc11*** | 245.6 | 175.7 | 134.3 | 322.5 | 342.9 | 264.2 | 318.5 | 157.4 | 180.7 |
| P09103 | ***P4hb*** | 27702.7 | 30812.3 | 9097.5 | 1963.2 | 993.4 | 1468.7 | 1541 | 1568.7 | 1742.6 |
| P07724 | ***Alb*** | 6935.7 | 17300 | 8307.9 | 688.6 | 464.4 | 1105.1 | 1378.9 | 2998.8 | 3254.1 |

| | | | | | | | | | | |
|---|---|---|---|---|---|---|---|---|---|---|
| P17742 | ***Ppia*** | 212.9 | 514.7 | 244.8 | 717.7 | 830.6 | 1301.6 | 658.6 | 2284.9 | 1573.8 |
| Q8BWQ1 | ***Ugt2a3*** | 1683.4 | 2921.1 | 1151.6 | 388.9 | 130.1 | 335.8 | 137 | 109.4 | 229.7 |
| P49935 | ***Ctsh*** | 474.2 | 1397.5 | 921 | 164.5 | 71.8 | 84.6 | 115.7 | 70.2 | 131.3 |
| P18760 | ***Cfl1*** | 259.7 | 1101.1 | 927.6 | 1397.4 | 2107.9 | 1811 | 1961.7 | 4652.8 | 3098.1 |
| P08003 | ***Pdia4*** | 7921.1 | 9823.8 | 3145.8 | 1131.4 | 531.5 | 916 | 817.9 | 734.1 | 895.2 |
| Q9JLJ2 | ***Aldh9a1*** | 1519.2 | 1564 | 794.2 | 461.3 | 421.1 | 635.7 | 623 | 602.8 | 683.5 |
| Q63880 | ***Ces3a*** | 1403.4 | 1872.2 | 483.8 | 6.6 | 15.8 | 11 | 25 | 8.9 | 24.3 |
| P46097 | ***Syt2*** | 33.6 | 61.6 | 84.8 | 221 | 304.2 | 541 | 1740.9 | 1299.1 | 4403.4 |
| Q61646 | ***Hp*** | 457.4 | 412.9 | 135.9 | 33.9 | 27.3 | 27.8 | 36.7 | 37.9 | 39.5 |
| P63017 | ***Hspa8*** | 1053.5 | 2971.6 | 2362.5 | 3847.2 | 4434 | 6094.6 | 4609.8 | 8929.2 | 11421.9 |
| P09411 | ***Pgk1*** | 118.3 | 571.3 | 466.5 | 717.6 | 1051.3 | 1241.5 | 655.7 | 2553.3 | 2051.5 |
| P62259 | ***Ywhae*** | 205.2 | 558.5 | 559.1 | 735.2 | 1104.3 | 1283.2 | 838.2 | 2293.1 | 2040.7 |
| Q8R1I1 | ***Uqcr10*** | 111.7 | 27.4 | 39.2 | 1358.6 | 698.7 | 490.2 | 432.5 | 898.8 | 509.8 |
| Q8JZN5 | ***Acad9*** | 531.8 | 576.8 | 128.9 | 852.8 | 905 | 791.6 | 975.4 | 709.1 | 716.4 |
| Q61885 | ***Mog*** | 2 | 17.4 | 0 | 122.7 | 214.3 | 68 | 303.7 | 687.4 | 451.6 |
| Q8BHN3 | ***Ganab*** | 277.3 | 511.7 | 241.3 | 107.3 | 64.8 | 95.2 | 76.3 | 88.8 | 99.3 |
| Q5FW57 | ***Gm4952*** | 1763.3 | 1012.3 | 564.7 | 72.1 | 81.3 | 74.9 | 64.9 | 38.7 | 91.8 |
| Q99K67 | ***Aass*** | 3791 | 1973.4 | 1301.2 | 147.5 | 186.5 | 154.2 | 162.3 | 100.8 | 187.8 |
| Q9D172 | ***D10Jhu81e*** | 1829.7 | 1860.6 | 2619.4 | 4136.1 | 3863.6 | 6783.9 | 3966 | 3481.6 | 3485.3 |
| P97450 | ***Atp5j*** | 5161.5 | 3077.8 | 1085.4 | 6938.7 | 7624.9 | 5929.6 | 7218.6 | 5853.7 | 4335.7 |
| P24270 | ***Cat*** | 7953.7 | 9123.6 | 2291.6 | 214.7 | 273.6 | 405.9 | 380.6 | 502.3 | 686.9 |
| P17717 | ***Ugt2b17*** | 1558.7 | 2972.6 | 886.7 | 8.2 | 22 | 11.4 | 27.2 | 7.7 | 38.2 |
| P00405 | ***Mtco2*** | 706.5 | 636.3 | 298.4 | 2081.4 | 1588.8 | 975 | 1742.6 | 969.2 | 942.7 |
| Q3UNZ8 | | 256 | 156 | 75.4 | 359.4 | 297.8 | 314 | 879 | 460.6 | 589.9 |
| Q9D8E6 | ***Rpl4*** | 1723.2 | 1582.4 | 3743.5 | 388.2 | 251.3 | 324.2 | 285 | 199 | 303.5 |
| Q8QZR3 | ***Ces2a*** | 2134.9 | 2710.7 | 651.5 | 49.8 | 67.7 | 58.6 | 66.1 | 85.4 | 120.9 |
| Q91XE0 | ***Glyat*** | 2343.4 | 1485.6 | 611.2 | 59.6 | 64.7 | 54.8 | 108.4 | 85.4 | 137.7 |
| O35488 | ***Slc27a2*** | 1645.7 | 2216.4 | 4972.2 | 69.6 | 71.4 | 52.2 | 61.1 | 45.6 | 72.4 |
| Q9D6U8 | ***Fam162a*** | 1071.8 | 451.2 | 952.1 | 1241.3 | 1839.5 | 1555.3 | 1380.8 | 1403 | 1573.4 |
| Q9D379 | ***Ephx1*** | 674.7 | 1389 | 462.9 | 57.9 | 51.9 | 61.3 | 92.4 | 123.1 | 127.7 |
| Q8JZU2 | ***Slc25a1*** | 1993.6 | 1129.1 | 878.9 | 476.1 | 383.1 | 255.5 | 847 | 479 | 740.1 |
| Q9WUM5 | ***Suclg1*** | 6753.7 | 2879.1 | 2826.6 | 6837 | 9424.7 | 10696 | 7243.1 | 5361.2 | 6338.2 |
| Q8CGK3 | ***Lonp1*** | 1865.8 | 1331.2 | 1349.3 | 1966.2 | 2397.7 | 2033.4 | 2015.1 | 1164.3 | 1542.8 |
| P17182 | ***Eno1*** | 284.1 | 2956.9 | 1567.7 | 3263.6 | 4243.5 | 4491.9 | 3355.1 | 9964.3 | 6225.3 |
| Q9DD20 | ***Mettl7b*** | 1643.4 | 2151.5 | 4981.5 | 27.2 | 20.4 | 30.1 | 15.6 | 8.7 | 26.4 |
| P14094 | ***Atp1b1*** | 194.3 | 1169.9 | 1001.1 | 2586.5 | 1562.9 | 1810.1 | 6963.4 | 5188.8 | 11504 |
| P62075 | ***Timm13*** | 433.9 | 164.2 | 46.5 | 524.2 | 669.5 | 498.3 | 621.4 | 527.8 | 430.2 |
| P43006 | ***Slc1a2*** | 21.1 | 91.4 | 406.5 | 436 | 665.5 | 533.7 | 1288.5 | 1549.1 | 2793.1 |
| P60202 | ***Plp1*** | 12 | 42.6 | 20.9 | 503.4 | 1061.7 | 321 | 989.2 | 2443.7 | 2284.3 |
| Q9JHS4 | ***Clpx*** | 199.5 | 112.7 | 124.4 | 65.4 | 79.5 | 68.4 | 66.6 | 32.2 | 46.5 |
| P63321 | ***Rala*** | 100.3 | 454.7 | 350.1 | 787.5 | 1015.3 | 527.8 | 850.2 | 2034.6 | 1174.4 |
| P35441 | ***Thbs1*** | 109 | 195.4 | 265.7 | 40.7 | 28.7 | 97.4 | 48.5 | 17.3 | 69.2 |
| A2AS89 | ***Agmat*** | 1364.4 | 1286 | 295.9 | 48 | 75.3 | 68.1 | 85.1 | 88.1 | 113.8 |
| P04370 | ***Mbp*** | 83 | 169.8 | 212.2 | 1568.2 | 3800 | 1359.8 | 4801.6 | 8637.6 | 8078.8 |
| O08795 | ***Prkcsh*** | 542.6 | 602.9 | 216.5 | 157.1 | 107.4 | 131.9 | 125.1 | 140.2 | 136.3 |
| Q9DBF1 | ***Aldh7a1*** | 2558.7 | 1735.6 | 891.9 | 326.9 | 375.9 | 591.5 | 424.8 | 432 | 765.2 |
| P25688 | ***Uox*** | 4924.4 | 6428 | 1231.8 | 85 | 100.6 | 86.8 | 115.1 | 106.3 | 170.5 |

| | | | | | | | | | | |
|---|---|---|---|---|---|---|---|---|---|---|
| Q99L04 | ***Dhrs1*** | 706.7 | 996.6 | 485.8 | 373.4 | 351 | 256.6 | 382.9 | 357.4 | 304.5 |
| P36552 | ***Cpox*** | 421.2 | 374.7 | 185.1 | 152.7 | 63.8 | 147.3 | 115.8 | 55.7 | 68 |
| P32020 | ***Scp2*** | 4301.4 | 16537.7 | 8886.2 | 278.4 | 444.6 | 672.5 | 503.3 | 761.2 | 893 |
| Q8CIM7 | ***Cyp2d26*** | 455.6 | 868.9 | 275.7 | 68.6 | 66.3 | 72.8 | 65.3 | 45.2 | 55.2 |
| Q8BJ64 | ***Chdh*** | 7311.3 | 4973.1 | 1628 | 343.1 | 284.7 | 209.7 | 310.4 | 174.4 | 190.5 |
| Q99LB7 | ***Sardh*** | 9597.2 | 4714.4 | 3047.2 | 514.7 | 578.9 | 771.8 | 693.6 | 885.2 | 1094.6 |
| Q8VCW8 | ***Acsf2*** | 5669.1 | 3084.4 | 2170.9 | 782.5 | 974.4 | 938.8 | 2012.9 | 1229.7 | 1598 |
| Q9D1Q6 | ***Erp44*** | 805.1 | 1175.2 | 390.3 | 259.9 | 136 | 186.3 | 159.4 | 163.7 | 156.7 |
| P27659 | ***Rpl3*** | 873.5 | 897.1 | 2200.7 | 221.1 | 124.9 | 201.7 | 157.1 | 89.2 | 151.5 |
| Q99MR8 | ***Mccc1*** | 1395.5 | 783.6 | 1246 | 515 | 731.5 | 661 | 784.5 | 530.6 | 810.1 |
| P56395 | ***Cyb5a*** | 3909.9 | 5342.7 | 1023.5 | 101.1 | 136.8 | 159.6 | 114.3 | 123 | 162.8 |
| Q8K4Z3 | ***Naxe*** | 192 | 119.5 | 60.9 | 320.5 | 286.7 | 606.7 | 219.8 | 325.9 | 293.1 |
| Q9WVA2 | ***Timm8a1*** | 946.8 | 660.6 | 93.5 | 1153.1 | 1529.3 | 1177 | 1461.3 | 1338.5 | 1064.2 |
| Q922Q1 | ***42796*** | 7302.5 | 7421.2 | 2154.2 | 1513.3 | 1022.6 | 859.5 | 1502.4 | 824.2 | 1215.9 |
| Q9QY76 | ***Vapb*** | 1220.9 | 992.6 | 781.1 | 758.7 | 596.5 | 510.9 | 664.4 | 849.9 | 771.5 |
| Q99M71 | ***Epdr1*** | 276.7 | 410 | 296.1 | 4185.5 | 906.9 | 3971.2 | 2027.1 | 620 | 3611.5 |
| Q91WN4 | ***Kmo*** | 2803.8 | 1415.1 | 678.7 | 41.9 | 46.1 | 66.5 | 47.8 | 43.8 | 54.3 |
| Q91W90 | ***Txndc5*** | 191.7 | 179.3 | 59.7 | 48.9 | 27 | 32 | 36.6 | 35.5 | 32 |
| Q8R4N0 | ***Clybl*** | 846.1 | 473 | 294.4 | 855 | 999.7 | 1196.5 | 1106 | 988.8 | 886.5 |
| O88531 | ***Ppt1*** | 242.6 | 805.3 | 397.7 | 2518.8 | 949.4 | 3240.9 | 1685.5 | 469.1 | 2653.9 |
| P11588 | ***Mup1*** | 6112.4 | 3095.1 | 1443.7 | 155.5 | 93.4 | 89 | 126 | 63.1 | 190.3 |
| P29758 | ***Oat*** | 6226.3 | 3319 | 2153.4 | 604.1 | 627.5 | 1218.1 | 513.1 | 472.6 | 741.7 |
| P50172 | ***Hsd11b1*** | 1065.4 | 2161.8 | 544.2 | 34.8 | 54.7 | 90.3 | 50.1 | 34.2 | 96.6 |
| P27773 | ***Pdia3*** | 13298.1 | 13797.5 | 4448.9 | 3611.3 | 1798.3 | 2955.7 | 2122.9 | 2917 | 2530.8 |
| Q9JKR6 | ***Hyou1*** | 1968.3 | 3411 | 1317.4 | 929 | 394.9 | 575.5 | 524.8 | 503.7 | 492.9 |
| Q9DBT9 | ***Dmgdh*** | 3705.9 | 1620.2 | 1200.7 | 269.2 | 261.6 | 223.8 | 226.3 | 138.3 | 280.9 |
| Q07417 | ***Acads*** | 2250.9 | 1372.9 | 645.4 | 277.9 | 241.1 | 292.8 | 364.6 | 238.4 | 338 |
| Q8BW75 | ***Maob*** | 7585.2 | 6879.9 | 2512.9 | 1987 | 1930.6 | 1062.3 | 2249.2 | 1796.5 | 1720.3 |
| P20852 | ***Cyp2a5*** | 540.7 | 1489.4 | 456.5 | 9.8 | 10.2 | 14.3 | 17.6 | 7.7 | 19.1 |
| P12970 | ***Rpl7a*** | 522.8 | 553.8 | 181.9 | 143.3 | 127.2 | 105.5 | 167.6 | 113.5 | 153.5 |
| Q921X9 | ***Pdia5*** | 2325.7 | 1809.1 | 480.3 | 213.7 | 89 | 243.9 | 162.4 | 68.6 | 240.2 |
| Q921G7 | ***Etfdh*** | 12613 | 10130.4 | 4196.3 | 2947.1 | 2901.4 | 2685.5 | 3670.5 | 2745 | 2753.1 |
| Q61171 | ***Prdx2*** | 162.8 | 302.9 | 107.3 | 438.5 | 528.5 | 1035.2 | 632.4 | 1328.8 | 1927.9 |
| O35490 | ***Bhmt*** | 733 | 3348.7 | 4777 | 46.8 | 61.3 | 86.7 | 52 | 34.1 | 64.6 |
| P56135 | ***Atp5j2*** | 807.7 | 592.4 | 279.6 | 2179.6 | 1745.3 | 838.6 | 1216.6 | 1532.2 | 785.2 |
| O70503 | ***Hsd17b12*** | 754.4 | 964.4 | 2151.6 | 270.6 | 234.8 | 205.4 | 282.7 | 321.6 | 316.1 |
| P37040 | ***Por*** | 2326.7 | 5639.1 | 1679 | 315.3 | 246.8 | 262.6 | 293.1 | 402.2 | 342.5 |
| P09671 | ***Sod2*** | 2600.7 | 927.5 | 754.1 | 4176.6 | 3364.5 | 8144.3 | 3284.7 | 3120 | 3345.6 |
| P14148 | ***Rpl7*** | 1839.3 | 2259.6 | 544.2 | 427 | 187.8 | 298.6 | 288.7 | 184.7 | 282.3 |
| Q6P3A8 | ***Bckdhb*** | 477.6 | 228.6 | 225.6 | 96.8 | 131.8 | 102 | 111.4 | 62.2 | 115.2 |
| Q91W64 | ***Cyp2c70*** | 780.8 | 1031.3 | 2879 | 8.8 | 10.6 | 6.9 | 12.4 | 0 | 20.9 |
| Q9CYR0 | ***Ssbp1*** | 300.3 | 130.1 | 34.4 | 302.6 | 365.7 | 356.1 | 306.9 | 246.7 | 193.3 |
| P18242 | ***Ctsd*** | 640.6 | 1772 | 1341.2 | 9030.9 | 2042.2 | 8250.2 | 4433.1 | 1440.3 | 7329.4 |
| Q91YQ5 | ***Rpn1*** | 4460.3 | 5388.9 | 1042.8 | 760.8 | 394.6 | 551.8 | 649.2 | 420.5 | 615.1 |
| P11725 | ***Otc*** | 10269 | 6156.5 | 1574 | 243.9 | 250 | 176.6 | 246 | 125.1 | 388.2 |
| P50544 | ***Acadvl*** | 5654.6 | 4547.4 | 1628.2 | 1134.5 | 1380.2 | 1005.5 | 1646.2 | 986.3 | 1292.7 |
| P21614 | ***Gc*** | 960.2 | 1206.8 | 134.4 | 27 | 30.7 | 40.9 | 36.2 | 42.1 | 49 |

| | | | | | | | | | | |
|---|---|---|---|---|---|---|---|---|---|---|
| Q8JZZ0 | ***Ugt3a2*** | 400.5 | 773.4 | 165.9 | 59.9 | 52.9 | 56.9 | 41.6 | 45.5 | 43.5 |
| P46656 | ***Fdx1*** | 941.7 | 476.6 | 172.8 | 40 | 33.3 | 41.3 | 41.9 | 21 | 38.2 |
| O54734 | ***Ddost*** | 2599.3 | 2915.3 | 522.3 | 548.2 | 220.8 | 271.8 | 363.3 | 237.2 | 274.3 |
| Q61207 | ***Psap*** | 371.3 | 1812.5 | 1184 | 9667.2 | 2415 | 14381.6 | 3145.5 | 1621.2 | 7818.4 |
| P20029 | ***Hspa5*** | 14835 | 20887.9 | 5487.4 | 5171.7 | 2429.5 | 3758.3 | 3304.4 | 3387.1 | 3703.6 |
| P16015 | ***Ca3*** | 73.8 | 731.9 | 742.1 | 25.1 | 36.4 | 36.2 | 49 | 66.2 | 79.8 |
| P10852 | ***Slc3a2*** | 27.8 | 174.4 | 237.6 | 236.2 | 319.1 | 340.3 | 711.9 | 1227.4 | 1486.2 |
| Q922R8 | ***Pdia6*** | 2837.5 | 3743.6 | 850 | 859.1 | 344.8 | 576.2 | 482.8 | 431.5 | 414.9 |
| Q9WTP7 | ***Ak3*** | 2065.7 | 1257.9 | 667 | 1971 | 2527.4 | 2328.9 | 2543.3 | 2756.5 | 2081.8 |
| Q8R086 | ***Suox*** | 859.6 | 463.7 | 187.8 | 76.6 | 55 | 110.1 | 127 | 83.2 | 114.4 |
| P38060 | ***Hmgcl*** | 3201.7 | 2127.6 | 1017.4 | 756 | 715.4 | 799.4 | 490.9 | 245.9 | 311 |
| Q9Z2I8 | ***Suclg2*** | 5325.2 | 3346.5 | 1703.1 | 928.4 | 1438 | 1178.9 | 1412.7 | 1183.2 | 1574.9 |
| P16546 | ***Sptan1*** | 173 | 577.1 | 908.8 | 1279.3 | 2607.9 | 1114.3 | 1260.2 | 2984.5 | 1585.9 |
| P03930 | ***Mtatp8*** | 1017 | 197.3 | 147.7 | 1303.3 | 989.7 | 987.9 | 628.4 | 971.3 | 726.1 |
| P14211 | ***Calr*** | 12028 | 10795.3 | 3217.9 | 3725.9 | 1606.9 | 2721.1 | 2134.9 | 2372.3 | 2222.6 |
| Q9CQV1 | ***Pam16*** | 390.1 | 259.6 | 120.5 | 541.2 | 498.1 | 346.9 | 380.7 | 300.3 | 195.4 |
| Q6XVG2 | ***Cyp2c54*** | 1833.7 | 2971.4 | 8671.6 | 64 | 74.2 | 64.3 | 69.7 | 76.1 | 104.8 |
| P47738 | ***Aldh2*** | 14827.4 | 9080.2 | 3366 | 2138.3 | 1987.7 | 2456.9 | 3179.1 | 2498.2 | 2582.9 |
| P42125 | ***Eci1*** | 4624.1 | 3755.2 | 1616.5 | 1418.7 | 1657.7 | 1397.3 | 2361.7 | 1902.9 | 1777.6 |
| Q9DCN2 | ***Cyb5r3*** | 2816.3 | 4014.9 | 953 | 1222.7 | 520.6 | 330.5 | 849.2 | 808.4 | 629.1 |
| Q9CRD0 | ***Ociad1*** | 420.6 | 264.2 | 164.6 | 510.2 | 512.5 | 368.3 | 470.1 | 297 | 287.1 |
| O35857 | ***Timm44*** | 935.6 | 1018 | 368.4 | 1162.1 | 1359.1 | 1122.7 | 1003 | 763 | 692.5 |
| P21279 | ***Gnaq*** | 26 | 156.1 | 106.8 | 221.3 | 201.2 | 140.3 | 195.5 | 596.4 | 311.7 |
| P60710 | ***Actb*** | 218.3 | 988.6 | 764.6 | 892.7 | 1369.2 | 1459.4 | 1252.4 | 2765.2 | 1970.1 |
| Q9DCU9 | ***Hoga1*** | 1574.1 | 547.8 | 387.7 | 83.8 | 94.8 | 103 | 88.8 | 52.9 | 114.5 |
| Q91Y97 | ***Aldob*** | 171.6 | 2022.3 | 1154.8 | 35.5 | 60.9 | 80.4 | 68.5 | 104.9 | 132.2 |
| P10854 | ***Hist1h2bm*** | 79.1 | 118.6 | 198 | 176.5 | 642.6 | 1001.9 | 662.1 | 377.2 | 508.7 |
| Q8CG76 | ***Akr7a2*** | 307 | 273.9 | 153.2 | 122.1 | 161.3 | 163.8 | 144.2 | 153.3 | 154.9 |
| P45952 | ***Acadm*** | 5944.2 | 4048.6 | 1473.2 | 1123.7 | 1329.3 | 1466.7 | 1227.5 | 1124.7 | 1252 |
| B1AR13 | ***Cisd3*** | 757.1 | 505.3 | 844 | 491.9 | 559.6 | 421.5 | 706.1 | 442.5 | 550.5 |
| P30115 | ***Gsta3*** | 90.9 | 716.8 | 407.8 | 47.2 | 60.3 | 62.5 | 62.9 | 140.8 | 140 |
| P35700 | ***Prdx1*** | 580.5 | 2821.1 | 3784.9 | 384.3 | 565.7 | 737.2 | 630.9 | 1268 | 1299.6 |
| Q9D1D4 | ***Tmed10*** | 931.4 | 1957.6 | 500 | 487.6 | 155 | 187.6 | 252.7 | 236.3 | 186.5 |
| P84228 | ***Hist1h3b; Hist1h3c; Hist1h3d; Hist1h3e; Hist1h3f; Hist2h3b; Hist2h3c1; Hist2h3c2*** | 417.8 | 924.7 | 556.9 | 766.4 | 1252.3 | 1792.1 | 1095.7 | 877.5 | 889.7 |
| P35564 | ***Canx*** | 2272.4 | 3797.4 | 1364.9 | 1431.8 | 816.2 | 974 | 1277.9 | 1411.9 | 1184.1 |
| Q8QZT1 | ***Acat1*** | 8144.6 | 6465.1 | 2824.2 | 8572.6 | 9255.4 | 8647.1 | 8124.1 | 6060.8 | 4668.1 |
| Q9R112 | ***Sqrdl*** | 345.4 | 181.7 | 874.9 | 95 | 70 | 54.6 | 137.1 | 56 | 98.6 |
| Q4LDG0 | ***Slc27a5*** | 419 | 617.3 | 2206.5 | 18.5 | 19.4 | 23.2 | 25.4 | 17 | 15 |
| Q7TMM9 | ***Tubb2a*** | 164.5 | 922.4 | 4336.6 | 3681 | 5462.1 | 4021.9 | 4380.5 | 10363.8 | 9497.6 |
| P12710 | ***Fabp1*** | 754.5 | 4638.1 | 1340.3 | 19.7 | 30.1 | 40.8 | 41.7 | 11.5 | 84.8 |
| Q9D7B6 | ***Acad8*** | 1052.3 | 750.1 | 798.2 | 534.3 | 748.6 | 694.3 | 588.6 | 355.3 | 521.8 |
| Q8BGA8 | ***Acsm5*** | 712.8 | 374.8 | 35.3 | 15.9 | 17.2 | 27.6 | 32.6 | 23.3 | 56.6 |
| Q91WS0 | ***Cisd1*** | 2500 | 1456 | 735.1 | 2511.7 | 2818.7 | 2264.4 | 2175.7 | 1941.5 | 1695.3 |

| | | | | | | | | | | |
|---|---|---|---|---|---|---|---|---|---|---|
| P16460 | ***Ass1*** | 844.6 | 8729.7 | 18750.7 | 96.7 | 192.6 | 192.4 | 145.7 | 160.3 | 239.9 |
| Q9Z0X1 | ***Aifm1*** | 8714 | 5989.4 | 2162 | 2066.4 | 2394.4 | 2234.1 | 2639.3 | 1445.1 | 1965.6 |
| Q9DBG6 | ***Rpn2*** | 538.2 | 821.6 | 74.4 | 161.7 | 79.5 | 85.8 | 126.2 | 73 | 107.2 |
| Q61176 | ***Arg1*** | 139.1 | 980.3 | 2609.2 | 33.9 | 44.3 | 60.1 | 31.9 | 36.8 | 48.3 |
| Q64433 | ***Hspe1*** | 5915.7 | 2239.7 | 567 | 6057.3 | 4158.5 | 8341.1 | 4623.1 | 4594.5 | 3149.5 |
| Q99PL5 | ***Rrbp1*** | 936.2 | 308.3 | 271.3 | 221.1 | 94.2 | 129.1 | 162.2 | 75.6 | 116.8 |
| P14152 | ***Mdh1*** | 69.9 | 408.2 | 243.1 | 315.8 | 439.5 | 527.1 | 251.8 | 1136.6 | 916.6 |
| Q9WUU7 | ***Ctsz*** | 577.6 | 2826.1 | 667.3 | 191.5 | 58.8 | 246.7 | 125.6 | 33.9 | 250.7 |
| Q8R0Y6 | ***Aldh1l1*** | 48.6 | 454.3 | 1027.4 | 46.4 | 50.4 | 57.4 | 54.9 | 104.4 | 92 |
| Q9WTP6 | ***Ak2*** | 3955.6 | 1989.4 | 297.7 | 399.9 | 310.1 | 439.9 | 461.5 | 354 | 399.1 |
| Q9WVD5 | ***Slc25a15*** | 732.6 | 365 | 10.2 | 36.9 | 39.3 | 34.6 | 51.2 | 28 | 60.6 |
| P14824 | ***Anxa6*** | 19.3 | 247.1 | 402.3 | 45.6 | 42.9 | 51.9 | 55.7 | 182.4 | 122.9 |
| P13707 | ***Gpd1*** | 66.1 | 119 | 47.8 | 110.6 | 121.1 | 104.9 | 127.1 | 233.8 | 232.4 |
| P20060 | ***Hexb*** | 51.1 | 252.1 | 148.4 | 494.4 | 133.6 | 399.9 | 215.9 | 79.5 | 313 |
| Q99KB8 | ***Hagh*** | 591.1 | 524.3 | 274.3 | 274.2 | 312 | 342.1 | 253.7 | 263.7 | 251.5 |
| P46638 | ***Rab11b*** | 205.5 | 508.4 | 192.5 | 637.9 | 475.7 | 384.5 | 480.8 | 1154.8 | 698.5 |
| UPSP:K22E_HUMAN | | 3052 | 7506.4 | 18900.6 | 3183.9 | 1691.6 | 2519.6 | 684.7 | 766.7 | 2930.4 |
| Q80XN0 | ***Bdh1*** | 4068.7 | 4814.5 | 1158.6 | 1912.8 | 1522.3 | 1361.5 | 1921.7 | 1376.6 | 1387.1 |
| P52196 | ***Tst*** | 3508.1 | 1597.7 | 1050.4 | 802.2 | 1115.4 | 729.7 | 1373.1 | 1244.6 | 1174.2 |
| Q80Y14 | ***Glrx5*** | 577.7 | 261.9 | 76.8 | 495.6 | 565.3 | 536.8 | 549.6 | 432.4 | 320.4 |
| Q8BMS4 | ***Coq3*** | 284.9 | 194 | 42 | 267.3 | 281.8 | 301.7 | 267.3 | 174 | 139.6 |
| P10639 | ***Txn*** | 364.4 | 1139 | 1942.8 | 415.7 | 374 | 544.5 | 480.5 | 726.1 | 643.6 |
| Q62261 | ***Sptbn1*** | 44 | 181.6 | 349 | 369.7 | 504.1 | 237.7 | 296 | 699.3 | 375.3 |
| P07309 | ***Ttr*** | 448.4 | 680.7 | 81.2 | 122.8 | 71.2 | 200 | 41.4 | 33.6 | 58.7 |
| Q9CXI5 | ***Manf*** | 660.3 | 564.5 | 89.7 | 211.7 | 111.1 | 176.8 | 130 | 141.6 | 131.4 |
| Q920A5 | ***Scpep1*** | 505.8 | 2185.8 | 458.2 | 260.1 | 84.7 | 209 | 168.4 | 37.5 | 233.4 |
| P24549 | ***Aldh1a1*** | 60.6 | 797.8 | 414.1 | 86.7 | 89.7 | 127.3 | 88.4 | 205.8 | 196.9 |
| Q921H9 | ***Coa7*** | 646.6 | 334.7 | 122.6 | 580.2 | 536.3 | 716.3 | 474 | 332.8 | 356 |
| P38647 | ***Hspa9*** | 9885.3 | 6491.8 | 2696.9 | 8682.9 | 9805.5 | 10103.1 | 9255.3 | 6154.7 | 6605.8 |
| P63038 | ***Hspd1*** | 16306.4 | 10159.4 | 4070.1 | 14830.8 | 15829.1 | 15782.3 | 14774.4 | 10771.8 | 8865.4 |
| Q8BVI4 | ***Qdpr*** | 55.7 | 95.8 | 85.8 | 82 | 107.7 | 140.9 | 83.8 | 233 | 210.8 |
| Q9Z2Z6 | ***Slc25a20*** | 1222.7 | 1208.1 | 89.2 | 273.6 | 276.2 | 308.7 | 377.7 | 253.3 | 362.5 |
| P00329 | ***Adh1*** | 250.5 | 4110.6 | 14019.7 | 73.4 | 74.3 | 93.5 | 92.3 | 63 | 102.7 |
| P46425 | ***Gstp2*** | 126 | 844.5 | 516.8 | 169.4 | 213.6 | 197.3 | 166.7 | 515.9 | 321.7 |
| Q9CQ62 | ***Decr1*** | 5108.4 | 3990.5 | 554.4 | 1131.7 | 1516.7 | 994.4 | 1831.7 | 1726 | 1520.7 |
| Q9EQ20 | ***Aldh6a1*** | 10084.8 | 6609.6 | 2485.8 | 3065.5 | 3292.8 | 3284.1 | 4152.2 | 2711.4 | 3103 |
| P28798 | ***Grn*** | 212.3 | 1487.1 | 468.2 | 175.5 | 72.6 | 222.5 | 80.1 | 50.5 | 135 |
| Q8CC88 | ***Vwa8*** | 250.2 | 157.7 | 680.5 | 117.3 | 155.6 | 123.8 | 185.6 | 95 | 159 |
| P50136 | ***Bckdha*** | 1035.3 | 472.8 | 249.7 | 213.2 | 287.7 | 253.3 | 258 | 168.1 | 217 |
| P47802 | ***Mtx1*** | 387.7 | 382.4 | 230.5 | 427.8 | 461.3 | 361.8 | 315.8 | 193.1 | 192.8 |
| P52825 | ***Cpt2*** | 1983.7 | 1331 | 268.7 | 520.1 | 588.3 | 339.9 | 828.8 | 604.2 | 605.3 |
| Q61644 | ***Pacsin1*** | 48.7 | 203 | 510.2 | 366 | 611.7 | 431.4 | 557.2 | 1274.6 | 1079.2 |
| Q9CQX2 | ***Cyb5b*** | 1878.9 | 1601.8 | 537.4 | 863.3 | 787.7 | 637.6 | 944.9 | 694.1 | 671.7 |
| Q99JB2 | ***Stoml2*** | 635.1 | 366.9 | 97.5 | 539.8 | 651.6 | 565 | 653.6 | 518.1 | 494.1 |
| O70439 | ***Stx7*** | 73.7 | 312.4 | 103.8 | 531 | 160.9 | 336.3 | 215.2 | 249.2 | 321.4 |
| Q61425 | ***Hadh*** | 4681.5 | 2903.6 | 799.8 | 1115.7 | 1245.8 | 1433.5 | 1614.3 | 1379.5 | 1583.1 |
| Q9EP89 | ***Lactb*** | 440.8 | 293.2 | 133.1 | 417.1 | 433.6 | 382.8 | 311.9 | 183.3 | 214.7 |

| | | | | | | | | | | |
|---|---|---|---|---|---|---|---|---|---|---|
| P26443 | ***Glud1*** | 7940.2 | 6663.1 | 2378.8 | 5630.7 | 9722.8 | 11315 | 11227.3 | 11882.3 | 17464.5 |
| P62821 | ***Rab1A*** | 292.7 | 450.1 | 270.9 | 232.5 | 310.5 | 214 | 244 | 438.9 | 406.3 |
| Q8R0N6 | ***Adhfe1*** | 89.8 | 61.1 | 20.6 | 31.1 | 27.2 | 32.7 | 38.6 | 27.1 | 33.9 |
| P15636 | | 4535.6 | 6485.1 | 24649.9 | 3755.8 | 3321.2 | 3071.2 | 2973.9 | 4486.4 | 3320.1 |
| Q9DCW4 | ***Etfb*** | 10251.5 | 9744.5 | 2348.9 | 4004.3 | 4396.4 | 3780.7 | 5030 | 3915.9 | 3580 |
| P31786 | ***Dbi*** | 937.8 | 940.1 | 562.4 | 414.1 | 484.3 | 832.1 | 442.1 | 1632.6 | 1028.3 |
| Q99KR7 | ***Ppif*** | 592 | 300.6 | 92.2 | 490.5 | 433.1 | 1301.1 | 508.3 | 676.8 | 854.5 |
| Q9DCM2 | ***Gstk1*** | 817.8 | 469.6 | 334.4 | 234.5 | 431.4 | 348.9 | 343.3 | 382.5 | 494 |
| Q9CR98 | ***Fam136a*** | 609.7 | 333.9 | 50 | 530.2 | 478.1 | 640.3 | 340 | 334 | 224.7 |
| Q6IFX2 | ***Krt42*** | 571.6 | 1369.4 | 2193.8 | 790.3 | 522.9 | 940.2 | 283.4 | 221.7 | 609.9 |
| Q99LP6 | ***Grpel1*** | 852.3 | 558.2 | 262.1 | 676.2 | 767.9 | 968.8 | 593.2 | 449.1 | 407.5 |
| Q62151 | ***Ager*** | 36.2 | 708.1 | 291.8 | 692.2 | 417.6 | 777.6 | 1132.4 | 501.6 | 1169 |
| UPSP:TRYP_PIG | | 1639.8 | 3231.6 | 14464.8 | 1324.2 | 1318.5 | 1346.4 | 921.6 | 1299.1 | 980.5 |
| P10126 | ***Eef1a1*** | 1805.7 | 5489.4 | 23199.3 | 1378.6 | 1718.2 | 2311.1 | 1631.8 | 2571.2 | 3242.1 |
| Q9CQN7 | ***Mrpl41*** | 170.6 | 123.8 | 15.7 | 152.3 | 206.7 | 140.8 | 136.1 | 82.2 | 88.3 |
| P07901 | ***Hsp90aa1*** | 50.1 | 267.9 | 387.3 | 286.7 | 344.7 | 542.2 | 402.2 | 956.4 | 1100.9 |
| Q99JY0 | ***Hadhb*** | 9708.8 | 6484.6 | 1599.6 | 2505 | 3747.1 | 2618.2 | 5067.9 | 3943.8 | 4395.2 |
| Q9CRB9 | ***Chchd3*** | 3885.5 | 2058.7 | 800.7 | 3449.5 | 4523.4 | 2668 | 4606.5 | 2102.1 | 2098.7 |
| P63030 | ***Mpc1*** | 462.9 | 243.4 | 185.7 | 504 | 484.7 | 298.8 | 397 | 446.7 | 281.5 |
| UPSP:K1CJ_HUMAN | | 2242.1 | 6898.6 | 22321.3 | 3466.7 | 1822 | 3637.7 | 884.1 | 921.1 | 2575 |
| Q9CQ01 | ***Rnaset2*** | 109.3 | 325.8 | 67.1 | 65.2 | 45.2 | 93.7 | 58.6 | 23.2 | 87.5 |
| Q8VDD5 | ***Myh9*** | 144.3 | 361.9 | 2774.4 | 67.8 | 43.5 | 92.4 | 55.6 | 38.7 | 57.8 |
| P21107 | ***Tpm3*** | 176.3 | 863 | 712.2 | 221 | 281.7 | 450.6 | 238.7 | 589.5 | 435.5 |
| Q922U2 | ***Krt5*** | 585.3 | 1860.6 | 6626.3 | 918.3 | 433.7 | 1049.4 | 180.3 | 193.6 | 684.8 |
| P00158 | ***Mt-Cyb*** | 668.9 | 411 | 878.2 | 1431.7 | 876.5 | 629 | 750.7 | 423.7 | 793.1 |
| P17047 | ***Lamp2*** | 113.6 | 1112.9 | 395.6 | 240.4 | 71.2 | 247.3 | 225.4 | 65 | 409.3 |
| P99027 | ***Rplp2*** | 819.8 | 927.5 | 203.5 | 426.4 | 271.8 | 440.1 | 370.4 | 307.2 | 327.6 |
| Q9WUR2 | ***Eci2*** | 1194.2 | 1053 | 361.2 | 419.1 | 672.2 | 584.5 | 636.7 | 556.1 | 678.7 |
| UPSP:K1CN_HUMAN | | 482.2 | 1717.9 | 7292.2 | 775.5 | 380.8 | 1056.5 | 202.4 | 211.5 | 575 |
| UPSP:K2C5_HUMAN | | 92.3 | 350.5 | 1058.2 | 158 | 68.3 | 263.2 | 33 | 55.1 | 135.6 |
| Q8K1Z0 | ***Coq9*** | 565 | 288.6 | 183.5 | 479.4 | 465.3 | 482.9 | 435.3 | 337.6 | 264.2 |
| O35465 | ***Fkbp8*** | 98 | 109.6 | 237.7 | 98.8 | 114.3 | 74.3 | 79.3 | 52.4 | 61.6 |
| Q91ZA3 | ***Pcca*** | 3095.7 | 2037.3 | 1230.3 | 2536.1 | 2861.1 | 2844.3 | 2429.9 | 1528.4 | 1742.4 |
| P09528 | ***Fth1*** | 74.8 | 403 | 363.6 | 155.2 | 146.1 | 185.7 | 148 | 279.8 | 332.8 |
| Q9CQI6 | ***Cotl1*** | 44.2 | 137.9 | 113 | 117.5 | 108.8 | 349.3 | 68.8 | 297 | 221.1 |
| Q9QYR9 | ***Acot2*** | 91.4 | 107.5 | 4.2 | 105.6 | 87.6 | 122.7 | 97.1 | 99.3 | 105.4 |
| Q99LC5 | ***Etfa*** | 8987.3 | 5556.4 | 1517.4 | 2627.4 | 3098.5 | 3105.6 | 3502.6 | 2531.7 | 2975 |
| Q9D0S9 | ***Hint2*** | 503.9 | 170.7 | 114 | 372 | 335.7 | 529.8 | 279.8 | 243.4 | 267.6 |
| UPSP:K1CI_HUMAN | | 2671.2 | 14887.4 | 93502.1 | 5615.4 | 2023.7 | 9276.7 | 992.5 | 1138.8 | 3661.3 |
| UPSP:K2C1_HUMAN | | 4622.4 | 20961.1 | 116672.9 | 9087.8 | 3566.7 | 14081.7 | 1627 | 1946.2 | 5836.4 |
| Q91W43 | ***Gldc*** | 521.1 | 243.4 | 133 | 168.2 | 220.8 | 119.9 | 64.5 | 32.3 | 50 |
| P70441 | ***Slc9a3r1*** | 30.8 | 249.2 | 201 | 89.3 | 95 | 80.5 | 73.2 | 220.7 | 125.4 |
| P16675 | ***Ctsa*** | 415.4 | 1466.1 | 318.4 | 424.5 | 124.6 | 419.8 | 173 | 67.6 | 284.3 |
| Q91VT4 | ***Cbr4*** | 644.4 | 418.7 | 305.9 | 370.9 | 355.7 | 318.5 | 377.4 | 217.7 | 301.4 |
| P60766 | ***Cdc42*** | 156.3 | 796 | 628.7 | 302.4 | 370.8 | 302.5 | 402.2 | 850.3 | 680.4 |
| P02088 | ***Hbb-b1*** | 1281.1 | 734.6 | 1059.9 | 753.4 | 1466 | 7941.7 | 1230.8 | 1229.8 | 9294.4 |

| | | | | | | | | | | |
|---|---|---|---|---|---|---|---|---|---|---|
| Q3ULD5 | ***Mccc2*** | 900.9 | 757 | 464.6 | 515.5 | 659.7 | 526.6 | 727.8 | 494.5 | 594.5 |
| Q3UJU9 | ***Rmdn3*** | 497.6 | 244.3 | 73.9 | 133.1 | 193.9 | 106.9 | 154.8 | 57.8 | 102.3 |
| P01868 | ***Ighg1*** | 3670.8 | 6584.2 | 44552.3 | 6069.4 | 3812.2 | 5140.7 | 4059.9 | 4007.1 | 5119.9 |
| Q60648 | ***Gm2a*** | 320 | 1329.6 | 481.2 | 427 | 183.8 | 528.3 | 190.5 | 113.7 | 368.1 |
| Q9D404 | ***Oxsm*** | 441.1 | 388.1 | 142.7 | 434.4 | 343.6 | 502.7 | 373.6 | 270.7 | 426.6 |
| Q9D6K5 | ***Synj2bp*** | 222.9 | 102.1 | 66.3 | 210.4 | 220.2 | 127.5 | 272.3 | 106.1 | 123.2 |
| P07356 | ***Anxa2*** | 84.5 | 1100.7 | 756.5 | 1137.8 | 725.6 | 1045.5 | 1959.9 | 855.9 | 1691.6 |
| P17426 | ***Ap2a1*** | 19.8 | 144.8 | 301.6 | 209 | 252.3 | 251.3 | 271.6 | 448.3 | 584.8 |
| P16332 | ***Mut*** | 1678.5 | 1071.7 | 463.4 | 725.8 | 723.4 | 727.7 | 745.6 | 440 | 465.4 |
| P55302 | ***Lrpap1*** | 486.5 | 545 | 237.5 | 428.9 | 206.5 | 302.5 | 249.9 | 287.3 | 231.1 |
| O08709 | ***Prdx6*** | 120.2 | 413.6 | 350.8 | 321.6 | 396.4 | 443.1 | 428.8 | 1577.7 | 921.6 |
| P51863 | ***Atp6v0d1*** | 203.8 | 1345.7 | 975.2 | 1359.9 | 981.2 | 1184.5 | 1072.7 | 2709.4 | 2037.8 |
| Q9WV54 | ***Asah1*** | 409.4 | 2304.8 | 795.5 | 2362.1 | 707.7 | 2885.5 | 1834.4 | 440.9 | 3650.1 |
| Q01853 | ***Vcp*** | 112.3 | 362.8 | 291.3 | 286.3 | 305.5 | 407 | 294.2 | 663.2 | 691.2 |
| O09159 | ***Man2b1*** | 85.4 | 351.3 | 19 | 66.1 | 32.6 | 74.2 | 49.1 | 32.1 | 89.2 |
| P16858 | ***Gapdh*** | 240.1 | 1583.3 | 4030.2 | 2354.5 | 3418 | 3237 | 3447.9 | 6328.6 | 5778.9 |
| O55143 | ***Atp2a2*** | 454.5 | 641.8 | 2030.6 | 888.5 | 436.3 | 389 | 522.5 | 527.8 | 487.2 |
| P01942 | ***Hba*** | 1188.2 | 374.4 | 608 | 410.1 | 857.9 | 4190.8 | 814 | 614.4 | 5624.3 |
| P51174 | ***Acadl*** | 3889.1 | 3319.4 | 739.5 | 1707.5 | 1939.3 | 1712.2 | 2830 | 2520.8 | 2366.6 |
| Q791V5 | ***Mtch2*** | 790 | 479.8 | 637.7 | 500.8 | 656.2 | 465.5 | 645.5 | 358.6 | 575.2 |
| P68369 | ***Tuba1a*** | 176 | 1129.5 | 7663.6 | 3779.9 | 5875 | 5791 | 5355.5 | 10933.3 | 12533 |
| Q99L13 | ***Hibadh*** | 2092 | 1664.5 | 737.1 | 1103.3 | 1146 | 1205 | 1430.6 | 1090.9 | 832.7 |
| O35459 | ***Ech1*** | 628 | 377.3 | 82.2 | 180.7 | 202.6 | 286.9 | 181.6 | 181.9 | 263.9 |
| P04943 | | 1263.2 | 2419.6 | 10841.2 | 2301.6 | 1806.9 | 2618.8 | 2212.1 | 2155.1 | 2279.8 |
| Q9QYG0 | ***Ndrg2*** | 21.8 | 135.2 | 304.5 | 67.2 | 77.5 | 104.2 | 129.9 | 306.9 | 281.5 |
| P67778 | ***Phb*** | 5573.2 | 3737.1 | 550 | 5240.6 | 4973.9 | 3586.1 | 3317.3 | 3025.1 | 2311 |
| P01837 | | 2408.3 | 4106.3 | 15011.6 | 4610.4 | 3035.2 | 3906.6 | 4344.7 | 3747.7 | 4056.2 |
| P08226 | ***Apoe*** | 363.9 | 1694.5 | 337.6 | 599.7 | 205.2 | 438.3 | 367.1 | 284.8 | 513.9 |
| Q91V41 | ***Rab14*** | 290.7 | 824.6 | 781.2 | 676.5 | 349.2 | 369.6 | 527.5 | 686 | 691.9 |
| O35114 | ***Scarb2*** | 497.8 | 3691.1 | 492.2 | 974.2 | 212.8 | 786.7 | 1004.7 | 138.7 | 1604.7 |
| Q8R164 | ***Bphl*** | 986.1 | 535.5 | 422.2 | 714.3 | 623.1 | 1218.6 | 667.2 | 472.1 | 653.4 |
| Q8K1R3 | ***Pnpt1*** | 162.4 | 257.1 | 935.5 | 210.8 | 293.2 | 259.7 | 240.2 | 165.6 | 206.1 |
| Q8VDN2 | ***Atp1a1*** | 455.7 | 4058.2 | 7800.1 | 2544.3 | 2362.9 | 2324.4 | 2213.2 | 5405.3 | 3634.2 |
| P06797 | ***Ctsl*** | 313.7 | 751.3 | 285 | 1028.8 | 200.7 | 800.9 | 447.7 | 115.6 | 590.5 |
| Q8QZS1 | ***Hibch*** | 814.7 | 451.7 | 414.2 | 410.8 | 283.9 | 612.2 | 437.9 | 387.4 | 428.3 |
| P62984 | ***Uba52*** | 503.8 | 1749.4 | 4517.4 | 1647.8 | 810.5 | 1633.6 | 1572.5 | 1602.3 | 2615.5 |
| P29416 | ***Hexa*** | 147.4 | 358.7 | 462.8 | 312.4 | 94.7 | 312.4 | 203.3 | 48.7 | 310.5 |
| Q9CQ22 | ***Lamtor1*** | 104.3 | 699.7 | 411 | 428.3 | 99.4 | 273.9 | 138 | 106.6 | 233.7 |
| P56213 | ***Gfer*** | 119.8 | 48 | 20.9 | 78.4 | 72 | 103.4 | 66.4 | 43.8 | 50.6 |
| P80317 | ***Cct6a*** | 23.2 | 232.6 | 296.3 | 117.7 | 113.4 | 149.8 | 142.1 | 214.9 | 254.5 |
| Q60864 | ***Stip1*** | 77.1 | 237.6 | 155.7 | 155.6 | 201.9 | 215 | 186.4 | 597.2 | 427.2 |
| O70318 | ***Epb41l2*** | 16.7 | 67.2 | 164.6 | 89.7 | 139.6 | 112.8 | 130.3 | 363.6 | 328.4 |
| Q9DBL1 | ***Acadsb*** | 319.1 | 181.3 | 76.9 | 186.9 | 291.3 | 255 | 279.6 | 188.3 | 277.6 |
| Q78IK4 | ***Apool*** | 698.9 | 513.3 | 210.4 | 418.6 | 413.3 | 291.1 | 476.8 | 307.8 | 278.8 |
| Q60634 | ***Flot2*** | 77 | 550.2 | 501.4 | 379.7 | 174.6 | 249.8 | 265.9 | 420.1 | 302.4 |
| Q9R013 | ***Ctsf*** | 89.8 | 316 | 256.5 | 385 | 90.5 | 425.8 | 162.4 | 41.9 | 310.9 |
| Q925I1 | ***Atad3*** | 2282.7 | 1361.5 | 1608.2 | 1506.7 | 1757.5 | 1432.8 | 1454.5 | 875.7 | 992.5 |

| | | | | | | | | | | |
|---|---|---|---|---|---|---|---|---|---|---|
| Q9DCV4 | ***Rmdn1*** | 376.1 | 173.6 | 55.6 | 235.1 | 257.2 | 292.7 | 231.9 | 227.6 | 266.2 |
| Q60936 | ***Coq8a*** | 169.2 | 61.5 | 35.7 | 66.9 | 65.7 | 56.8 | 67.7 | 31.2 | 39.2 |
| P11499 | ***Hsp90ab1*** | 233.9 | 1277.9 | 3250.6 | 692.5 | 1006.2 | 1398.5 | 1181.9 | 3077.9 | 2598.1 |
| P10649 | ***Gstm1*** | 70.1 | 574 | 203.7 | 201.6 | 181.1 | 190.9 | 193.9 | 755.6 | 431.1 |
| P08228 | ***Sod1*** | 409.7 | 970.1 | 376.4 | 366.6 | 302 | 682.4 | 305.1 | 873.6 | 806.4 |
| O89023 | ***Tpp1*** | 618.7 | 2481.7 | 731.8 | 2046.6 | 599.8 | 2747.4 | 974.7 | 285 | 1935.1 |
| Q9DCM0 | ***Ethe1*** | 1081.9 | 806 | 455.4 | 634.2 | 605.7 | 767.4 | 852.1 | 556.8 | 559.2 |
| Q8R0F8 | ***Fahd1*** | 530 | 406.6 | 312 | 355.9 | 423 | 350.7 | 408.2 | 286.4 | 329.8 |
| Q8C5H8 | ***Nadk2*** | 957.1 | 581.3 | 352 | 449.3 | 454.4 | 655.3 | 686.1 | 554.1 | 710.5 |
| Q8K4F5 | ***Abhd11*** | 418.6 | 406.2 | 1341.2 | 458.7 | 465.9 | 691.7 | 504.9 | 475.3 | 550 |
| O35129 | ***Phb2*** | 5882 | 3120.1 | 728.3 | 4731.3 | 4475.7 | 3161.3 | 3967.9 | 2746.1 | 2231.2 |
| Q99MN9 | ***Pccb*** | 1742.4 | 1386.7 | 785.6 | 1409.7 | 1455.1 | 1517.2 | 1286.2 | 858.1 | 883.6 |
| P68368 | ***Tuba4a*** | 26.1 | 256.3 | 2181.9 | 316.4 | 552.7 | 459.9 | 517.1 | 967.8 | 1007.9 |
| Q9CZS1 | ***Aldh1b1*** | 1642.8 | 751.6 | 301.1 | 602.1 | 626.5 | 806.9 | 393.6 | 443.4 | 362.2 |
| Q9ER88 | ***Dap3*** | 63 | 222.9 | 38.2 | 164.8 | 146.5 | 111.6 | 115.5 | 86.2 | 76.6 |
| Q99LB2 | ***Dhrs4*** | 1540.7 | 1372.3 | 362.1 | 892.1 | 1046.4 | 719.4 | 1361.9 | 1127.6 | 1221.6 |
| P62827 | ***Ran*** | 47.2 | 76.1 | 224.5 | 91.6 | 150.8 | 200.7 | 142.3 | 195.8 | 270.2 |
| P84091 | ***Ap2m1*** | 22.5 | 178.1 | 281.9 | 241.5 | 206.4 | 152.4 | 303.4 | 479.5 | 574.1 |
| O35643 | ***Ap1b1*** | 76.9 | 519.9 | 1752.5 | 567.4 | 567 | 474.6 | 677.1 | 993 | 1219.5 |
| Q8K2B3 | ***Sdha*** | 9254.4 | 6158.4 | 3394.6 | 6473.8 | 7076 | 7807.2 | 8600.6 | 5054.8 | 6319.3 |
| Q9DBG3 | ***Ap2b1*** | 51.8 | 819.8 | 830.8 | 332.3 | 426.4 | 588.8 | 373.4 | 768.4 | 726.4 |
| O08756 | ***Hsd17b10*** | 867.2 | 947.4 | 226.7 | 723.9 | 611.2 | 379.4 | 702.4 | 406 | 446.8 |
| P48036 | ***Anxa5*** | 81.5 | 411.9 | 185.5 | 244.4 | 218.9 | 347.1 | 321.4 | 1441.2 | 956.3 |
| Q8BH95 | ***Echs1*** | 4894.1 | 3380.5 | 1051.9 | 3269.1 | 3927.6 | 3540.3 | 4129 | 3179.3 | 3265.3 |
| P63158 | ***Hmgb1*** | 434.8 | 1082.2 | 459 | 781.5 | 692.3 | 764 | 449.2 | 721.9 | 586.7 |
| Q9CQA3 | ***Sdhb*** | 8777.7 | 4335.2 | 2392.4 | 5369.6 | 6294.7 | 6100.3 | 6116.9 | 4031.8 | 4234.3 |
| P54071 | ***Idh2*** | 2010.1 | 2108.2 | 640.9 | 1411 | 1387.6 | 1440.2 | 1608.3 | 1142.8 | 1188.7 |
| P68134 | ***Acta1*** | 1643.7 | 10424.1 | 12003.5 | 5174.2 | 7597.8 | 7734.5 | 7875 | 15400.8 | 10960.1 |
| G5E829 | ***Atp2b1*** | 95.5 | 232.4 | 1306 | 641 | 739.6 | 663.1 | 769.5 | 1443.2 | 1571.7 |
| P10605 | ***Ctsb*** | 503 | 1859.5 | 811.9 | 1152.7 | 235.1 | 1246.3 | 618.1 | 191.4 | 1072.4 |
| Q99J99 | ***Mpst*** | 1029 | 358.9 | 133.2 | 383.2 | 494.4 | 362.3 | 415.2 | 268.2 | 195 |
| Q9D023 | ***Mpc2*** | 874.9 | 800.9 | 84.2 | 698.6 | 559.7 | 208.1 | 520.5 | 507.8 | 364.9 |
| P03911 | ***Mtnd4*** | 79.5 | 122.2 | 143.6 | 164.8 | 120.9 | 87.3 | 75.9 | 80.9 | 68.9 |
| Q8BUV3 | ***Gphn*** | 7 | 56.1 | 142.4 | 60.8 | 96.9 | 85.8 | 147.4 | 220.5 | 276.1 |
| P26041 | ***Msn*** | 714.8 | 2912.5 | 1005.6 | 1329.9 | 1275.7 | 1446.3 | 1062.9 | 1716.3 | 1310.8 |
| P68372 | ***Tubb4b*** | 25.6 | 186.4 | 1344.1 | 421.2 | 419 | 383.1 | 490.3 | 913.7 | 744.7 |
| Q99JR1 | ***Sfxn1*** | 1619.2 | 624.2 | 320.9 | 1115.9 | 1046.5 | 724.2 | 551.6 | 425.6 | 313.9 |
| P70699 | ***Gaa*** | 139.8 | 725.6 | 288.6 | 579.2 | 103.5 | 314.7 | 229.3 | 84.1 | 285.6 |
| Q9CQN1 | ***Trap1*** | 793.8 | 630.2 | 506 | 578.6 | 687 | 602.9 | 615.1 | 388.4 | 460.7 |
| P27573 | ***Mpz*** | 55 | 95.6 | 157.4 | 109.4 | 94.9 | 83.3 | 1176.7 | 562.1 | 4098.7 |

| | | | | | | | | | | |
|---|---|---|---|---|---|---|---|---|---|---|
| P62806 | ***Hist1h4a; Hist1h4b; Hist1h4c; Hist1h4d; Hist1h4f; Hist1h4h; Hist1h4i; Hist1h4j; Hist1h4k; Hist1h4m; Hist2h4a; Hist4h4*** | 72.9 | 119.4 | 684.6 | 138.1 | 405.9 | 476 | 423.6 | 291.4 | 380.8 |
| Q8BP40 | ***Acp6*** | 438.1 | 277 | 39.6 | 317.5 | 179.8 | 332.7 | 291.5 | 171.1 | 231.2 |
| O55125 | ***Nipsnap1*** | 5247.3 | 2243.5 | 1440.3 | 2900.7 | 3209.9 | 3503.6 | 2119.5 | 1399 | 1374 |
| Q6PB66 | ***Lrpprc*** | 860.4 | 876.1 | 1394.9 | 1139.8 | 995.4 | 1096.9 | 760 | 571.4 | 601.1 |
| Q9JHI5 | ***Ivd*** | 2609.6 | 1338.9 | 243.3 | 1107.2 | 1151.9 | 1564.3 | 980.1 | 928.9 | 1061.7 |
| P15105 | ***Glul*** | 207.9 | 1212.6 | 1848.7 | 578.5 | 1016.1 | 1397 | 730.7 | 1178.3 | 1587.7 |
| P03995 | ***Gfap*** | 137.7 | 446.4 | 1355.3 | 548.9 | 524.6 | 1058.9 | 1284.9 | 1968.4 | 2903.3 |
| P10637 | ***Mapt*** | 57.3 | 207.5 | 671.9 | 274.4 | 437.2 | 308.2 | 301.5 | 726.6 | 541.6 |
| Q571F8 | ***Gls2*** | 837.3 | 311.4 | 300 | 527 | 432.8 | 552.4 | 296.6 | 158.7 | 185.9 |
| Q8BFZ3 | ***Actbl2*** | 204.9 | 986.1 | 1466.8 | 743.3 | 870.8 | 937.1 | 1035.4 | 1573.7 | 1488.2 |
| P11438 | ***Lamp1*** | 41.9 | 469.7 | 249.2 | 305 | 77.6 | 337.6 | 325.8 | 79.3 | 558.6 |
| P51150 | ***Rab7a*** | 372.4 | 787.3 | 299.6 | 704.4 | 419.4 | 385.2 | 317.7 | 506.8 | 479.8 |
| Q9CPY7 | ***Lap3*** | 673.6 | 617.6 | 187.1 | 502.8 | 543.9 | 471.5 | 586.8 | 650.7 | 563.5 |
| Q68FD5 | ***Cltc*** | 99.7 | 373.5 | 1440.5 | 684.5 | 530.4 | 780.3 | 1356.8 | 1556.9 | 3126.8 |
| Q9D1G1 | ***Rab1b*** | 346.2 | 737.8 | 352.4 | 569.4 | 496 | 394.8 | 701 | 893.1 | 905.5 |
| P22315 | ***Fech*** | 1239.4 | 666.4 | 622.3 | 745.9 | 925 | 825.8 | 798.2 | 504 | 551.8 |
| P50171 | ***Hsd17b8*** | 233.6 | 223.2 | 71.4 | 156 | 190.4 | 175.3 | 187.4 | 134.1 | 161.7 |
| P06151 | ***Ldha*** | 50.9 | 342.9 | 233.4 | 157.7 | 227.2 | 232.9 | 165.5 | 486.5 | 407.6 |
| O08917 | ***Flot1*** | 54.9 | 293.3 | 205.8 | 274.4 | 109.9 | 174.2 | 177 | 274.5 | 215.4 |
| Q2TPA8 | ***Hsdl2*** | 293.7 | 303.4 | 43.1 | 200.8 | 225.4 | 216.8 | 250 | 259.2 | 265.8 |

Table S3: Gene Ontology analyses of proteome: Liver vs Brain

| | Liver | Brain |
|---|---|---|
| Total intensity of all peptides | 1,062,212 | 1,086,811 |
| Cytosolic | 20.90% | 20.90% |
| Endoplasmic Reticulum | 18.90% | 4.10% |
| Plasma Membrane | 11.40% | 16.40% |
| Cytoskeletal | 4.80% | 5.30% |
| Endolysosomal | 3.50% | 4.90% |
| Mitochondrial | 52.50% | 75.20% |
| Total intensity of mitochondrial proteins | 557,174 | 817,049 |
| OMM | 9.80% | 9.80% |
| IMM | 65.90% | 64.30% |
| IMMS | 5.10% | 7.40% |
| Matrix | 42.30% | 27.80% |

**Table S4: Immunoisolated proteome of basal and starved Liver Avs**

Only proteins that were detected with more than 3 unique peptides are listed.

The numerical values are peptide intensity normalized to total intensity of each sample.

| uniprot | gene symbol | Intensity of Basal1 | Intensity of Basal2 | Intensity of Basal3 | Intensity of Starved1 | Intensity of Starved2 | Intensity of Starved3 |
|---|---|---|---|---|---|---|---|
| O89017 | ***Lgmn*** | 59.7 | 109.7 | 85.8 | 193.4 | 215.4 | 206 |
| P10605 | ***Ctsb*** | 119.9 | 398.3 | 475 | 1407 | 1455.4 | 1791.7 |
| Q61646 | ***Hp*** | 144.6 | 134.2 | 320.2 | 632.6 | 738.2 | 695.4 |
| P18242 | ***Ctsd*** | 1388.6 | 1642.6 | 3310.6 | 5600.3 | 5384 | 5600.1 |
| Q99LB7 | ***Sardh*** | 771.9 | 681.2 | 687.7 | 1134.1 | 985.9 | 1212.2 |
| P11276 | ***Fn1*** | 91.8 | 120.1 | 95.9 | 255.4 | 273.9 | 192.9 |
| Q9Z0J0 | ***Npc2*** | 89.2 | 54.4 | 148.5 | 455.4 | 304.1 | 489.5 |
| P20918 | ***Plg*** | 110.8 | 199.5 | 222.3 | 326.4 | 368.9 | 351.1 |
| Q8BFR4 | ***Gns*** | 24.1 | 22.7 | 48.7 | 87.5 | 95.2 | 128.7 |
| Q920A5 | ***Scpep1*** | 106.4 | 326.5 | 235 | 639.9 | 553.6 | 782.1 |
| P25688 | ***Uox*** | 958.7 | 826 | 818.6 | 2166 | 1751.4 | 1498.2 |
| Q91YW3 | ***Dnajc3*** | 139.4 | 181.9 | 225.5 | 295 | 341.3 | 304.1 |
| P07309 | ***Ttr*** | 157.9 | 206.3 | 439.5 | 630.3 | 709.2 | 749.8 |
| P97821 | ***Ctsc*** | 28 | 80.7 | 172.9 | 331.3 | 269.9 | 304.1 |
| P62751 | ***Rpl23a*** | 323.6 | 330.6 | 537.6 | 1080.2 | 922.4 | 752 |
| P06909 | ***Cfh*** | 125.3 | 134.9 | 155.7 | 226.6 | 297.2 | 226.3 |
| O88531 | ***Ppt1*** | 234 | 399.9 | 452.6 | 786.1 | 753 | 1043.8 |
| Q9WUU7 | ***Ctsz*** | 127.6 | 573.9 | 471.2 | 1210.3 | 1148.7 | 1764.7 |
| O89023 | ***Tpp1*** | 330.6 | 659.3 | 1034.5 | 1411.1 | 1768.1 | 1834.9 |
| Q9CR57 | ***Rpl14*** | 121.8 | 128.5 | 147 | 226.1 | 182.9 | 183.5 |
| Q9ESB3 | ***Hrg*** | 16.4 | 30 | 44 | 81 | 110.6 | 70.5 |
| Q9WVJ3 | ***Cpq*** | 2.6 | 23 | 69.2 | 293.7 | 181.7 | 401.7 |
| Q61838 | ***Pzp*** | 225.2 | 609.8 | 641.1 | 1085.8 | 934.9 | 1149.4 |
| Q9D1D4 | ***Tmed10*** | 55.2 | 96.1 | 97.8 | 128.8 | 146.3 | 152.2 |
| Q61207 | ***Psap*** | 536 | 2163.8 | 2051.6 | 3490.5 | 3503.7 | 4367.6 |
| Q05117 | ***Acp5*** | 64.6 | 170.7 | 133.8 | 275.7 | 220.3 | 297.9 |
| Q9Z0M5 | ***Lipa*** | 158.7 | 542.8 | 531.8 | 938.8 | 839.9 | 870.6 |
| P33267 | ***Cyp2f2*** | 590.3 | 958.7 | 923.6 | 1801 | 1488.8 | 1271.4 |
| Q9R013 | ***Ctsf*** | 177.8 | 355.2 | 639 | 811.7 | 982.5 | 925 |
| Q3TCN2 | ***Plbd2*** | 14.1 | 23.8 | 57.2 | 146.4 | 100.6 | 207.3 |
| P32261 | ***Serpinc1*** | 56.2 | 92.5 | 118.7 | 143.2 | 188.1 | 169.5 |
| P12265 | ***Gusb*** | 40.1 | 73.4 | 94.3 | 257.4 | 177 | 365.4 |
| Q07797 | ***Lgals3bp*** | 11.8 | 27 | 47 | 108.4 | 83.5 | 165 |
| P51410 | ***Rpl9*** | 42.3 | 63.7 | 81.2 | 132.3 | 123.6 | 95.3 |
| Q00897 | ***Serpina1d*** | 440.2 | 652.5 | 1403.4 | 1680 | 2279.4 | 2560.4 |
| P47911 | ***Rpl6*** | 444.7 | 522 | 532.8 | 964.1 | 749 | 687.9 |
| O70404 | ***Vamp8*** | 25.4 | 96.8 | 80.4 | 153.7 | 131.6 | 145.7 |
| O09159 | ***Man2b1*** | 81.1 | 237.5 | 291.5 | 676.8 | 459.6 | 894.4 |
| P23953 | ***Ces1c*** | 65 | 99.9 | 216.5 | 357 | 314.8 | 256.9 |
| O88587 | ***Comt*** | 169.6 | 210.4 | 315.2 | 529.4 | 410.7 | 381.6 |
| Q9ET22 | ***Dpp7*** | 6.7 | 27.1 | 85 | 208.2 | 196.8 | 395.2 |
| P49935 | ***Ctsh*** | 100.8 | 325.2 | 498.6 | 1369.4 | 994.4 | 2185.9 |
| Q00896 | ***Serpina1c*** | 56.2 | 87.8 | 205.3 | 280.2 | 288.9 | 435.4 |

| | | | | | | | |
|---|---|---|---|---|---|---|---|
| P24369 | ***Ppib*** | 758.7 | 785.1 | 752.7 | 1285.4 | 1691.1 | 1066.7 |
| Q9CXI5 | ***Manf*** | 238.2 | 255.5 | 317.7 | 384.3 | 547.2 | 411.9 |
| Q91X72 | ***Hpx*** | 120.5 | 77 | 155.2 | 193.4 | 345.7 | 280 |
| Q00623 | ***Apoa1*** | 166 | 440.3 | 699 | 923.7 | 1557.4 | 1103.9 |
| Q80XN0 | ***Bdh1*** | 956.5 | 1045.6 | 473.6 | 305.3 | 199.9 | 282.2 |
| O55142 | ***Rpl35a*** | 166 | 121.4 | 146.7 | 236 | 237.8 | 178.3 |
| P28665 | ***Mug1*** | 260.7 | 531 | 771.7 | 1057.6 | 937.8 | 1351 |
| Q921G7 | ***Etfdh*** | 1861.4 | 1759.3 | 871 | 516.5 | 468.7 | 617.5 |
| P12970 | ***Rpl7a*** | 298.5 | 304.1 | 339.8 | 588.8 | 403.8 | 457.3 |
| P14869 | ***Rplp0*** | 187 | 180.9 | 239.4 | 368.9 | 296.5 | 264.4 |
| Q9Z2I0 | ***Letm1*** | 222.6 | 271.2 | 138.5 | 76.9 | 64.3 | 116.9 |
| Q9DBG1 | ***Cyp27a1*** | 939.5 | 852.7 | 481.9 | 319.7 | 242.7 | 404.4 |
| Q922Q1 | ***2-Mar*** | 1018.1 | 982.3 | 608.5 | 553 | 315.3 | 441.7 |
| P04939 | ***Mup3*** | 99.9 | 41.5 | 221.4 | 292 | 683.5 | 480.6 |
| P51881 | ***Slc25a5*** | 2741.3 | 2290.5 | 1304.9 | 753 | 715.3 | 1053 |
| Q9WUZ9 | ***Entpd5*** | 182.3 | 195.9 | 391.3 | 491.7 | 488.8 | 413.9 |
| P07759 | ***Serpina3k*** | 68.6 | 121 | 282.7 | 309.4 | 496.7 | 616.7 |
| Q60991 | ***Cyp7b1*** | 51.9 | 62.7 | 88.3 | 121.2 | 111.8 | 183.2 |
| Q8BJ64 | ***Chdh*** | 780.5 | 661.4 | 350.5 | 213.7 | 218.6 | 258.3 |
| Q9D023 | ***Mpc2*** | 581.7 | 549.7 | 299.8 | 240.8 | 90.8 | 232.8 |
| Q9DBW0 | ***Cyp4v2*** | 43.7 | 85 | 85.4 | 166 | 114 | 118.3 |
| P06797 | ***Ctsl*** | 48.5 | 140.8 | 155.7 | 247.8 | 481.1 | 285.2 |
| P28798 | ***Grn*** | 81.9 | 391.9 | 357.3 | 663.4 | 632.2 | 1140.2 |
| P47962 | ***Rpl5*** | 82.9 | 71.5 | 102.6 | 144.5 | 159 | 106.1 |
| Q9CR62 | ***Slc25a11*** | 134.7 | 145.1 | 75.3 | 56.9 | 41.5 | 65.6 |
| P07724 | ***Alb*** | 905.8 | 2849.5 | 6159.3 | 6990.3 | 9294.7 | 7598.8 |
| P24270 | ***Cat*** | 1337.6 | 1236.9 | 1428.4 | 3297.7 | 2282.2 | 1905.4 |
| Q9Z2Z6 | ***Slc25a20*** | 502.5 | 597 | 287.4 | 194.3 | 108.3 | 252 |
| P08228 | ***Sod1*** | 66.6 | 212.4 | 1643.5 | 2187.7 | 1957.4 | 1975.2 |
| P50429 | ***Arsb*** | 22.7 | 46 | 56.1 | 149.6 | 97.9 | 247 |
| Q8BGA8 | ***Acsm5*** | 73.5 | 98.4 | 59.4 | 34.8 | 36.6 | 54 |
| Q99KV1 | ***Dnajb11*** | 170.7 | 160.6 | 197.5 | 238.6 | 370.3 | 262.8 |
| P08226 | ***Apoe*** | 185.5 | 557.3 | 1027.3 | 1152.3 | 1355.1 | 1812.7 |
| Q64521 | ***Gpd2*** | 76.4 | 77.6 | 45.3 | 21.7 | 29.6 | 45.7 |
| P35979 | ***Rpl12*** | 251.5 | 255.2 | 424.9 | 582.2 | 605.8 | 413.2 |
| P07356 | ***Anxa2*** | 214 | 451.3 | 287 | 673.7 | 1074.7 | 565.5 |
| O70362 | ***Gpld1*** | 44.1 | 95.5 | 141.9 | 216.1 | 185.4 | 356.2 |
| P17047 | ***Lamp2*** | 253.8 | 1451.2 | 1504.5 | 2229.9 | 1949 | 2610 |
| Q8R086 | ***Suox*** | 224.8 | 180.9 | 102.6 | 62.6 | 75.6 | 82.6 |
| O08677 | ***Kng1*** | 51.7 | 83.6 | 126 | 143.9 | 236.8 | 160.8 |
| Q8CGK3 | ***Lonp1*** | 282.5 | 273 | 136.3 | 92.7 | 95.7 | 125.9 |
| Q8BH59 | ***Slc25a12*** | 510.6 | 540 | 272.8 | 223.1 | 143.8 | 251.1 |
| Q8BMS1 | ***Hadha*** | 3988 | 4383.6 | 1960.9 | 1268 | 1446.1 | 1716.4 |
| Q9WU79 | ***Prodh*** | 292.1 | 307.4 | 157.5 | 139.7 | 82.2 | 138.5 |
| O88668 | ***Creg1*** | 22.6 | 121.5 | 96.9 | 245.3 | 143.2 | 331.4 |
| Q06890 | ***Clu*** | 43.8 | 107.1 | 101.7 | 172.1 | 421.7 | 244.9 |
| Q9QXX4 | ***Slc25a13*** | 2782.8 | 2635.1 | 1158.5 | 911.5 | 653.5 | 992.8 |

| | | | | | | | |
|---|---|---|---|---|---|---|---|
| E9Q414 | ***Apob*** | 35.5 | 96.8 | 88.9 | 128.1 | 112.5 | 156.6 |
| P67984 | ***Rpl22*** | 71.7 | 69.2 | 77.3 | 138.9 | 106.2 | 87.8 |
| P16675 | ***Ctsa*** | 298.1 | 609.2 | 426.7 | 908.2 | 722.1 | 1420.8 |
| Q61335 | ***Bcap31*** | 122.6 | 248.4 | 223.8 | 549.3 | 424.4 | 278.4 |
| Q9DBT9 | ***Dmgdh*** | 241.4 | 204.1 | 251.4 | 312.3 | 305.4 | 450.8 |
| P11438 | ***Lamp1*** | 69.4 | 430.5 | 380.5 | 657.1 | 498.6 | 815.8 |
| P29391 | ***Ftl1*** | 20.2 | 79.8 | 147.9 | 189.6 | 161.3 | 276.4 |
| Q8VEM8 | ***Slc25a3*** | 668.5 | 669.6 | 290.7 | 239.4 | 133 | 266.2 |
| Q9D6M3 | ***Slc25a22*** | 271.9 | 206.9 | 120.3 | 101.2 | 58.5 | 96.7 |
| P52825 | ***Cpt2*** | 1105.2 | 913.5 | 420.6 | 286.5 | 269.2 | 353.3 |
| Q60605 | ***Myl6*** | 18.2 | 110.4 | 201.9 | 187.4 | 429.9 | 419.1 |
| P47915 | ***Rpl29*** | 75.2 | 66 | 89.7 | 128.8 | 101.9 | 93.6 |
| P27659 | ***Rpl3*** | 116.2 | 155.5 | 140.4 | 246.8 | 168.9 | 187.8 |
| P01027 | ***C3*** | 775.3 | 1941.1 | 1689.8 | 3906.9 | 2605.4 | 2315.5 |
| P50544 | ***Acadvl*** | 2325.4 | 1901.2 | 906.7 | 685 | 576.6 | 772.9 |
| P53026 | ***Rpl10a*** | 216.2 | 224.8 | 249.4 | 538.3 | 345.2 | 318.2 |
| Q91WN4 | ***Kmo*** | 741.2 | 613.5 | 344.2 | 289.5 | 234.8 | 306.8 |
| Q9WVD5 | ***Slc25a15*** | 338.9 | 275.2 | 132.2 | 105.6 | 90.7 | 103.3 |
| P15636 | ***LysC*** | 7910.3 | 11419.2 | 5368.5 | 3328.7 | 4153.2 | 4261 |
| Q9CZM2 | ***Rpl15*** | 65.1 | 84.5 | 76.7 | 141.7 | 97.3 | 99.3 |
| Q9CQ01 | ***Rnaset2*** | 69.4 | 175.8 | 188.4 | 380.4 | 233.5 | 569.7 |
| P97742 | ***Cpt1a*** | 1272.6 | 1511.8 | 713.5 | 680.5 | 465.7 | 608.6 |
| Q9JLJ2 | ***Aldh9a1*** | 259.6 | 329.5 | 395.3 | 182.6 | 213.9 | 263.5 |
| P51660 | ***Hsd17b4*** | 300.5 | 445.2 | 351 | 950.3 | 664.5 | 483.3 |
| Q91WG0 | ***Ces2c*** | 138.8 | 171.6 | 416.4 | 398.5 | 485.7 | 495.4 |
| Q8VCM7 | ***Fgg*** | 156.4 | 378.2 | 408.3 | 447.1 | 899 | 669.5 |
| Q8CAQ8 | ***Immt*** | 661 | 498 | 286.6 | 181.1 | 190 | 281.7 |
| P54869 | ***Hmgcs2*** | 4270.2 | 4053.5 | 1830.6 | 1506 | 1412.4 | 1734.6 |
| Q9Z0X1 | ***Aifm1*** | 1454.6 | 1141.7 | 577.3 | 388.2 | 465.1 | 533.3 |
| P21614 | ***Gc*** | 256.8 | 461.2 | 716.3 | 737.6 | 1170.6 | 813.8 |
| P17439 | ***Gba*** | 251.3 | 854.4 | 703.6 | 1063.7 | 963 | 1581.6 |
| P32020 | ***Scp2*** | 545.9 | 1848.5 | 967.4 | 8938.2 | 9586 | 1850.1 |
| O35448 | ***Ppt2*** | 64.8 | 183.5 | 240.3 | 268.2 | 277.3 | 408.7 |
| O55022 | ***Pgrmc1*** | 375.1 | 752 | 797.4 | 1089.5 | 1207.8 | 819.7 |
| P62717 | ***Rpl18a*** | 258.8 | 341.2 | 354.5 | 565.5 | 362 | 472.3 |
| P62897 | ***Cycs*** | 1674.8 | 1193.7 | 467.2 | 265.4 | 266.2 | 416.4 |
| P26150 | ***Hsd3b3*** | 65 | 132.9 | 120.2 | 181.3 | 134.4 | 172.6 |
| Q8JZU2 | ***Slc25a1*** | 301.9 | 275.8 | 137.9 | 120.2 | 78.6 | 148.8 |
| Q9QZD8 | ***Slc25a10*** | 443 | 356.5 | 188.5 | 158.2 | 116.4 | 191.9 |
| Q9D8E6 | ***Rpl4*** | 384.2 | 432.4 | 426.1 | 800.6 | 525.5 | 529.5 |
| Q9D379 | ***Ephx1*** | 210.3 | 499.6 | 579.4 | 828.4 | 760.3 | 585.4 |
| Q8BW75 | ***Maob*** | 2259.5 | 2419 | 922.8 | 990.1 | 695.5 | 712.6 |
| Q9DCU9 | ***Hoga1*** | 100.8 | 51.2 | 91.1 | 129.8 | 100.4 | 149.8 |
| Q8CHT0 | ***Aldh4a1*** | 2537.7 | 2040 | 836.9 | 679.6 | 640.6 | 782.6 |
| Q9R0H0 | ***Acox1*** | 635 | 746 | 639 | 1753.6 | 1086.3 | 894.9 |
| Q91VR2 | ***Atp5c1*** | 1265.5 | 801.8 | 529.4 | 345.8 | 349.9 | 474.6 |
| Q921I1 | ***Tf*** | 1002.3 | 1089.3 | 2174.5 | 2097.2 | 2551.5 | 2213.4 |

| | | | | | | | |
|---|---|---|---|---|---|---|---|
| P14148 | ***Rpl7*** | 328.8 | 359.1 | 367.1 | 677.3 | 421.3 | 468.5 |
| P54071 | ***Idh2*** | 1449.2 | 1280.2 | 464.8 | 391.8 | 362.6 | 484.9 |
| P34914 | ***Ephx2*** | 84.3 | 157.5 | 201.1 | 268.4 | 196.9 | 230.1 |
| Q91VA0 | ***Acsm1*** | 454.5 | 441.4 | 183.1 | 174.6 | 132.6 | 199.5 |
| Q921H8 | ***Acaa1a*** | 113.9 | 263.8 | 154.8 | 928.7 | 967 | 211.4 |
| P20029 | ***Hspa5*** | 2803.5 | 3884.5 | 6559.5 | 6492.9 | 10933.4 | 7063.5 |
| Q02819 | ***Nucb1*** | 134.8 | 214.3 | 252.9 | 259.7 | 395.7 | 286 |
| P68134 | ***Acta1*** | 449 | 1731.4 | 2577.9 | 2176.7 | 4029.3 | 4984.5 |
| P60710 | ***Actb*** | 72.1 | 218.8 | 378 | 331.6 | 518.6 | 742.6 |
| Q99JY0 | ***Hadhb*** | 3104.6 | 2082.1 | 1153.9 | 782.4 | 852 | 1148.4 |
| P56382 | ***Atp5e*** | 322.7 | 184.7 | 141.2 | 91.5 | 103.3 | 112.8 |
| Q05920 | ***Pc*** | 4340.1 | 2961.2 | 1363.2 | 1162.4 | 975.8 | 1217.2 |
| Q9D0M3 | ***Cyc1*** | 521.5 | 365.8 | 217.2 | 145.3 | 190.4 | 216.3 |
| Q9WVA2 | ***Timm8a1*** | 578.7 | 332.3 | 171.9 | 65 | 157.6 | 119.9 |
| Q8VCU1 | ***Ces3b*** | 264.3 | 316.7 | 605.3 | 660.6 | 708.2 | 536.1 |
| P11588 | ***Mup1*** | 1985.2 | 986.6 | 5310.9 | 3831.6 | 8217.6 | 8051.5 |
| Q9CQ69 | ***Uqcrq*** | 482.5 | 315.7 | 174.2 | 113.3 | 152 | 159.5 |
| E9PV24 | ***Fga*** | 306.9 | 810.6 | 815.1 | 845.6 | 2042.7 | 1374.7 |
| P27046 | ***Man2a1*** | 18.4 | 57.5 | 28.6 | 51.9 | 60.4 | 85.3 |
| Q8BWT1 | ***Acaa2*** | 6507 | 4263.2 | 2272.9 | 1535.7 | 1958.6 | 2090.1 |
| P01878 | | 44.1 | 330.9 | 192.2 | 499.6 | 261.4 | 496.4 |
| Q9QWR8 | ***Naga*** | 9.4 | 63.5 | 53.8 | 107.7 | 57.3 | 149.2 |
| P29699 | ***Ahsg*** | 56.1 | 162.9 | 133.7 | 172.7 | 340.4 | 202.6 |
| P51150 | ***Rab7a*** | 81.3 | 179.9 | 130.7 | 215.9 | 193.6 | 169.9 |
| Q8BMF4 | ***Dlat*** | 447.5 | 346.3 | 193.2 | 148.2 | 153.1 | 223.2 |
| Q9CXZ1 | ***Ndufs4*** | 566.8 | 257.7 | 246 | 118 | 126.1 | 188.2 |
| P46638 | ***Rab11b*** | 20.4 | 72.6 | 81.3 | 84.7 | 109.6 | 101 |
| Q9D1Q6 | ***Erp44*** | 310.5 | 398.9 | 483.7 | 535 | 711.7 | 467.9 |
| Q64FW2 | ***Retsat*** | 24.6 | 49 | 41.5 | 83.2 | 63.3 | 45.5 |
| Q922R8 | ***Pdia6*** | 375.2 | 544.4 | 735 | 717.9 | 1030.1 | 752 |
| Q8CHQ9 | ***Cml2*** | 105.9 | 238.3 | 196 | 289.1 | 222.5 | 297.5 |
| Q9CQQ7 | ***Atp5f1*** | 1457.3 | 917.2 | 468.6 | 328.3 | 407.4 | 421.5 |
| Q60759 | ***Gcdh*** | 1523 | 1107.2 | 484.4 | 411.5 | 412.3 | 512.4 |
| P70699 | ***Gaa*** | 88.7 | 511.1 | 216.2 | 598.9 | 419.2 | 904.5 |
| P42125 | ***Eci1*** | 1077.3 | 894 | 369.9 | 329.4 | 328.4 | 430.5 |
| Q5FW60 | ***Mup20*** | 77.9 | 51.6 | 105.6 | 108.7 | 297 | 166.7 |
| Q8VDD5 | ***Myh9*** | 51.2 | 328.8 | 343 | 332.3 | 520.9 | 723.3 |
| Q8K0E8 | ***Fgb*** | 159.3 | 462 | 475.3 | 476.7 | 1206.4 | 770.8 |
| P16406 | ***Enpep*** | 14.7 | 141.2 | 163.4 | 208 | 161.5 | 267.8 |
| Q99PG0 | ***Aadac*** | 475.3 | 803.6 | 773.5 | 1548.6 | 1011.2 | 841 |
| P97450 | ***Atp5j*** | 3765.6 | 1724.4 | 1087.3 | 477.7 | 865.2 | 606.3 |
| P61255 | ***Rpl26*** | 62.4 | 61 | 82 | 139.4 | 80.6 | 93.3 |
| Q9DCZ4 | ***Apoo*** | 195.9 | 134.3 | 62.9 | 42 | 57.9 | 71.2 |
| P52480 | ***Pkm*** | 26 | 122.8 | 155.6 | 125 | 244.4 | 258.3 |
| O09167 | ***Rpl21*** | 217.9 | 200.2 | 259 | 499 | 249.6 | 347.9 |
| P05202 | ***Got2*** | 609.6 | 483.6 | 666.5 | 761.5 | 675.1 | 1086.8 |
| Q9CQN1 | ***Trap1*** | 413.7 | 345 | 129.9 | 136.4 | 119 | 154.6 |

| | | | | | | | |
|---|---|---|---|---|---|---|---|
| Q9CZ13 | ***Uqcrc1*** | 1639.8 | 958.6 | 568.8 | 388.9 | 502 | 525 |
| Q99LC5 | ***Etfa*** | 2625.6 | 1891.6 | 903 | 715.2 | 932 | 977.5 |
| P22315 | ***Fech*** | 100.9 | 73.2 | 40.1 | 32.7 | 41.8 | 41.2 |
| Q5FW57 | ***Gm4952*** | 378.1 | 211.9 | 120.9 | 90.7 | 110.5 | 94.1 |
| P56391 | ***Cox6b1*** | 1661.7 | 950.5 | 537.1 | 357.6 | 438.1 | 539.5 |
| P01872 | ***Ighm*** | 195.9 | 197.3 | 306.6 | 521 | 414.5 | 245.4 |
| Q9WV54 | ***Asah1*** | 199.1 | 820.7 | 643.7 | 759.2 | 977.1 | 1031.5 |
| Q9WTP6 | ***Ak2*** | 1203.5 | 620.5 | 376.6 | 234.5 | 271.5 | 347 |
| P52503 | ***Ndufs6*** | 569.3 | 255.4 | 182.1 | 85.4 | 138.7 | 133.3 |
| Q9D172 | ***D10Jhu81e*** | 482.8 | 477.5 | 320.3 | 228.3 | 222.1 | 401.4 |
| Q8VCH0 | ***Acaa1b*** | 80.1 | 232.9 | 130.3 | 915.9 | 643.8 | 137.7 |
| Q8C7E7 | ***Stbd1*** | 29.5 | 67.7 | 60.4 | 120 | 170.4 | 56.2 |
| Q9DB05 | ***Napa*** | 36.9 | 102.4 | 100.9 | 106.8 | 119.2 | 167.8 |
| Q8BWF0 | ***Aldh5a1*** | 313.2 | 186.2 | 91.4 | 77.1 | 84.5 | 84.2 |
| Q8K2B3 | ***Sdha*** | 3816.5 | 2047.5 | 1179 | 778.6 | 1023.9 | 1065.3 |
| Q91VS7 | ***Mgst1*** | 464.6 | 457.6 | 524.1 | 864.6 | 625.2 | 516.6 |
| Q3ULD5 | ***Mccc2*** | 147.4 | 95.2 | 59.4 | 58.5 | 42.5 | 61.7 |
| P41105 | ***Rpl28*** | 122.2 | 188.2 | 233.2 | 316.2 | 215.1 | 245.1 |
| Q03265 | ***Atp5a1*** | 8767 | 4990.9 | 3693.2 | 2327.4 | 3485.1 | 3376.1 |
| P11352 | ***Gpx1*** | 381.1 | 249.9 | 449.4 | 496.4 | 501.1 | 420.4 |
| P00186 | ***Cyp1a2*** | 205.9 | 559.5 | 461.3 | 858.1 | 463.4 | 728.4 |
| P51863 | ***Atp6v0d1*** | 40 | 265.1 | 147.4 | 278.8 | 217.6 | 360.1 |
| Q9D3D9 | ***Atp5d*** | 513.8 | 265 | 211.8 | 113.9 | 136.9 | 224.7 |
| Q8BFR5 | ***Tufm*** | 1394.1 | 882.5 | 501 | 417.9 | 466.1 | 536.1 |
| P14211 | ***Calr*** | 2368 | 2030.2 | 3893.5 | 3631.1 | 5054.7 | 3524 |
| Q99MR8 | ***Mccc1*** | 341.2 | 219.3 | 123.6 | 125.5 | 105.7 | 125.5 |
| Q9EQ20 | ***Aldh6a1*** | 2419.3 | 1450.3 | 717.5 | 649.1 | 585.7 | 802.6 |
| Q9JKR6 | ***Hyou1*** | 629.8 | 1285.4 | 1416.6 | 1480.1 | 2126.8 | 1421.1 |
| Q91V41 | ***Rab14*** | 37.2 | 101.1 | 79.6 | 132.4 | 107.4 | 91.5 |
| Q8VCW8 | ***Acsf2*** | 742.2 | 602.4 | 220 | 216.6 | 221.3 | 314.7 |
| O35129 | ***Phb2*** | 1491.8 | 807.6 | 465.7 | 334.6 | 397.2 | 481.6 |
| B5X0G2 | ***Mup17*** | 49.5 | 10.6 | 126.5 | 82.4 | 152.5 | 238.2 |
| Q9CPQ1 | ***Cox6c*** | 1228 | 820.1 | 415.7 | 343.3 | 442 | 476 |
| Q5XG73 | ***Acbd5*** | 58.4 | 82.3 | 64.1 | 181.5 | 119.1 | 70.5 |
| Q6P3A8 | ***Bckdhb*** | 333.2 | 141.6 | 91 | 74.7 | 47.5 | 69.3 |
| Q99L04 | ***Dhrs1*** | 183 | 233.9 | 212.1 | 278.6 | 230.2 | 230.3 |
| Q9CY27 | ***Tecr*** | 153.9 | 361.9 | 230.6 | 435.5 | 433.2 | 279.1 |
| Q91ZA3 | ***Pcca*** | 784.2 | 576.9 | 345.4 | 318.7 | 274.8 | 433.8 |
| P18760 | ***Cfl1*** | 43.5 | 147.4 | 288.1 | 201.8 | 347.7 | 376.5 |
| P99028 | ***Uqcrh*** | 1432.6 | 613.3 | 459.8 | 240.9 | 371.9 | 364.8 |
| Q9DC69 | ***Ndufa9*** | 154.3 | 94.5 | 49.9 | 37.6 | 50.7 | 57.1 |
| Q8JZZ0 | ***Ugt3a2*** | 242.8 | 424.8 | 332.8 | 518 | 473.5 | 360.1 |
| Q9DCW4 | ***Etfb*** | 3369 | 3156.7 | 1254.7 | 1184.1 | 1728.7 | 1446.3 |
| P19253 | ***Rpl13a*** | 68.1 | 93 | 109.7 | 159.1 | 97.9 | 120.6 |
| Q9CPP6 | ***Ndufa5*** | 416.4 | 166.1 | 152.2 | 73.9 | 109.9 | 121.8 |
| P35486 | ***Pdha1*** | 222.2 | 144.6 | 88.6 | 65.8 | 58.4 | 118.7 |
| Q9DB20 | ***Atp5o*** | 1107.4 | 532.4 | 415.6 | 272 | 360.2 | 361.4 |

| | | | | | | | |
|---|---|---|---|---|---|---|---|
| Q7TNG8 | ***Ldhd*** | 396.6 | 233.3 | 104.8 | 89.8 | 82.5 | 138.4 |
| Q4LDG0 | ***Slc27a5*** | 677.6 | 1314.2 | 1101.9 | 2150.4 | 1306.6 | 1299.9 |
| Q9CQ75 | ***Ndufa2*** | 213.4 | 115.8 | 73.4 | 38.7 | 81.9 | 69.1 |
| Q8C5H8 | ***Nadk2*** | 246.3 | 149.4 | 73.7 | 58.8 | 86.7 | 76.2 |
| Q9CR68 | ***Uqcrfs1*** | 2118 | 827.1 | 587.6 | 293.9 | 416.8 | 491.4 |
| Q9CRB9 | ***Chchd3*** | 437.7 | 201.4 | 167.5 | 103.9 | 129.4 | 154.2 |
| Q8BSY0 | ***Asph*** | 70.7 | 79 | 57.6 | 87.7 | 83.6 | 72.9 |
| Q922D8 | ***Mthfd1*** | 221.2 | 478.4 | 733.8 | 1352.1 | 1006.2 | 487.9 |
| P67778 | ***Phb*** | 1962.2 | 806.3 | 631.7 | 369.5 | 504 | 499 |
| Q8R143 | ***Pttg1ip*** | 10.9 | 60.9 | 81.4 | 74.7 | 80.4 | 129.4 |
| Q6ZWV3 | ***Rpl10*** | 109.2 | 182.5 | 182.4 | 297.1 | 193.3 | 190 |
| Q8QZT1 | ***Acat1*** | 1634.5 | 1009.8 | 609.1 | 547.7 | 564 | 689.3 |
| Q60597 | ***Ogdh*** | 334.7 | 197.8 | 108.8 | 91.3 | 87.3 | 137.5 |
| P02089 | ***Hbb-b2*** | 26.4 | 35.1 | 34.3 | 37 | 87.4 | 46.8 |
| O08547 | ***Sec22b*** | 41.6 | 93.7 | 68.6 | 117.8 | 101 | 76.9 |
| Q9CW42 | ***42795*** | 538 | 518.2 | 260.2 | 341 | 169.1 | 309.5 |
| Q9Z2I9 | ***Sucla2*** | 780.2 | 467.9 | 236.8 | 226.3 | 208.5 | 289 |
| Q62425 | ***Ndufa4*** | 2430.5 | 885.4 | 667.9 | 348 | 529.5 | 459.8 |
| Q8BK48 | ***Ces2e*** | 138.2 | 226.9 | 388.3 | 383.8 | 463 | 311.2 |
| P12787 | ***Cox5a*** | 1362.3 | 605.1 | 450.8 | 268.3 | 399.7 | 400.8 |
| Q8R1I1 | ***Uqcr10*** | 518.2 | 200.8 | 161.7 | 108.6 | 115.1 | 123.1 |
| Q99LC3 | ***Ndufa10*** | 1845.4 | 859.4 | 537.8 | 334.1 | 463.8 | 567 |
| Q9DCX2 | ***Atp5h*** | 4109.4 | 1500.8 | 1281.1 | 702.5 | 849.8 | 1049.6 |
| Q9Z2I8 | ***Suclg2*** | 1400.1 | 682 | 335.9 | 250.3 | 345.4 | 347.3 |
| P29758 | ***Oat*** | 534.1 | 297.1 | 525.9 | 720.8 | 486 | 1204.4 |
| Q8C196 | ***Cps1*** | 41642.2 | 23667.6 | 8335.2 | 9279.6 | 11386.5 | 7801.9 |
| P97872 | ***Fmo5*** | 610.5 | 1307.2 | 1715.5 | 2462.8 | 1440.9 | 1785.6 |
| P50136 | ***Bckdha*** | 534.9 | 214.4 | 155.8 | 105.1 | 116.4 | 133.5 |
| P50171 | ***Hsd17b8*** | 181.2 | 79.7 | 56.1 | 40.8 | 47.4 | 49.7 |
| Q91VD9 | ***Ndufs1*** | 915 | 454.5 | 312.8 | 218 | 252.1 | 347.8 |
| P38060 | ***Hmgcl*** | 204.7 | 137.4 | 94.4 | 72 | 117.5 | 85.3 |
| P56395 | ***Cyb5a*** | 1254.5 | 1325.2 | 2085.4 | 2512.7 | 3354.4 | 1531.9 |
| Q9CQR4 | ***Acot13*** | 651.4 | 305.3 | 169.9 | 140.9 | 158.4 | 164.1 |
| Q9CQA3 | ***Sdhb*** | 2175.7 | 993.8 | 618.4 | 421.4 | 646.1 | 532.1 |
| Q91XE8 | ***Tmem205*** | 206.4 | 495.6 | 386.3 | 656.5 | 495.3 | 432 |
| P26039 | ***Tln1*** | 20.7 | 133.9 | 121 | 106.4 | 173.3 | 215.9 |
| Q8K0C4 | ***Cyp51a1*** | 93.1 | 242.7 | 190.4 | 262 | 261.1 | 215.8 |
| P28230 | ***Gjb1*** | 13.1 | 57.2 | 105.1 | 100.6 | 80.2 | 186.2 |
| Q9DCS9 | ***Ndufb10*** | 236 | 96.9 | 69.6 | 35.5 | 63.1 | 65.2 |
| P97807 | ***Fh*** | 513.8 | 274.1 | 187.3 | 141.3 | 151.8 | 223.5 |
| Q9CQH3 | ***Ndufb5*** | 439.4 | 233.2 | 138.4 | 95.6 | 131.8 | 171.4 |
| P56480 | ***Atp5b*** | 6564.2 | 2792.5 | 2551.2 | 1363.7 | 2358.2 | 2172.5 |
| P58710 | ***Gulo*** | 230.6 | 582.3 | 494.9 | 760.6 | 512.7 | 610 |
| Q99KF1 | ***Tmed9*** | 129.3 | 302.3 | 231.9 | 293.8 | 316.2 | 283.1 |
| P63101 | ***Ywhaz*** | 13.6 | 52 | 128.1 | 76.3 | 139.6 | 183.4 |
| P38647 | ***Hspa9*** | 2471.9 | 1592.2 | 1026.9 | 874.8 | 930.7 | 1308.3 |
| P08003 | ***Pdia4*** | 1628 | 1947.1 | 3091.2 | 2842.9 | 4487.3 | 2625 |

| | | | | | | | |
|---|---|---|---|---|---|---|---|
| Q921X9 | ***Pdia5*** | 517.1 | 496.9 | 260.9 | 622.2 | 738.2 | 443.1 |
| Q8BH95 | ***Echs1*** | 2973.4 | 1320.6 | 785.4 | 587.9 | 785.7 | 778.7 |
| P99027 | ***Rplp2*** | 210.3 | 179.3 | 337 | 346.1 | 744.7 | 297.1 |
| Q64176 | ***Ces1e*** | 326.5 | 523.1 | 760.4 | 954.2 | 804.2 | 582.5 |
| P37040 | ***Por*** | 479.6 | 1336.2 | 1187.1 | 2302.9 | 1709.9 | 1004.2 |
| P63038 | ***Hspd1*** | 10589.4 | 5003.7 | 2882.1 | 2173.5 | 3226.2 | 2936 |
| Q91W90 | ***Txndc5*** | 116.4 | 152.9 | 185.5 | 176.5 | 209.9 | 170.3 |
| Q9DB77 | ***Uqcrc2*** | 7286.4 | 3065 | 1867.7 | 1384.7 | 1753 | 1919.1 |
| Q80XL6 | ***Acad11*** | 73 | 153.1 | 90.5 | 200.4 | 248 | 92.9 |
| P53395 | ***Dbt*** | 519.2 | 226.2 | 151.4 | 98.6 | 160.7 | 145.5 |
| P19536 | ***Cox5b*** | 1728.9 | 945.5 | 573.5 | 303.1 | 871.1 | 428.2 |
| O08749 | ***Dld*** | 615.6 | 398 | 244.5 | 222.3 | 197.2 | 339.6 |
| P01029 | ***C4b*** | 27.9 | 53.9 | 38.2 | 51 | 56 | 47.1 |
| Q9EQ06 | ***Hsd17b11*** | 122.1 | 222.1 | 202.3 | 400.2 | 190 | 252 |
| Q9CQX2 | ***Cyb5b*** | 604.7 | 676.4 | 397.5 | 503.6 | 340.7 | 426.8 |
| Q9QXE0 | ***Hacl1*** | 81.1 | 189.4 | 181.1 | 302.1 | 212.8 | 163.4 |
| Q9DCY0 | ***Keg1*** | 573.9 | 166.2 | 128.8 | 64.8 | 110.2 | 86.9 |
| Q91ZX7 | ***Lrp1*** | 19.7 | 90.3 | 99.3 | 104.2 | 86.6 | 176.4 |
| Q99LP6 | ***Grpel1*** | 387.7 | 190.3 | 139.7 | 117.8 | 98.1 | 170.5 |
| Q9WUM5 | ***Suclg1*** | 1467.2 | 775.9 | 471.2 | 402.2 | 543.9 | 516.2 |
| P62075 | ***Timm13*** | 787.3 | 218.3 | 211.7 | 109.9 | 134.4 | 168.2 |
| P09528 | ***Fth1*** | 41.2 | 183.3 | 433 | 375.7 | 309 | 507.7 |
| Q9DBL1 | ***Acadsb*** | 321.6 | 167.4 | 84 | 76.3 | 78.6 | 122.8 |
| Q9WTP7 | ***Ak3*** | 2611.9 | 940.4 | 605.4 | 430.5 | 517.2 | 632.9 |
| Q9D6J6 | ***Ndufv2*** | 618.1 | 265.7 | 234.9 | 135 | 207.5 | 249.7 |
| Q07417 | ***Acads*** | 388.1 | 290.2 | 171.7 | 158.5 | 188.2 | 231.3 |
| Q9DD20 | ***Mettl7b*** | 1047.7 | 1096.9 | 1121.2 | 1615 | 1225 | 1084.5 |
| Q8R164 | ***Bphl*** | 938.7 | 417.9 | 194 | 175.1 | 217.2 | 248 |
| P51174 | ***Acadl*** | 739.1 | 864.8 | 270 | 347.8 | 352.2 | 426.8 |
| P41216 | ***Acsl1*** | 1037.6 | 1891.4 | 1330.9 | 2605 | 1738.5 | 1572.2 |
| P62702 | ***Rps4x*** | 10.2 | 90.1 | 82 | 97.8 | 136.2 | 74.1 |
| Q91X77 | ***Cyp2c50*** | 28 | 60.4 | 66.6 | 123.8 | 47.5 | 85 |
| Q61694 | ***Hsd3b5*** | 325 | 288.6 | 231.6 | 201.1 | 91.8 | 293.2 |
| O55125 | ***Nipsnap1*** | 361 | 128.7 | 116.9 | 81.3 | 93.6 | 111.8 |
| Q61425 | ***Hadh*** | 2265.1 | 951.5 | 580.3 | 553 | 589.2 | 610 |
| P62918 | ***Rpl8*** | 311 | 473.1 | 375.9 | 733.8 | 394.4 | 480.6 |
| Q9WUR2 | ***Eci2*** | 222.3 | 177.7 | 112.2 | 158.1 | 108 | 98.7 |
| Q99KB8 | ***Hagh*** | 152.3 | 107.9 | 151.9 | 82.1 | 115.3 | 132 |
| P17717 | ***Ugt2b17*** | 322.3 | 687.4 | 549.2 | 906.3 | 596.4 | 631.2 |
| Q9D2G2 | ***Dlst*** | 791.7 | 310.4 | 253.5 | 166.2 | 219.1 | 283 |
| Q60648 | ***Gm2a*** | 93.2 | 334.2 | 317.2 | 366.2 | 294.4 | 424.3 |
| P28843 | ***Dpp4*** | 68.7 | 627.4 | 668.7 | 747.9 | 512.9 | 1269.8 |
| Q8K3J1 | ***Ndufs8*** | 158.6 | 52.9 | 54.8 | 33.7 | 49.7 | 44.9 |
| Q63880 | ***Ces3a*** | 413.2 | 439.8 | 968.2 | 1102.7 | 697.2 | 865.7 |
| Q8QZR3 | ***Ces2a*** | 897.4 | 1210.5 | 2440.5 | 2628.7 | 2239 | 1757.9 |
| Q05421 | ***Cyp2e1*** | 866.8 | 2781.5 | 2453.1 | 4543.4 | 2352.4 | 2702.1 |
| Q9D6Y7 | ***Msra*** | 65.9 | 37.4 | 78.1 | 25.1 | 47.5 | 52.7 |

| | | | | | | | |
|---|---|---|---|---|---|---|---|
| Q9D855 | ***Uqcrb*** | 1338.6 | 423.4 | 382.4 | 208.9 | 423.7 | 304.4 |
| P52196 | ***Tst*** | 1882 | 642.8 | 376.8 | 371.8 | 327.2 | 446.4 |
| Q64459 | ***Cyp3a11*** | 572.7 | 2328.4 | 1714.3 | 2559.6 | 1897.2 | 2230.5 |
| P26443 | ***Glud1*** | 9814.6 | 2908.1 | 3882.2 | 2501.5 | 3123.9 | 2805.7 |
| UPSP:TRYP_PIG | | 1577 | 2288.6 | 1806.6 | 465.6 | 2265.1 | 643.7 |
| Q9DCJ5 | ***Ndufa8*** | 954.3 | 353.8 | 289.6 | 180.3 | 286.5 | 319.4 |
| P27773 | ***Pdia3*** | 2509.9 | 2158.4 | 3225.4 | 2819.3 | 4653.2 | 2940.5 |
| Q99K67 | ***Aass*** | 415 | 393.5 | 141.7 | 197.6 | 209.6 | 213 |
| P55096 | ***Abcd3*** | 677.1 | 1209.9 | 772.4 | 1859.4 | 980.8 | 1043.1 |
| Q91XE0 | ***Glyat*** | 1761 | 650.3 | 386.1 | 353.1 | 425.4 | 446.4 |
| P11725 | ***Otc*** | 2755.1 | 2017.6 | 838.8 | 1092.8 | 1221.4 | 1213.2 |
| O35114 | ***Scarb2*** | 157.7 | 1184.6 | 723.2 | 1383.2 | 737.1 | 1270.6 |
| Q8CFX1 | ***H6pd*** | 88 | 180.8 | 207.2 | 236.8 | 220.1 | 171.5 |
| O35488 | ***Slc27a2*** | 490.9 | 1077.7 | 926.1 | 1790.4 | 946.1 | 968.3 |
| Q91WL5 | ***Cyp4a12a*** | 32.5 | 158.3 | 187.9 | 205 | 124.1 | 338.4 |
| P35564 | ***Canx*** | 388.2 | 844.8 | 645.5 | 972.8 | 716.2 | 755.9 |
| Q9DCT2 | ***Ndufs3*** | 350.5 | 139.7 | 110.6 | 90.6 | 100.1 | 129.8 |
| Q9D1R9 | ***Rpl34*** | 115.9 | 118.8 | 111.1 | 201.3 | 116.5 | 126.3 |
| P20108 | ***Prdx3*** | 428.6 | 297 | 269 | 259.3 | 234.7 | 305.4 |
| Q9CZS1 | ***Aldh1b1*** | 535.6 | 188.9 | 79.6 | 90.2 | 134.8 | 73.3 |
| Q8BHN3 | ***Ganab*** | 410.8 | 583.2 | 725 | 899.7 | 676.7 | 604.5 |
| P47738 | ***Aldh2*** | 3140.5 | 3082.3 | 1875.6 | 1743.7 | 1798.1 | 2709.3 |
| P35505 | ***Fah*** | 79.8 | 88.3 | 206.5 | 58.2 | 54.9 | 105.2 |
| Q64433 | ***Hspe1*** | 1304.2 | 1124 | 797.1 | 611.3 | 1083.4 | 814.5 |
| P62204 | ***Calm1; Calm2; Calm3*** | 10.6 | 50.5 | 153.2 | 67.6 | 160.1 | 181.8 |
| P56654 | ***Cyp2c37*** | 550.8 | 1457.3 | 1344.5 | 2454.8 | 1119.8 | 1506.2 |
| P09103 | ***P4hb*** | 3229.1 | 3087.1 | 5975 | 5100.9 | 7951.1 | 4342.3 |
| UPSP:K2C1_HUMAN | | 2328.7 | 4233.8 | 5576.5 | 3206.9 | 2491.9 | 3032.2 |
| Q8VCI0 | ***Plbd1*** | 49.9 | 139.4 | 96 | 126.7 | 100.8 | 179.1 |
| Q99LY9 | ***Ndufs5*** | 300.8 | 70.1 | 105.6 | 54.5 | 75.1 | 93.9 |
| P61922 | ***Abat*** | 75.1 | 102.7 | 33.2 | 40.5 | 43.1 | 55.4 |
| Q6XVG2 | ***Cyp2c54*** | 179.6 | 487.2 | 381.1 | 741.3 | 331.1 | 494 |
| P19783 | ***Cox4i1*** | 1039.9 | 338.4 | 188.9 | 165.4 | 284 | 202.7 |
| P56593 | ***Cyp2a12*** | 564 | 1318.8 | 888.5 | 1673.3 | 1128.6 | 1005.3 |
| Q9CQ62 | ***Decr1*** | 1096.5 | 401.9 | 229 | 246.6 | 275.3 | 296.1 |
| Q9CZU6 | ***Cs*** | 433 | 609.1 | 276.5 | 264.7 | 215.5 | 437.6 |
| Q99J99 | ***Mpst*** | 178 | 44.4 | 26.3 | 21.1 | 26.4 | 37.5 |
| P08113 | ***Hsp90b1*** | 2362 | 2823 | 4101.9 | 3578.9 | 4655.5 | 3311 |
| P45878 | ***Fkbp2*** | 85 | 59.3 | 104.9 | 117.5 | 96.4 | 88.2 |
| Q64458 | ***Cyp2c29*** | 556.2 | 1847.4 | 1380.4 | 2326.6 | 1399.5 | 1605.1 |
| P35441 | ***Thbs1*** | 57.4 | 228.1 | 205.8 | 102.1 | 385 | 342.9 |
| Q99JI6 | ***Rap1b*** | 48.1 | 173.1 | 249.5 | 196.1 | 198.7 | 296.8 |
| Q9DB15 | ***Mrpl12*** | 175.8 | 65.4 | 54.6 | 34.8 | 63.2 | 67.8 |
| Q8CIM7 | ***Cyp2d26*** | 319.2 | 979.4 | 942.6 | 1711.6 | 1052.2 | 677.8 |
| Q9DBM2 | ***Ehhadh*** | 391.2 | 731.7 | 357.6 | 1449.4 | 733.8 | 383.9 |
| P54116 | ***Stom*** | 33.7 | 162.3 | 103.7 | 152 | 107.4 | 177.6 |
| O35887 | ***Calu*** | 227.7 | 314.6 | 316.9 | 267 | 417.8 | 344.4 |

| | | | | | | | |
|---|---|---|---|---|---|---|---|
| P20852 | ***Cyp2a5*** | 172.2 | 786.7 | 534 | 1201.7 | 598.7 | 565.9 |
| P19157 | ***Gstp1*** | 36.8 | 242.3 | 1084.8 | 343 | 1076.9 | 1349.5 |
| Q9Z1P6 | ***Ndufa7*** | 134.3 | 64.6 | 45.7 | 36.9 | 55.6 | 63.4 |
| Q9JM62 | ***Reep6*** | 63.5 | 122.1 | 129 | 255.3 | 154.4 | 78.7 |
| P51658 | ***Hsd17b2*** | 188.3 | 485.6 | 316.4 | 574.4 | 376 | 379.6 |
| Q9D1G1 | ***Rab1b*** | 117.3 | 291.3 | 272.1 | 262.4 | 259.2 | 359.3 |
| P47963 | ***Rpl13*** | 197 | 210.4 | 331.7 | 299.7 | 268.2 | 310 |
| P09671 | ***Sod2*** | 453.7 | 237 | 309.9 | 346.4 | 375 | 588.6 |
| Q6ZQI3 | ***Mlec*** | 96.8 | 118.3 | 68.8 | 144.5 | 93.1 | 110.7 |
| Q99MZ7 | ***Pecr*** | 392 | 308.2 | 232.3 | 510.8 | 365 | 297.5 |
| Q9CQZ5 | ***Ndufa6*** | 257.4 | 74.1 | 59.1 | 44.9 | 72.9 | 74.7 |
| P57780 | ***Actn4*** | 20 | 53 | 139.1 | 74.1 | 91.3 | 250.9 |
| P24456 | ***Cyp2d10*** | 221.1 | 630.4 | 734 | 1141.2 | 666.7 | 521.5 |
| UPSP:K1CI_HUMAN | | 1353.1 | 2038.6 | 3522.1 | 1558 | 1163.2 | 2095.2 |
| P58252 | ***Eef2*** | 123 | 202.2 | 279.3 | 256.9 | 234.6 | 250.2 |
| Q3UEG6 | ***Agxt2*** | 173.9 | 182.7 | 137.1 | 121.5 | 116.3 | 180.8 |
| Q8VCT4 | ***Ces1d*** | 1049.6 | 1569.3 | 2786.6 | 3161.6 | 2769.1 | 1568.9 |
| Q9CPT4 | ***Mydgf*** | 80.9 | 66 | 204.3 | 139.7 | 231.7 | 137.8 |
| Q9D051 | ***Pdhb*** | 392.1 | 189 | 162.7 | 136.6 | 158.6 | 227 |
| P62806 | ***Hist1h4a; Hist1h4b; Hist1h4c; Hist1h4d; Hist1h4f; Hist1h4h; Hist1h4i; Hist1h4j; Hist1h4k; Hist1h4m; Hist2h4a; Hist4h4*** | 80.4 | 86.7 | 72.4 | 69.6 | 93 | 118.7 |
| Q9EP75 | ***Cyp4f14*** | 29.3 | 110.1 | 71.5 | 133.7 | 80.8 | 79.4 |
| O88844 | ***Idh1*** | 31.5 | 122.1 | 357.4 | 259.8 | 454.6 | 161 |
| Q99KI0 | ***Aco2*** | 917.4 | 611.3 | 304.1 | 359 | 363 | 575.2 |
| Q9JMA7 | ***Cyp3a41a; Cyp3a41b*** | 200.9 | 1367.4 | 412 | 680.8 | 85.5 | 65 |
| Q924Z4 | ***Cers2*** | 60.2 | 120.1 | 85.2 | 145.1 | 100.7 | 88.1 |
| Q63836 | ***Selenbp2*** | 47.3 | 164.3 | 692.7 | 371 | 381.9 | 844.4 |
| O88451 | ***Rdh7*** | 445.9 | 1253.3 | 1355.7 | 1985 | 1375.4 | 858.8 |
| O08795 | ***Prkcsh*** | 258.7 | 377.9 | 467.7 | 403.2 | 674.9 | 343.6 |
| Q8VCC2 | ***Ces1*** | 682.5 | 995.3 | 1429.9 | 1748.7 | 1378.7 | 872.6 |
| P04943 | | 1422.3 | 2311.3 | 1347.6 | 1585.1 | 1165.7 | 1454.8 |
| P16460 | ***Ass1*** | 426.1 | 3990.1 | 6768.2 | 13685.3 | 3452.6 | 4007.2 |
| Q3UV17 | ***Krt76*** | 200.1 | 416.7 | 555.4 | 342.6 | 268.9 | 283.1 |
| P01942 | ***Hba*** | 1606.5 | 246.5 | 1154.1 | 254.4 | 459.4 | 1076.1 |
| P97501 | ***Fmo3*** | 99.9 | 523.1 | 272.4 | 339.1 | 118.7 | 46.7 |
| UPSP:K1CJ_HUMAN | | 1243.2 | 2591.7 | 2829.8 | 1825.5 | 2078.3 | 1384.9 |
| O70503 | ***Hsd17b12*** | 98.7 | 211.9 | 145.5 | 229.1 | 165.9 | 161.2 |
| Q9D0F3 | ***Lman1*** | 226.2 | 913.4 | 644.5 | 796.4 | 815.3 | 688.6 |
| P83882 | ***Rpl36a*** | 109.4 | 102.8 | 136.6 | 183.5 | 91.7 | 145.6 |
| Q9WV55 | ***Vapa*** | 123.3 | 179.1 | 103.5 | 191.5 | 142.8 | 141.1 |

| | | | | | | | |
|---|---|---|---|---|---|---|---|
| P99029 | ***Prdx5*** | 482.6 | 346.8 | 566 | 289.6 | 373.6 | 508.4 |
| Q01339 | ***Apoh*** | 97.7 | 208.9 | 320.2 | 198.5 | 285.9 | 325.3 |
| Q62452 | ***Ugt1a9*** | 88.1 | 207.5 | 112.6 | 242.9 | 127.8 | 160.1 |
| P13707 | ***Gpd1*** | 30 | 121.5 | 215.4 | 189.1 | 147.2 | 164.2 |
| P26043 | ***Rdx*** | 27.4 | 94.3 | 177.6 | 87.9 | 172.1 | 163.5 |
| Q91W64 | ***Cyp2c70*** | 371.1 | 908.7 | 547.7 | 947.2 | 516.5 | 863.7 |
| P45952 | ***Acadm*** | 753.2 | 426 | 324.7 | 344.5 | 372.2 | 462.4 |
| O54734 | ***Ddost*** | 289.9 | 275.3 | 163.8 | 312.5 | 334.6 | 212.8 |
| Q9QXT0 | ***Cnpy2*** | 100.3 | 112.7 | 240.6 | 179.7 | 219.4 | 168 |
| Q91YH5 | ***Atl3*** | 32.2 | 88.1 | 72.4 | 124 | 55.3 | 74.9 |
| Q9WVL0 | ***Gstz1*** | 58.5 | 144 | 232.7 | 117.7 | 56.1 | 133.4 |
| Q61830 | ***Mrc1*** | 29.3 | 132.5 | 80.2 | 99.3 | 64.4 | 182.8 |
| P50516 | ***Atp6v1a*** | 29.6 | 97.7 | 136.1 | 72.2 | 111.5 | 177.8 |
| P08249 | ***Mdh2*** | 2837.4 | 942 | 793.9 | 818.5 | 897.1 | 1340.6 |
| P18572 | ***Bsg*** | 84.2 | 427.8 | 790.9 | 195.3 | 276.7 | 362.1 |
| P62984 | ***Uba52*** | 186.5 | 659.7 | 1115.5 | 1151.8 | 471 | 1088.5 |
| O35459 | ***Ech1*** | 289.6 | 125.5 | 217.6 | 187.8 | 144.5 | 192.8 |
| P14094 | ***Atp1b1*** | 42.4 | 878.5 | 1658.5 | 276 | 324.1 | 887.5 |
| O55143 | ***Atp2a2*** | 202.1 | 405.6 | 295.8 | 423.7 | 267 | 373 |
| Q8JZR0 | ***Acsl5*** | 527.6 | 1298.7 | 974.4 | 1506.1 | 997.5 | 915.3 |
| P41317 | ***Mbl2*** | 35.7 | 59.9 | 63.4 | 55.2 | 86.2 | 48.4 |
| Q9R092 | ***Hsd17b6*** | 322.4 | 998.5 | 1021.3 | 1595.7 | 840.2 | 673.6 |
| Q9CQV8 | ***Ywhab*** | 25 | 122.4 | 290.3 | 140.1 | 192.2 | 292.2 |
| O88833 | ***Cyp4a10*** | 20.5 | 207 | 123.5 | 312.6 | 162.8 | 60.6 |
| Q9CZX8 | ***Rps19*** | 47.6 | 125.6 | 286.6 | 183.4 | 255.1 | 173.5 |
| P62814 | ***Atp6v1b2*** | 29.7 | 113.5 | 145.2 | 74 | 124.7 | 185.2 |
| Q08857 | ***Cd36*** | 33.5 | 325.3 | 323.1 | 162.9 | 132.2 | 188.9 |
| P47740 | ***Aldh3a2*** | 525.6 | 1047.5 | 762.9 | 1315.7 | 787.8 | 718 |
| P01837 | | 2903.2 | 3186.8 | 2007.6 | 2198.3 | 3024.1 | 1894.3 |
| P10126 | ***Eef1a1*** | 782.7 | 2646.3 | 3296.2 | 3278.3 | 2555.9 | 2478.8 |
| Q8BU14 | ***Sec62*** | 58.9 | 89.9 | 67.8 | 102.4 | 67 | 74.9 |
| Q8VHE0 | ***Sec63*** | 145.7 | 229.4 | 97.8 | 213.1 | 159.8 | 181.1 |
| Q9EQH2 | ***Erap1*** | 41.3 | 71.8 | 87.6 | 104.1 | 51.3 | 85.4 |
| Q9QYG0 | ***Ndrg2*** | 21.5 | 244.1 | 918 | 106 | 141.1 | 391.6 |
| Q91X83 | ***Mat1a*** | 31 | 396.5 | 564.4 | 210.1 | 222.4 | 258.9 |
| P24638 | ***Acp2*** | 38.6 | 345.9 | 175.3 | 266.1 | 148.4 | 346.6 |
| Q9CPY7 | ***Lap3*** | 60.8 | 89.2 | 127.6 | 57 | 64.6 | 109 |
| P02088 | ***Hbb-b1*** | 2736.5 | 592.2 | 3425.1 | 354.9 | 1041.3 | 3148.6 |
| P01868 | ***Ighg1*** | 1619.3 | 2677.8 | 1757.3 | 1727.3 | 1920.2 | 1786.8 |
| P15105 | ***Glul*** | 146.1 | 526.4 | 1723.2 | 1143.7 | 1163.8 | 961.4 |
| P26041 | ***Msn*** | 345.3 | 580.2 | 437.6 | 366.8 | 596.7 | 581 |
| O08692 | ***Ngp*** | 103.4 | 50.5 | 40.6 | 88.8 | 101 | 49.4 |
| O35728 | ***Cyp4a14*** | 27.2 | 139.5 | 85.6 | 289.3 | 88.6 | 26.9 |
| Q9JJL3 | ***Slco1b2*** | 17.8 | 323.5 | 764.7 | 147.5 | 157.5 | 392.3 |
| P62270 | ***Rps18*** | 20.3 | 61.4 | 97.1 | 88.3 | 62.4 | 69.4 |
| P14246 | ***Slc2a2*** | 46.8 | 524.4 | 1573.9 | 267.5 | 228.7 | 798.8 |
| Q9QWL7 | ***Krt17*** | 196.9 | 336.1 | 668.5 | 423 | 239.9 | 280.1 |

| | | | | | | | |
|---|---|---|---|---|---|---|---|
| Q60634 | ***Flot2*** | 13.9 | 133.9 | 80.6 | 91.1 | 77.7 | 123.7 |
| Q9DCP2 | ***Slc38a3*** | 3.1 | 128.7 | 214.1 | 59.1 | 59 | 120.9 |
| O08601 | ***Mttp*** | 788.6 | 1529 | 2126.4 | 2312.4 | 1376.1 | 1515.7 |
| Q60931 | ***Vdac3*** | 273.5 | 211.7 | 172.2 | 167 | 148.1 | 268.1 |
| Q64481 | ***Cyp3a16*** | 150.4 | 505.2 | 389.7 | 506.5 | 343.5 | 375.5 |
| Q8VDN2 | ***Atp1a1*** | 93.6 | 1613 | 2798.8 | 695.5 | 660.8 | 1801.9 |
| Q8K2C9 | ***Hacd3*** | 39.5 | 113.9 | 75.7 | 118.9 | 74.8 | 75.4 |
| Q9CYH2 | ***Fam213a*** | 62.3 | 94.9 | 50.6 | 94.9 | 72.1 | 65.3 |
| O08914 | ***Faah*** | 49.6 | 88 | 58.2 | 127.7 | 50.1 | 59 |
| O35423 | ***Agxt*** | 61.3 | 234 | 65.8 | 123.7 | 71.3 | 78.1 |
| P40936 | ***Inmt*** | 36.6 | 112.9 | 507.9 | 114.9 | 144.9 | 181.4 |
| P46978 | ***Stt3a*** | 162.3 | 190.6 | 87.4 | 157.7 | 107.9 | 125.7 |
| Q8C165 | ***Pm20d1*** | 22.2 | 41.7 | 45.5 | 51 | 42.1 | 29.9 |
| Q60930 | ***Vdac2*** | 532.1 | 612.4 | 396.4 | 430.1 | 367.9 | 606.8 |
| P34927 | ***Asgr1*** | 23.7 | 395.4 | 401 | 138.5 | 128.6 | 348.9 |
| Q9JIL4 | ***Pdzk1*** | 14.9 | 147.8 | 452.2 | 91.6 | 133.1 | 204.5 |
| Q8BWQ1 | ***Ugt2a3*** | 339.6 | 782.9 | 446.9 | 896.5 | 478.3 | 463.4 |
| P29416 | ***Hexa*** | 54.6 | 199.8 | 106.3 | 139 | 140.7 | 139.2 |
| P01899 | ***H2-D1*** | 53.7 | 128.2 | 175.8 | 127.1 | 111.8 | 173.3 |
| O08807 | ***Prdx4*** | 217.9 | 504.5 | 1210.5 | 843.4 | 832 | 659.2 |
| O35604 | ***Npc1*** | 259.9 | 462.8 | 338.6 | 473.5 | 242.6 | 474 |
| P56657 | ***Cyp2c40*** | 202.1 | 919.8 | 427.5 | 723.9 | 258.2 | 234.5 |
| Q8R0Y6 | ***Aldh1l1*** | 94.8 | 1096.5 | 2694.2 | 714.2 | 831.5 | 1364.9 |
| P20060 | ***Hexb*** | 69.7 | 297.8 | 178.9 | 226.2 | 149.4 | 261.3 |
| P11609 | ***Cd1d1*** | 20.1 | 234.3 | 328.8 | 162.9 | 108.5 | 197.2 |
| P11679 | ***Krt8*** | 203.6 | 243.2 | 763.1 | 348.2 | 74.4 | 520.9 |
| P50285 | ***Fmo1*** | 449.3 | 1401.6 | 1141.9 | 1734.9 | 926.8 | 812 |
| P62880 | ***Gnb2*** | 47.3 | 268.6 | 436.2 | 228.2 | 244.7 | 432.8 |
| P00329 | ***Adh1*** | 46.7 | 842.6 | 1664.9 | 2344.2 | 523.1 | 579.7 |
| Q922Q8 | ***Lrrc59*** | 193.4 | 238.8 | 120 | 301.9 | 218.3 | 105.6 |
| Q9QXD6 | ***Fbp1*** | 14.5 | 78.2 | 213.3 | 67 | 59.9 | 112.3 |
| UPSP:K22E_HUMAN | | 1315.6 | 2909.8 | 2982.3 | 2266.9 | 2655.9 | 1605.6 |
| Q01768 | ***Nme2*** | 117.2 | 165.4 | 521.4 | 142 | 214.5 | 300.3 |
| P62259 | ***Ywhae*** | 10.9 | 56.4 | 137.1 | 60.9 | 79.2 | 106.3 |
| P63017 | ***Hspa8*** | 150.5 | 347.5 | 983.2 | 349.2 | 589.3 | 849.9 |
| P63242 | ***Eif5a*** | 85 | 94.5 | 227.4 | 131.8 | 156.1 | 168.2 |
| P01901 | ***H2-K1*** | 55.1 | 230.9 | 399.2 | 205.7 | 280.8 | 307 |
| Q9JHI5 | ***Ivd*** | 430.8 | 317.8 | 208.9 | 253.3 | 281.6 | 357.2 |
| Q9DCN2 | ***Cyb5r3*** | 755.9 | 1210.6 | 680.1 | 1012.4 | 759.9 | 700.7 |
| Q61171 | ***Prdx2*** | 57.1 | 95 | 247 | 85.3 | 145.3 | 232.9 |
| Q91Y97 | ***Aldob*** | 134.7 | 726.5 | 3240.5 | 658.5 | 1275.7 | 1303.6 |
| P05213 | ***Tuba1b*** | 49.6 | 728.5 | 805.2 | 394.7 | 514.1 | 922 |
| P17742 | ***Ppia*** | 48.4 | 123.5 | 826.2 | 232.6 | 340.5 | 663.2 |
| Q91XD4 | ***Ftcd*** | 56.7 | 162.9 | 319.6 | 120.6 | 142.4 | 211.8 |
| Q9DBG6 | ***Rpn2*** | 396.4 | 684 | 279.7 | 670.3 | 396.8 | 410.4 |
| Q9QZW0 | ***Atp11c*** | 55.2 | 572.2 | 909 | 302.8 | 236 | 767.4 |
| P14824 | ***Anxa6*** | 46.4 | 291.4 | 1046.5 | 191.8 | 256.4 | 683.5 |

| | | | | | | | |
|---|---|---|---|---|---|---|---|
| Q01853 | ***Vcp*** | 15.6 | 76.4 | 185.8 | 68.6 | 104.5 | 145.3 |
| Q8QZS1 | ***Hibch*** | 79.4 | 111.3 | 73.8 | 67.8 | 90.7 | 119.5 |
| P16858 | ***Gapdh*** | 42.5 | 539 | 1176.5 | 352.6 | 820.7 | 842.8 |
| P12790 | ***Cyp2b9*** | 47.4 | 227.8 | 117.7 | 364.3 | 75.7 | 34.6 |
| P11499 | ***Hsp90ab1*** | 42.6 | 287.3 | 765.3 | 206.4 | 332.2 | 412.3 |
| P12791 | ***Cyp2b10*** | 87 | 505.6 | 275.6 | 669.8 | 249 | 84.2 |
| Q99LB2 | ***Dhrs4*** | 590.6 | 233.9 | 211.5 | 350.1 | 359.4 | 247.3 |
| P16331 | ***Pah*** | 0 | 76 | 145.1 | 77.4 | 78.5 | 90.8 |
| O08709 | ***Prdx6*** | 23.2 | 96.7 | 427.2 | 103.1 | 161.6 | 207.8 |
| P31786 | ***Dbi*** | 154.3 | 361.5 | 1420.1 | 299.6 | 751.8 | 652.1 |
| P50172 | ***Hsd11b1*** | 1090.4 | 2330.1 | 1649.4 | 2473.4 | 1696.9 | 1183.6 |
| P35700 | ***Prdx1*** | 242 | 904.8 | 2410.6 | 1246.3 | 1505.2 | 1161.6 |
| Q61176 | ***Arg1*** | 117.7 | 487.1 | 2139 | 422.3 | 829.6 | 1135.6 |
| Q78JT3 | ***Haao*** | 5.3 | 85.3 | 356.9 | 69.7 | 121.8 | 199.6 |
| P36552 | ***Cpox*** | 148 | 143.1 | 146 | 166.2 | 62.3 | 189.8 |
| P16015 | ***Ca3*** | 15.2 | 173.7 | 1405.1 | 298.8 | 575 | 948 |
| O08573 | ***Lgals9*** | 62.8 | 252.4 | 769.4 | 323.9 | 179.3 | 475.1 |
| O35490 | ***Bhmt*** | 254.6 | 857.9 | 2557.8 | 813.6 | 1094.5 | 1434.9 |
| UPSP:K1CX_HUMAN | | 31.2 | 68.7 | 138.2 | 121.1 | 54.7 | 44.3 |
| Q91YQ5 | ***Rpn1*** | 774.5 | 1153.4 | 478.5 | 984.2 | 813.5 | 704.8 |
| Q91YI0 | ***Asl*** | 4.2 | 160.5 | 306.8 | 125 | 176.6 | 209.1 |
| Q63886 | ***Ugt1a1*** | 756.7 | 1698.6 | 798.8 | 1578 | 867.4 | 651.7 |
| P24549 | ***Aldh1a1*** | 132.2 | 1076.4 | 3145 | 906 | 909.8 | 2196.6 |
| Q60932 | ***Vdac1*** | 1155.3 | 745.8 | 678.2 | 724.5 | 809 | 995.3 |
| P61620 | ***Sec61a1*** | 43.8 | 64.6 | 23.2 | 54.2 | 45 | 35.9 |
| Q61009 | ***Scarb1*** | 7.6 | 100.8 | 167.7 | 69.5 | 58.5 | 134.4 |
| P06151 | ***Ldha*** | 10.8 | 157.7 | 405 | 153.9 | 198.1 | 251.1 |
| Q64435 | ***Ugt1a6*** | 773.5 | 1314 | 625.4 | 1158.9 | 904.4 | 587.4 |
| Q9DBF1 | ***Aldh7a1*** | 257.4 | 388.1 | 311.2 | 311.7 | 260 | 398 |
| P49429 | ***Hpd*** | 75.6 | 260.1 | 1117.3 | 357.6 | 458.6 | 561.9 |
| P12710 | ***Fabp1*** | 99.2 | 359.6 | 3811.2 | 668.2 | 1480.9 | 1835.8 |
| Q01279 | ***Egfr*** | 10.4 | 134.5 | 225.5 | 65.9 | 75.4 | 211.6 |
| P17182 | ***Eno1*** | 102.3 | 519.6 | 1929.7 | 468.6 | 790.4 | 1163 |
| P52760 | ***Hrsp12*** | 14.9 | 44.3 | 425.4 | 99.2 | 156.9 | 201.2 |
| P50247 | ***Ahcy*** | 67.1 | 697.3 | 2626.9 | 755.5 | 1206 | 1576.8 |
| Q9DBH5 | ***Lman2*** | 51.1 | 197.3 | 124.1 | 137 | 120.2 | 107.6 |
| Q7TMM9 | ***Tubb2a*** | 18.5 | 263.1 | 307.2 | 135.5 | 148 | 323.3 |
| P52430 | ***Pon1*** | 310.4 | 746.5 | 579.7 | 652.1 | 465.4 | 496.4 |
| Q9QY76 | ***Vapb*** | 302 | 391.4 | 229.7 | 360 | 319.6 | 234.1 |
| P09411 | ***Pgk1*** | 18.2 | 92.6 | 320.5 | 72 | 142.1 | 201.7 |
| Q64464 | ***Cyp3a13*** | 257.8 | 645.8 | 411.8 | 581.4 | 313.6 | 400.6 |
| O09173 | ***Hgd*** | 8.6 | 82.4 | 201 | 86.2 | 97.7 | 116.1 |
| Q9DCM2 | ***Gstk1*** | 224.6 | 116.8 | 99 | 160.3 | 168.5 | 105.7 |
| Q9QXF8 | ***Gnmt*** | 16.1 | 148.4 | 440.2 | 128.6 | 195.1 | 263.9 |
| Q9QXZ6 | ***Slco1a1*** | 18.4 | 283.8 | 687.7 | 139.1 | 182.6 | 638.7 |
| P10649 | ***Gstm1*** | 32.2 | 171.1 | 903.1 | 235.5 | 389.8 | 449.3 |
| P14152 | ***Mdh1*** | 21.3 | 89.9 | 414.6 | 119.5 | 179.7 | 240.1 |

| | | | | | | | |
|---|---|---|---|---|---|---|---|
| Q64374 | ***Rgn*** | 43.8 | 187.6 | 1519.8 | 439.5 | 654.9 | 705.5 |
| P17751 | ***Tpi1*** | 10.4 | 86.7 | 440.9 | 83.5 | 201.6 | 239.7 |
| Q64442 | ***Sord*** | 24.5 | 274.2 | 1099.5 | 303.5 | 508.1 | 617.2 |
| Q8VEK0 | ***Tmem30a*** | 25.4 | 90.5 | 145 | 60.3 | 50.4 | 146.2 |
| P70441 | ***Slc9a3r1*** | 19.4 | 105.9 | 288.5 | 64.7 | 144.1 | 199.3 |
| P30115 | ***Gsta3*** | 26.7 | 147.2 | 874.8 | 206.2 | 397 | 434.2 |
| P70694 | ***Akr1c6*** | 54.6 | 318.5 | 1355.9 | 356.3 | 598.6 | 783.9 |
| Q99PL5 | ***Rrbp1*** | 523.2 | 304.4 | 304 | 410.1 | 478.8 | 244.5 |

**Table S5: Immunoisolated proteome of basal and starved Brain AVs**

Only proteins that were detected with more than 3 unique peptides are listed.

The numerical values are peptide intensity normalized to total intensity of each sample.

| uniprot | gene symbol | Intensity of Basal1 | Intensity of Basal2 | Intensity of Basal3 | Intensity of Starved1 | Intensity of Starved2 | Intensity of Starved3 |
|---|---|---|---|---|---|---|---|
| Q922B1 | ***Macrod1*** | 188.8 | 185.4 | 202.7 | 195.8 | 201.5 | 211.4 |
| Q99L13 | ***Hibadh*** | 479.8 | 483.3 | 350.9 | 428.9 | 445.7 | 440 |
| Q9QYR9 | ***Acot2*** | 179.5 | 100.9 | 103.8 | 175.6 | 163.4 | 176.3 |
| Q99LC5 | ***Etfa*** | 1913.1 | 2364 | 2050.1 | 1977.8 | 1976.4 | 1910.1 |
| Q8R4N0 | ***Clybl*** | 357.2 | 373.6 | 354.4 | 373.8 | 396.4 | 386.6 |
| Q99KR7 | ***Ppif*** | 296.2 | 193.6 | 399.5 | 472.2 | 442.4 | 456.5 |
| P08249 | ***Mdh2*** | 9506.5 | 7292.5 | 11790.1 | 12055.6 | 11614.7 | 12353.4 |
| P29758 | ***Oat*** | 247.2 | 280.1 | 313 | 539.4 | 582.4 | 538 |
| Q9JHI5 | ***Ivd*** | 682.8 | 810 | 691.3 | 972.6 | 1116 | 1131.3 |
| Q8QZS1 | ***Hibch*** | 477.6 | 327.8 | 554.8 | 575.6 | 599 | 647.6 |
| P08228 | ***Sod1*** | 397.2 | 320.5 | 883.8 | 1020.3 | 1314.6 | 644.5 |
| P09671 | ***Sod2*** | 2229 | 1887.6 | 3006.4 | 3518.4 | 3669.5 | 3690 |
| Q9JKC6 | ***Cend1*** | 4325.3 | 3478.2 | 3699.4 | 2254.1 | 2018.6 | 2346.7 |
| P54071 | ***Idh2*** | 762 | 1057.7 | 672.7 | 924 | 991.9 | 1160.6 |
| Q9WTP7 | ***Ak3*** | 1796.9 | 1769.2 | 2073.3 | 1523.5 | 1626.3 | 1771.8 |
| Q9D0K2 | ***Oxct1*** | 2007.8 | 1759.4 | 1982.5 | 2696.3 | 2516.5 | 2798.6 |
| P47738 | ***Aldh2*** | 2238.9 | 2087.3 | 1123.3 | 2125.8 | 2118.4 | 2292.8 |
| Q8CC88 | ***Vwa8*** | 36.6 | 54.5 | 21.5 | 28.4 | 28.2 | 38.4 |
| Q9D5T0 | ***Atad1*** | 117.5 | 155.8 | 68 | 136.1 | 145.6 | 161.1 |
| Q9JLZ3 | ***Auh*** | 1782.6 | 2818.6 | 2943.4 | 1680.6 | 1744 | 2191.3 |
| P61922 | ***Abat*** | 1238.1 | 1499.2 | 949.4 | 1593.9 | 1147.4 | 1514.1 |
| Q9CZP5 | ***Bcs1l*** | 183.7 | 200.1 | 153 | 119.9 | 104 | 147.7 |
| Q9CQV1 | ***Pam16*** | 176 | 195.5 | 254.7 | 115.8 | 131.6 | 137.1 |
| Q61425 | ***Hadh*** | 575.2 | 614 | 752.9 | 592.3 | 733.5 | 705.9 |
| P51174 | ***Acadl*** | 731.6 | 966.1 | 501.7 | 810.9 | 955.5 | 985.1 |
| Q8BK72 | ***Mrps27*** | 265.9 | 237.2 | 132.2 | 153.5 | 144.3 | 191.1 |
| Q9Z0J0 | ***Npc2*** | 55.9 | 29.9 | 91.7 | 141.5 | 83.5 | 72.7 |
| Q8C5H8 | ***Nadk2*** | 127.2 | 119.7 | 115 | 130.7 | 136.4 | 158.8 |
| Q9CR21 | ***Ndufab1*** | 4531.1 | 3094 | 3230.5 | 2182.3 | 1871.1 | 2327.9 |
| Q9CZS1 | ***Aldh1b1*** | 200.7 | 184.6 | 172.6 | 189.3 | 145.2 | 202.9 |
| Q64433 | ***Hspe1*** | 4660.2 | 2440.2 | 4859.5 | 3577.4 | 4460.1 | 3996.1 |
| Q8BMF3 | ***Me3*** | 512.6 | 475.7 | 417.1 | 613.7 | 394.7 | 723 |
| Q06185 | ***Atp5i*** | 2365.9 | 2179.2 | 3440.4 | 1274.8 | 1177.3 | 1614.4 |
| Q8C163 | ***Exog*** | 542 | 556.1 | 690.3 | 427.2 | 364.8 | 492.7 |
| Q9DC61 | ***Pmpca*** | 169.6 | 212.9 | 103.3 | 132.6 | 136.9 | 209.5 |
| Q8R164 | ***Bphl*** | 199.9 | 228.6 | 366.6 | 495.5 | 386.2 | 583.5 |
| Q99M87 | ***Dnaja3*** | 514.5 | 578.6 | 450.6 | 277 | 328.5 | 402.7 |
| Q9CPV4 | ***Glod4*** | 295.5 | 258.2 | 398.2 | 565.1 | 571 | 464 |
| Q8VCW8 | ***Acsf2*** | 335.2 | 437 | 203.1 | 358 | 373.9 | 442.6 |
| O35143 | ***Atpif1*** | 2698.6 | 2261.7 | 3693.7 | 1347.4 | 1519.8 | 1816.1 |
| P07724 | ***Alb*** | 582.7 | 336 | 545.8 | 722.8 | 1153.9 | 784.6 |

| | | | | | | | |
|---|---|---|---|---|---|---|---|
| Q9D172 | ***D10Jhu81e*** | 2183.2 | 1771.5 | 2381.1 | 2811.9 | 2778.9 | 3377.4 |
| P26443 | ***Glud1*** | 6587.5 | 10559.6 | 10247.6 | 7419.4 | 7498.6 | 8947.5 |
| Q9CQA3 | ***Sdhb*** | 4370.5 | 5294 | 5675.9 | 3258.1 | 3050.2 | 4082.6 |
| Q9DCN2 | ***Cyb5r3*** | 275.2 | 254 | 168.1 | 203.3 | 209.2 | 298.2 |
| Q9EQ20 | ***Aldh6a1*** | 1906.6 | 2087.8 | 1216.5 | 1656.8 | 1847.7 | 1980.9 |
| P46638 | ***Rab11b*** | 261 | 200.1 | 221.4 | 291.4 | 312.8 | 253 |
| Q9D0S9 | ***Hint2*** | 134 | 95.7 | 164.2 | 199.4 | 222.1 | 172.4 |
| Q80Y14 | ***Glrx5*** | 403 | 371.4 | 461.4 | 302 | 311.2 | 342.7 |
| O35387 | ***Hax1*** | 67.3 | 46.3 | 65.7 | 42.9 | 44.1 | 53.3 |
| Q8VEM8 | ***Slc25a3*** | 4440.2 | 3705.5 | 3085.1 | 2649.5 | 1645.6 | 3295.8 |
| P42125 | ***Eci1*** | 969.8 | 1035.9 | 839.7 | 732.3 | 823.7 | 892.7 |
| Q9DBL1 | ***Acadsb*** | 524.1 | 592.4 | 347.7 | 579.5 | 475.3 | 603.6 |
| P10107 | ***Anxa1*** | 620.6 | 377.9 | 574.3 | 872.7 | 494 | 370.8 |
| Q61102 | ***Abcb7*** | 496 | 534.9 | 314.5 | 401.9 | 339.4 | 517.2 |
| O35459 | ***Ech1*** | 48.5 | 47.3 | 52.9 | 61.6 | 86.7 | 92.1 |
| Q9DCM0 | ***Ethe1*** | 346 | 315.9 | 291.1 | 319.2 | 307.2 | 348.6 |
| P56213 | ***Gfer*** | 135.4 | 123.3 | 196.6 | 89.9 | 98.9 | 94.8 |
| Q99L04 | ***Dhrs1*** | 268.2 | 349.4 | 309.2 | 253.7 | 219.4 | 285.9 |
| P62984 | ***Uba52*** | 499 | 261.9 | 462.7 | 873.6 | 843 | 438.9 |
| Q9Z2I0 | ***Letm1*** | 1511.2 | 1394.5 | 1394.3 | 1309 | 738.5 | 1475.8 |
| Q8BH59 | ***Slc25a12*** | 4511.9 | 4625.2 | 2441.1 | 3630.5 | 2351.8 | 4356.2 |
| Q8CGK3 | ***Lonp1*** | 824.5 | 1178 | 529.5 | 787.6 | 622.5 | 944.9 |
| P11725 | ***Otc*** | 21.4 | 25.1 | 36.7 | 40.6 | 19 | 24.3 |
| P08226 | ***Apoe*** | 326.5 | 165.2 | 349.1 | 339.5 | 423.6 | 270.9 |
| Q9JLJ2 | ***Aldh9a1*** | 140.8 | 150.3 | 97.5 | 165 | 182.9 | 189.1 |
| Q922H2 | ***Pdk3*** | 197.5 | 231.3 | 125.8 | 176.1 | 148.2 | 244.1 |
| Q8QZT1 | ***Acat1*** | 4558.7 | 4993.9 | 4835.9 | 3748.6 | 3479 | 4081.4 |
| Q91VN4 | ***Chchd6*** | 1444.2 | 1518.1 | 1245.3 | 1118 | 1021.5 | 1341.2 |
| P45952 | ***Acadm*** | 571.6 | 563.6 | 620 | 545 | 654.6 | 568.3 |
| P56135 | ***Atp5j2*** | 513 | 330.2 | 416.8 | 111.6 | 143.4 | 165.4 |
| Q9ET22 | ***Dpp7*** | 33.7 | 0 | 44.9 | 139.9 | 84.6 | 45.6 |
| P70663 | ***Sparcl1*** | 143.6 | 33.5 | 123.2 | 221.1 | 203.9 | 96 |
| Q8BKZ9 | ***Pdhx*** | 2043.6 | 2075 | 1972.4 | 1518.9 | 1251.1 | 1893.9 |
| Q9CQ92 | ***Fis1*** | 309.1 | 407.9 | 443.4 | 429.4 | 324.2 | 334.7 |
| Q9DCX2 | ***Atp5h*** | 8235.5 | 7405.5 | 9533.6 | 4863.1 | 4391.4 | 5438.2 |
| Q9Z2I8 | ***Suclg2*** | 173.8 | 342.5 | 181 | 237.4 | 234.5 | 326.4 |
| P06797 | ***Ctsl*** | 612.4 | 97.4 | 537.3 | 558.4 | 272 | 243.5 |
| Q03265 | ***Atp5a1*** | 23580.4 | 26206.9 | 30073.5 | 18832.4 | 14687.8 | 23386.6 |
| P48962 | ***Slc25a4*** | 4002.9 | 3892.3 | 3066.2 | 3181.9 | 2352.2 | 3709.8 |
| Q3V3R1 | ***Mthfd1l*** | 239.6 | 256.4 | 131 | 200.6 | 139.8 | 232.5 |
| Q9CQQ7 | ***Atp5f1*** | 5954.4 | 6157.3 | 5355 | 4486.2 | 3054 | 5511.3 |
| Q61207 | ***Psap*** | 5807.1 | 963.4 | 4280.1 | 13956.2 | 5877.9 | 4007.9 |
| P07356 | ***Anxa2*** | 493.2 | 343.3 | 498.1 | 812.1 | 418.5 | 337.6 |
| Q9DBF1 | ***Aldh7a1*** | 271.4 | 312.8 | 213.7 | 375 | 348.9 | 373.4 |
| P19783 | ***Cox4i1*** | 1939.7 | 1706 | 1451.9 | 1032.8 | 947.5 | 1337.1 |
| Q9R0X4 | ***Acot9*** | 245.7 | 291.8 | 188.2 | 239.5 | 192.8 | 313.6 |
| P85094 | ***Isoc2a*** | 158 | 132 | 148 | 140.9 | 185.3 | 192.3 |

| | | | | | | | |
|---|---|---|---|---|---|---|---|
| Q8JZQ2 | ***Afg3l2*** | 1129.9 | 1277.8 | 1058.5 | 991.7 | 885.2 | 1359.5 |
| Q80U63 | ***Mfn2*** | 181.1 | 166.8 | 89.2 | 153.2 | 151.1 | 183 |
| Q07417 | ***Acads*** | 98.1 | 93.7 | 103.3 | 95.8 | 109.7 | 102.2 |
| Q791T5 | ***Mtch1*** | 180.2 | 164 | 155.3 | 131.3 | 130.3 | 176.9 |
| Q9CQH3 | ***Ndufb5*** | 3652.6 | 4093.4 | 3357.4 | 2563.8 | 2160.9 | 3544.5 |
| P10605 | ***Ctsb*** | 988.7 | 184.1 | 651.9 | 1553 | 740.9 | 484.1 |
| O35114 | ***Scarb2*** | 879.6 | 200.7 | 318.7 | 1021.7 | 482 | 457 |
| O89023 | ***Tpp1*** | 1304.4 | 344.1 | 1264.2 | 2092.2 | 896.6 | 672.7 |
| P56480 | ***Atp5b*** | 13819.9 | 11514.6 | 17490.7 | 9419.1 | 7196.6 | 10066 |
| P70699 | ***Gaa*** | 647 | 73.4 | 178.9 | 542.9 | 269.7 | 153.1 |
| Q5IRJ6 | ***Slc30a9*** | 146.9 | 141.6 | 120.2 | 129.7 | 92.1 | 142.7 |
| P84086 | ***Cplx2*** | 281.5 | 293.1 | 400.2 | 338.7 | 467.2 | 341.4 |
| P63038 | ***Hspd1*** | 13863.2 | 13692.1 | 15301 | 11099.9 | 9936.5 | 11465.6 |
| Q04519 | ***Smpd1*** | 212.5 | 49 | 157.9 | 391.4 | 142.1 | 142.9 |
| UPSP:K1CI_HUMAN | | 1522.4 | 1361.5 | 6645.5 | 3521.6 | 1286.3 | 3193.7 |
| Q8CHT0 | ***Aldh4a1*** | 331.7 | 373.2 | 280.4 | 297.9 | 322.7 | 257.5 |
| P50429 | ***Arsb*** | 173.4 | 48.1 | 116.4 | 452.6 | 207.9 | 173 |
| P16675 | ***Ctsa*** | 239.7 | 57 | 112.4 | 259.1 | 144.1 | 96.6 |
| Q9CWZ7 | ***Napg*** | 941.8 | 899.8 | 1140.2 | 2389.9 | 2724.1 | 1203.8 |
| Q9R013 | ***Ctsf*** | 243.9 | 59.1 | 221.4 | 448.5 | 160.9 | 133.2 |
| Q9D2G2 | ***Dlst*** | 3106.8 | 2949.4 | 2764.2 | 1947.8 | 2070.7 | 2594 |
| Q9D6M3 | ***Slc25a22*** | 2237.9 | 2534.8 | 2080 | 2000.3 | 1339.7 | 2807.3 |
| Q9D6J5 | ***Ndufb8*** | 952.1 | 842.7 | 860.1 | 684 | 447 | 790.6 |
| Q3UMR5 | ***Mcu*** | 424.9 | 497.6 | 306.1 | 430.9 | 249 | 520.6 |
| Q64332 | ***Syn2*** | 143.4 | 212.7 | 170.7 | 819.9 | 598.2 | 316.8 |
| P18242 | ***Ctsd*** | 8688 | 1592.3 | 6645 | 11784.1 | 4478.7 | 4152.5 |
| P67778 | ***Phb*** | 3100.3 | 3440.7 | 4066 | 2448.1 | 2061.8 | 3125.4 |
| Q9DB05 | ***Napa*** | 329.7 | 208.7 | 241.6 | 652.4 | 679 | 313.2 |
| P47802 | ***Mtx1*** | 215.2 | 214.8 | 197.6 | 190.7 | 105 | 246.4 |
| Q9DCB8 | ***Isca2*** | 165.1 | 131.7 | 177.5 | 144.4 | 139.3 | 165.5 |
| P52196 | ***Tst*** | 306.8 | 502.3 | 248.8 | 302.3 | 300.4 | 367.5 |
| P20060 | ***Hexb*** | 823.5 | 243.5 | 418 | 979.8 | 395.4 | 424 |
| Q9DBT9 | ***Dmgdh*** | 57.1 | 51.1 | 57.2 | 54.3 | 50.1 | 82.4 |
| Q8BX10 | ***Pgam5*** | 328.2 | 342.9 | 268.1 | 235.2 | 219.6 | 369.1 |
| Q9CQJ8 | ***Ndufb9*** | 626 | 753.3 | 736.8 | 433.9 | 419.2 | 530.9 |
| Q9WV54 | ***Asah1*** | 874.3 | 253.6 | 1149.3 | 1988.6 | 595.5 | 715.2 |
| P62806 | ***Hist1h4a; Hist1h4b; Hist1h4c; Hist1h4d; Hist1h4f; Hist1h4h; Hist1h4i; Hist1h4j; Hist1h4k; Hist1h4m; Hist2h4a; Hist4h4*** | 65.1 | 200.7 | 358.9 | 242.3 | 46.4 | 394.2 |
| Q00493 | ***Cpe*** | 218.4 | 86.5 | 175.4 | 473.1 | 495.9 | 218.6 |
| Q60648 | ***Gm2a*** | 556.9 | 148.1 | 498.2 | 551 | 330.4 | 261.4 |

| | | | | | | | |
|---|---|---|---|---|---|---|---|
| O88531 | ***Ppt1*** | 1955.5 | 416.1 | 1319.9 | 2304.4 | 851.2 | 905.4 |
| Q9DCZ4 | ***Apoo*** | 696.2 | 811.3 | 960.8 | 621.2 | 572 | 930 |
| Q9CZW5 | ***Tomm70*** | 2763.8 | 3265.5 | 2453.6 | 2779.7 | 2045.4 | 3532.3 |
| P41216 | ***Acsl1*** | 135.3 | 154 | 79.5 | 120.6 | 107.8 | 124.1 |
| P28663 | ***Napb*** | 236 | 171.9 | 181.6 | 459.5 | 517.7 | 250.6 |
| Q9Z2Z6 | ***Slc25a20*** | 84.8 | 78.9 | 92 | 61.6 | 60.8 | 96.6 |
| Q9CZ13 | ***Uqcrc1*** | 5193.6 | 4592.4 | 3310.5 | 3219.4 | 2589.6 | 4008.5 |
| P16332 | ***Mut*** | 193.9 | 166.6 | 113.8 | 149.8 | 123.5 | 195.5 |
| UPSP:K22E_HUMAN | | 1543.9 | 499.7 | 2441.1 | 2564 | 1220.1 | 1543.6 |
| Q9D051 | ***Pdhb*** | 5336.3 | 5585.6 | 5228.3 | 4222.1 | 3050 | 4929.9 |
| Q9EQI8 | ***Mrpl46*** | 81.1 | 111 | 75.3 | 88.2 | 69.8 | 94.5 |
| O35405 | ***Pld3*** | 927 | 236.7 | 516.7 | 1122.3 | 424.7 | 441.8 |
| P26040 | ***Ezr*** | 993.3 | 827.4 | 1205 | 646.8 | 675.1 | 474.6 |
| Q99KI0 | ***Aco2*** | 7928.9 | 9801 | 6436.4 | 11046.8 | 8560.5 | 11282.6 |
| Q99M71 | ***Epdr1*** | 1874.3 | 315.4 | 1172.2 | 2336.2 | 880.1 | 836 |
| Q9CQ75 | ***Ndufa2*** | 1539.3 | 1673.6 | 2023.7 | 1006.7 | 946.9 | 1299.4 |
| P68368 | ***Tuba4a*** | 452.3 | 924.4 | 250.6 | 1329.1 | 1823.4 | 836.3 |
| Q99LP6 | ***Grpel1*** | 463.3 | 472.2 | 514.7 | 532.3 | 466.1 | 588.5 |
| Q8K2B3 | ***Sdha*** | 4993.7 | 5243.4 | 2860.3 | 3716.8 | 3283.1 | 4011.4 |
| Q8BIJ6 | ***Iars2*** | 403.2 | 446.7 | 230.5 | 342.4 | 251.4 | 409.7 |
| O35129 | ***Phb2*** | 1623 | 1952.3 | 2098.7 | 1440.2 | 1250.7 | 2013.3 |
| Q9CZR8 | ***Tsfm*** | 166.6 | 279.3 | 236.7 | 180.8 | 177.4 | 230 |
| D3Z7P3 | ***Gls*** | 1922 | 2036.1 | 1230.6 | 1638.7 | 1283.1 | 2196.4 |
| UPSP:K1CJ_HUMAN | | 1536.7 | 471.1 | 2588.2 | 2435.6 | 1042.2 | 1409.7 |
| O35857 | ***Timm44*** | 193.3 | 240.5 | 180.2 | 174.6 | 146.5 | 229 |
| Q99JR1 | ***Sfxn1*** | 282.9 | 313.9 | 252 | 245.2 | 200.6 | 287.8 |
| P15105 | ***Glul*** | 827 | 1230.1 | 721.8 | 1839.7 | 2715.3 | 1708.4 |
| Q9CPQ1 | ***Cox6c*** | 2347.2 | 2488.4 | 2658.5 | 1829.3 | 1748.6 | 2138.9 |
| O88741 | ***Gdap1*** | 1045.3 | 1049 | 770.5 | 828.5 | 534.3 | 1086.1 |
| Q8BFR5 | ***Tufm*** | 5537.3 | 8015.2 | 4863.3 | 6398.6 | 4948.8 | 6824.2 |
| Q9DB20 | ***Atp5o*** | 5337.2 | 6347.2 | 8835.3 | 4392.2 | 3699.2 | 5113.1 |
| Q9DC70 | ***Ndufs7*** | 858.2 | 1013.4 | 650 | 593.4 | 485.6 | 852.2 |
| P29416 | ***Hexa*** | 299.6 | 81.2 | 141.7 | 354.7 | 134.2 | 136.1 |
| Q60932 | ***Vdac1*** | 9871.6 | 10714.5 | 8406.1 | 9773.8 | 7881.7 | 9942.9 |
| Q9WVA2 | ***Timm8a1*** | 201.4 | 258.9 | 279.7 | 140.2 | 120.1 | 165.4 |
| Q3UV17 | ***Krt76*** | 340.7 | 119.3 | 873.8 | 472.8 | 199.1 | 324.2 |
| Q9CQC7 | ***Ndufb4*** | 1119.4 | 1199.6 | 1401.7 | 897.9 | 663.4 | 980.9 |
| O88384 | ***Vti1b*** | 84.7 | 28.5 | 70.4 | 151.3 | 75.7 | 59.5 |
| Q9WUU7 | ***Ctsz*** | 16.4 | 0 | 21.6 | 26 | 15 | 3.1 |
| Q91WD5 | ***Ndufs2*** | 3476 | 2823.5 | 2244.5 | 2698.6 | 1408 | 3182.4 |
| Q9D1K2 | ***Atp6v1f*** | 204.4 | 146.2 | 237.5 | 257.2 | 253.6 | 160.6 |
| P38647 | ***Hspa9*** | 6041.9 | 5746 | 4974.6 | 4983.2 | 5301.1 | 5384.8 |
| P97807 | ***Fh*** | 2505.4 | 3338.6 | 2081.2 | 2523.4 | 2925.4 | 2887.5 |
| Q9D1L0 | ***Chchd2*** | 120.1 | 125.9 | 125.3 | 117.3 | 133.8 | 118.3 |
| Q8BMF4 | ***Dlat*** | 3475.3 | 3912.4 | 2877.7 | 2927.1 | 2111.8 | 3298 |
| UPSP:K2C1_HUMAN | | 3114.7 | 1641.3 | 9700.2 | 4944.5 | 2123.4 | 4276.9 |

| | | | | | | | |
|---|---|---|---|---|---|---|---|
| Q8BWT1 | ***Acaa2*** | 701.5 | 1213.7 | 703.2 | 898.4 | 780.4 | 874.2 |
| Q8CAQ8 | ***Immt*** | 4235.9 | 7192.5 | 3279.6 | 5892.6 | 4818.6 | 7647.8 |
| Q9CQ62 | ***Decr1*** | 403.6 | 726.5 | 464.6 | 439.1 | 478.3 | 585.1 |
| Q920A5 | ***Scpep1*** | 192.4 | 38.8 | 72.8 | 164.3 | 79.6 | 40.6 |
| Q9CYR0 | ***Ssbp1*** | 433.5 | 471.9 | 448.8 | 415.5 | 376.8 | 528.9 |
| Q8K0D5 | ***Gfm1*** | 132.4 | 126.6 | 62.1 | 118.1 | 111.4 | 125.6 |
| UPSP:K1CN_HUMAN | | 1012.5 | 372.1 | 2339.9 | 1466.5 | 582 | 867.6 |
| Q9CQR4 | ***Acot13*** | 993.3 | 1091.3 | 1318.4 | 1088.7 | 839 | 1193.4 |
| A2ASZ8 | ***Slc25a25*** | 260.6 | 314.3 | 213 | 233.8 | 147.6 | 280.4 |
| Q62425 | ***Ndufa4*** | 4484.1 | 4464.1 | 5839.5 | 2808.5 | 2286.1 | 3001.6 |
| Q9CR62 | ***Slc25a11*** | 936.5 | 1115.8 | 602.1 | 847.1 | 654.9 | 1223.8 |
| P50516 | ***Atp6v1a*** | 1214.9 | 1018.8 | 917.3 | 1836.6 | 1853.8 | 1445.6 |
| P62075 | ***Timm13*** | 67.1 | 87.6 | 80.5 | 55.2 | 51.9 | 77.2 |
| Q8VEA4 | ***Chchd4*** | 108.1 | 91.6 | 135.4 | 122.4 | 98.6 | 136 |
| O54734 | ***Ddost*** | 103.7 | 57.5 | 38.5 | 140.2 | 209.4 | 115.8 |
| O08756 | ***Hsd17b10*** | 288.8 | 477.1 | 527 | 278.8 | 310.9 | 355.9 |
| Q791V5 | ***Mtch2*** | 472.4 | 652.3 | 314.6 | 557.5 | 412.2 | 711.2 |
| Q9CQZ5 | ***Ndufa6*** | 1859.2 | 2176.9 | 2654.1 | 1473 | 1099.8 | 1772.2 |
| Q9DC69 | ***Ndufa9*** | 876.8 | 933.6 | 650.6 | 910.3 | 478.2 | 1225.4 |
| Q80XN0 | ***Bdh1*** | 715.5 | 849.1 | 923.7 | 659.3 | 450.4 | 905.3 |
| Q91V61 | ***Sfxn3*** | 1334 | 1377.6 | 1147.6 | 1094.3 | 740.6 | 1277.5 |
| P35564 | ***Canx*** | 642.5 | 344.7 | 203.5 | 622.3 | 1048.3 | 574 |
| P99024 | ***Tubb5*** | 183.9 | 289.2 | 115 | 442.4 | 545.1 | 339.9 |
| Q9D0M3 | ***Cyc1*** | 4423 | 4952 | 4056.8 | 2885.7 | 2193.8 | 3152.4 |
| P63011 | ***Rab3a*** | 350.6 | 605.8 | 404.3 | 661.8 | 621.4 | 692.2 |
| P56394 | ***Cox17*** | 148.2 | 110.3 | 193.9 | 195.4 | 179.1 | 177.2 |
| P22315 | ***Fech*** | 547 | 641.7 | 560.4 | 495.3 | 368.9 | 559.6 |
| P56391 | ***Cox6b1*** | 3574.8 | 2944.4 | 3363.7 | 3145.1 | 1825.7 | 3230.2 |
| O35643 | ***Ap1b1*** | 180.3 | 241.1 | 154.1 | 333.5 | 356.2 | 328.1 |
| P05202 | ***Got2*** | 3846.8 | 3744.6 | 5569.8 | 6101.4 | 4581.4 | 7163.4 |
| P84228 | ***Hist1h3b; Hist1h3c; Hist1h3d; Hist1h3e; Hist1h3f; Hist2h3b; Hist2h3c1; Hist2h3c2*** | 461.2 | 845.5 | 1621.1 | 677.4 | 248.2 | 1266.3 |
| Q8BH95 | ***Echs1*** | 1808.4 | 2124.5 | 2148 | 1693.8 | 1618 | 2030.3 |
| P52503 | ***Ndufs6*** | 3149.2 | 2601.9 | 3448.6 | 1772.5 | 1394.9 | 1917.9 |
| O88696 | ***Clpp*** | 114.7 | 108.8 | 91.1 | 80.5 | 80.6 | 95.2 |
| P35486 | ***Pdha1*** | 4597.2 | 4923.6 | 4489.7 | 3347.1 | 2828.5 | 4238 |
| UPSP:K2CA_HUMAN | | 136.1 | 53.2 | 313.2 | 261.6 | 106.6 | 182.9 |
| Q9CQZ6 | ***Ndufb3*** | 852.4 | 1088.7 | 1294.9 | 858.7 | 642.4 | 916 |
| P11588 | ***Mup1*** | 66.6 | 85.8 | 116.8 | 76.9 | 109.1 | 90.9 |
| Q60759 | ***Gcdh*** | 191.2 | 239.5 | 135 | 176.9 | 180.9 | 235.3 |
| Q9CQX8 | ***Mrps36*** | 1798.3 | 1916.4 | 2440 | 1194.8 | 1011 | 1272.7 |
| Q80U23 | ***Snph*** | 144.9 | 193.1 | 86.3 | 186.9 | 214.6 | 192.1 |
| Q9Z2I9 | ***Sucla2*** | 4257.9 | 5427 | 2696.4 | 3937.2 | 2885 | 3945.7 |

| | | | | | | | |
|---|---|---|---|---|---|---|---|
| Q91VR2 | ***Atp5c1*** | 5102.4 | 5270 | 5221.6 | 3890 | 3638.4 | 6087.1 |
| P10854 | ***Hist1h2bm*** | 42.3 | 181.9 | 411.7 | 175.8 | 29.2 | 432.9 |
| Q9DB15 | ***Mrpl12*** | 845.3 | 1008.4 | 883.9 | 718.2 | 608.5 | 879 |
| P50518 | ***Atp6v1e1*** | 871.3 | 719.6 | 1274.2 | 978.1 | 1204.5 | 730.3 |
| O70325 | ***Gpx4*** | 236.6 | 287.8 | 220.3 | 222.4 | 220.7 | 247.9 |
| Q9CYH2 | ***Fam213a*** | 152.1 | 203.8 | 122 | 158.5 | 207.4 | 191.1 |
| P03930 | ***Mtatp8*** | 1096.3 | 1379.6 | 1324.2 | 908.1 | 527.1 | 1157.9 |
| P46460 | ***Nsf*** | 821.9 | 829.3 | 559 | 2190 | 2652.6 | 966.6 |
| Q01853 | ***Vcp*** | 105.5 | 130 | 164.8 | 199.2 | 357.3 | 208.2 |
| O55125 | ***Nipsnap1*** | 440.9 | 578.2 | 813.2 | 501.5 | 441.8 | 665.4 |
| P51881 | ***Slc25a5*** | 3247.4 | 3945.9 | 2834 | 3386.3 | 2323.1 | 4440.3 |
| Q8BMS1 | ***Hadha*** | 1154.9 | 2140.4 | 1029.7 | 1352.4 | 1255.3 | 1960.6 |
| P20108 | ***Prdx3*** | 635.8 | 654.2 | 628.1 | 861.9 | 677.4 | 902 |
| Q78IK4 | ***Apool*** | 246.5 | 299.4 | 235.3 | 157.8 | 156 | 250.7 |
| Q9D1G1 | ***Rab1b*** | 224.8 | 330.4 | 204.2 | 317.2 | 401.3 | 326.4 |
| Q925I1 | ***Atad3*** | 188 | 272.5 | 180.9 | 202.3 | 201.1 | 297.8 |
| P62077 | ***Timm8b*** | 236 | 246.5 | 242 | 147.8 | 153.4 | 184 |
| P58281 | ***Opa1*** | 1744.8 | 2600.2 | 1387 | 2200.3 | 1756.5 | 2902.9 |
| Q91V41 | ***Rab14*** | 168.5 | 109.1 | 118.8 | 208.6 | 239.4 | 237.5 |
| O35658 | ***C1qbp*** | 192.5 | 148 | 172.2 | 123.1 | 105.5 | 127.5 |
| Q9QY76 | ***Vapb*** | 282 | 245 | 177.4 | 329.2 | 557.4 | 260.3 |
| Q9CQ54 | ***Ndufc2*** | 778 | 1382.6 | 817.1 | 869.5 | 736.4 | 1196.3 |
| Q99LX0 | ***Park7*** | 232.6 | 193.7 | 325.4 | 257.9 | 425.3 | 250.4 |
| Q63880 | ***Ces3a*** | 26.6 | 17.5 | 17.2 | 37.3 | 36.4 | 36.3 |
| P51660 | ***Hsd17b4*** | 27.4 | 44.2 | 42.1 | 37.9 | 55.6 | 43.9 |
| P97742 | ***Cpt1a*** | 64.6 | 94.4 | 55.8 | 64.1 | 76.2 | 84.7 |
| P06880 | ***Gh1*** | 20 | 16.5 | 106.9 | 445.7 | 212.6 | 80.8 |
| Q9WUR9 | ***Ak4*** | 707.8 | 769.6 | 1120.5 | 607.1 | 577.2 | 778.1 |
| P11352 | ***Gpx1*** | 109.2 | 132.6 | 163.6 | 133.8 | 114.8 | 190.9 |
| Q9DB60 | ***Fam213b*** | 80 | 118.5 | 88.1 | 74.7 | 86.4 | 104.2 |
| P56382 | ***Atp5e*** | 822.4 | 553.6 | 1146.3 | 468.9 | 677 | 355.8 |
| P58252 | ***Eef2*** | 30.9 | 43.7 | 35.9 | 57.4 | 53.8 | 44 |
| Q8C196 | ***Cps1*** | 411.2 | 282.3 | 354.6 | 278.7 | 297.8 | 146.6 |
| Q7TMM9 | ***Tubb2a*** | 1568.5 | 3064 | 1049.1 | 4465.2 | 5823.1 | 3103.9 |
| Q78J03 | ***Msrb2*** | 156 | 140.1 | 186.7 | 137.7 | 182 | 158.3 |
| Q8R1S0 | ***Coq6*** | 157.5 | 204.9 | 194.4 | 164.2 | 122.6 | 198.3 |
| Q921I1 | ***Tf*** | 199.1 | 106.6 | 153.1 | 197.8 | 359.3 | 177.5 |
| P30275 | ***Ckmt1*** | 5594.5 | 4833 | 8739.4 | 5599.4 | 4371.3 | 6418.4 |
| Q9QZ23 | ***Nfu1*** | 160.3 | 135 | 173.2 | 215 | 140.1 | 205.8 |
| O08585 | ***Clta*** | 390.5 | 377.3 | 670.1 | 721.1 | 944.6 | 629 |
| P57759 | ***Erp29*** | 162.9 | 104.4 | 159.3 | 135 | 229.3 | 128.8 |
| P18872 | ***Gnao1*** | 1067.1 | 1006.1 | 529.8 | 955 | 1925.2 | 768.6 |
| P14094 | ***Atp1b1*** | 2214 | 2329.5 | 1135 | 3103.2 | 4085.9 | 2144 |
| P21279 | ***Gnaq*** | 75.2 | 66.6 | 59.2 | 97.8 | 114.2 | 59.6 |
| Q8VDN2 | ***Atp1a1*** | 467.7 | 466.6 | 325.8 | 575.8 | 1127.8 | 460.5 |
| O09111 | ***Ndufb11*** | 1510.6 | 1551.3 | 1295.1 | 1133.2 | 829 | 1225.3 |
| Q60634 | ***Flot2*** | 105.2 | 52.9 | 68.6 | 196.7 | 102.6 | 74.4 |

| | | | | | | | |
|---|---|---|---|---|---|---|---|
| P50544 | ***Acadvl*** | 307.6 | 482 | 205.7 | 301.7 | 269.5 | 406.8 |
| Q9CXT8 | ***Pmpcb*** | 931.6 | 1023.2 | 749.1 | 795.4 | 592 | 908.6 |
| P68372 | ***Tubb4b*** | 328.8 | 328 | 177.6 | 478.9 | 552.3 | 341.1 |
| P62874 | ***Gnb1*** | 191.9 | 226 | 93.4 | 229 | 433.3 | 138.2 |
| Q6PB66 | ***Lrpprc*** | 847 | 1137 | 530.4 | 922.1 | 594.7 | 1180.5 |
| Q921G7 | ***Etfdh*** | 1263.3 | 1431.3 | 1040.6 | 827.1 | 752.4 | 1166.5 |
| Q9Z1P6 | ***Ndufa7*** | 1686.1 | 2256.2 | 2336 | 1292.9 | 1236.1 | 1506.1 |
| Q8R3Q6 | ***Ccdc58*** | 60.8 | 59.8 | 86.3 | 52.7 | 37.5 | 48.9 |
| Q64521 | ***Gpd2*** | 3329.8 | 4236.7 | 2711.1 | 3091.5 | 2552.7 | 4126.2 |
| O88935 | ***Syn1*** | 752.7 | 1376.6 | 1576.1 | 1966.1 | 2465.5 | 1720.2 |
| P01837 | | 7435.6 | 3016.6 | 3032.5 | 2984.3 | 3206.9 | 1427.3 |
| Q9R0P5 | ***Dstn*** | 139 | 242.5 | 337.3 | 312 | 457.8 | 216.8 |
| O88441 | ***Mtx2*** | 160.1 | 244.2 | 99.4 | 219.6 | 180.9 | 290.8 |
| P04943 | | 2322.5 | 1565.7 | 1541.4 | 1554.3 | 1339.3 | 863.6 |
| Q60597 | ***Ogdh*** | 5975.5 | 7207.2 | 3505 | 5442.2 | 3760.9 | 5936.2 |
| P62897 | ***Cycs*** | 4236 | 3812.3 | 6005.7 | 1992 | 1769.2 | 1966 |
| P01831 | ***Thy1*** | 277.5 | 303 | 113 | 161.6 | 484.8 | 123.8 |
| Q9DCT2 | ***Ndufs3*** | 2295.1 | 2461.7 | 1749.2 | 1945.4 | 1646.4 | 2453.2 |
| Q9D855 | ***Uqcrb*** | 2299.6 | 2498.4 | 2839.6 | 1738.6 | 1584.7 | 1872.3 |
| P70296 | ***Pebp1*** | 110.7 | 148.9 | 189.5 | 290.7 | 625.1 | 214.2 |
| Q9DCW4 | ***Etfb*** | 1923.3 | 1991.8 | 2428.9 | 1633.6 | 1509.3 | 1756.1 |
| Q9DBG3 | ***Ap2b1*** | 102.8 | 131.2 | 98.7 | 204.5 | 204.2 | 189.9 |
| Q9ERD7 | ***Tubb3*** | 70.6 | 188.1 | 74.7 | 243.1 | 385.7 | 195.3 |
| Q9D6F9 | ***Tubb4a*** | 77.2 | 150.8 | 63.5 | 195.4 | 293.7 | 157.5 |
| Q8BW75 | ***Maob*** | 788.9 | 999.3 | 370.9 | 721.7 | 624.7 | 943.7 |
| Q8C1B7 | ***42989*** | 290.1 | 404.4 | 204.5 | 429.4 | 828.6 | 417.3 |
| Q9CZL5 | ***Pcbd2*** | 148.1 | 110.9 | 115.4 | 72.7 | 105.1 | 79.8 |
| Q9WUR2 | ***Eci2*** | 210.7 | 277.5 | 219.9 | 244.4 | 238.2 | 308.8 |
| P60710 | ***Actb*** | 351.8 | 453.6 | 219.9 | 515 | 763.5 | 473.6 |
| P21107 | ***Tpm3*** | 127 | 97.7 | 196.6 | 87.2 | 217.7 | 119.4 |
| P04370 | ***Mbp*** | 628.3 | 1584.3 | 903.4 | 1367.8 | 4935.9 | 1022.8 |
| Q9DBJ1 | ***Pgam1*** | 94.5 | 110.3 | 108.3 | 212.9 | 496.1 | 146.6 |
| P31786 | ***Dbi*** | 249.1 | 252.4 | 537.4 | 474.3 | 831.9 | 372.4 |
| P97450 | ***Atp5j*** | 6506.2 | 5410.1 | 8416.3 | 3874.7 | 2957.1 | 3849.8 |
| P08551 | ***Nefl*** | 367.7 | 506.6 | 551.6 | 639.7 | 695.5 | 438.5 |
| Q9CPQ8 | ***Atp5l*** | 419.2 | 648.1 | 766.4 | 599.5 | 337.4 | 835.2 |
| O08539 | ***Bin1*** | 136.6 | 244.4 | 225.3 | 250.1 | 535.1 | 190.5 |
| Q91YQ5 | ***Rpn1*** | 152.4 | 55.2 | 47.1 | 142.5 | 229.3 | 115.7 |
| P62814 | ***Atp6v1b2*** | 980.8 | 819.5 | 759.1 | 1269 | 1410.6 | 1063.9 |
| Q04447 | ***Ckb*** | 785.8 | 1209.4 | 1200.4 | 2124.9 | 3518.2 | 1825.7 |
| P61264 | ***Stx1b*** | 628.6 | 628 | 504.9 | 799 | 1112 | 719.5 |
| O70439 | ***Stx7*** | 379.1 | 83.8 | 289 | 455.6 | 219.6 | 233.3 |
| P07309 | ***Ttr*** | 172.9 | 75.8 | 388.2 | 246 | 349.1 | 137.5 |
| P60202 | ***Plp1*** | 208.8 | 650.1 | 265.6 | 684 | 2310.1 | 452.2 |
| Q91YT0 | ***Ndufv1*** | 2740.7 | 3452.6 | 2533.5 | 2436.4 | 1660 | 3203.5 |
| P02088 | ***Hbb-b1*** | 599.8 | 1051.2 | 3951.1 | 1118.2 | 543.1 | 2624.2 |
| P17183 | ***Eno2*** | 447.6 | 680.8 | 578.3 | 1230.8 | 2687.7 | 925.6 |

| | | | | | | | |
|---|---|---|---|---|---|---|---|
| Q02819 | ***Nucb1*** | 85.3 | 16.8 | 76.2 | 61.5 | 253.4 | 38.1 |
| Q9QYG0 | ***Ndrg2*** | 102.3 | 144.9 | 94 | 321.9 | 385.2 | 212.8 |
| Q9ES97 | ***Rtn3*** | 168.3 | 182.7 | 156.5 | 258.2 | 413.1 | 214.1 |
| O55042 | ***Snca*** | 163.9 | 316.2 | 379.3 | 457.1 | 809.4 | 409.4 |
| Q9CQV8 | ***Ywhab*** | 609 | 733.9 | 655.2 | 1616.2 | 2815.8 | 1028.2 |
| P18760 | ***Cfl1*** | 273.2 | 441.9 | 403.7 | 537.3 | 891.4 | 338.5 |
| Q61885 | ***Mog*** | 111.4 | 235.1 | 177.6 | 178.1 | 533.2 | 137.2 |
| Q9D6J6 | ***Ndufv2*** | 2461.4 | 2569 | 2894.3 | 1837.6 | 1476.9 | 2249.4 |
| Q6PIE5 | ***Atp1a2*** | 1258.1 | 1758.9 | 1238.8 | 2097.9 | 3083.3 | 2057.1 |
| Q8BMK4 | ***Ckap4*** | 91.3 | 28.3 | 38.1 | 99.5 | 177.5 | 69.2 |
| P14152 | ***Mdh1*** | 271.4 | 413.3 | 394.1 | 896.4 | 1547.7 | 617.7 |
| Q9CR98 | ***Fam136a*** | 686.8 | 379.8 | 954.7 | 418 | 403.8 | 408.1 |
| Q8BHN3 | ***Ganab*** | 86.5 | 41.7 | 26.2 | 94.3 | 141.9 | 75.5 |
| Q60930 | ***Vdac2*** | 2898 | 3662.3 | 1932.4 | 4161.2 | 2814.3 | 3909.2 |
| P63328 | ***Ppp3ca*** | 50.1 | 51 | 31.6 | 49.4 | 136.8 | 32 |
| P08553 | ***Nefm*** | 143.9 | 270.8 | 244.5 | 376.1 | 346 | 237 |
| Q8CI94 | ***Pygb*** | 58.1 | 88.1 | 58.6 | 91.6 | 82.5 | 110.6 |
| P50396 | ***Gdi1*** | 66.7 | 107 | 46.6 | 178.8 | 442.2 | 145.6 |
| Q64458 | ***Cyp2c29*** | 16.6 | 7.4 | 19 | 12.1 | 25.4 | 11.4 |
| Q9CQ69 | ***Uqcrq*** | 893.4 | 1373.9 | 1109.5 | 1126 | 834.9 | 1262.2 |
| Q9DCU9 | ***Hoga1*** | 17.4 | 20 | 13.8 | 10.6 | 18.1 | 13.8 |
| Q9ESW4 | ***Agk*** | 128 | 180.5 | 99.4 | 147.3 | 114.2 | 197.8 |
| O08553 | ***Dpysl2*** | 1189.9 | 1567.4 | 1071.9 | 1699.6 | 3211.3 | 1150.2 |
| Q9DCJ5 | ***Ndufa8*** | 2768.2 | 2998.5 | 3185 | 2154.8 | 1702.3 | 2543.9 |
| Q9CXW2 | ***Mrps22*** | 415 | 502.9 | 302.3 | 378.7 | 328.1 | 443.4 |
| P62821 | ***Rab1A*** | 166.1 | 181.2 | 124.7 | 198.2 | 296.4 | 174.9 |
| Q61171 | ***Prdx2*** | 336.1 | 394.7 | 707.5 | 600.8 | 783.8 | 696 |
| Q6IRU5 | ***Cltb*** | 270.7 | 256.3 | 427.4 | 447.4 | 687 | 358.9 |
| P05064 | ***Aldoa*** | 1193.1 | 1747.8 | 1220.6 | 2058.5 | 5454.6 | 1775.3 |
| Q9Z2Q6 | ***42983*** | 981.6 | 1207.6 | 694.9 | 1295.1 | 2658.5 | 1276.9 |
| P09411 | ***Pgk1*** | 346.7 | 464.1 | 516.4 | 931 | 1826.3 | 680.2 |
| P62073 | ***Timm10*** | 577.4 | 433.2 | 671.5 | 290.8 | 287.4 | 272.8 |
| Q7TMF3 | ***Ndufa12*** | 1194.2 | 1570 | 1456.1 | 1095.8 | 775.5 | 1206.2 |
| O08599 | ***Stxbp1*** | 982.4 | 1464.7 | 718.2 | 1303.8 | 2170 | 1150.6 |
| Q60931 | ***Vdac3*** | 1551.8 | 2014.8 | 1099.3 | 1971.7 | 1147.4 | 2006.6 |
| O55022 | ***Pgrmc1*** | 139.2 | 110.4 | 92.6 | 157.9 | 314.2 | 136.8 |
| P24369 | ***Ppib*** | 161.5 | 91.5 | 112 | 170.6 | 283.2 | 121.5 |
| P00920 | ***Ca2*** | 120.3 | 210.6 | 198.7 | 292.1 | 718.3 | 264.1 |
| Q05816 | ***Fabp5*** | 63 | 69.4 | 101.6 | 105.6 | 227.1 | 98.5 |
| Q91VM9 | ***Ppa2*** | 636.2 | 696.6 | 610.5 | 601.5 | 532.7 | 659.9 |
| Q99LY9 | ***Ndufs5*** | 799.7 | 1222.8 | 1161.5 | 1033.4 | 754.8 | 1215.9 |
| P97872 | ***Fmo5*** | 15.1 | 15.6 | 9.4 | 20.9 | 24.2 | 14.3 |
| Q99JY0 | ***Hadhb*** | 1806.6 | 2600 | 1940.6 | 1601.4 | 1746.9 | 2112.6 |
| Q9CZU6 | ***Cs*** | 3312.4 | 2466.5 | 3196.5 | 4206.7 | 2195 | 4663 |
| Q6GQS1 | ***Slc25a23*** | 195 | 221.5 | 149.3 | 167.7 | 136.1 | 244.1 |
| P32020 | ***Scp2*** | 172.7 | 206.5 | 473.2 | 317.1 | 377.3 | 255.4 |
| Q9CPU0 | ***Glo1*** | 62.8 | 65.5 | 101 | 128.3 | 226.4 | 125.1 |

| | | | | | | | |
|---|---|---|---|---|---|---|---|
| Q8BMS4 | ***Coq3*** | 185.6 | 175.8 | 167.5 | 167.9 | 129.2 | 193.1 |
| P51150 | ***Rab7a*** | 168.1 | 114.4 | 140.6 | 207.1 | 211 | 150.4 |
| Q8BWF0 | ***Aldh5a1*** | 3479.8 | 3371.4 | 2478.1 | 2197.1 | 1867.9 | 2429.9 |
| Q9CZ42 | ***Naxd*** | 278.9 | 404 | 517.2 | 336.7 | 284.9 | 401.3 |
| Q99PL5 | ***Rrbp1*** | 180.1 | 104 | 146.3 | 182.6 | 362.7 | 150.3 |
| Q9R0K7 | ***Atp2b2*** | 131.3 | 175 | 97.8 | 208.8 | 369 | 206.2 |
| P01942 | ***Hba*** | 325.7 | 777.6 | 2770.4 | 697.6 | 318.7 | 1561.7 |
| O55131 | ***42985*** | 300.2 | 291.1 | 212.2 | 316 | 754.4 | 303.2 |
| Q9D7P6 | ***Iscu*** | 166.4 | 179.6 | 242.9 | 149.2 | 130.1 | 157.1 |
| P43006 | ***Slc1a2*** | 253.6 | 261.2 | 160.3 | 342.6 | 748.8 | 277.3 |
| P16858 | ***Gapdh*** | 1013.6 | 1536.5 | 1398 | 2024.9 | 2966.4 | 1587.8 |
| Q9CR68 | ***Uqcrfs1*** | 5553.9 | 6883.1 | 7296.8 | 4472.5 | 3556.3 | 4992.6 |
| Q9D3D9 | ***Atp5d*** | 1022 | 738.4 | 1140.2 | 632.3 | 462.7 | 669.9 |
| P35700 | ***Prdx1*** | 677.8 | 770.9 | 769.3 | 1080.5 | 1109.9 | 1121.1 |
| Q9CR61 | ***Ndufb7*** | 874.2 | 1160.2 | 1084.9 | 879.3 | 737.4 | 983.5 |
| P68134 | ***Acta1*** | 2984 | 3648.2 | 2524.3 | 4745.7 | 6361.8 | 4486.3 |
| Q99LC3 | ***Ndufa10*** | 2880.2 | 3638.8 | 2446.1 | 2816.3 | 2205.7 | 3201.6 |
| Q9DB77 | ***Uqcrc2*** | 6763.7 | 8654.1 | 6759.1 | 5850.9 | 4780.7 | 7130.8 |
| O55126 | ***Gbas*** | 1100.3 | 1711 | 1726.7 | 1421.9 | 1043.8 | 1498.8 |
| Q7TQF7 | ***Amph*** | 473.4 | 837.9 | 639.7 | 737.2 | 1318.7 | 693.7 |
| Q9QXV0 | ***Pcsk1n*** | 325.3 | 208.7 | 327.2 | 292.5 | 377.1 | 244 |
| Q62261 | ***Sptbn1*** | 162 | 328.8 | 143.3 | 244.5 | 546.7 | 284 |
| Q3UJU9 | ***Rmdn3*** | 84.4 | 101.9 | 32.8 | 80 | 100.7 | 104.6 |
| Q8R1I1 | ***Uqcr10*** | 547.3 | 596.5 | 1172.7 | 767.7 | 378.3 | 719.9 |
| Q61644 | ***Pacsin1*** | 767.8 | 1104.2 | 818.6 | 872.5 | 1851 | 892.8 |
| P10649 | ***Gstm1*** | 235 | 313.1 | 300.6 | 574.8 | 846.5 | 448.2 |
| Q8JZN5 | ***Acad9*** | 312.2 | 354.7 | 161.9 | 351.6 | 195.6 | 453.3 |
| Q9ER00 | ***Stx12*** | 140.3 | 43.6 | 104.7 | 177.2 | 128.6 | 80 |
| Q80WJ7 | ***Mtdh*** | 121.5 | 28.7 | 49.1 | 91.2 | 226.8 | 86.9 |
| Q8R086 | ***Suox*** | 57 | 60.4 | 61.1 | 76.6 | 103.1 | 98.2 |
| Q9DCS9 | ***Ndufb10*** | 1895.3 | 2174.1 | 2235.8 | 1687.2 | 1340.3 | 1902.9 |
| P10637 | ***Mapt*** | 504.5 | 583 | 547.7 | 729.8 | 1619.7 | 443.9 |
| Q9Z1S5 | ***42981*** | 411 | 578.3 | 335.8 | 544.9 | 1096.1 | 534.7 |
| P12787 | ***Cox5a*** | 4965.2 | 4813.3 | 5557.3 | 3724.5 | 2870.3 | 4231.4 |
| P68373 | ***Tuba1c*** | 1821.5 | 3079.2 | 929.1 | 4115.7 | 6646.1 | 3512.9 |
| P43024 | ***Cox6a1*** | 247.5 | 178.4 | 205.3 | 186 | 197.7 | 250.8 |
| P62880 | ***Gnb2*** | 721.9 | 991.3 | 361 | 928.3 | 1995.5 | 679.5 |
| Q8BP92 | ***Rcn2*** | 330.9 | 201.3 | 416 | 306 | 537.8 | 282 |
| O35887 | ***Calu*** | 275.8 | 91.2 | 215.2 | 126.3 | 324.2 | 86.6 |
| Q8BFP9 | ***Pdk1*** | 198.2 | 269.2 | 141.9 | 239.5 | 179.1 | 297 |
| Q9CXI5 | ***Manf*** | 110.8 | 55.6 | 89.8 | 129.7 | 274.9 | 68.3 |
| P31650 | ***Slc6a11*** | 695.9 | 888.2 | 563.7 | 1040.7 | 1806.9 | 969 |
| Q9D6R2 | ***Idh3a*** | 6650.6 | 7059.8 | 5707.4 | 5917.3 | 5137.8 | 6538.7 |
| Q7TQD2 | ***Tppp*** | 523.6 | 554.5 | 659.8 | 571.9 | 1294.9 | 337.9 |
| P52480 | ***Pkm*** | 959.4 | 1220.5 | 1202.9 | 2323.9 | 3652.4 | 1764.3 |
| Q68FD5 | ***Cltc*** | 238.3 | 264.5 | 276.4 | 623.2 | 660.8 | 655.2 |
| P53810 | ***Pitpna*** | 99.1 | 109.6 | 125.2 | 133.5 | 193.3 | 118.4 |

| | | | | | | | |
|---|---|---|---|---|---|---|---|
| Q62420 | ***Sh3gl2*** | 294 | 560.7 | 552 | 461.4 | 759.9 | 443.4 |
| P17182 | ***Eno1*** | 1882.5 | 2151.1 | 2002.3 | 3505.4 | 6444.9 | 2424.5 |
| P62259 | ***Ywhae*** | 454.8 | 610.8 | 556.4 | 1048.1 | 2213.3 | 882.3 |
| Q9QXX4 | ***Slc25a13*** | 173.3 | 145 | 115.1 | 165.7 | 118.8 | 194.2 |
| P46660 | ***Ina*** | 196.1 | 244.2 | 253.2 | 311.2 | 346.6 | 181.4 |
| Q3UIU2 | ***Ndufb6*** | 118.5 | 117.4 | 110.4 | 133 | 124.8 | 199.2 |
| P50136 | ***Bckdha*** | 94.4 | 114.9 | 76.1 | 74.2 | 82.1 | 83.1 |
| P20357 | ***Map2*** | 123.5 | 144.7 | 82.4 | 158.5 | 221.9 | 152.1 |
| P00158 | ***Mt-Cyb*** | 456.7 | 328.9 | 321.4 | 372.7 | 335.5 | 436.2 |
| P48771 | ***Cox7a2*** | 1180.7 | 1429.3 | 1412.3 | 1258.8 | 770.8 | 1235.1 |
| P50171 | ***Hsd17b8*** | 86.5 | 112.4 | 122.3 | 93.5 | 82.2 | 133.8 |
| Q3ULD5 | ***Mccc2*** | 197.4 | 231 | 145.5 | 162.3 | 135.8 | 220.9 |
| Q59J78 | ***Ndufaf2*** | 468.6 | 584.6 | 659.7 | 350.4 | 313.6 | 367.5 |
| Q99LB7 | ***Sardh*** | 128.3 | 137.6 | 168.7 | 198 | 196.7 | 194.3 |
| P61982 | ***Ywhag*** | 307.3 | 285.5 | 287.3 | 479.1 | 1150.9 | 434.4 |
| P17710 | ***Hk1*** | 5897.9 | 6019.6 | 4093.7 | 5619 | 3363.5 | 7514.7 |
| Q9QYA2 | ***Tomm40*** | 188.8 | 222.7 | 184.5 | 206.1 | 196.9 | 334.5 |
| P14211 | ***Calr*** | 2783.3 | 869.9 | 1701.8 | 1877.5 | 4231.5 | 1273.9 |
| Q9CPP6 | ***Ndufa5*** | 2283.6 | 2720.7 | 2921.2 | 2090.8 | 1626.4 | 2536.5 |
| Q8R429 | ***Atp2a1*** | 65.5 | 56.7 | 44 | 93.4 | 213.7 | 85.1 |
| P17742 | ***Ppia*** | 506.1 | 463 | 958 | 863.1 | 1759.2 | 745.5 |
| Q8K0T0 | ***Rtn1*** | 222.4 | 202.5 | 173 | 385.9 | 800.1 | 290.6 |
| Q9Z0X1 | ***Aifm1*** | 956.7 | 1229.6 | 813.6 | 983.3 | 673.5 | 1116 |
| P09103 | ***P4hb*** | 1651.5 | 693.8 | 1193.8 | 1130.6 | 2559.9 | 854.4 |
| Q99JB2 | ***Stoml2*** | 474.8 | 591.8 | 586.1 | 424.2 | 406 | 564.7 |
| P70404 | ***Idh3g*** | 1616.7 | 2407.6 | 1836.1 | 2201.8 | 1979.1 | 2542.3 |
| P46096 | ***Syt1*** | 316.9 | 417.8 | 148.4 | 458.2 | 712.7 | 407.2 |
| P05201 | ***Got1*** | 54 | 74.1 | 71.9 | 155.4 | 281.5 | 119.8 |
| P11798 | ***Camk2a*** | 439.1 | 470.4 | 166.9 | 834.3 | 1548.1 | 549.6 |
| O08709 | ***Prdx6*** | 107.5 | 143.8 | 184.2 | 326.1 | 461.8 | 255.8 |
| P63044 | ***Vamp2*** | 254.5 | 268.5 | 249.1 | 451.6 | 340.3 | 361.4 |
| Q64516 | ***Gk*** | 419.5 | 520.7 | 261.5 | 491.7 | 309.5 | 647.3 |
| G5E829 | ***Atp2b1*** | 121.4 | 157 | 88.3 | 219.7 | 412.7 | 174.2 |
| P00405 | ***Mtco2*** | 1400.9 | 1431.7 | 951.4 | 1919.9 | 948.2 | 2279.9 |
| P17426 | ***Ap2a1*** | 121.3 | 194.3 | 103 | 276 | 269.8 | 286.7 |
| Q9D7A8 | ***Armc1*** | 100.1 | 143.9 | 153.7 | 148.6 | 137.2 | 159.2 |
| P16015 | ***Ca3*** | 13.3 | 5.9 | 9.6 | 14.9 | 16.9 | 16.4 |
| Q9D273 | ***Mmab*** | 261.2 | 341.9 | 455.9 | 200.4 | 203.7 | 283.4 |
| Q9R1Q8 | ***Tagln3*** | 95.3 | 108.9 | 182.4 | 195.4 | 346.4 | 139.3 |
| P10126 | ***Eef1a1*** | 604.7 | 821.9 | 795.9 | 1238 | 1506.1 | 999.2 |
| Q99PJ0 | ***Ntm*** | 203.1 | 248.7 | 103.1 | 209.6 | 470 | 179.4 |
| P47934 | ***Crat*** | 167.7 | 226.4 | 119 | 149.1 | 108.7 | 202.8 |
| O08917 | ***Flot1*** | 57.5 | 23.4 | 38.7 | 74.3 | 48.6 | 42.8 |
| O88451 | ***Rdh7*** | 14.5 | 8.5 | 12.9 | 21.4 | 29.6 | 14.9 |
| P55302 | ***Lrpap1*** | 167.4 | 79.2 | 122.9 | 169.6 | 332.8 | 124 |
| P54869 | ***Hmgcs2*** | 50.5 | 40.5 | 28.6 | 42.6 | 53.3 | 25.5 |
| P16546 | ***Sptan1*** | 753.1 | 2029.4 | 546.7 | 1444.4 | 3370.8 | 1555.8 |

| | | | | | | | |
|---|---|---|---|---|---|---|---|
| P16460 | ***Ass1*** | 66.4 | 93.6 | 65.3 | 106.2 | 146.8 | 68.7 |
| Q2TPA8 | ***Hsdl2*** | 108.9 | 160.7 | 153.1 | 106.7 | 129.8 | 149 |
| Q99MR8 | ***Mccc1*** | 204.4 | 318.5 | 144.8 | 257.5 | 210.8 | 277.1 |
| P70441 | ***Slc9a3r1*** | 92.2 | 85.8 | 96.7 | 112.3 | 195.5 | 63.2 |
| Q4VAE3 | ***Tmem65*** | 64.8 | 111.1 | 103.3 | 148.4 | 81.9 | 133.5 |
| Q9JK42 | ***Pdk2*** | 132.6 | 175.1 | 91.4 | 153.6 | 114.3 | 180.8 |
| Q3UHB1 | ***Nt5dc3*** | 382.4 | 493.4 | 339.7 | 541.7 | 352.2 | 717.6 |
| P27773 | ***Pdia3*** | 3176.2 | 1237.4 | 2540.2 | 2227.3 | 5321.6 | 1730.3 |
| O35490 | ***Bhmt*** | 25.8 | 26.3 | 36.1 | 30.4 | 47.9 | 32.8 |
| Q9ERS2 | ***Ndufa13*** | 1291.1 | 1567.9 | 1304.3 | 1258.1 | 759.5 | 1299.1 |
| Q9D6U8 | ***Fam162a*** | 148 | 294.1 | 256.8 | 249.5 | 178.5 | 300.2 |
| Q91VT4 | ***Cbr4*** | 81.6 | 123.1 | 111.1 | 102.6 | 83.9 | 151.9 |
| Q9WV98 | ***Timm9*** | 1840.3 | 1629.9 | 1836.5 | 1207.6 | 1150.8 | 1348.3 |
| P24270 | ***Cat*** | 103.9 | 84.4 | 145.7 | 136.2 | 195.6 | 102.4 |
| P14231 | ***Atp1b2*** | 225.4 | 371.9 | 192.5 | 375.2 | 604.2 | 334.6 |
| P63101 | ***Ywhaz*** | 241.8 | 328.9 | 264.4 | 501.1 | 1229.2 | 407.5 |
| P26645 | ***Marcks*** | 342.5 | 468 | 518.9 | 485 | 811.5 | 449.5 |
| Q925N0 | ***Sfxn5*** | 1046 | 1433.1 | 774.4 | 878.9 | 817.7 | 1407.3 |
| P08003 | ***Pdia4*** | 719.2 | 319.8 | 493.6 | 616.6 | 1231.8 | 505.5 |
| P54227 | ***Stmn1*** | 180.9 | 117.8 | 229.6 | 121.8 | 471.1 | 144.5 |
| Q8K3J1 | ***Ndufs8*** | 948 | 1084.7 | 1101.4 | 832.3 | 615.5 | 942.9 |
| Q9CQN1 | ***Trap1*** | 413.9 | 551.4 | 268.1 | 454.3 | 373.2 | 564.9 |
| Q9D1Q6 | ***Erp44*** | 222.7 | 80.8 | 153.2 | 210.7 | 496.2 | 156.3 |
| P07901 | ***Hsp90aa1*** | 155.1 | 152.3 | 106.8 | 273.3 | 435.9 | 206.6 |
| Q9Z1G4 | ***Atp6v0a1*** | 316 | 298.7 | 194.5 | 473.4 | 494.8 | 515.6 |
| P47857 | ***Pfkm*** | 106.5 | 213.8 | 119 | 269.4 | 352.2 | 263.1 |
| P10639 | ***Txn*** | 136.1 | 103.6 | 201.7 | 151.8 | 233.2 | 139.6 |
| Q9EP89 | ***Lactb*** | 124 | 158.2 | 81.6 | 136.4 | 75.8 | 162 |
| P01868 | ***Ighg1*** | 9512.2 | 5631.5 | 4621.7 | 5057.7 | 5860.3 | 3003.8 |
| P84091 | ***Ap2m1*** | 116.9 | 149.6 | 98.8 | 178.5 | 263 | 190.9 |
| Q05920 | ***Pc*** | 2399 | 3099.8 | 1286.6 | 2137.3 | 1952.1 | 2690.7 |
| O08749 | ***Dld*** | 4732.7 | 6381.6 | 4122.3 | 5386.9 | 4629.6 | 6282.7 |
| Q60864 | ***Stip1*** | 73 | 87.6 | 104.9 | 96.8 | 173.3 | 94.7 |
| Q8BFZ9 | ***Erlin2*** | 151.8 | 101.1 | 62.7 | 146.1 | 280.4 | 133.1 |
| P99027 | ***Rplp2*** | 179.5 | 111.2 | 225.4 | 257 | 361.1 | 134 |
| Q8R366 | ***Igsf8*** | 80 | 71.6 | 44.9 | 89.2 | 226.3 | 44.2 |
| Q64133 | ***Maoa*** | 355.6 | 359.3 | 201.9 | 274.9 | 251.6 | 383.6 |
| P38060 | ***Hmgcl*** | 391.5 | 369.5 | 302.6 | 300.7 | 273.9 | 324.5 |
| P13595 | ***Ncam1*** | 420.9 | 532.1 | 181.7 | 488.2 | 1067.1 | 398.1 |
| P68254 | ***Ywhaq*** | 161.4 | 181 | 141.9 | 250.9 | 507.8 | 218.6 |
| Q9WUM5 | ***Suclg1*** | 3629.1 | 5048.5 | 4668.3 | 4314 | 3694.4 | 4243 |
| Q9CPY7 | ***Lap3*** | 246.5 | 324.2 | 230.4 | 275.9 | 322.7 | 324.8 |
| P47911 | ***Rpl6*** | 213.4 | 97.2 | 132.1 | 252.2 | 306.1 | 195.4 |
| Q91VD9 | ***Ndufs1*** | 8113.8 | 9037.2 | 5018.3 | 7215.2 | 5062.4 | 7964.8 |
| Q9DCS3 | ***Mecr*** | 297.4 | 307.2 | 308.7 | 271.7 | 234.3 | 311.5 |
| Q7TSJ2 | ***Map6*** | 160.5 | 159.4 | 95.6 | 257.2 | 451.8 | 134.8 |
| Q6PIC6 | ***Atp1a3*** | 5204.3 | 6125.2 | 3686.8 | 7763.5 | 12489 | 7115.8 |

| | | | | | | | |
|---|---|---|---|---|---|---|---|
| Q91WS0 | ***Cisd1*** | 2075 | 2530.8 | 2959 | 2204.7 | 2182.6 | 2513.9 |
| P52825 | ***Cpt2*** | 200.6 | 324.5 | 178.8 | 179.5 | 179.5 | 229.5 |
| Q99KB8 | ***Hagh*** | 166.8 | 170.4 | 217.8 | 149.8 | 185.7 | 145.8 |
| P52760 | ***Hrsp12*** | 51.7 | 56.3 | 92.5 | 64.9 | 127.4 | 89.5 |
| P17665 | ***Cox7c*** | 1898.9 | 1990.3 | 1962.6 | 1180.4 | 926.9 | 1478.1 |
| P19536 | ***Cox5b*** | 3686.6 | 3058.4 | 3812.9 | 2241.8 | 1829.7 | 2494.7 |
| P51863 | ***Atp6v0d1*** | 314 | 288.5 | 200.8 | 431.4 | 370.5 | 414.6 |
| Q91WC3 | ***Acsl6*** | 237.5 | 380 | 167.7 | 234.8 | 302.2 | 409.2 |
| P16330 | ***Cnp*** | 1091.4 | 2239.3 | 972.9 | 1994.6 | 6094.2 | 1763.3 |
| Q91XE0 | ***Glyat*** | 28.7 | 12.5 | 21.6 | 20.6 | 20 | 9 |
| Q8BJ64 | ***Chdh*** | 58.9 | 58 | 28.5 | 35 | 42.7 | 49.7 |
| P63001 | ***Rac1*** | 99.9 | 119.4 | 89.6 | 176.9 | 193 | 119.3 |
| Q9CRB9 | ***Chchd3*** | 1979.5 | 2200.1 | 1906 | 1585.9 | 1433.1 | 2106.6 |
| O55143 | ***Atp2a2*** | 123.8 | 86.7 | 66.9 | 153.1 | 298.6 | 127.4 |
| P10852 | ***Slc3a2*** | 74.8 | 80.7 | 57.5 | 114.3 | 200.5 | 93.6 |
| Q01768 | ***Nme2*** | 141.3 | 173.4 | 389.7 | 367 | 489.1 | 375.4 |
| Q8BK30 | ***Ndufv3*** | 628.1 | 423.7 | 520.6 | 301.4 | 243.9 | 297.9 |
| Q8JZU2 | ***Slc25a1*** | 100.7 | 117.4 | 77.8 | 94 | 88.6 | 138.6 |
| Q91ZA3 | ***Pcca*** | 1684.8 | 1977.7 | 966.4 | 1657.3 | 1453.1 | 1942.1 |
| Q8K1Z0 | ***Coq9*** | 377.7 | 356.7 | 489.3 | 289.3 | 195.9 | 294.5 |
| P12960 | ***Cntn1*** | 258.2 | 211.1 | 98.4 | 209.6 | 425.9 | 155.4 |
| Q8VDQ8 | ***Sirt2*** | 68.2 | 137.9 | 74.2 | 74.8 | 374.4 | 66.2 |
| P16125 | ***Ldhb*** | 207 | 331 | 240.3 | 593 | 717.5 | 510.1 |
| O70251 | ***Eef1b*** | 131.7 | 63.9 | 122.6 | 110.1 | 156.7 | 69.5 |
| P53395 | ***Dbt*** | 581.9 | 796.7 | 473.1 | 449.8 | 513.4 | 632.3 |
| P97427 | ***Crmp1*** | 482.9 | 745.3 | 482.9 | 751.7 | 1468 | 567.6 |
| P60879 | ***Snap25*** | 607.9 | 859.1 | 740.3 | 974.2 | 1787.7 | 760.7 |
| P11499 | ***Hsp90ab1*** | 414.3 | 494.6 | 299.9 | 688.9 | 1024.9 | 551.7 |
| Q99MN9 | ***Pccb*** | 696.9 | 839.7 | 553.5 | 731.8 | 570.3 | 861.8 |
| Q8VCT4 | ***Ces1d*** | 14.5 | 2.9 | 19.8 | 12.9 | 17.7 | 8.7 |
| P40630 | ***Tfam*** | 497.2 | 524.5 | 500.5 | 461.7 | 428.7 | 572.1 |
| Q61548 | ***Snap91*** | 247.9 | 360.9 | 228 | 478.4 | 745.3 | 471.4 |
| Q99K67 | ***Aass*** | 25.8 | 25.2 | 24.5 | 20.6 | 19.3 | 20.2 |
| P62204 | ***Calm1; Calm2; Calm3*** | 633.7 | 694.2 | 944.2 | 789.2 | 1817.9 | 603.3 |
| P39053 | ***Dnm1*** | 653.9 | 1278.1 | 656.9 | 1261.6 | 1686.3 | 1117.3 |
| Q9DCV4 | ***Rmdn1*** | 59.1 | 92.7 | 66.3 | 59.9 | 75.8 | 78.9 |
| Q9Z1J3 | ***Nfs1*** | 378.2 | 564.1 | 365.5 | 450.3 | 359.9 | 561.4 |
| P68510 | ***Ywhah*** | 150.1 | 168.7 | 145.7 | 266.8 | 589.3 | 236.3 |
| Q9CXZ1 | ***Ndufs4*** | 1840.4 | 1806.4 | 1967.9 | 1420.7 | 1069.6 | 1504.9 |
| P47754 | ***Capza2*** | 105.6 | 113 | 86 | 175.4 | 232.3 | 149 |
| Q8VE33 | ***Gdap1l1*** | 192.8 | 163.9 | 168.1 | 196 | 147.1 | 239.2 |
| P08113 | ***Hsp90b1*** | 2391.2 | 774.9 | 997.5 | 1676.7 | 3732.4 | 1140.1 |
| Q9Z0P4 | ***Palm*** | 71.6 | 106.4 | 57.7 | 96.8 | 243 | 90.6 |
| Q922Q1 | ***42796*** | 480.4 | 615.8 | 356.3 | 445.9 | 397 | 551.7 |
| Q9JKR6 | ***Hyou1*** | 670.3 | 339 | 470.4 | 550.6 | 1129.5 | 401.8 |
| P63017 | ***Hspa8*** | 2451.5 | 2499.3 | 2685.7 | 4456.6 | 6592.2 | 3746.6 |

| | | | | | | | |
|---|---|---|---|---|---|---|---|
| P17751 | ***Tpi1*** | 699.5 | 706.4 | 830.4 | 955.1 | 2244.3 | 781.3 |
| Q9D7B6 | ***Acad8*** | 118 | 154.5 | 85.3 | 98.1 | 109.3 | 134.6 |
| Q8BGH2 | ***Samm50*** | 415.3 | 585.7 | 342.6 | 645.9 | 338.9 | 680.9 |
| Q9WU79 | ***Prodh*** | 90.2 | 105.2 | 67 | 80 | 60.3 | 100.9 |
| P99028 | ***Uqcrh*** | 1523.4 | 1315.6 | 1960.1 | 1045.1 | 934.6 | 1201.6 |
| Q91XV3 | ***Basp1*** | 3145.1 | 3781.2 | 2166.6 | 2817.2 | 8701.4 | 2166.2 |
| Q9CQX2 | ***Cyb5b*** | 464.6 | 461.4 | 321.1 | 406.4 | 445.1 | 523.8 |
| O08795 | ***Prkcsh*** | 265.7 | 101.2 | 207.1 | 162.5 | 401 | 144.8 |
| P06837 | ***Gap43*** | 951.8 | 829.7 | 471.7 | 735 | 2402.2 | 650.3 |
| Q60598 | ***Cttn*** | 34.7 | 47.8 | 61.2 | 46.3 | 57.3 | 52.7 |
| P05063 | ***Aldoc*** | 340.7 | 370.1 | 351.4 | 656.8 | 1088.5 | 667.2 |
| Q62277 | ***Syp*** | 363.7 | 430.9 | 196.6 | 463.2 | 425.6 | 489.6 |
| Q3TC72 | ***Fahd2*** | 825.4 | 1002.4 | 641.8 | 772.2 | 637.9 | 838.2 |
| P25688 | ***Uox*** | 7.5 | 7.6 | 20.6 | 13.8 | 18.9 | 6.2 |
| P99029 | ***Prdx5*** | 3933.5 | 3307.8 | 4819.7 | 3277.2 | 2463.1 | 3227.5 |
| P06151 | ***Ldha*** | 49.4 | 94.9 | 37.4 | 116.5 | 172.6 | 107.2 |
| P06745 | ***Gpi*** | 50.8 | 40.2 | 50.8 | 98.2 | 159.7 | 81.9 |
| Q9WV55 | ***Vapa*** | 148 | 142.7 | 98.8 | 213.8 | 399.6 | 192.2 |
| Q922R8 | ***Pdia6*** | 353.6 | 140.5 | 237.6 | 272.7 | 538.8 | 193.6 |
| P20029 | ***Hspa5*** | 4737.9 | 1866.4 | 3188.1 | 4153.2 | 9026.3 | 3150.6 |
